## Supplemental Table 1 for "Splicing factor SRSF1 is essential for homing of precursor spermatogonial stem cells in mice"

**Table S1. SRSF1 peak-containing genes identified through CLIP-seq.**

| GeneID | geneSymbol | GeneID | geneSymbol |
|---|---|---|---|
| ENSMUSG00000000028 | *Cdc45* | ENSMUSG00000037487 | *Ubr5* |
| ENSMUSG00000000037 | *Scml2* | ENSMUSG00000037519 | *Ppfia1* |
| ENSMUSG00000000125 | *Wnt3* | ENSMUSG00000037531 | *Mrpl47* |
| ENSMUSG00000000127 | *Fer* | ENSMUSG00000037533 | *Rapgef6* |
| ENSMUSG00000000131 | *Xpo6* | ENSMUSG00000037536 | *Fbxo34* |
| ENSMUSG00000000282 | *Mnt* | ENSMUSG00000037608 | *Bclaf1* |
| ENSMUSG00000000339 | *Rtca* | ENSMUSG00000037617 | *Spag1* |
| ENSMUSG00000000365 | *Rnf17* | ENSMUSG00000037622 | *Wdtc1* |
| ENSMUSG00000000399 | *Ndufa9* | ENSMUSG00000037624 | *Kcnk2* |
| ENSMUSG00000000411 | *Tssk3* | ENSMUSG00000037627 | *Rgs22* |
| ENSMUSG00000000420 | *Galnt1* | ENSMUSG00000037628 | *Cdkn3* |
| ENSMUSG00000000563 | *Atp5pb* | ENSMUSG00000037683 | *Armc3* |
| ENSMUSG00000000568 | *Hnrnpd* | ENSMUSG00000037689 | *Tmem247* |
| ENSMUSG00000000605 | *Clcn4* | ENSMUSG00000037697 | *Ddhd1* |
| ENSMUSG00000000631 | *Myo18a* | ENSMUSG00000037716 | *Ccdc33* |
| ENSMUSG00000000738 | *Spg7* | ENSMUSG00000037736 | *Limch1* |
| ENSMUSG00000000740 | *Rpl13* | ENSMUSG00000037742 | *Eef1a1* |
| ENSMUSG00000000787 | *Ddx3x* | ENSMUSG00000037787 | *Coa8* |
| ENSMUSG00000000794 | *Kcnn3* | ENSMUSG00000037801 | *Iqch* |
| ENSMUSG00000000804 | *Usp32* | ENSMUSG00000037808 | *Fam76b* |
| ENSMUSG00000000811 | *Txnrd3* | ENSMUSG00000037852 | *Cpe* |
| ENSMUSG00000000823 | *Zfp512b* | ENSMUSG00000037857 | *Nufip2* |
| ENSMUSG00000000838 | *Fmr1* | ENSMUSG00000037876 | *Jmjd1c* |
| ENSMUSG00000000915 | *Hip1r* | ENSMUSG00000037894 | *H2az1* |
| ENSMUSG00000001016 | *Ilf2* | ENSMUSG00000037926 | *Ssh2* |
| ENSMUSG00000001017 | *Chtop* | ENSMUSG00000037933 | *Bicd2* |
| ENSMUSG00000001052 | *Sec24b* | ENSMUSG00000037935 | *Smarce1* |
| ENSMUSG00000001089 | *Luzp1* | ENSMUSG00000037958 | *Nsrp1* |
| ENSMUSG00000001105 | *Ift20* | ENSMUSG00000037971 | *1110032A03Rik* |
| ENSMUSG00000001120 | *Pcbp3* | ENSMUSG00000037979 | *Ccdc92* |
| ENSMUSG00000001151 | *Pcnt* | ENSMUSG00000037993 | *Dhx38* |
| ENSMUSG00000001156 | *Mxd1* | ENSMUSG00000037999 | *Arap2* |
| ENSMUSG00000001158 | *Snrnp27* | ENSMUSG00000038002 | *Cramp1* |
| ENSMUSG00000001175 | *Calm1* | ENSMUSG00000038010 | *Ccdc138* |
| ENSMUSG00000001211 | *Agpat3* | ENSMUSG00000038011 | *Dnah10* |
| ENSMUSG00000001228 | *Uhrf1* | ENSMUSG00000038015 | *Prm2* |
| ENSMUSG00000001247 | *Lsr* | ENSMUSG00000038022 | *Mindy4* |
| ENSMUSG00000001248 | *Gramd1a* | ENSMUSG00000038025 | *Phf2* |
| ENSMUSG00000001289 | *Pfdn5* | ENSMUSG00000038026 | *Kcnj9* |
| ENSMUSG00000001305 | *Rrp15* | ENSMUSG00000038039 | *Gcc2* |
| ENSMUSG00000001366 | *Fbxo9* | ENSMUSG00000038042 | *Ptpdc1* |
| ENSMUSG00000001383 | *Zmat2* | ENSMUSG00000038056 | *Kmt2c* |
| ENSMUSG00000001415 | *Smg5* | ENSMUSG00000038057 | *Dbil5* |
| ENSMUSG00000001416 | *Cct3* | ENSMUSG00000038069 | *Cdkn2aip* |
| ENSMUSG00000001435 | *Col18a1* | ENSMUSG00000038070 | *Cntln* |
| ENSMUSG00000001440 | *Kpnb1* | ENSMUSG00000038084 | *Opa1* |
| ENSMUSG00000001467 | *Cyp51* | ENSMUSG00000038085 | *Cnbd2* |
| ENSMUSG00000001472 | *Tcf25* | ENSMUSG00000038095 | *Sbno1* |
| ENSMUSG00000001525 | *Tubb5* | ENSMUSG00000038116 | *Phf20* |
| ENSMUSG00000001542 | *Ell2* | ENSMUSG00000038126 | *Mphosph9* |
| ENSMUSG00000001558 | *Klhl10* | ENSMUSG00000038127 | *Ccdc50* |
| ENSMUSG00000001569 | *Nom1* | ENSMUSG00000038128 | *Camk4* |
| ENSMUSG00000001632 | *Brpf1* | ENSMUSG00000038143 | *Stox2* |
| ENSMUSG00000001666 | *Ddt* | ENSMUSG00000038165 | *1700022A21Rik* |
| ENSMUSG00000001674 | *Ddx18* | ENSMUSG00000038170 | *Pde4dip* |
| ENSMUSG00000001687 | *Ubl3* | ENSMUSG00000038180 | *Spag4* |

| | | | |
|---|---|---|---|
| ENSMUSG00000001767 | *Crnkl1* | ENSMUSG00000038187 | *Btbd10* |
| ENSMUSG00000001783 | *Rtcb* | ENSMUSG00000038199 | *Iqca1l* |
| ENSMUSG00000001794 | *Capns1* | ENSMUSG00000038240 | *Pdss2* |
| ENSMUSG00000001829 | *Clpb* | ENSMUSG00000038241 | *Cep250* |
| ENSMUSG00000001833 | *Septin7* | ENSMUSG00000038244 | *Mical2* |
| ENSMUSG00000001910 | *Nacc1* | ENSMUSG00000038246 | *Fam50b* |
| ENSMUSG00000001948 | *Spa17* | ENSMUSG00000038252 | *Ncapd2* |
| ENSMUSG00000002017 | *Fam98a* | ENSMUSG00000038271 | *Iffo1* |
| ENSMUSG00000002031 | *Ift46* | ENSMUSG00000038274 | *Fau* |
| ENSMUSG00000002052 | *Supt6* | ENSMUSG00000038279 | *Nop2* |
| ENSMUSG00000002055 | *Spag5* | ENSMUSG00000038290 | *Smg6* |
| ENSMUSG00000002076 | *Hsf2bp* | ENSMUSG00000093815 | None |
| ENSMUSG00000002102 | *Psmc3* | ENSMUSG00000038292 | *Ccdc155* |
| ENSMUSG00000002107 | *Celf2* | ENSMUSG00000038298 | *Pdzk1* |
| ENSMUSG00000002129 | *Sf3a1* | ENSMUSG00000038335 | *Tsr1* |
| ENSMUSG00000002190 | *Clgn* | ENSMUSG00000038344 | *Txlng* |
| ENSMUSG00000002210 | *Smg9* | ENSMUSG00000038351 | *Sgsm2* |
| ENSMUSG00000002265 | *Peg3* | ENSMUSG00000038366 | *Lasp1* |
| ENSMUSG00000002307 | *Daxx* | ENSMUSG00000038369 | *Ncoa6* |
| ENSMUSG00000002308 | *Cd320* | ENSMUSG00000038371 | *Sbf2* |
| ENSMUSG00000002319 | *Ipo4* | ENSMUSG00000038374 | *Rbm8a* |
| ENSMUSG00000002324 | *Rec8* | ENSMUSG00000038384 | *Setd1b* |
| ENSMUSG00000002372 | *Ranbp3* | ENSMUSG00000038388 | *Pals2* |
| ENSMUSG00000002409 | *Dyrk1b* | ENSMUSG00000038398 | *Upf3a* |
| ENSMUSG00000002413 | *Braf* | ENSMUSG00000038406 | *Scaf1* |
| ENSMUSG00000002428 | *Hltf* | ENSMUSG00000038408 | *1700018A04Rik* |
| ENSMUSG00000002455 | *Prpf6* | ENSMUSG00000038437 | *Mllt6* |
| ENSMUSG00000002486 | *Tchp* | ENSMUSG00000038453 | *Srcin1* |
| ENSMUSG00000002489 | *Tiam1* | ENSMUSG00000038467 | *Chmp4b* |
| ENSMUSG00000002524 | *Puf60* | ENSMUSG00000038481 | *Cdk19* |
| ENSMUSG00000002546 | *Golga2* | ENSMUSG00000038495 | *Otud7b* |
| ENSMUSG00000002608 | *Ccdc97* | ENSMUSG00000038503 | *Mesd* |
| ENSMUSG00000002625 | *Akap8l* | ENSMUSG00000038510 | *Rpf2* |
| ENSMUSG00000002658 | *Gtf2f1* | ENSMUSG00000038518 | *Jarid2* |
| ENSMUSG00000002732 | *Fkbp7* | ENSMUSG00000038523 | *1700003F12Rik* |
| ENSMUSG00000002748 | *Baz1b* | ENSMUSG00000038535 | *Zfp280d* |
| ENSMUSG00000002768 | *Mea1* | ENSMUSG00000038538 | *Ubn2* |
| ENSMUSG00000002833 | *Hdgfl2* | ENSMUSG00000038546 | *Ranbp9* |
| ENSMUSG00000002835 | *Chaf1a* | ENSMUSG00000038564 | *Ift172* |
| ENSMUSG00000002948 | *Map2k7* | ENSMUSG00000038587 | *Akap12* |
| ENSMUSG00000002949 | *Timm44* | ENSMUSG00000038594 | *Cep85l* |
| ENSMUSG00000002985 | *Apoe* | ENSMUSG00000038611 | *Phrf1* |
| ENSMUSG00000002996 | *Hbp1* | ENSMUSG00000038615 | *Nfe2l1* |
| ENSMUSG00000003033 | *Ap1m1* | ENSMUSG00000038619 | *Ensa* |
| ENSMUSG00000003037 | *Rab8a* | ENSMUSG00000038637 | *Lrrc56* |
| ENSMUSG00000003039 | *Fam32a* | ENSMUSG00000038664 | *Herc1* |
| ENSMUSG00000003068 | *Stk11* | ENSMUSG00000038668 | *Lpar1* |
| ENSMUSG00000003119 | *Cdk12* | ENSMUSG00000038683 | *Pak1ip1* |
| ENSMUSG00000003123 | *Lipe* | ENSMUSG00000038690 | *Atp5j2* |
| ENSMUSG00000003134 | *Tbc1d8* | ENSMUSG00000038691 | *Mbd3l1* |
| ENSMUSG00000003153 | *Slc2a3* | ENSMUSG00000038696 | *Mapkap1* |
| ENSMUSG00000003184 | *Irf3* | ENSMUSG00000038708 | *Golga4* |
| ENSMUSG00000003200 | *Sh3gl1* | ENSMUSG00000038722 | *bud31* |
| ENSMUSG00000003226 | *Ranbp2* | ENSMUSG00000038729 | *Pakap* |
| ENSMUSG00000003234 | *Abcf3* | ENSMUSG00000038733 | *Wdr26* |
| ENSMUSG00000003235 | *Eif2b5* | ENSMUSG00000038759 | *Nup205* |
| ENSMUSG00000003269 | *Cyth2* | ENSMUSG00000038762 | *Abcf1* |
| ENSMUSG00000003316 | *Glg1* | ENSMUSG00000038766 | *Gabpb2* |

| | | | |
|---|---|---|---|
| ENSMUSG00000003341 | *Atp8b3* | ENSMUSG00000038774 | *Ascc3* |
| ENSMUSG00000003345 | *Csnk1g2* | ENSMUSG00000038776 | *Ephx1* |
| ENSMUSG00000003354 | *Ccdc65* | ENSMUSG00000038784 | *Cnot4* |
| ENSMUSG00000003360 | *Ddx23* | ENSMUSG00000038797 | *Zscan2* |
| ENSMUSG00000003402 | *Prkcsh* | ENSMUSG00000038806 | *Sde2* |
| ENSMUSG00000003421 | *Nosip* | ENSMUSG00000038836 | *Agbl3* |
| ENSMUSG00000003423 | *Pih1d1* | ENSMUSG00000038844 | *Kif16b* |
| ENSMUSG00000003429 | *Rps11* | ENSMUSG00000038866 | *Zcchc2* |
| ENSMUSG00000003435 | *Supt5* | ENSMUSG00000038871 | *Bpgm* |
| ENSMUSG00000003437 | *Paf1* | ENSMUSG00000038895 | *Zfp653* |
| ENSMUSG00000003527 | *Ess2* | ENSMUSG00000038900 | *Rpl12* |
| ENSMUSG00000003531 | *Dgcr6* | ENSMUSG00000038909 | *Kat7* |
| ENSMUSG00000003604 | *Aven* | ENSMUSG00000038914 | *Dido1* |
| ENSMUSG00000003660 | *Snrnp200* | ENSMUSG00000038925 | *E330034G19Rik* |
| ENSMUSG00000003731 | *Kpna6* | ENSMUSG00000038932 | *Tcfl5* |
| ENSMUSG00000003778 | *Brd8* | ENSMUSG00000038943 | *Prc1* |
| ENSMUSG00000003779 | *Kif20a* | ENSMUSG00000038967 | *Pdk2* |
| ENSMUSG00000003810 | *Mast2* | ENSMUSG00000038973 | *Cldnd2* |
| ENSMUSG00000003814 | *Calr* | ENSMUSG00000038976 | *Ppp1r9b* |
| ENSMUSG00000003824 | *Syce2* | ENSMUSG00000038987 | *Cfap157* |
| ENSMUSG00000003847 | *Nfat5* | ENSMUSG00000038994 | *H1f9* |
| ENSMUSG00000003923 | *Tfam* | ENSMUSG00000038997 | *Asb17* |
| ENSMUSG00000003970 | *Rpl8* | ENSMUSG00000039007 | *Cpq* |
| ENSMUSG00000004032 | *Gstm5* | ENSMUSG00000039031 | *Arhgap18* |
| ENSMUSG00000004040 | *Stat3* | ENSMUSG00000039033 | *Tasp1* |
| ENSMUSG00000004070 | *Hmox2* | ENSMUSG00000039047 | *Pigk* |
| ENSMUSG00000004096 | *Cwc15* | ENSMUSG00000039050 | *Osbpl2* |
| ENSMUSG00000004099 | *Dnmt1* | ENSMUSG00000039067 | *Psmd7* |
| ENSMUSG00000004127 | *Trmt10a* | ENSMUSG00000039068 | *Zzz3* |
| ENSMUSG00000004207 | *Psap* | ENSMUSG00000039099 | *Wdr93* |
| ENSMUSG00000004347 | *Pde1c* | ENSMUSG00000039100 | *Marchf6* |
| ENSMUSG00000004364 | *Cul3* | ENSMUSG00000039103 | *Nexn* |
| ENSMUSG00000004455 | *Ppp1cc* | ENSMUSG00000039105 | *Atp6v1g1* |
| ENSMUSG00000004460 | *Dnajb11* | ENSMUSG00000039110 | *Mycbpap* |
| ENSMUSG00000004508 | *Gab2* | ENSMUSG00000039148 | *Sart1* |
| ENSMUSG00000004535 | *Tax1bp1* | ENSMUSG00000039156 | *Stim2* |
| ENSMUSG00000004591 | *Pkn2* | ENSMUSG00000039159 | *Ube2h* |
| ENSMUSG00000004610 | *Etfb* | ENSMUSG00000039166 | *Akap7* |
| ENSMUSG00000004637 | *Wwox* | ENSMUSG00000039168 | *Dap* |
| ENSMUSG00000004642 | *Slbp* | ENSMUSG00000039176 | *Polg* |
| ENSMUSG00000004667 | *Polr2e* | ENSMUSG00000039179 | *Tekt5* |
| ENSMUSG00000004771 | *Rab11a* | ENSMUSG00000039187 | *Fanci* |
| ENSMUSG00000004798 | *Ulk2* | ENSMUSG00000039205 | *Ciz1* |
| ENSMUSG00000004843 | *Chmp2b* | ENSMUSG00000039210 | *Gpatch2* |
| ENSMUSG00000004865 | *Srpk1* | ENSMUSG00000039218 | *Srrm2* |
| ENSMUSG00000004895 | *Prcc* | ENSMUSG00000039219 | *Arid4b* |
| ENSMUSG00000004897 | *Hdgf* | ENSMUSG00000039220 | *Ppp1r10* |
| ENSMUSG00000004929 | *Thop1* | ENSMUSG00000039224 | *D1Pas1* |
| ENSMUSG00000004980 | *Hnrnpa2b1* | ENSMUSG00000039254 | *Pomt1* |
| ENSMUSG00000005045 | *Chd5* | ENSMUSG00000039262 | *Prrc2b* |
| ENSMUSG00000005102 | *Eif2ak4* | ENSMUSG00000039286 | *Fndc3b* |
| ENSMUSG00000005103 | *Wdr1* | ENSMUSG00000039298 | *Cdk5rap2* |
| ENSMUSG00000005131 | *4930550C14Rik* | ENSMUSG00000039330 | *Tsga10ip* |
| ENSMUSG00000005198 | *Polr2a* | ENSMUSG00000039335 | *Spata16* |
| ENSMUSG00000005237 | *Dnah2* | ENSMUSG00000039391 | *Ccdc81* |
| ENSMUSG00000005299 | *Letm1* | ENSMUSG00000039396 | *Neil3* |
| ENSMUSG00000005362 | *Crbn* | ENSMUSG00000039449 | *Prpf18* |
| ENSMUSG00000005370 | *Msh6* | ENSMUSG00000039456 | *Morc3* |

| | | | |
|---|---|---|---|
| ENSMUSG00000005371 | *Fbxo11* | ENSMUSG00000039473 | *Ubn1* |
| ENSMUSG00000005374 | *Tbl2* | ENSMUSG00000039477 | *Tnrc18* |
| ENSMUSG00000005417 | *Mprip* | ENSMUSG00000039492 | *Ccdc27* |
| ENSMUSG00000005474 | *Myl10* | ENSMUSG00000039501 | *Znfx1* |
| ENSMUSG00000005481 | *Ddx39a* | ENSMUSG00000039523 | *Cep104* |
| ENSMUSG00000005483 | *Dnajb1* | ENSMUSG00000039530 | *Tusc3* |
| ENSMUSG00000005566 | *Trim28* | ENSMUSG00000039531 | *Zup1* |
| ENSMUSG00000005575 | *Ube2m* | ENSMUSG00000039536 | *Stau1* |
| ENSMUSG00000005609 | *Ctr9* | ENSMUSG00000039540 | *4921524L21Rik* |
| ENSMUSG00000005610 | *Eif4g2* | ENSMUSG00000039543 | *Cfap70* |
| ENSMUSG00000005615 | *Pcyt1a* | ENSMUSG00000039555 | *Cylc2* |
| ENSMUSG00000005625 | *Psmd4* | ENSMUSG00000039585 | *Myo9a* |
| ENSMUSG00000005687 | *Bcas2* | ENSMUSG00000039615 | *Stub1* |
| ENSMUSG00000005698 | *Ctcf* | ENSMUSG00000039629 | *Strip2* |
| ENSMUSG00000005732 | *Ranbp1* | ENSMUSG00000039630 | *Hnrnpu* |
| ENSMUSG00000005803 | *Sqor* | ENSMUSG00000039632 | *Odad3* |
| ENSMUSG00000005823 | *Gpr108* | ENSMUSG00000039660 | *Spout1* |
| ENSMUSG00000005881 | *Ergic3* | ENSMUSG00000039684 | *Gm5422* |
| ENSMUSG00000005882 | *Uqcc1* | ENSMUSG00000039703 | *Nploc4* |
| ENSMUSG00000005886 | *Ncoa2* | ENSMUSG00000039704 | *Lmbrd2* |
| ENSMUSG00000005936 | *Kctd20* | ENSMUSG00000039716 | *Dock3* |
| ENSMUSG00000005981 | *Trap1* | ENSMUSG00000039735 | *Fnbp1l* |
| ENSMUSG00000005983 | *1700037C18Rik* | ENSMUSG00000039737 | *Prkrip1* |
| ENSMUSG00000006005 | *Tpr* | ENSMUSG00000039738 | *Slx4* |
| ENSMUSG00000006024 | *Napa* | ENSMUSG00000039742 | *Garin1b* |
| ENSMUSG00000006058 | *Snf8* | ENSMUSG00000039756 | *Dnttip2* |
| ENSMUSG00000006169 | *Clint1* | ENSMUSG00000039763 | *Dnajc28* |
| ENSMUSG00000006191 | *Cdkal1* | ENSMUSG00000039781 | *Cep131* |
| ENSMUSG00000006215 | *Zbtb17* | ENSMUSG00000039782 | *Cpeb2* |
| ENSMUSG00000006276 | *Eps15l1* | ENSMUSG00000039801 | *Cplane1* |
| ENSMUSG00000006299 | *Aamp* | ENSMUSG00000039804 | *Ncoa5* |
| ENSMUSG00000006307 | *Kmt2b* | ENSMUSG00000039828 | *Wdr70* |
| ENSMUSG00000006310 | *Zbtb32* | ENSMUSG00000039831 | *Arhgap29* |
| ENSMUSG00000006333 | *Rps9* | ENSMUSG00000039852 | *Rere* |
| ENSMUSG00000006335 | *Tfpt* | ENSMUSG00000039879 | *Heca* |
| ENSMUSG00000006527 | *Sfmbt1* | ENSMUSG00000039891 | *Txlnb* |
| ENSMUSG00000006576 | *Slc4a3* | ENSMUSG00000039943 | *Plcb4* |
| ENSMUSG00000006599 | *Gtf2h1* | ENSMUSG00000039953 | *Clstn1* |
| ENSMUSG00000006676 | *Usp19* | ENSMUSG00000039958 | *Etfbkmt* |
| ENSMUSG00000006715 | *Gmnn* | ENSMUSG00000039959 | *Hip1* |
| ENSMUSG00000006740 | *Kif5b* | ENSMUSG00000039963 | *Ccdc40* |
| ENSMUSG00000006784 | *Odad4* | ENSMUSG00000039977 | *Deup1* |
| ENSMUSG00000006906 | *Stambp* | ENSMUSG00000039990 | *Edrf1* |
| ENSMUSG00000006930 | *Hap1* | ENSMUSG00000040003 | *Magi2* |
| ENSMUSG00000006932 | *Ctnnb1* | ENSMUSG00000040013 | *Fkbp6* |
| ENSMUSG00000006941 | *Eif1b* | ENSMUSG00000040040 | *Ift88* |
| ENSMUSG00000006998 | *Psmd2* | ENSMUSG00000040054 | *Baz2a* |
| ENSMUSG00000007029 | *Vars* | ENSMUSG00000040084 | *Bub1b* |
| ENSMUSG00000007033 | *Hspa1l* | ENSMUSG00000040097 | *Flywch1* |
| ENSMUSG00000007035 | *Msh5* | ENSMUSG00000040111 | *Gramd1b* |
| ENSMUSG00000007080 | *Pole* | ENSMUSG00000040123 | *Zmym5* |
| ENSMUSG00000007097 | *Atp1a2* | ENSMUSG00000040140 | *Tdrd6* |
| ENSMUSG00000007107 | *Atp1a4* | ENSMUSG00000040174 | *Alkbh3* |
| ENSMUSG00000007122 | *Casq1* | ENSMUSG00000040195 | *Nemp1* |
| ENSMUSG00000007440 | *Pcdha10* | ENSMUSG00000040219 | *Ttc12* |
| ENSMUSG00000007656 | *Arpp19* | ENSMUSG00000040220 | *Gas8* |
| ENSMUSG00000007721 | *Ccdc124* | ENSMUSG00000040225 | *Prrc2c* |
| ENSMUSG00000007739 | *Cct4* | ENSMUSG00000040231 | *Syngr4* |

| | | | |
|---|---|---|---|
| ENSMUSG00000007827 | *Ankrd26* | ENSMUSG00000040242 | *Fgfr1op2* |
| ENSMUSG00000007836 | *Hnrnpa0* | ENSMUSG00000040249 | *Lrp1* |
| ENSMUSG00000007850 | *Hnrnph1* | ENSMUSG00000040250 | *Ints13* |
| ENSMUSG00000007867 | *Ift43* | ENSMUSG00000040260 | *Daam2* |
| ENSMUSG00000007907 | *Cabs1* | ENSMUSG00000040263 | *Klhdc4* |
| ENSMUSG00000008028 | *1700008O03Rik* | ENSMUSG00000040265 | *Dnm3* |
| ENSMUSG00000008200 | *Fnbp4* | ENSMUSG00000040269 | *Mrps28* |
| ENSMUSG00000008301 | *Phax* | ENSMUSG00000040276 | *Pacsin1* |
| ENSMUSG00000008307 | *1700109H08Rik* | ENSMUSG00000040297 | *Suco* |
| ENSMUSG00000008333 | *Snrpb2* | ENSMUSG00000040312 | *Cchcr1* |
| ENSMUSG00000008348 | *Ubc* | ENSMUSG00000040325 | *Dcaf1* |
| ENSMUSG00000008373 | *Prpf31* | ENSMUSG00000040351 | *Ankib1* |
| ENSMUSG00000008429 | *Herpud2* | ENSMUSG00000040359 | *Ufl1* |
| ENSMUSG00000008575 | *Nfib* | ENSMUSG00000040364 | *Sec1* |
| ENSMUSG00000008683 | *Rps15a* | ENSMUSG00000040365 | *Trim41* |
| ENSMUSG00000008690 | *Ncaph2* | ENSMUSG00000040370 | *Etfrf1* |
| ENSMUSG00000008730 | *Hipk1* | ENSMUSG00000040383 | *Aqr* |
| ENSMUSG00000008763 | *Man1a2* | ENSMUSG00000040390 | *Map3k10* |
| ENSMUSG00000008822 | *Acyp1* | ENSMUSG00000040407 | *Akap9* |
| ENSMUSG00000008859 | *Rala* | ENSMUSG00000040412 | *Elapor1* |
| ENSMUSG00000008958 | *Vps72* | ENSMUSG00000040414 | *Slc25a28* |
| ENSMUSG00000008976 | *Gabpa* | ENSMUSG00000040420 | *Cdh18* |
| ENSMUSG00000009013 | *Dynll1* | ENSMUSG00000040423 | *Rc3h1* |
| ENSMUSG00000009030 | *Pdcl* | ENSMUSG00000040451 | *Sgms1* |
| ENSMUSG00000009073 | *Nf2* | ENSMUSG00000040459 | *Arglu1* |
| ENSMUSG00000009079 | *Ewsr1* | ENSMUSG00000040462 | *Os9* |
| ENSMUSG00000009093 | *Gstt4* | ENSMUSG00000040463 | *Mybbp1a* |
| ENSMUSG00000009470 | *Tnpo1* | ENSMUSG00000040473 | *Cfap69* |
| ENSMUSG00000009549 | *Srp14* | ENSMUSG00000040481 | *Bptf* |
| ENSMUSG00000009575 | *Cbx5* | ENSMUSG00000040489 | *Sox30* |
| ENSMUSG00000009596 | *Taf7l* | ENSMUSG00000040524 | *Zfp609* |
| ENSMUSG00000009628 | *Tex15* | ENSMUSG00000040541 | *Tmem225* |
| ENSMUSG00000009630 | *Ppp2cb* | ENSMUSG00000040548 | *Tex2* |
| ENSMUSG00000009670 | *Tex11* | ENSMUSG00000040549 | *Ckap5* |
| ENSMUSG00000009734 | *Pou6f2* | ENSMUSG00000040550 | *Otud6b* |
| ENSMUSG00000009741 | *Ubp1* | ENSMUSG00000040562 | *Gstm2* |
| ENSMUSG00000009907 | *Vps4b* | ENSMUSG00000040565 | *Btaf1* |
| ENSMUSG00000009927 | *Rps25* | ENSMUSG00000040591 | *Cstpp1* |
| ENSMUSG00000009941 | *Nxf2* | ENSMUSG00000040629 | *Mael* |
| ENSMUSG00000010044 | *Zmynd10* | ENSMUSG00000040640 | *Erc2* |
| ENSMUSG00000010095 | *Slc3a2* | ENSMUSG00000040649 | *Rimklb* |
| ENSMUSG00000010097 | *Nxf1* | ENSMUSG00000040651 | *Tasor* |
| ENSMUSG00000010136 | *Pifo* | ENSMUSG00000040659 | *Efhd2* |
| ENSMUSG00000010277 | *2610507B11Rik* | ENSMUSG00000040661 | *Rad54l2* |
| ENSMUSG00000010342 | *Tex14* | ENSMUSG00000040667 | *Nup88* |
| ENSMUSG00000010362 | *Rdm1* | ENSMUSG00000040690 | *Col16a1* |
| ENSMUSG00000010376 | *Nedd8* | ENSMUSG00000040720 | *Virma* |
| ENSMUSG00000010392 | *Gosr1* | ENSMUSG00000040725 | *Hnrnpul1* |
| ENSMUSG00000010406 | *Mrpl52* | ENSMUSG00000040729 | *Cep126* |
| ENSMUSG00000010435 | *Spaca7* | ENSMUSG00000040731 | *Eif4h* |
| ENSMUSG00000010453 | *Kansl3* | ENSMUSG00000040761 | *Spen* |
| ENSMUSG00000010505 | *Myt1* | ENSMUSG00000040782 | *Cop1* |
| ENSMUSG00000010608 | *Rbm25* | ENSMUSG00000040785 | *Ttc3* |
| ENSMUSG00000010755 | *Cars* | ENSMUSG00000040795 | *Iqcc* |
| ENSMUSG00000010796 | *Asz1* | ENSMUSG00000040812 | *Agbl2* |
| ENSMUSG00000011154 | *Cfap161* | ENSMUSG00000040813 | *Tex264* |
| ENSMUSG00000011158 | *Brf1* | ENSMUSG00000040824 | *Snrpd2* |
| ENSMUSG00000011306 | *Sugp1* | ENSMUSG00000040829 | *Zmynd15* |

| | | | |
|---|---|---|---|
| ENSMUSG00000011349 | *Dmrtc2* | ENSMUSG00000040838 | *Gm11639* |
| ENSMUSG00000011350 | *Gm5893* | ENSMUSG00000040850 | *Psme4* |
| ENSMUSG00000011751 | *Sptbn4* | ENSMUSG00000040852 | *Plekhh2* |
| ENSMUSG00000011831 | *Evi5* | ENSMUSG00000040859 | *Bsdc1* |
| ENSMUSG00000011832 | *Evi5l* | ENSMUSG00000040860 | *Crocc* |
| ENSMUSG00000011877 | *Git1* | ENSMUSG00000040866 | *Rsph6a* |
| ENSMUSG00000011958 | *Bnip2* | ENSMUSG00000040875 | *Osbpl10* |
| ENSMUSG00000012076 | *Brms1l* | ENSMUSG00000040888 | *Gfer* |
| ENSMUSG00000012126 | *Ubxn11* | ENSMUSG00000040919 | *4930505A04Rik* |
| ENSMUSG00000012211 | *Tex22* | ENSMUSG00000040936 | *Ulk4* |
| ENSMUSG00000012296 | *Tjap1* | ENSMUSG00000041009 | *Iqcf4* |
| ENSMUSG00000012443 | *Kif11* | ENSMUSG00000041020 | *Map7d2* |
| ENSMUSG00000012609 | *Ttll5* | ENSMUSG00000041028 | *Ghitm* |
| ENSMUSG00000013076 | *Amotl1* | ENSMUSG00000041057 | *Wdr43* |
| ENSMUSG00000013091 | *Tmem190* | ENSMUSG00000041096 | *Tspyl2* |
| ENSMUSG00000013155 | *Enkd1* | ENSMUSG00000041112 | *Elmo1* |
| ENSMUSG00000013236 | *Ptprs* | ENSMUSG00000041133 | *Smc1a* |
| ENSMUSG00000013663 | *Pten* | ENSMUSG00000041141 | *Pnma8a* |
| ENSMUSG00000013787 | *Ehmt2* | ENSMUSG00000041144 | *Dnah7b* |
| ENSMUSG00000014074 | *Rnf168* | ENSMUSG00000041161 | *Otud3* |
| ENSMUSG00000014077 | *Chp1* | ENSMUSG00000041164 | *Zmiz2* |
| ENSMUSG00000014195 | *Dnajc7* | ENSMUSG00000041168 | *Lonp1* |
| ENSMUSG00000014226 | *Cacybp* | ENSMUSG00000041180 | *Hectd2* |
| ENSMUSG00000014232 | *Cluap1* | ENSMUSG00000041203 | *Trir* |
| ENSMUSG00000014426 | *Map3k4* | ENSMUSG00000041219 | *Arhgap11a* |
| ENSMUSG00000014504 | *Srp19* | ENSMUSG00000041225 | *Arhgap12* |
| ENSMUSG00000014551 | *Mrps25* | ENSMUSG00000041229 | *Phf8* |
| ENSMUSG00000014668 | *Chfr* | ENSMUSG00000041236 | *Vps41* |
| ENSMUSG00000014782 | *Plekhg4* | ENSMUSG00000041238 | *Rbbp8* |
| ENSMUSG00000014850 | *Msh3* | ENSMUSG00000041245 | *Wnk3* |
| ENSMUSG00000014905 | *Dnajb9* | ENSMUSG00000041255 | *Tmco5b* |
| ENSMUSG00000014956 | *Ppp1cb* | ENSMUSG00000041268 | *Dmxl2* |
| ENSMUSG00000015002 | *Efr3a* | ENSMUSG00000041278 | *Ttc1* |
| ENSMUSG00000015087 | *Rabl6* | ENSMUSG00000041297 | *Cdk13* |
| ENSMUSG00000015090 | *Ptgds* | ENSMUSG00000041298 | *Katnal1* |
| ENSMUSG00000015092 | *Edf1* | ENSMUSG00000041301 | *Cftr* |
| ENSMUSG00000015095 | *Fbxw5* | ENSMUSG00000041303 | *Gtf3c3* |
| ENSMUSG00000015120 | *Ube2i* | ENSMUSG00000041323 | *Ak7* |
| ENSMUSG00000015149 | *Sirt2* | ENSMUSG00000041328 | *Pcf11* |
| ENSMUSG00000015165 | *Hnrnpl* | ENSMUSG00000041343 | *Ankrd42* |
| ENSMUSG00000015176 | *Nolc1* | ENSMUSG00000041353 | *Tmem29* |
| ENSMUSG00000015217 | *Hmgb3* | ENSMUSG00000041360 | *Pum3* |
| ENSMUSG00000015222 | *Map2* | ENSMUSG00000041361 | *Myzap* |
| ENSMUSG00000015243 | *Abca1* | ENSMUSG00000041362 | *Shtn1* |
| ENSMUSG00000015305 | *Sash1* | ENSMUSG00000041375 | *Ccdc9* |
| ENSMUSG00000015357 | *Clpx* | ENSMUSG00000041408 | *Wapl* |
| ENSMUSG00000015365 | *Mov10l1* | ENSMUSG00000041453 | *Rpl21* |
| ENSMUSG00000015597 | *Zfp318* | ENSMUSG00000041459 | *Tardbp* |
| ENSMUSG00000015599 | *Ttbk1* | ENSMUSG00000041536 | *Serpina3a* |
| ENSMUSG00000015656 | *Hspa8* | ENSMUSG00000041540 | *Sox5* |
| ENSMUSG00000015672 | *Mrpl32* | ENSMUSG00000041556 | *Fbxo2* |
| ENSMUSG00000015697 | *Setdb1* | ENSMUSG00000041560 | *Nop53* |
| ENSMUSG00000015733 | *Capza2* | ENSMUSG00000041565 | *L3mbtl4* |
| ENSMUSG00000015748 | *Prpf3* | ENSMUSG00000041566 | *Tssk1* |
| ENSMUSG00000015749 | *Anp32e* | ENSMUSG00000041570 | *Camsap2* |
| ENSMUSG00000015757 | *Ppil4* | ENSMUSG00000041596 | *Nlrp5-ps* |
| ENSMUSG00000015804 | *Med28* | ENSMUSG00000041623 | *D11Wsu47e* |
| ENSMUSG00000015837 | *Sqstm1* | ENSMUSG00000041629 | *Fam104a* |

| | | | |
|---|---|---|---|
| ENSMUSG00000015880 | *Ncapg* | ENSMUSG00000041638 | *Gcn1* |
| ENSMUSG00000015882 | *Lcorl* | ENSMUSG00000041642 | *Kif21b* |
| ENSMUSG00000015937 | *Macroh2a1* | ENSMUSG00000041645 | *Ddx24* |
| ENSMUSG00000015942 | *Gtf2ird2* | ENSMUSG00000041654 | *Slc39a11* |
| ENSMUSG00000015961 | *Adss* | ENSMUSG00000041673 | *Lrrc18* |
| ENSMUSG00000015962 | *1700016C15Rik* | ENSMUSG00000041685 | *Fcho2* |
| ENSMUSG00000015971 | *Actr8* | ENSMUSG00000041712 | *Ubr7* |
| ENSMUSG00000015980 | *Lrrc27* | ENSMUSG00000041716 | *Gm20604* |
| ENSMUSG00000016018 | *Mtrex* | ENSMUSG00000041720 | *Pi4ka* |
| ENSMUSG00000016239 | *Lonrf3* | ENSMUSG00000041733 | *Coq5* |
| ENSMUSG00000016458 | *Wt1* | ENSMUSG00000041740 | *Rnf10* |
| ENSMUSG00000016487 | *Ppfibp1* | ENSMUSG00000041757 | *Plekha6* |
| ENSMUSG00000016526 | *Dyrk3* | ENSMUSG00000041763 | *Tpp2* |
| ENSMUSG00000016554 | *Eif3d* | ENSMUSG00000041777 | *Cir1* |
| ENSMUSG00000016619 | *Nup50* | ENSMUSG00000041781 | *Cpsf2* |
| ENSMUSG00000016626 | *Nlrp14* | ENSMUSG00000041809 | *Efhc1* |
| ENSMUSG00000016664 | *Pacsin2* | ENSMUSG00000041815 | *Poldip3* |
| ENSMUSG00000016921 | *Srsf6* | ENSMUSG00000041837 | *Pdcd7* |
| ENSMUSG00000016982 | *Pom121l2* | ENSMUSG00000041841 | *Rpl37* |
| ENSMUSG00000017049 | *Ccdc70* | ENSMUSG00000041846 | *Ppp4r3a* |
| ENSMUSG00000017119 | *Nbr1* | ENSMUSG00000041852 | *Tcf20* |
| ENSMUSG00000017132 | *Cyth1* | ENSMUSG00000041859 | *Mcm3* |
| ENSMUSG00000017146 | *Brca1* | ENSMUSG00000041870 | *Ankrd13a* |
| ENSMUSG00000017264 | *Exosc10* | ENSMUSG00000041895 | *Wipi1* |
| ENSMUSG00000017291 | *Taok1* | ENSMUSG00000041912 | *Tdrkh* |
| ENSMUSG00000017299 | *Dnttip1* | ENSMUSG00000041923 | *Nol4* |
| ENSMUSG00000017376 | *Nlk* | ENSMUSG00000041935 | *Rimoc1* |
| ENSMUSG00000017404 | *Rpl19* | ENSMUSG00000041992 | *Rapgef5* |
| ENSMUSG00000017466 | *Timp2* | ENSMUSG00000042043 | *Tbca* |
| ENSMUSG00000017478 | *Zc3h18* | ENSMUSG00000042050 | *Dync2i1* |
| ENSMUSG00000017485 | *Top2b* | ENSMUSG00000042079 | *Hnrnpf* |
| ENSMUSG00000017548 | *Suz12* | ENSMUSG00000042096 | *Dao* |
| ENSMUSG00000017639 | *Rab11fip4* | ENSMUSG00000042133 | *Ppig* |
| ENSMUSG00000017707 | *Serinc3* | ENSMUSG00000042138 | *Msantd2* |
| ENSMUSG00000017720 | *Trp53tg5* | ENSMUSG00000042155 | *Klhl23* |
| ENSMUSG00000017781 | *Pitpna* | ENSMUSG00000042156 | *Dzip1* |
| ENSMUSG00000017831 | *Rab5a* | ENSMUSG00000042189 | *Tekt3* |
| ENSMUSG00000017837 | *Nkiras2* | ENSMUSG00000042197 | *Zfp451* |
| ENSMUSG00000017843 | *Ppp2r5c* | ENSMUSG00000042200 | *Cdrt4* |
| ENSMUSG00000017861 | *Mybl2* | ENSMUSG00000042207 | *Kdm5b* |
| ENSMUSG00000017978 | *Cadps2* | ENSMUSG00000042208 | *Sanbr* |
| ENSMUSG00000017999 | *Ddx27* | ENSMUSG00000042211 | *Fbxo38* |
| ENSMUSG00000018001 | *Cyth3* | ENSMUSG00000042213 | *Zfand4* |
| ENSMUSG00000018040 | *Rrp7a* | ENSMUSG00000042302 | *Ehbp1* |
| ENSMUSG00000018171 | *Vmp1* | ENSMUSG00000042305 | *Tmem183a* |
| ENSMUSG00000018189 | *Uchl5* | ENSMUSG00000042308 | *Setd1a* |
| ENSMUSG00000018196 | *Glrx2* | ENSMUSG00000042323 | *Pbrm1* |
| ENSMUSG00000018209 | *Stk4* | ENSMUSG00000042331 | *Specc1* |
| ENSMUSG00000018287 | *Spag7* | ENSMUSG00000042354 | *Gnl3* |
| ENSMUSG00000018322 | *Tomm34* | ENSMUSG00000042390 | *Gatad2b* |
| ENSMUSG00000018326 | *Ywhab* | ENSMUSG00000042423 | *Fbrs* |
| ENSMUSG00000018372 | *Cep95* | ENSMUSG00000042444 | *Mindy2* |
| ENSMUSG00000018379 | *Srsf1* | ENSMUSG00000042446 | *Zmym4* |
| ENSMUSG00000018395 | *Kif3a* | ENSMUSG00000042447 | *Mios* |
| ENSMUSG00000018412 | *Kansl1* | ENSMUSG00000042476 | *Abcb4* |
| ENSMUSG00000018415 | *Gid4* | ENSMUSG00000042487 | *Leo1* |
| ENSMUSG00000018425 | *Dhx40* | ENSMUSG00000042489 | *Clspn* |
| ENSMUSG00000018428 | *Akap1* | ENSMUSG00000042502 | *Cd2bp2* |

| | | | |
|---|---|---|---|
| ENSMUSG00000018474 | *Chd3* | ENSMUSG00000042508 | *Dmtf1* |
| ENSMUSG00000018481 | *Appbp2* | ENSMUSG00000042520 | *Ubap2l* |
| ENSMUSG00000018501 | *Ncor1* | ENSMUSG00000042525 | *4933428M09Rik* |
| ENSMUSG00000018509 | *Cenpv* | ENSMUSG00000042541 | *Sem1* |
| ENSMUSG00000018541 | *Cwc25* | ENSMUSG00000042548 | *Asxl1* |
| ENSMUSG00000018548 | *Trim37* | ENSMUSG00000042554 | *Zp3r* |
| ENSMUSG00000018554 | *Ybx2* | ENSMUSG00000042557 | *Sin3a* |
| ENSMUSG00000018567 | *Gabarap* | ENSMUSG00000042564 | *Fam227a* |
| ENSMUSG00000018570 | *2810408A11Rik* | ENSMUSG00000042599 | *Kdm7a* |
| ENSMUSG00000018572 | *Phf23* | ENSMUSG00000042606 | *Hirip3* |
| ENSMUSG00000018593 | *Sparc* | ENSMUSG00000042616 | *Oscp1* |
| ENSMUSG00000018651 | *Tada2a* | ENSMUSG00000042625 | *Safb2* |
| ENSMUSG00000018666 | *Cbx1* | ENSMUSG00000042644 | *Itpr3* |
| ENSMUSG00000018669 | *Cdk5rap3* | ENSMUSG00000042650 | *Alkbh5* |
| ENSMUSG00000018677 | *Slc25a39* | ENSMUSG00000042678 | *Myo15* |
| ENSMUSG00000018697 | *Aatf* | ENSMUSG00000042682 | *Selenok* |
| ENSMUSG00000018707 | *Dync1h1* | ENSMUSG00000042688 | *Mapk6* |
| ENSMUSG00000018736 | *Ndel1* | ENSMUSG00000042699 | *Dhx9* |
| ENSMUSG00000018765 | *Fxr2* | ENSMUSG00000042700 | *Sipa1l1* |
| ENSMUSG00000018796 | *Acsl1* | ENSMUSG00000042707 | *Dnali1* |
| ENSMUSG00000018830 | *Myh11* | ENSMUSG00000042708 | *Shcbp1l* |
| ENSMUSG00000018844 | *Fndc8* | ENSMUSG00000042726 | *Trafd1* |
| ENSMUSG00000018846 | *Pank3* | ENSMUSG00000042744 | *Hectd4* |
| ENSMUSG00000018848 | *Rars* | ENSMUSG00000042750 | *Bex2* |
| ENSMUSG00000018921 | *Pelp1* | ENSMUSG00000042772 | *Smg7* |
| ENSMUSG00000018974 | *Sart3* | ENSMUSG00000042790 | *Rnf214* |
| ENSMUSG00000019027 | *Dnah1* | ENSMUSG00000042851 | *Zc3h6* |
| ENSMUSG00000019132 | *BC005537* | ENSMUSG00000042961 | *Egflam* |
| ENSMUSG00000019139 | *Isyna1* | ENSMUSG00000043003 | *Rasef* |
| ENSMUSG00000019254 | *Ppp1r12c* | ENSMUSG00000043020 | *Dnai3* |
| ENSMUSG00000019278 | *Dpep1* | ENSMUSG00000043036 | *Ccdc63* |
| ENSMUSG00000019303 | *Psmc3ip* | ENSMUSG00000043050 | *Tnp2* |
| ENSMUSG00000019370 | *Calm3* | ENSMUSG00000043059 | *Zfp513* |
| ENSMUSG00000019428 | *Fkbp8* | ENSMUSG00000043067 | *Dpy19l1* |
| ENSMUSG00000019432 | *Ddx39b* | ENSMUSG00000043183 | *Simc1* |
| ENSMUSG00000019467 | *Arhgef25* | ENSMUSG00000043192 | *Gpi-ps* |
| ENSMUSG00000019471 | *Cdc37* | ENSMUSG00000043223 | *Gm4835* |
| ENSMUSG00000019494 | *Cops6* | ENSMUSG00000043241 | *Upf2* |
| ENSMUSG00000019505 | *Ubb* | ENSMUSG00000043319 | *Cox8c* |
| ENSMUSG00000019564 | *Arid3a* | ENSMUSG00000043323 | *Fbrsl1* |
| ENSMUSG00000019578 | *Ubxn6* | ENSMUSG00000043336 | *Filip1l* |
| ENSMUSG00000019659 | *Ccdc12* | ENSMUSG00000043384 | *Gprasp1* |
| ENSMUSG00000019689 | *Fmc1* | ENSMUSG00000043411 | *Usp48* |
| ENSMUSG00000019699 | *Akt3* | ENSMUSG00000043424 | *Eif3j2* |
| ENSMUSG00000019710 | *Mrpl24* | ENSMUSG00000043429 | *Ccdc185* |
| ENSMUSG00000019715 | *Gle1* | ENSMUSG00000043445 | *Pgp* |
| ENSMUSG00000019732 | *Calr3* | ENSMUSG00000043468 | *Adam30* |
| ENSMUSG00000019738 | *Polr2i* | ENSMUSG00000043483 | *Gm6863* |
| ENSMUSG00000019777 | *Hdac2* | ENSMUSG00000043535 | *Setx* |
| ENSMUSG00000019782 | *Rwdd1* | ENSMUSG00000043541 | *Casc1* |
| ENSMUSG00000019795 | *Pcmt1* | ENSMUSG00000043633 | *Fam221b* |
| ENSMUSG00000019802 | *Sec63* | ENSMUSG00000043635 | *Adamts3* |
| ENSMUSG00000019813 | *Cep57l1* | ENSMUSG00000043716 | *Rpl7* |
| ENSMUSG00000019814 | *Ltv1* | ENSMUSG00000043719 | *Col6a6* |
| ENSMUSG00000019818 | *Cd164* | ENSMUSG00000043733 | *Ptpn11* |
| ENSMUSG00000019820 | *Utrn* | ENSMUSG00000043873 | *Chil5* |
| ENSMUSG00000019837 | *Gtf3c6* | ENSMUSG00000043889 | *Gm8399* |
| ENSMUSG00000019841 | *Rev3l* | ENSMUSG00000043909 | *Trp53bp1* |

| | | | |
|---|---|---|---|
| ENSMUSG00000019843 | *Fyn* | ENSMUSG00000043913 | *Ccdc60* |
| ENSMUSG00000019856 | *Fam184a* | ENSMUSG00000043940 | *Wdfy3* |
| ENSMUSG00000019861 | *Gopc* | ENSMUSG00000043945 | *Adam6a* |
| ENSMUSG00000019876 | *Pkib* | ENSMUSG00000043962 | *Thrap3* |
| ENSMUSG00000019892 | *Lrriq1* | ENSMUSG00000043972 | *Opn5* |
| ENSMUSG00000019906 | *Lin7a* | ENSMUSG00000043987 | *Cep164* |
| ENSMUSG00000019907 | *Ppp1r12a* | ENSMUSG00000043998 | *Mgat2* |
| ENSMUSG00000019917 | *Septin10* | ENSMUSG00000044042 | *Fmn1* |
| ENSMUSG00000019923 | *Zwint* | ENSMUSG00000044068 | *Zrsr1* |
| ENSMUSG00000019933 | *Mrln* | ENSMUSG00000044072 | *Eml6* |
| ENSMUSG00000019943 | *Atp2b1* | ENSMUSG00000044083 | *Efcab8* |
| ENSMUSG00000019945 | *Cabcoco1* | ENSMUSG00000044084 | *Spem2* |
| ENSMUSG00000019966 | *Kitl* | ENSMUSG00000044098 | *Rsbn1* |
| ENSMUSG00000019969 | *Psen1* | ENSMUSG00000044117 | *Bmerb1* |
| ENSMUSG00000019971 | *Cep290* | ENSMUSG00000044122 | *Proca1* |
| ENSMUSG00000019977 | *Hbs1l* | ENSMUSG00000044141 | *E130201H02Rik* |
| ENSMUSG00000019978 | *Epb41l2* | ENSMUSG00000044206 | *Vsig4* |
| ENSMUSG00000019986 | *Ahi1* | ENSMUSG00000044211 | *Gm7887* |
| ENSMUSG00000019996 | *Map7* | ENSMUSG00000044224 | *Dnajc21* |
| ENSMUSG00000020015 | *Cdk17* | ENSMUSG00000044252 | *Osbpl1a* |
| ENSMUSG00000020020 | *Usp44* | ENSMUSG00000044268 | *Gm4895* |
| ENSMUSG00000020022 | *Ndufa12* | ENSMUSG00000044285 | *Ubb-ps* |
| ENSMUSG00000020024 | *Cep83* | ENSMUSG00000044308 | *Ubr3* |
| ENSMUSG00000020029 | *Nudt4* | ENSMUSG00000044340 | *Phlpp1* |
| ENSMUSG00000020037 | *Rfx4* | ENSMUSG00000044349 | *Snhg11* |
| ENSMUSG00000020048 | *Hsp90b1* | ENSMUSG00000044362 | *Ccdc89* |
| ENSMUSG00000020059 | *Sycp3* | ENSMUSG00000044468 | *Tent5c* |
| ENSMUSG00000020064 | *Herc4* | ENSMUSG00000044471 | *Lncpint* |
| ENSMUSG00000020069 | *Hnrnph3* | ENSMUSG00000044475 | *Ascc1* |
| ENSMUSG00000020074 | *Ccar1* | ENSMUSG00000044526 | *Znrf4* |
| ENSMUSG00000020075 | *Ddx21* | ENSMUSG00000044533 | *Rps2* |
| ENSMUSG00000020076 | *Ddx50* | ENSMUSG00000044566 | *Cage1* |
| ENSMUSG00000020088 | *Sar1a* | ENSMUSG00000044581 | *4932415D10Rik* |
| ENSMUSG00000020102 | *Slc16a7* | ENSMUSG00000044627 | *Swi5* |
| ENSMUSG00000020114 | *Cand1* | ENSMUSG00000044647 | *Csrnp3* |
| ENSMUSG00000020115 | *Tbk1* | ENSMUSG00000044701 | *Il27* |
| ENSMUSG00000020116 | *Pno1* | ENSMUSG00000044751 | *Atp5pb-ps* |
| ENSMUSG00000020124 | *Usp15* | ENSMUSG00000044757 | *Gm6430* |
| ENSMUSG00000020130 | *Tbc1d15* | ENSMUSG00000044783 | *Hjurp* |
| ENSMUSG00000020131 | *Pcsk4* | ENSMUSG00000044787 | *Spata32* |
| ENSMUSG00000020156 | *Pwwp3a* | ENSMUSG00000044791 | *Setd2* |
| ENSMUSG00000020164 | *Kcnmb4os1* | ENSMUSG00000044795 | *Cyb5d1* |
| ENSMUSG00000020184 | *Mdm2* | ENSMUSG00000044854 | *1700056E22Rik* |
| ENSMUSG00000020189 | *Osbpl8* | ENSMUSG00000044948 | *Cfap43* |
| ENSMUSG00000020191 | *Spata48* | ENSMUSG00000044949 | *Ubtd2* |
| ENSMUSG00000020193 | *Zpbp* | ENSMUSG00000045004 | *Spata21* |
| ENSMUSG00000020196 | *Cabin1* | ENSMUSG00000045031 | *Cetn4* |
| ENSMUSG00000020198 | *Ap3d1* | ENSMUSG00000045036 | *Tmem232* |
| ENSMUSG00000020212 | *Mdm1* | ENSMUSG00000045048 | *Gm9795* |
| ENSMUSG00000020214 | *Glipr1l2* | ENSMUSG00000045078 | *Rnf216* |
| ENSMUSG00000020224 | *Llph* | ENSMUSG00000045095 | *Magi1* |
| ENSMUSG00000020248 | *Nfyb* | ENSMUSG00000045103 | *Dmd* |
| ENSMUSG00000020250 | *Txnrd1* | ENSMUSG00000045104 | *Ldhb-ps* |
| ENSMUSG00000020260 | *Pofut2* | ENSMUSG00000045210 | *Vcpip1* |
| ENSMUSG00000020263 | *Appl2* | ENSMUSG00000045211 | *Nudt18* |
| ENSMUSG00000020270 | *Smim23* | ENSMUSG00000045252 | *Zfp574* |
| ENSMUSG00000020272 | *Stk10* | ENSMUSG00000045257 | *Morn2* |
| ENSMUSG00000020283 | *Pex13* | ENSMUSG00000045273 | *Cenph* |

| | | | |
|---|---|---|---|
| ENSMUSG00000020286 | *1700093K21Rik* | ENSMUSG00000045275 | *Lca5l* |
| ENSMUSG00000020290 | *Xpo1* | ENSMUSG00000045328 | *Cenpe* |
| ENSMUSG00000020307 | *Cdc34* | ENSMUSG00000045330 | *Magec2* |
| ENSMUSG00000020311 | *Erlec1* | ENSMUSG00000045350 | *Fam186a* |
| ENSMUSG00000020315 | *Sptbn1* | ENSMUSG00000045409 | *Trim39* |
| ENSMUSG00000020317 | *Theg* | ENSMUSG00000045411 | *2410002F23Rik* |
| ENSMUSG00000020319 | *Wdpcp* | ENSMUSG00000045427 | *Hnrnph2* |
| ENSMUSG00000020321 | *Mdh1* | ENSMUSG00000045466 | *Zfp956* |
| ENSMUSG00000020328 | *Nudcd2* | ENSMUSG00000045467 | *Ttll13* |
| ENSMUSG00000020330 | *Hmmr* | ENSMUSG00000045482 | *Trrap* |
| ENSMUSG00000020332 | *Meikin* | ENSMUSG00000045521 | *Tssk2* |
| ENSMUSG00000020333 | *Acsl6* | ENSMUSG00000045573 | *Penk* |
| ENSMUSG00000020349 | *Ppp2ca* | ENSMUSG00000045624 | *Esf1* |
| ENSMUSG00000020358 | *Hnrnpab* | ENSMUSG00000045672 | *Col27a1* |
| ENSMUSG00000020361 | *Hspa4* | ENSMUSG00000045690 | *Wdr89* |
| ENSMUSG00000020368 | *Canx* | ENSMUSG00000045709 | *Smkr-ps* |
| ENSMUSG00000020380 | *Rad50* | ENSMUSG00000045763 | *Basp1* |
| ENSMUSG00000020385 | *Clk4* | ENSMUSG00000045794 | *Lkaaear1* |
| ENSMUSG00000020389 | *Cdkl3* | ENSMUSG00000045795 | *Whamm* |
| ENSMUSG00000020401 | *Garin3* | ENSMUSG00000045799 | *Gm9800* |
| ENSMUSG00000020409 | *Slu7* | ENSMUSG00000045835 | *Hdgfl1* |
| ENSMUSG00000020412 | *Ascc2* | ENSMUSG00000045886 | *Pam16l* |
| ENSMUSG00000020430 | *Pes1* | ENSMUSG00000045896 | *Paip2b* |
| ENSMUSG00000020434 | *4921536K21Rik* | ENSMUSG00000045915 | *Ccdc42* |
| ENSMUSG00000020435 | *Osbp2* | ENSMUSG00000045928 | *4933440M02Rik* |
| ENSMUSG00000020441 | *2310033P09Rik* | ENSMUSG00000045942 | *BC049762* |
| ENSMUSG00000020454 | *Eif4enif1* | ENSMUSG00000045962 | *Wnk1* |
| ENSMUSG00000020456 | *Ogdh* | ENSMUSG00000045969 | *Ing1* |
| ENSMUSG00000020462 | *Cfap36* | ENSMUSG00000045983 | *Eif4g1* |
| ENSMUSG00000020463 | *Ppp4r3b* | ENSMUSG00000046010 | *Zfp830* |
| ENSMUSG00000020472 | *Zkscan17* | ENSMUSG00000046056 | *Sbsn* |
| ENSMUSG00000020475 | *Pgam2* | ENSMUSG00000046062 | *Ppp1r15b* |
| ENSMUSG00000020476 | *Dbnl* | ENSMUSG00000046085 | *4931422A03Rik* |
| ENSMUSG00000020481 | *Ankrd36* | ENSMUSG00000046111 | *Cep295* |
| ENSMUSG00000020483 | *Dynll2* | ENSMUSG00000046138 | *9930021J03Rik* |
| ENSMUSG00000020486 | *Septin4* | ENSMUSG00000046173 | *Pabpc6* |
| ENSMUSG00000020491 | *2810021J22Rik* | ENSMUSG00000046192 | *Iqub* |
| ENSMUSG00000020496 | *Rnf187* | ENSMUSG00000046196 | *Ttc39d* |
| ENSMUSG00000020515 | *Cnot8* | ENSMUSG00000046229 | *Scand1* |
| ENSMUSG00000020516 | *Rps6kb1* | ENSMUSG00000046230 | *Vps13a* |
| ENSMUSG00000020520 | *Galnt10* | ENSMUSG00000046337 | *Fam178b* |
| ENSMUSG00000020525 | *Ppm1d* | ENSMUSG00000046338 | *Gpat2* |
| ENSMUSG00000020530 | *Ggnbp2* | ENSMUSG00000046341 | *Gm11223* |
| ENSMUSG00000020532 | *Acaca* | ENSMUSG00000046351 | *Zfp322a* |
| ENSMUSG00000020562 | *Efcab10* | ENSMUSG00000046372 | *Gm5590* |
| ENSMUSG00000020571 | *Pdia6* | ENSMUSG00000046434 | *Hnrnpa1* |
| ENSMUSG00000020580 | *Rock2* | ENSMUSG00000046440 | *Gm5564* |
| ENSMUSG00000020593 | *Lpin1* | ENSMUSG00000046447 | *Camk2n1* |
| ENSMUSG00000020594 | *Pum2* | ENSMUSG00000046516 | *Cox17* |
| ENSMUSG00000020608 | *Smc6* | ENSMUSG00000046573 | *Lyrm4* |
| ENSMUSG00000020610 | *Amz2* | ENSMUSG00000046585 | *Cfap58* |
| ENSMUSG00000020612 | *Prkar1a* | ENSMUSG00000046717 | *Igbp1b* |
| ENSMUSG00000020617 | *1700012B07Rik* | ENSMUSG00000046722 | *Cdc42se1* |
| ENSMUSG00000020622 | *Nt5c1b* | ENSMUSG00000046723 | *Adam24* |
| ENSMUSG00000020636 | *Allc* | ENSMUSG00000046727 | *Cystm1* |
| ENSMUSG00000020639 | *Pfn4* | ENSMUSG00000046750 | *Selenov* |
| ENSMUSG00000020640 | *Itsn2* | ENSMUSG00000046753 | *Ccdc66* |
| ENSMUSG00000020652 | *Cenpo* | ENSMUSG00000046755 | *Kif2b* |

| | | | |
|---|---|---|---|
| ENSMUSG00000020654 | *Adcy3* | ENSMUSG00000046782 | *Ttc6* |
| ENSMUSG00000020657 | *Dnajc27* | ENSMUSG00000046841 | *Ckap4* |
| ENSMUSG00000020671 | *Rab10* | ENSMUSG00000046846 | *Spesp1* |
| ENSMUSG00000020677 | *Ddx52* | ENSMUSG00000046865 | *Fbl* |
| ENSMUSG00000020680 | *Taf15* | ENSMUSG00000046934 | *Csl* |
| ENSMUSG00000020681 | *Ace* | ENSMUSG00000046942 | *Mageb16* |
| ENSMUSG00000020687 | *Cdc27* | ENSMUSG00000046957 | *Spz1* |
| ENSMUSG00000020690 | *Efcab3* | ENSMUSG00000046958 | *4930432E11Rik* |
| ENSMUSG00000020694 | *Tlk2* | ENSMUSG00000047021 | *Cfap65* |
| ENSMUSG00000020697 | *Lig3* | ENSMUSG00000047022 | *Mipol1* |
| ENSMUSG00000020705 | *Ddx42* | ENSMUSG00000047025 | *Ccer1* |
| ENSMUSG00000020706 | *Ftsj3* | ENSMUSG00000047036 | *Zfp445* |
| ENSMUSG00000020708 | *Psmc5* | ENSMUSG00000047104 | *Pbp2* |
| ENSMUSG00000020719 | *Ddx5* | ENSMUSG00000047108 | *Dnajb7* |
| ENSMUSG00000020720 | *Psmd12* | ENSMUSG00000047126 | *Cltc* |
| ENSMUSG00000020721 | *Helz* | ENSMUSG00000047129 | *1700113H08Rik* |
| ENSMUSG00000020728 | *Cep112* | ENSMUSG00000047141 | *Zfp654* |
| ENSMUSG00000020737 | *Jpt1* | ENSMUSG00000047193 | *Dync2h1* |
| ENSMUSG00000020743 | *Mif4gd* | ENSMUSG00000047248 | *C2cd3* |
| ENSMUSG00000020745 | *Pafah1b1* | ENSMUSG00000047361 | *Gm973* |
| ENSMUSG00000020755 | *Sap30bp* | ENSMUSG00000047369 | *Dnah14* |
| ENSMUSG00000020775 | *Mrpl38* | ENSMUSG00000047417 | *Rexo1* |
| ENSMUSG00000020776 | *Fbf1* | ENSMUSG00000047446 | *Arl4a* |
| ENSMUSG00000020783 | *Ncbp3* | ENSMUSG00000047454 | *Gphn* |
| ENSMUSG00000020792 | *Exoc7* | ENSMUSG00000047466 | *8030462N17Rik* |
| ENSMUSG00000020799 | *Tekt1* | ENSMUSG00000047502 | *Mroh7* |
| ENSMUSG00000020802 | *Ube2o* | ENSMUSG00000047509 | *Gm6776* |
| ENSMUSG00000020807 | *4933427D14Rik* | ENSMUSG00000047514 | *Tspyl1* |
| ENSMUSG00000020814 | *Mxra7* | ENSMUSG00000047518 | *Slfnl1* |
| ENSMUSG00000020817 | *Rabep1* | ENSMUSG00000047528 | *Flacc1* |
| ENSMUSG00000020823 | *Sec14l1* | ENSMUSG00000047547 | *Cltb* |
| ENSMUSG00000020827 | *Mink1* | ENSMUSG00000047619 | *Ddi1* |
| ENSMUSG00000020844 | *Nxn* | ENSMUSG00000047648 | *Fbxo30* |
| ENSMUSG00000020849 | *Ywhae* | ENSMUSG00000047649 | *Cd3eap* |
| ENSMUSG00000020850 | *Prpf8* | ENSMUSG00000047674 | *Pdha2* |
| ENSMUSG00000020859 | *Spag9* | ENSMUSG00000047675 | *Rps8* |
| ENSMUSG00000020863 | *Luc7l3* | ENSMUSG00000047694 | *Yipf6* |
| ENSMUSG00000020867 | *Spata20* | ENSMUSG00000047696 | *Ccdc144b* |
| ENSMUSG00000020869 | *Lrrc59* | ENSMUSG00000047720 | *Clec2m* |
| ENSMUSG00000020870 | *Cdc34b* | ENSMUSG00000047777 | *Phf13* |
| ENSMUSG00000020878 | *Lrrc46* | ENSMUSG00000047810 | *Ccdc88b* |
| ENSMUSG00000020882 | *Cacnb1* | ENSMUSG00000047821 | *Trim16* |
| ENSMUSG00000020893 | *Per1* | ENSMUSG00000047841 | *Fndc11* |
| ENSMUSG00000020900 | *Myh10* | ENSMUSG00000047866 | *Lonp2* |
| ENSMUSG00000020903 | *Stx8* | ENSMUSG00000047878 | *A4galt* |
| ENSMUSG00000020904 | *Cfap52* | ENSMUSG00000047888 | *Tnrc6b* |
| ENSMUSG00000020914 | *Top2a* | ENSMUSG00000047905 | *Gm8566* |
| ENSMUSG00000020923 | *Ubtf* | ENSMUSG00000047907 | *Tshz2* |
| ENSMUSG00000020925 | *Ccdc43* | ENSMUSG00000047940 | *Stpg2* |
| ENSMUSG00000020929 | *Eftud2* | ENSMUSG00000047945 | *Marcksl1* |
| ENSMUSG00000020936 | *Nmt1* | ENSMUSG00000047995 | *Cypt4* |
| ENSMUSG00000020945 | *Lyzl6* | ENSMUSG00000048000 | *Gigyf2* |
| ENSMUSG00000020954 | *Strn3* | ENSMUSG00000048003 | *Catsper4* |
| ENSMUSG00000020962 | *Gtf2a1* | ENSMUSG00000048077 | *H1f7* |
| ENSMUSG00000020964 | *Sel1l* | ENSMUSG00000048096 | *Lmod1* |
| ENSMUSG00000020978 | *Klhdc2* | ENSMUSG00000048100 | *Taf13* |
| ENSMUSG00000020982 | *Nemf* | ENSMUSG00000048118 | *Arid4a* |
| ENSMUSG00000020986 | *Sec23a* | ENSMUSG00000048154 | *Kmt2d* |

| | | | |
|---|---|---|---|
| ENSMUSG00000020993 | *Trappc6b* | ENSMUSG00000048222 | *Mfap1b* |
| ENSMUSG00000020994 | *Pnn* | ENSMUSG00000048230 | *Fbxo43* |
| ENSMUSG00000021003 | *Galc* | ENSMUSG00000048249 | *Crebrf* |
| ENSMUSG00000021007 | *Spata7* | ENSMUSG00000048271 | *Rbm33* |
| ENSMUSG00000021012 | *Zc3h14* | ENSMUSG00000048385 | *Scrt1* |
| ENSMUSG00000021022 | *Ppp2r3c* | ENSMUSG00000048410 | *Zfp407* |
| ENSMUSG00000021027 | *Ralgapa1* | ENSMUSG00000048411 | *Gm597* |
| ENSMUSG00000021036 | *Sptlc2* | ENSMUSG00000048416 | *Mlf1* |
| ENSMUSG00000021037 | *Ahsa1* | ENSMUSG00000048445 | *Ccdc57* |
| ENSMUSG00000021039 | *Snw1* | ENSMUSG00000048478 | *Spata33* |
| ENSMUSG00000021051 | *Ppp2r5e* | ENSMUSG00000048520 | *Fbxl13* |
| ENSMUSG00000021056 | *Tex21* | ENSMUSG00000048578 | *Mlec* |
| ENSMUSG00000021061 | *Sptb* | ENSMUSG00000048592 | *Gm5946* |
| ENSMUSG00000021065 | *Fut8* | ENSMUSG00000048602 | *Morc2b* |
| ENSMUSG00000021066 | *Atl1* | ENSMUSG00000048647 | *Exd1* |
| ENSMUSG00000021067 | *Sav1* | ENSMUSG00000048652 | *Samd13* |
| ENSMUSG00000021078 | *Tomm20l* | ENSMUSG00000048655 | *Ccdc169* |
| ENSMUSG00000021086 | *Ccdc175* | ENSMUSG00000048661 | *Lemd3* |
| ENSMUSG00000021090 | *Lrrc9* | ENSMUSG00000048686 | *Hmgb4* |
| ENSMUSG00000021096 | *Ppm1a* | ENSMUSG00000048696 | *Mex3d* |
| ENSMUSG00000021097 | *Clmn* | ENSMUSG00000048701 | *Ccdc6* |
| ENSMUSG00000021109 | *Hif1a* | ENSMUSG00000048707 | *Tprn* |
| ENSMUSG00000021111 | *Papola* | ENSMUSG00000048731 | *Ggnbp1* |
| ENSMUSG00000021112 | *Pals1* | ENSMUSG00000048732 | *Klhl11* |
| ENSMUSG00000021113 | *Snapc1* | ENSMUSG00000048787 | *Dcun1d3* |
| ENSMUSG00000021114 | *Atp6v1d* | ENSMUSG00000048794 | *Cfap100* |
| ENSMUSG00000021115 | *Vrk1* | ENSMUSG00000048799 | *Cep120* |
| ENSMUSG00000021116 | *Eif2s1* | ENSMUSG00000048865 | *Arhgap30* |
| ENSMUSG00000021124 | *Vti1b* | ENSMUSG00000048874 | *Phf3* |
| ENSMUSG00000021134 | *Srsf5* | ENSMUSG00000048878 | *Hexim1* |
| ENSMUSG00000021139 | *Gm20498* | ENSMUSG00000048911 | *Rnf24* |
| ENSMUSG00000021140 | *Pcnx* | ENSMUSG00000048922 | *Cdca2* |
| ENSMUSG00000021143 | *Pacs2* | ENSMUSG00000048930 | *Tada3* |
| ENSMUSG00000021149 | *Gtpbp4* | ENSMUSG00000048967 | *Yjefn3* |
| ENSMUSG00000021156 | *Zmynd11* | ENSMUSG00000049091 | *Sephs2* |
| ENSMUSG00000021178 | *Psmc1* | ENSMUSG00000049106 | *Dcaf5* |
| ENSMUSG00000021179 | *Nrde2* | ENSMUSG00000049123 | *Catsperg2* |
| ENSMUSG00000021182 | *Ccdc88c* | ENSMUSG00000049160 | *Tex50* |
| ENSMUSG00000021188 | *Trip11* | ENSMUSG00000049230 | *Myef2l* |
| ENSMUSG00000021189 | *Atxn3* | ENSMUSG00000049295 | *Zfp219* |
| ENSMUSG00000021192 | *Golga5* | ENSMUSG00000049327 | *Kmt5a* |
| ENSMUSG00000021194 | *Chga* | ENSMUSG00000049401 | *Ogfr* |
| ENSMUSG00000021196 | *Pfkp* | ENSMUSG00000049470 | *Aff4* |
| ENSMUSG00000021203 | *Otub2* | ENSMUSG00000049550 | *Clip1* |
| ENSMUSG00000021218 | *Gdi2* | ENSMUSG00000049553 | *Polr1a* |
| ENSMUSG00000021221 | *Dpf3* | ENSMUSG00000049571 | *Cfap46* |
| ENSMUSG00000021234 | *Fam161b* | ENSMUSG00000049658 | *Bdp1* |
| ENSMUSG00000021244 | *Ylpm1* | ENSMUSG00000049659 | *Aftph* |
| ENSMUSG00000021248 | *Tmed10* | ENSMUSG00000049676 | *Catsperg1* |
| ENSMUSG00000021258 | *Ccnk* | ENSMUSG00000049739 | *Zfp646* |
| ENSMUSG00000021262 | *Evl* | ENSMUSG00000049760 | *Micos13* |
| ENSMUSG00000021264 | *Yy1* | ENSMUSG00000049792 | *Bag5* |
| ENSMUSG00000021268 | *Meg3* | ENSMUSG00000049800 | *Sertad2* |
| ENSMUSG00000021270 | *Hsp90aa1* | ENSMUSG00000049807 | *Arhgap23* |
| ENSMUSG00000021273 | *Fdft1* | ENSMUSG00000049811 | *Fam161a* |
| ENSMUSG00000021279 | *Cdc42bpb* | ENSMUSG00000049916 | *2610318N02Rik* |
| ENSMUSG00000021281 | *Tnfaip2* | ENSMUSG00000049940 | *Pgrmc2* |
| ENSMUSG00000021282 | *Eif5* | ENSMUSG00000049957 | *Ccdc137* |

| | | | |
|---|---|---|---|
| ENSMUSG00000021288 | *Klc1* | ENSMUSG00000049971 | *Glt1d1* |
| ENSMUSG00000021301 | *Hecw1* | ENSMUSG00000050017 | *Pitpnb* |
| ENSMUSG00000021311 | *Mtr* | ENSMUSG00000050035 | *Fhl4* |
| ENSMUSG00000021338 | *Carmil1* | ENSMUSG00000050050 | *Ccdc158* |
| ENSMUSG00000021339 | *Mrs2* | ENSMUSG00000050089 | *Akap4* |
| ENSMUSG00000021363 | *Mak* | ENSMUSG00000050107 | *Haspin* |
| ENSMUSG00000021366 | *Hivep1* | ENSMUSG00000050114 | *Prdx6b* |
| ENSMUSG00000021374 | *Nup153* | ENSMUSG00000050122 | *Vwa3b* |
| ENSMUSG00000021377 | *Dek* | ENSMUSG00000050141 | *Fam205c* |
| ENSMUSG00000021379 | *Id4* | ENSMUSG00000050150 | *Slc9b1* |
| ENSMUSG00000021384 | *Susd3* | ENSMUSG00000050213 | *Snip1* |
| ENSMUSG00000021392 | *Nol8* | ENSMUSG00000050243 | *Gm5446* |
| ENSMUSG00000021413 | *Prpf4b* | ENSMUSG00000050299 | *Gm9843* |
| ENSMUSG00000021414 | *Fam217a* | ENSMUSG00000050379 | *Septin6* |
| ENSMUSG00000021415 | *4933417A18Rik* | ENSMUSG00000050530 | *Fam171a1* |
| ENSMUSG00000021420 | *Fars2* | ENSMUSG00000050553 | *Gk2* |
| ENSMUSG00000021427 | *Ssr1* | ENSMUSG00000050555 | *Hyls1* |
| ENSMUSG00000021428 | *Riok1* | ENSMUSG00000050565 | *Tor1aip2* |
| ENSMUSG00000021431 | *Snrnp48* | ENSMUSG00000050592 | *Fam78a* |
| ENSMUSG00000021476 | *Habp4* | ENSMUSG00000050612 | *Txndc2* |
| ENSMUSG00000021477 | *Ctsl* | ENSMUSG00000050623 | *Catsperz* |
| ENSMUSG00000021488 | *Nsd1* | ENSMUSG00000050668 | *Gpatch11* |
| ENSMUSG00000021494 | *Ddx41* | ENSMUSG00000050677 | *Ccdc96* |
| ENSMUSG00000021499 | *Catsper3* | ENSMUSG00000050685 | *Ccdc54* |
| ENSMUSG00000021500 | *Ddx46* | ENSMUSG00000050702 | *4930563M21Rik* |
| ENSMUSG00000021501 | *Caml* | ENSMUSG00000050860 | *Phospho1* |
| ENSMUSG00000021534 | *1700001L19Rik* | ENSMUSG00000050900 | *Gm7327* |
| ENSMUSG00000021537 | *Cetn3* | ENSMUSG00000050944 | *Efcab5* |
| ENSMUSG00000021546 | *Hnrnpk* | ENSMUSG00000050957 | *Insl6* |
| ENSMUSG00000021549 | *Rasa1* | ENSMUSG00000050994 | *Adgb* |
| ENSMUSG00000021552 | *Gkap1* | ENSMUSG00000050996 | *Cetn1* |
| ENSMUSG00000021557 | *Agtpbp1* | ENSMUSG00000051036 | *Ttc24* |
| ENSMUSG00000021577 | *Sdha* | ENSMUSG00000051111 | *Sv2c* |
| ENSMUSG00000021585 | *Cast* | ENSMUSG00000051149 | *Adnp* |
| ENSMUSG00000021595 | *Nsun2* | ENSMUSG00000051166 | *Eml5* |
| ENSMUSG00000021596 | *Mctp1* | ENSMUSG00000051185 | *Fam174a* |
| ENSMUSG00000021598 | *Med10* | ENSMUSG00000051223 | *Bzw1* |
| ENSMUSG00000021615 | *Xrcc4* | ENSMUSG00000051255 | *Gm6563* |
| ENSMUSG00000021619 | *Atg10* | ENSMUSG00000051257 | *Trap1a* |
| ENSMUSG00000021635 | *Rad17* | ENSMUSG00000051276 | *Actrt2* |
| ENSMUSG00000021639 | *Gtf2h2* | ENSMUSG00000051285 | *Pcmtd1* |
| ENSMUSG00000021643 | *Serf1* | ENSMUSG00000051306 | *Usp42* |
| ENSMUSG00000021668 | *Polk* | ENSMUSG00000051316 | *Taf7* |
| ENSMUSG00000021669 | *Cert1* | ENSMUSG00000051343 | *Rab11fip5* |
| ENSMUSG00000021670 | *Hmgcr* | ENSMUSG00000051403 | *Ppp1r37* |
| ENSMUSG00000021671 | *Poc5* | ENSMUSG00000051427 | *Ccdc157* |
| ENSMUSG00000021676 | *Iqgap2* | ENSMUSG00000051435 | *Fhad1* |
| ENSMUSG00000021681 | *Aggf1* | ENSMUSG00000051451 | *Crebzf* |
| ENSMUSG00000021686 | *Ap3b1* | ENSMUSG00000051586 | *Mical3* |
| ENSMUSG00000021690 | *Jmy* | ENSMUSG00000051618 | *Ubqln3* |
| ENSMUSG00000021693 | *Kif2a* | ENSMUSG00000051648 | *Kctd19* |
| ENSMUSG00000021699 | *Pde4d* | ENSMUSG00000051674 | *Dcun1d4* |
| ENSMUSG00000021709 | *Erbin* | ENSMUSG00000051695 | *Pcbp1* |
| ENSMUSG00000021712 | *Trim23* | ENSMUSG00000051728 | *Fam243* |
| ENSMUSG00000021713 | *Ppwd1* | ENSMUSG00000051732 | *Pabpc2* |
| ENSMUSG00000021715 | *Cwc27* | ENSMUSG00000051736 | *Fam229b* |
| ENSMUSG00000021716 | *Srek1ip1* | ENSMUSG00000051768 | *Xrcc1* |
| ENSMUSG00000021733 | *Slc4a7* | ENSMUSG00000051811 | *Cox6b2* |

| | | | |
|---|---|---|---|
| ENSMUSG00000021737 | *Psmd6* | ENSMUSG00000051900 | *Abca16* |
| ENSMUSG00000021741 | *Gm5457* | ENSMUSG00000051910 | *Sox6* |
| ENSMUSG00000021748 | *Pdhb* | ENSMUSG00000051934 | *Spats2* |
| ENSMUSG00000021754 | *Map3k1* | ENSMUSG00000051956 | *Rnf133* |
| ENSMUSG00000021758 | *Ddx4* | ENSMUSG00000051989 | *Smim11* |
| ENSMUSG00000021767 | *Kat6b* | ENSMUSG00000052056 | *Zfp217* |
| ENSMUSG00000021771 | *Vdac2* | ENSMUSG00000052062 | *Pard3b* |
| ENSMUSG00000021775 | *Nr1d2* | ENSMUSG00000052075 | *1700029F12Rik* |
| ENSMUSG00000021790 | *Dydc1* | ENSMUSG00000052105 | *Mtcl1* |
| ENSMUSG00000021791 | *Dydc2* | ENSMUSG00000052137 | *Rbm12b2* |
| ENSMUSG00000021807 | *Rtraf* | ENSMUSG00000052144 | *Ppp4r2* |
| ENSMUSG00000021811 | *Dnajc9* | ENSMUSG00000052221 | *Ppp1r36* |
| ENSMUSG00000021819 | *Zswim8* | ENSMUSG00000052253 | *Zfp622* |
| ENSMUSG00000021823 | *Vcl* | ENSMUSG00000052273 | *Dnah3* |
| ENSMUSG00000021832 | *Psmc6* | ENSMUSG00000052276 | *Ostn* |
| ENSMUSG00000021838 | *Samd4* | ENSMUSG00000052293 | *Taf9* |
| ENSMUSG00000021843 | *Ktn1* | ENSMUSG00000052296 | *Ppp6r1* |
| ENSMUSG00000021846 | *Peli2* | ENSMUSG00000052337 | *Immt* |
| ENSMUSG00000021850 | *ccdc198* | ENSMUSG00000052372 | *Il1rapl1* |
| ENSMUSG00000021870 | *Slmap* | ENSMUSG00000052387 | *Trpm3* |
| ENSMUSG00000021879 | *Dnah12* | ENSMUSG00000052397 | *Ezr* |
| ENSMUSG00000021891 | *Mettl6* | ENSMUSG00000052407 | *Ccdc171* |
| ENSMUSG00000021892 | *Sh3bp5* | ENSMUSG00000052428 | *Tmco1* |
| ENSMUSG00000021901 | *Bap1* | ENSMUSG00000052469 | *Tcp10c* |
| ENSMUSG00000021902 | *Phf7* | ENSMUSG00000052488 | *Cherp* |
| ENSMUSG00000021911 | *Parg* | ENSMUSG00000052525 | *Spdya* |
| ENSMUSG00000021916 | *Glt8d1* | ENSMUSG00000052539 | *Magi3* |
| ENSMUSG00000021918 | *Nek4* | ENSMUSG00000052566 | *Hook2* |
| ENSMUSG00000021936 | *Mapk8* | ENSMUSG00000052613 | *Pcdh15* |
| ENSMUSG00000021938 | *Pspc1* | ENSMUSG00000052642 | *Lypd9* |
| ENSMUSG00000021940 | *Ptpn20* | ENSMUSG00000052676 | *Zmat1* |
| ENSMUSG00000021945 | *Zmym2* | ENSMUSG00000052698 | *Tln2* |
| ENSMUSG00000021948 | *Prkcd* | ENSMUSG00000052713 | *Zfp608* |
| ENSMUSG00000021958 | *Pinx1* | ENSMUSG00000052727 | *Map1b* |
| ENSMUSG00000021959 | *Lats2* | ENSMUSG00000052783 | *Grk4* |
| ENSMUSG00000021963 | *Sap18* | ENSMUSG00000052794 | *1700030K09Rik* |
| ENSMUSG00000021966 | *Prss52* | ENSMUSG00000052812 | *Atad2b* |
| ENSMUSG00000021972 | *Hmbox1* | ENSMUSG00000052825 | *Gm9892* |
| ENSMUSG00000021977 | *1700129C05Rik* | ENSMUSG00000052833 | *Sae1* |
| ENSMUSG00000021983 | *Atp8a2* | ENSMUSG00000052861 | *Dnah6* |
| ENSMUSG00000021986 | *Amer2* | ENSMUSG00000052906 | *Ubxn8* |
| ENSMUSG00000021987 | *Mtmr6* | ENSMUSG00000052915 | *Msl1* |
| ENSMUSG00000021997 | *Lrrc63* | ENSMUSG00000052926 | *Rnaseh2a* |
| ENSMUSG00000022000 | *Zc3h13* | ENSMUSG00000053012 | *Krcc1* |
| ENSMUSG00000022010 | *Tsc22d1* | ENSMUSG00000053030 | *Spink2* |
| ENSMUSG00000022013 | *Dnajc15* | ENSMUSG00000053038 | *Gm6180* |
| ENSMUSG00000022014 | *Epsti1* | ENSMUSG00000053110 | *Yap1* |
| ENSMUSG00000022016 | *Akap11* | ENSMUSG00000053111 | *Fank1* |
| ENSMUSG00000022019 | *Tdrd3* | ENSMUSG00000053119 | *Chmp3* |
| ENSMUSG00000022023 | *Wbp4* | ENSMUSG00000053153 | *Spag16* |
| ENSMUSG00000022024 | *Sugt1* | ENSMUSG00000053184 | *Spaca3* |
| ENSMUSG00000022032 | *Scara5* | ENSMUSG00000053199 | *Arhgap20* |
| ENSMUSG00000022035 | *Ccdc25* | ENSMUSG00000053205 | *Styx* |
| ENSMUSG00000022037 | *Clu* | ENSMUSG00000053286 | *Trmt1l* |
| ENSMUSG00000022039 | *Adam2* | ENSMUSG00000053289 | *Ddx10* |
| ENSMUSG00000022064 | *Pibf1* | ENSMUSG00000053293 | *Pom121* |
| ENSMUSG00000022085 | *Pebp4* | ENSMUSG00000053332 | *Gas5* |
| ENSMUSG00000022089 | *Bin3* | ENSMUSG00000053375 | *Atp6v1e2* |

| | | | |
|---|---|---|---|
| ENSMUSG00000022091 | *Sorbs3* | ENSMUSG00000053453 | *Thoc7* |
| ENSMUSG00000022092 | *Ppp3cc* | ENSMUSG00000053470 | *Kdm3a* |
| ENSMUSG00000022105 | *Rb1* | ENSMUSG00000053477 | *Tcf4* |
| ENSMUSG00000022108 | *Itm2b* | ENSMUSG00000053508 | *Gtsf2* |
| ENSMUSG00000022110 | *Sucla2* | ENSMUSG00000053510 | *Nrd1* |
| ENSMUSG00000022111 | *Uchl3* | ENSMUSG00000053536 | *Cstf2t* |
| ENSMUSG00000022119 | *Rbm26* | ENSMUSG00000053553 | *3110082I17Rik* |
| ENSMUSG00000022136 | *Dnajc3* | ENSMUSG00000053624 | *Gykl1* |
| ENSMUSG00000022141 | *Nipbl* | ENSMUSG00000053754 | *Chd8* |
| ENSMUSG00000022148 | *Fyb* | ENSMUSG00000053768 | *Chchd3* |
| ENSMUSG00000022151 | *Ttc33* | ENSMUSG00000053769 | *Lysmd1* |
| ENSMUSG00000022155 | *Mroh2b* | ENSMUSG00000053783 | *1700016K19Rik* |
| ENSMUSG00000022177 | *Haus4* | ENSMUSG00000053838 | *Nudcd3* |
| ENSMUSG00000022185 | *Acin1* | ENSMUSG00000053841 | *Txlna* |
| ENSMUSG00000022191 | *Drosha* | ENSMUSG00000053877 | *Srcap* |
| ENSMUSG00000022194 | *Pabpn1* | ENSMUSG00000053896 | *4933409G03Rik* |
| ENSMUSG00000022197 | *Pdzd2* | ENSMUSG00000053898 | *Ech1* |
| ENSMUSG00000022200 | *Golph3* | ENSMUSG00000054003 | *Tdrd9* |
| ENSMUSG00000022201 | *Zfr* | ENSMUSG00000054051 | *Ercc6* |
| ENSMUSG00000022205 | *Sub1* | ENSMUSG00000054079 | *Utp18* |
| ENSMUSG00000022214 | *Dcaf11* | ENSMUSG00000054150 | *Syne3* |
| ENSMUSG00000022216 | *Psme1* | ENSMUSG00000054199 | *Gon4l* |
| ENSMUSG00000022234 | *Cct5* | ENSMUSG00000054237 | *Fra10ac1* |
| ENSMUSG00000022236 | *Ropn1l* | ENSMUSG00000054277 | *Arfgap3* |
| ENSMUSG00000022246 | *Rai14* | ENSMUSG00000054302 | *Eapp* |
| ENSMUSG00000022247 | *Brix1* | ENSMUSG00000054320 | *Lrrc36* |
| ENSMUSG00000022249 | *Ttc23l* | ENSMUSG00000054405 | *Dnajc8* |
| ENSMUSG00000022255 | *Mtdh* | ENSMUSG00000054428 | *Atpif1* |
| ENSMUSG00000022263 | *Trio* | ENSMUSG00000054452 | *Tle5* |
| ENSMUSG00000022269 | *Marchf11* | ENSMUSG00000054455 | *Vapb* |
| ENSMUSG00000022272 | *Myo10* | ENSMUSG00000054611 | *Kdm2a* |
| ENSMUSG00000022280 | *Rnf19a* | ENSMUSG00000054717 | *Hmgb2* |
| ENSMUSG00000022283 | *Pabpc1* | ENSMUSG00000054727 | *1700013H16Rik* |
| ENSMUSG00000022285 | *Ywhaz* | ENSMUSG00000054728 | *Phactr1* |
| ENSMUSG00000022306 | *Zfpm2* | ENSMUSG00000054752 | *Fsd1l* |
| ENSMUSG00000022307 | *Oxr1* | ENSMUSG00000054766 | *Set* |
| ENSMUSG00000022312 | *Eif3h* | ENSMUSG00000054863 | *Tafa5* |
| ENSMUSG00000022314 | *Rad21* | ENSMUSG00000054885 | *4930578G10Rik* |
| ENSMUSG00000022329 | *Stk3* | ENSMUSG00000054901 | *Arhgef33* |
| ENSMUSG00000022332 | *Khdrbs3* | ENSMUSG00000054909 | *Wbscr25* |
| ENSMUSG00000022339 | *Ebag9* | ENSMUSG00000055024 | *Ep300* |
| ENSMUSG00000022360 | *Atad2* | ENSMUSG00000055065 | *Ddx17* |
| ENSMUSG00000022364 | *Tbc1d31* | ENSMUSG00000055078 | *Gabra5* |
| ENSMUSG00000022375 | *Lrrc6* | ENSMUSG00000055148 | *Klf2* |
| ENSMUSG00000022377 | *Asap1* | ENSMUSG00000055204 | *Ankrd17* |
| ENSMUSG00000022387 | *Brd1* | ENSMUSG00000055214 | *Pld5* |
| ENSMUSG00000022388 | *Ttll8* | ENSMUSG00000055296 | *Tmem245* |
| ENSMUSG00000022389 | *Tef* | ENSMUSG00000055319 | *Sec23ip* |
| ENSMUSG00000022390 | *Zc3h7b* | ENSMUSG00000055322 | *Tns1* |
| ENSMUSG00000022391 | *Rangap1* | ENSMUSG00000055334 | *Snupn* |
| ENSMUSG00000022394 | *L3mbtl2* | ENSMUSG00000055421 | *Pcdh9* |
| ENSMUSG00000022403 | *St13* | ENSMUSG00000055436 | *Srsf11* |
| ENSMUSG00000022419 | *Deptor* | ENSMUSG00000055531 | *Cpsf6* |
| ENSMUSG00000022420 | *Dnal4* | ENSMUSG00000055602 | *Tcp10b* |
| ENSMUSG00000022427 | *Tomm22* | ENSMUSG00000055629 | *B4galnt4* |
| ENSMUSG00000022428 | *Cby1* | ENSMUSG00000055660 | *Mettl4* |
| ENSMUSG00000022429 | *Dmc1* | ENSMUSG00000055692 | *Tmem191c* |
| ENSMUSG00000022431 | *Ribc2* | ENSMUSG00000055760 | *Gemin6* |

| | | | |
|---|---|---|---|
| ENSMUSG00000022432 | *Smc1b* | ENSMUSG00000055762 | *Eef1d* |
| ENSMUSG00000022439 | *Parvg* | ENSMUSG00000055817 | *Mta3* |
| ENSMUSG00000022441 | *Efcab6* | ENSMUSG00000055826 | *Tescl* |
| ENSMUSG00000022443 | *Myh9* | ENSMUSG00000055850 | *Rnf181* |
| ENSMUSG00000022450 | *Ndufa6* | ENSMUSG00000055862 | *Izumo4* |
| ENSMUSG00000022452 | *Smdt1* | ENSMUSG00000055891 | *Ubl4b* |
| ENSMUSG00000022455 | *Wbp2nl* | ENSMUSG00000055897 | *Ppp4r1l-ps* |
| ENSMUSG00000022466 | *Rpap3* | ENSMUSG00000056018 | *Ccdc7b* |
| ENSMUSG00000022471 | *Xrcc6* | ENSMUSG00000056050 | *Mia3* |
| ENSMUSG00000022472 | *Desi1* | ENSMUSG00000056076 | *Eif3b* |
| ENSMUSG00000022501 | *Prm1* | ENSMUSG00000056131 | *Pgm3* |
| ENSMUSG00000022507 | *1810013L24Rik* | ENSMUSG00000056158 | *Car10* |
| ENSMUSG00000022518 | *4930562C15Rik* | ENSMUSG00000056185 | *Snx32* |
| ENSMUSG00000022521 | *Crebbp* | ENSMUSG00000056201 | *Cfl1* |
| ENSMUSG00000022529 | *Zfp263* | ENSMUSG00000056211 | *R3hdm1* |
| ENSMUSG00000022536 | *Glyr1* | ENSMUSG00000056216 | *Cebpg* |
| ENSMUSG00000022538 | *Lsg1* | ENSMUSG00000056260 | *Lrif1* |
| ENSMUSG00000022542 | *Septin12* | ENSMUSG00000056267 | *Cep70* |
| ENSMUSG00000022557 | *Bop1* | ENSMUSG00000056342 | *Usp34* |
| ENSMUSG00000022558 | *Mroh1* | ENSMUSG00000056394 | *Lig1* |
| ENSMUSG00000022564 | *Grina* | ENSMUSG00000056429 | *Tgoln1* |
| ENSMUSG00000022565 | *Plec* | ENSMUSG00000056436 | *Cyct* |
| ENSMUSG00000022568 | *Scrib* | ENSMUSG00000056476 | *Med12l* |
| ENSMUSG00000022579 | *Gpihbp1* | ENSMUSG00000056494 | *Cngb3* |
| ENSMUSG00000022601 | *Zbtb11* | ENSMUSG00000056508 | *1700001K19Rik* |
| ENSMUSG00000022602 | *Arc* | ENSMUSG00000056509 | *Gm9999* |
| ENSMUSG00000022603 | *Mroh4* | ENSMUSG00000056515 | *Rab31* |
| ENSMUSG00000022607 | *Ptk2* | ENSMUSG00000056531 | *Ccdc18* |
| ENSMUSG00000022619 | *Mapk8ip2* | ENSMUSG00000056536 | *Pign* |
| ENSMUSG00000022621 | *Rabl2* | ENSMUSG00000056553 | *Ptprn2* |
| ENSMUSG00000022622 | *Acr* | ENSMUSG00000056598 | *Drc3* |
| ENSMUSG00000022629 | *Kif21a* | ENSMUSG00000056608 | *Chd9* |
| ENSMUSG00000022635 | *Zcrb1* | ENSMUSG00000056612 | *Ppp1r14b* |
| ENSMUSG00000022641 | *Bbx* | ENSMUSG00000056617 | *4931429L15Rik* |
| ENSMUSG00000022656 | *Nectin3* | ENSMUSG00000056692 | *Ilrun* |
| ENSMUSG00000022663 | *Atg3* | ENSMUSG00000056735 | *A930024E05Rik* |
| ENSMUSG00000022678 | *Nde1* | ENSMUSG00000056763 | *Cspp1* |
| ENSMUSG00000022680 | *Pdxdc1* | ENSMUSG00000056770 | *Setd3* |
| ENSMUSG00000022685 | *Parn* | ENSMUSG00000056771 | *Gm10010* |
| ENSMUSG00000022701 | *Ccdc191* | ENSMUSG00000056772 | *Rps6-ps2* |
| ENSMUSG00000022710 | *Usp7* | ENSMUSG00000056815 | *Gm6812* |
| ENSMUSG00000022715 | *Tmem114* | ENSMUSG00000056832 | *Ttc26* |
| ENSMUSG00000022718 | *Dgcr8* | ENSMUSG00000056851 | *Pcbp2* |
| ENSMUSG00000022723 | *Crybg3* | ENSMUSG00000056877 | *Rps15a-ps1* |
| ENSMUSG00000022765 | *Snap29* | ENSMUSG00000056899 | *Immp2l* |
| ENSMUSG00000022768 | *Ccdc116* | ENSMUSG00000056912 | *1700017N19Rik* |
| ENSMUSG00000022770 | *Dlg1* | ENSMUSG00000056919 | *Cep162* |
| ENSMUSG00000022771 | *Ppil2* | ENSMUSG00000056952 | *Tatdn2* |
| ENSMUSG00000022774 | *Ncbp2* | ENSMUSG00000056962 | *Jmjd6* |
| ENSMUSG00000022787 | *Wdr53* | ENSMUSG00000056987 | *Garin2* |
| ENSMUSG00000022789 | *Dnm1l* | ENSMUSG00000056999 | *Ide* |
| ENSMUSG00000022790 | *Igsf11* | ENSMUSG00000057000 | *Nxf3* |
| ENSMUSG00000022791 | *Tnk2* | ENSMUSG00000057047 | *1700010B08Rik* |
| ENSMUSG00000022800 | *Fyttd1* | ENSMUSG00000057072 | *Spata45* |
| ENSMUSG00000022805 | *Cfap91* | ENSMUSG00000057110 | *Cntrl* |
| ENSMUSG00000022808 | *Snx4* | ENSMUSG00000057113 | *Npm1* |
| ENSMUSG00000022811 | *Zfp148* | ENSMUSG00000057116 | *Semp2l2a* |
| ENSMUSG00000022832 | *Ropn1* | ENSMUSG00000057132 | *Rpgrip1* |

| | | | |
|---|---|---|---|
| ENSMUSG00000022837 | *Iqcb1* | ENSMUSG00000057133 | *Chd6* |
| ENSMUSG00000022855 | *Senp2* | ENSMUSG00000057147 | *Dph6* |
| ENSMUSG00000022858 | *Tra2b* | ENSMUSG00000057156 | *Homez* |
| ENSMUSG00000022867 | *Usp25* | ENSMUSG00000057176 | *Ccdc189* |
| ENSMUSG00000022883 | *Robo1* | ENSMUSG00000057236 | *Rbbp4* |
| ENSMUSG00000022884 | *Eif4a2* | ENSMUSG00000057246 | *BC051142* |
| ENSMUSG00000022889 | *Mrpl39* | ENSMUSG00000057265 | *Bbof1* |
| ENSMUSG00000022892 | *App* | ENSMUSG00000057286 | *St6galnac2* |
| ENSMUSG00000022897 | *Dyrk1a* | ENSMUSG00000057322 | *Rpl38* |
| ENSMUSG00000022911 | *Arl13b* | ENSMUSG00000057335 | *Cep170* |
| ENSMUSG00000022914 | *Brwd1* | ENSMUSG00000057363 | *Uxs1* |
| ENSMUSG00000022957 | *Itsn1* | ENSMUSG00000057367 | *Birc2* |
| ENSMUSG00000022961 | *Son* | ENSMUSG00000057388 | *Mrpl18* |
| ENSMUSG00000022972 | *Cfap298* | ENSMUSG00000057406 | *Nsd2* |
| ENSMUSG00000022974 | *Paxbp1* | ENSMUSG00000057411 | *Antkmt* |
| ENSMUSG00000022982 | *Sod1* | ENSMUSG00000057421 | *Las1l* |
| ENSMUSG00000022995 | *Enah* | ENSMUSG00000057572 | *Zbtb8os* |
| ENSMUSG00000023007 | *Prpf40b* | ENSMUSG00000057596 | *Trim30d* |
| ENSMUSG00000023010 | *Tmbim6* | ENSMUSG00000057637 | *Prdm2* |
| ENSMUSG00000023027 | *Atf1* | ENSMUSG00000057649 | *Brd9* |
| ENSMUSG00000023032 | *Slc4a8* | ENSMUSG00000057738 | *Sptan1* |
| ENSMUSG00000023055 | *Calcoco1* | ENSMUSG00000057805 | *1700084M14Rik* |
| ENSMUSG00000023072 | *Cep89* | ENSMUSG00000057816 | *Cfap299* |
| ENSMUSG00000023084 | *Lrrc71* | ENSMUSG00000057858 | *Fam204a* |
| ENSMUSG00000023088 | *Abcc1* | ENSMUSG00000058006 | *Mdn1* |
| ENSMUSG00000023104 | *Rfc2* | ENSMUSG00000058046 | *4933430I17Rik* |
| ENSMUSG00000023106 | *Denr* | ENSMUSG00000058064 | *Gm10036* |
| ENSMUSG00000023118 | *Sympk* | ENSMUSG00000058126 | *Tpm3-rs7* |
| ENSMUSG00000023150 | *Ivns1abp* | ENSMUSG00000058135 | *Gstm1* |
| ENSMUSG00000023170 | *Gps2* | ENSMUSG00000058145 | *Adamts17* |
| ENSMUSG00000023175 | *Bsg* | ENSMUSG00000058153 | *Sez6l* |
| ENSMUSG00000023257 | *Cypt1* | ENSMUSG00000058183 | *Mmel1* |
| ENSMUSG00000023272 | *Creld2* | ENSMUSG00000058230 | *Arhgap35* |
| ENSMUSG00000023350 | *Phf8-ps* | ENSMUSG00000058297 | *Spock2* |
| ENSMUSG00000023456 | *Tpi1* | ENSMUSG00000058301 | *Upf1* |
| ENSMUSG00000023467 | *Tulp2* | ENSMUSG00000058318 | *Phf21a* |
| ENSMUSG00000023505 | *Cdca3* | ENSMUSG00000058325 | *Dock1* |
| ENSMUSG00000023805 | *Synj2* | ENSMUSG00000058388 | *Phtf1* |
| ENSMUSG00000023806 | *Rsph3b* | ENSMUSG00000058398 | *Prss43* |
| ENSMUSG00000023830 | *Igf2r* | ENSMUSG00000058546 | *Rpl23a* |
| ENSMUSG00000023852 | *Chd1* | ENSMUSG00000058558 | *Rpl5* |
| ENSMUSG00000023868 | *Pde10a* | ENSMUSG00000058567 | *Gm2531* |
| ENSMUSG00000023873 | *1700010I14Rik* | ENSMUSG00000058626 | *Capn11* |
| ENSMUSG00000023883 | *Phf10* | ENSMUSG00000058638 | *Zfp110* |
| ENSMUSG00000023919 | *Cenpq* | ENSMUSG00000058643 | *Speer4f1* |
| ENSMUSG00000023923 | *Tbc1d5* | ENSMUSG00000058655 | *Eif4b* |
| ENSMUSG00000023930 | *Crisp2* | ENSMUSG00000058670 | *Dmtf1l* |
| ENSMUSG00000023931 | *Efhb* | ENSMUSG00000058690 | *Ccser2* |
| ENSMUSG00000023932 | *Cdc5l* | ENSMUSG00000058709 | *Egln2* |
| ENSMUSG00000023938 | *Aars2* | ENSMUSG00000058761 | *Rnf169* |
| ENSMUSG00000023944 | *Hsp90ab1* | ENSMUSG00000058799 | *Nap1l1* |
| ENSMUSG00000023952 | *Gtpbp2* | ENSMUSG00000058816 | *Ppp1r2-ps3* |
| ENSMUSG00000023977 | *Ubr2* | ENSMUSG00000058835 | *Abi1* |
| ENSMUSG00000023984 | *Gm20517* | ENSMUSG00000058881 | *Zfp516* |
| ENSMUSG00000024002 | *Brd4* | ENSMUSG00000058925 | *Ccdc192* |
| ENSMUSG00000024006 | *Stk38* | ENSMUSG00000059005 | *Hnrnpa3* |
| ENSMUSG00000024033 | *Rsph1* | ENSMUSG00000059040 | *Eno1b* |
| ENSMUSG00000024044 | *Epb41l3* | ENSMUSG00000059089 | *Fcgr4* |

| | | | |
|---|---|---|---|
| ENSMUSG00000024045 | *Akap8* | ENSMUSG00000059114 | *Lrrc74a* |
| ENSMUSG00000024049 | *Myom1* | ENSMUSG00000059119 | *Nap1l4* |
| ENSMUSG00000024056 | *Ndc80* | ENSMUSG00000059150 | *Rpl26-ps4* |
| ENSMUSG00000024059 | *Clip4* | ENSMUSG00000059182 | *Skap2* |
| ENSMUSG00000024067 | *Dpy30* | ENSMUSG00000059195 | *Gm12715* |
| ENSMUSG00000024068 | *Spast* | ENSMUSG00000059208 | *Hnrnpm* |
| ENSMUSG00000024073 | *Birc6* | ENSMUSG00000059248 | *Septin9* |
| ENSMUSG00000024077 | *Strn* | ENSMUSG00000059263 | *Usp47* |
| ENSMUSG00000024079 | *Eif2ak2* | ENSMUSG00000059273 | *Zc3h4* |
| ENSMUSG00000024081 | *Cebpz* | ENSMUSG00000059278 | *Naa38* |
| ENSMUSG00000024083 | *Pja2* | ENSMUSG00000059288 | *Cdyl* |
| ENSMUSG00000024085 | *Man2a1* | ENSMUSG00000059323 | *Tonsl* |
| ENSMUSG00000024091 | *Vapa* | ENSMUSG00000059395 | *Nkapl* |
| ENSMUSG00000024095 | *Hnrnpll* | ENSMUSG00000059474 | *Mbtd1* |
| ENSMUSG00000024096 | *Ralbp1* | ENSMUSG00000059482 | *Cfap418* |
| ENSMUSG00000024097 | *Srsf7* | ENSMUSG00000059498 | *Fcgr3* |
| ENSMUSG00000024099 | *Ndufv2* | ENSMUSG00000059518 | *Znhit1* |
| ENSMUSG00000024104 | *Washc2* | ENSMUSG00000059540 | *Tcea2* |
| ENSMUSG00000024109 | *Nrxn1* | ENSMUSG00000059554 | *Ccdc28a* |
| ENSMUSG00000024154 | *Gtf2a1l* | ENSMUSG00000059645 | *Gm7361* |
| ENSMUSG00000024158 | *Hagh* | ENSMUSG00000059659 | *Gm10069* |
| ENSMUSG00000024163 | *Mapk8ip3* | ENSMUSG00000059690 | *1700020N15Rik* |
| ENSMUSG00000024169 | *Ift140* | ENSMUSG00000059708 | *Akap17b* |
| ENSMUSG00000024175 | *Tekt4* | ENSMUSG00000059751 | *Rps3a3* |
| ENSMUSG00000024182 | *Axin1* | ENSMUSG00000059810 | *Rgs3* |
| ENSMUSG00000024187 | *Fam234a* | ENSMUSG00000059834 | *Sclt1* |
| ENSMUSG00000024188 | *Luc7l* | ENSMUSG00000059854 | *Hydin* |
| ENSMUSG00000024193 | *Phf1* | ENSMUSG00000059891 | *Tsks* |
| ENSMUSG00000024209 | *Acsbg3* | ENSMUSG00000059920 | *4930453N24Rik* |
| ENSMUSG00000024213 | *Nudt3* | ENSMUSG00000059921 | *Unc5c* |
| ENSMUSG00000024218 | *Taf11* | ENSMUSG00000059939 | *9430015G10Rik* |
| ENSMUSG00000024222 | *Fkbp5* | ENSMUSG00000059970 | *Hspa2* |
| ENSMUSG00000024236 | *Svil* | ENSMUSG00000059974 | *Ntm* |
| ENSMUSG00000024238 | *Zeb1* | ENSMUSG00000059981 | *Taok2* |
| ENSMUSG00000024242 | *Map4k3* | ENSMUSG00000060036 | *Rpl3* |
| ENSMUSG00000024253 | *Dync2li1* | ENSMUSG00000060126 | *Tpt1* |
| ENSMUSG00000024259 | *Slc25a46* | ENSMUSG00000060128 | *Gm2710* |
| ENSMUSG00000024260 | *Sap130* | ENSMUSG00000060143 | *Gm10076* |
| ENSMUSG00000024270 | *Slc39a6* | ENSMUSG00000060147 | *Serpinb6a* |
| ENSMUSG00000024277 | *Mapre2* | ENSMUSG00000060176 | *Kif27* |
| ENSMUSG00000024286 | *Ccny* | ENSMUSG00000060206 | *Zfp462* |
| ENSMUSG00000024287 | *Thoc1* | ENSMUSG00000060227 | *Golm2* |
| ENSMUSG00000024290 | *Rock1* | ENSMUSG00000060260 | *Pwwp2b* |
| ENSMUSG00000024294 | *Mib1* | ENSMUSG00000060261 | *Gtf2i* |
| ENSMUSG00000024306 | *Ccdc178* | ENSMUSG00000060268 | *Armh1* |
| ENSMUSG00000024309 | *Pfdn6* | ENSMUSG00000060279 | *Ap2a1* |
| ENSMUSG00000024312 | *Wdr46* | ENSMUSG00000060373 | *Hnrnpc* |
| ENSMUSG00000024317 | *Rnf138* | ENSMUSG00000060407 | *Cyp2a12* |
| ENSMUSG00000024335 | *Brd2* | ENSMUSG00000060445 | *Sycp2* |
| ENSMUSG00000024346 | *Pfdn1* | ENSMUSG00000060450 | *Rnf14* |
| ENSMUSG00000024350 | *Dnajc18* | ENSMUSG00000060467 | *Gm10080* |
| ENSMUSG00000024352 | *Spata24* | ENSMUSG00000060475 | *Wtap* |
| ENSMUSG00000024357 | *Sil1* | ENSMUSG00000060538 | *Tmem219* |
| ENSMUSG00000024359 | *Hspa9* | ENSMUSG00000060579 | *Fhit* |
| ENSMUSG00000024360 | *Etf1* | ENSMUSG00000060600 | *Eno3* |
| ENSMUSG00000024369 | *Nelfe* | ENSMUSG00000060679 | *Mrps9* |
| ENSMUSG00000024382 | *Ercc3* | ENSMUSG00000060708 | *Bloc1s4* |
| ENSMUSG00000024384 | *Iws1* | ENSMUSG00000060715 | *1700019A02Rik* |

| | | | |
|---|---|---|---|
| ENSMUSG00000024387 | *Csnk2b* | ENSMUSG00000060726 | *Tmsb15a* |
| ENSMUSG00000024392 | *Bag6* | ENSMUSG00000060739 | *Nsa2* |
| ENSMUSG00000024393 | *Prrc2a* | ENSMUSG00000060771 | *Tsga10* |
| ENSMUSG00000024397 | *Aif1* | ENSMUSG00000060811 | *Gm10088* |
| ENSMUSG00000024400 | *Wdr33* | ENSMUSG00000060843 | *Ctnna3* |
| ENSMUSG00000024404 | *Riok3* | ENSMUSG00000060923 | *Acyp2* |
| ENSMUSG00000024422 | *Dhx16* | ENSMUSG00000060938 | *Rpl26* |
| ENSMUSG00000024425 | *Ndfip1* | ENSMUSG00000060985 | *Tdrd5* |
| ENSMUSG00000024429 | *Gnl1* | ENSMUSG00000060989 | *Gm11847* |
| ENSMUSG00000024430 | *Cabyr* | ENSMUSG00000061024 | *Rrs1* |
| ENSMUSG00000024456 | *Diaph1* | ENSMUSG00000061028 | *Clasrp* |
| ENSMUSG00000024457 | *Trim26* | ENSMUSG00000061032 | *Rrp1* |
| ENSMUSG00000024474 | *Ik* | ENSMUSG00000061046 | *Haghl* |
| ENSMUSG00000024483 | *Ankhd1* | ENSMUSG00000061062 | *Hdac1-ps* |
| ENSMUSG00000024487 | *Yipf5* | ENSMUSG00000061099 | *Gapdhs* |
| ENSMUSG00000024491 | *Rbm27* | ENSMUSG00000061104 | *Sap18b* |
| ENSMUSG00000024498 | *Tcerg1* | ENSMUSG00000061130 | *Ppm1b* |
| ENSMUSG00000024501 | *Dpysl3* | ENSMUSG00000061136 | *Prpf40a* |
| ENSMUSG00000024507 | *Hsd17b4* | ENSMUSG00000061186 | *Sfmbt2* |
| ENSMUSG00000024513 | *Mbd2* | ENSMUSG00000061244 | *Exoc5* |
| ENSMUSG00000024527 | *Afg3l2* | ENSMUSG00000061288 | *Taok3* |
| ENSMUSG00000024532 | *1700034E13Rik* | ENSMUSG00000061298 | *Agbl4* |
| ENSMUSG00000024539 | *Ptpn2* | ENSMUSG00000061306 | *Slc38a10* |
| ENSMUSG00000024542 | *Cep192* | ENSMUSG00000061322 | *Dnai1* |
| ENSMUSG00000024560 | *Cxxc1* | ENSMUSG00000061411 | *Nol4l* |
| ENSMUSG00000024566 | *Atp9b* | ENSMUSG00000061458 | *Nol10* |
| ENSMUSG00000024576 | *Csnk1a1* | ENSMUSG00000061518 | *Cox5b* |
| ENSMUSG00000024583 | *Txnl1* | ENSMUSG00000061525 | *4921509C19Rik* |
| ENSMUSG00000024587 | *Nars* | ENSMUSG00000061533 | *Cep128* |
| ENSMUSG00000024590 | *Lmnb1* | ENSMUSG00000061589 | *Dot1l* |
| ENSMUSG00000024608 | *Rps14* | ENSMUSG00000061607 | *Mdc1* |
| ENSMUSG00000024614 | *Tmx3* | ENSMUSG00000061633 | *Ndufb11b* |
| ENSMUSG00000024654 | *Asrgl1* | ENSMUSG00000061650 | *Med9* |
| ENSMUSG00000024658 | *Gm9750* | ENSMUSG00000061665 | *Cd2ap* |
| ENSMUSG00000024660 | *Incenp* | ENSMUSG00000061666 | *Gdpd1* |
| ENSMUSG00000024661 | *Fth1* | ENSMUSG00000061684 | *Rpl21-ps8* |
| ENSMUSG00000024663 | *Rab3il1* | ENSMUSG00000061689 | *Dlgap4* |
| ENSMUSG00000024687 | *Osbp* | ENSMUSG00000061751 | *Kalrn* |
| ENSMUSG00000024695 | *Zfp91* | ENSMUSG00000061755 | *Bod1l* |
| ENSMUSG00000024740 | *Ddb1* | ENSMUSG00000061759 | *Armt1* |
| ENSMUSG00000024749 | *Tmc1* | ENSMUSG00000061787 | *Rps17* |
| ENSMUSG00000024759 | *Atl3* | ENSMUSG00000061802 | *Odad2* |
| ENSMUSG00000024772 | *Ehd1* | ENSMUSG00000061848 | *Gm5805* |
| ENSMUSG00000024773 | *Atg2a* | ENSMUSG00000061859 | *Patj* |
| ENSMUSG00000024780 | *Cdc37l1* | ENSMUSG00000061882 | *Ccdc62* |
| ENSMUSG00000024786 | *Majin* | ENSMUSG00000061923 | *Odf1* |
| ENSMUSG00000024795 | *Kif20b* | ENSMUSG00000061928 | *Dsg1b* |
| ENSMUSG00000024805 | *Pcgf5* | ENSMUSG00000061950 | *Ppp4r1* |
| ENSMUSG00000024812 | *Tjp2* | ENSMUSG00000061983 | *Rps12* |
| ENSMUSG00000024841 | *Eif1ad* | ENSMUSG00000062014 | *Gmfb* |
| ENSMUSG00000024843 | *Chka* | ENSMUSG00000062028 | *Irgc1* |
| ENSMUSG00000024853 | *Sf3b2* | ENSMUSG00000062075 | *Lmnb2* |
| ENSMUSG00000024862 | *Klc2* | ENSMUSG00000062078 | *Qki* |
| ENSMUSG00000024878 | *Cbwd1* | ENSMUSG00000062083 | *Rpl13a-ps1* |
| ENSMUSG00000024885 | *Aldh3b1* | ENSMUSG00000062154 | *Tex33* |
| ENSMUSG00000024897 | *Apba1* | ENSMUSG00000062190 | *Lancl2* |
| ENSMUSG00000024900 | *Cpt1a* | ENSMUSG00000062202 | *Btbd9* |
| ENSMUSG00000024908 | *Ppp6r3* | ENSMUSG00000062203 | *Gspt1* |

| | | | |
|---|---|---|---|
| ENSMUSG00000024913 | *Lrp5* | ENSMUSG00000062209 | *Erbb4* |
| ENSMUSG00000024914 | *Drap1* | ENSMUSG00000062224 | *4933411G06Rik* |
| ENSMUSG00000024921 | *Smarca2* | ENSMUSG00000062234 | *Gak* |
| ENSMUSG00000024927 | *Rela* | ENSMUSG00000062270 | *Morf4l1* |
| ENSMUSG00000024937 | *Ehbp1l1* | ENSMUSG00000062296 | *Trank1* |
| ENSMUSG00000024943 | *Smc5* | ENSMUSG00000062328 | *Rpl17* |
| ENSMUSG00000024947 | *Men1* | ENSMUSG00000062353 | *Gm15772* |
| ENSMUSG00000024949 | *Sf1* | ENSMUSG00000062438 | *Adam1b* |
| ENSMUSG00000024957 | *Kcnk4* | ENSMUSG00000062554 | *Gm12751* |
| ENSMUSG00000024958 | *Gpr137* | ENSMUSG00000062588 | *Gm6104* |
| ENSMUSG00000024959 | *Bad* | ENSMUSG00000062604 | *Srpk2* |
| ENSMUSG00000024960 | *Plcb3* | ENSMUSG00000062611 | *Rps3a2* |
| ENSMUSG00000024963 | *Dnajc4* | ENSMUSG00000062627 | *Mysm1* |
| ENSMUSG00000024966 | *Stip1* | ENSMUSG00000062732 | *Lypd4* |
| ENSMUSG00000024969 | *Mark2* | ENSMUSG00000062761 | *Zfp512* |
| ENSMUSG00000024973 | *Plaat5* | ENSMUSG00000062846 | *Gm14176* |
| ENSMUSG00000024974 | *Smc3* | ENSMUSG00000062859 | *Tcp11* |
| ENSMUSG00000024975 | *Pdcd4* | ENSMUSG00000062866 | *Phactr2* |
| ENSMUSG00000024983 | *Vti1a* | ENSMUSG00000062939 | *Stat4* |
| ENSMUSG00000024989 | *Cep55* | ENSMUSG00000062949 | *Atp11c* |
| ENSMUSG00000024991 | *Eif3a* | ENSMUSG00000062980 | *Cped1* |
| ENSMUSG00000024999 | *Noc3l* | ENSMUSG00000062995 | *Ica1* |
| ENSMUSG00000025001 | *Hells* | ENSMUSG00000063063 | *Ctnna2* |
| ENSMUSG00000025016 | *Tm9sf3* | ENSMUSG00000063077 | *Kif1b* |
| ENSMUSG00000025019 | *Lcor* | ENSMUSG00000063129 | *Aldoart2* |
| ENSMUSG00000025025 | *Mxi1* | ENSMUSG00000063163 | *Speer2* |
| ENSMUSG00000025027 | *Xpnpep1* | ENSMUSG00000063200 | *Nol7* |
| ENSMUSG00000025035 | *Arl3* | ENSMUSG00000063229 | *Ldha* |
| ENSMUSG00000025041 | *Nt5c2* | ENSMUSG00000063273 | *Naa15* |
| ENSMUSG00000025044 | *Msr1* | ENSMUSG00000063281 | *Zfp35* |
| ENSMUSG00000025047 | *Pdcd11* | ENSMUSG00000063328 | *Gm9396* |
| ENSMUSG00000025049 | *Taf5* | ENSMUSG00000063334 | *Krr1* |
| ENSMUSG00000025050 | *Pcgf6* | ENSMUSG00000063358 | *Mapk1* |
| ENSMUSG00000025060 | *Slk* | ENSMUSG00000063409 | *Lrrc43* |
| ENSMUSG00000025066 | *Sfr1* | ENSMUSG00000063410 | *Stk24* |
| ENSMUSG00000025081 | *Tdrd1* | ENSMUSG00000063450 | *Syne2* |
| ENSMUSG00000025103 | *Btbd1* | ENSMUSG00000063511 | *Snrnp70* |
| ENSMUSG00000025104 | *Hdgfl3* | ENSMUSG00000063524 | *Eno1* |
| ENSMUSG00000025130 | *P4hb* | ENSMUSG00000063550 | *Nup98* |
| ENSMUSG00000025132 | *Arhgdia* | ENSMUSG00000063568 | *Jazf1* |
| ENSMUSG00000025137 | *Pcyt2* | ENSMUSG00000063646 | *Jakmip1* |
| ENSMUSG00000025151 | *Maged1* | ENSMUSG00000063652 | *Slc22a21* |
| ENSMUSG00000025155 | *Dus1l* | ENSMUSG00000063663 | *Brwd3* |
| ENSMUSG00000025156 | *Gps1* | ENSMUSG00000063754 | *Gm10136* |
| ENSMUSG00000025171 | *Ubtd1* | ENSMUSG00000063800 | *Prpf38a* |
| ENSMUSG00000025176 | *Hoga1* | ENSMUSG00000063804 | *Lin28b* |
| ENSMUSG00000025203 | *Scd2* | ENSMUSG00000063810 | *Alms1* |
| ENSMUSG00000025204 | *Ndufb8* | ENSMUSG00000063820 | *Arl9* |
| ENSMUSG00000025220 | *Oga* | ENSMUSG00000063870 | *Chd4* |
| ENSMUSG00000025228 | *Actr1a* | ENSMUSG00000063873 | *Slc24a3* |
| ENSMUSG00000025231 | *Sufu* | ENSMUSG00000063882 | *Uqcrh* |
| ENSMUSG00000025234 | *Arih1* | ENSMUSG00000063889 | *Crem* |
| ENSMUSG00000025241 | *Fyco1* | ENSMUSG00000063902 | *Gm7964* |
| ENSMUSG00000025261 | *Huwe1* | ENSMUSG00000063932 | *Poteg* |
| ENSMUSG00000025288 | *4933436I01Rik* | ENSMUSG00000063952 | *Brpf3* |
| ENSMUSG00000025289 | *Prdx4* | ENSMUSG00000063972 | *Nr6a1* |
| ENSMUSG00000025290 | *Rps24* | ENSMUSG00000064061 | *Dzip3* |
| ENSMUSG00000025326 | *Ube3a* | ENSMUSG00000064063 | *BC048507* |

| | | | |
|---|---|---|---|
| ENSMUSG00000025337 | *Sbds* | ENSMUSG00000064068 | *Mtx1* |
| ENSMUSG00000025357 | *Dgka* | ENSMUSG00000064128 | *Cenpj* |
| ENSMUSG00000025358 | *Cdk2* | ENSMUSG00000064137 | *Rhox8* |
| ENSMUSG00000025362 | *Rps26* | ENSMUSG00000064138 | *Fam172a* |
| ENSMUSG00000025364 | *Pa2g4* | ENSMUSG00000064145 | *Arih2* |
| ENSMUSG00000025369 | *Smarcc2* | ENSMUSG00000064181 | *Rab3ip* |
| ENSMUSG00000025371 | *Chmp6* | ENSMUSG00000064193 | *Gm4735* |
| ENSMUSG00000025372 | *Baiap2* | ENSMUSG00000064202 | *Spata6l* |
| ENSMUSG00000025393 | *Atp5b* | ENSMUSG00000064280 | *Ccdc146* |
| ENSMUSG00000025404 | *R3hdm2* | ENSMUSG00000064281 | *Rpl19-ps1* |
| ENSMUSG00000025408 | *Ddit3* | ENSMUSG00000064294 | *Aox3* |
| ENSMUSG00000025410 | *Dctn2* | ENSMUSG00000064299 | *4921528I07Rik* |
| ENSMUSG00000025413 | *Ttc4* | ENSMUSG00000064302 | *Clasp1* |
| ENSMUSG00000025420 | *Katnal2* | ENSMUSG00000064307 | *Lrrc51* |
| ENSMUSG00000025428 | *Atp5a1* | ENSMUSG00000064339 | *mt-Rnr2* |
| ENSMUSG00000025437 | *Usp33* | ENSMUSG00000064341 | *mt-Nd1* |
| ENSMUSG00000025439 | *Clns1a* | ENSMUSG00000064344 | *mt-Tm* |
| ENSMUSG00000025451 | *Paip1* | ENSMUSG00000064345 | *mt-Nd2* |
| ENSMUSG00000025453 | *Nnt* | ENSMUSG00000064346 | *mt-Tw* |
| ENSMUSG00000025474 | *Tubgcp2* | ENSMUSG00000064351 | *mt-Co1* |
| ENSMUSG00000025477 | *Inpp5a* | ENSMUSG00000064354 | *mt-Co2* |
| ENSMUSG00000025480 | *Syce1* | ENSMUSG00000064367 | *mt-Nd5* |
| ENSMUSG00000025487 | *Psmd13* | ENSMUSG00000064368 | *mt-Nd6* |
| ENSMUSG00000025508 | *Rplp2* | ENSMUSG00000064370 | *mt-Cytb* |
| ENSMUSG00000025532 | *Crcp* | ENSMUSG00000064372 | *mt-Tp* |
| ENSMUSG00000025571 | *Tnrc6c* | ENSMUSG00000064682 | *Gm25813* |
| ENSMUSG00000025584 | *Pde8a* | ENSMUSG00000064694 | *Gm24146* |
| ENSMUSG00000025592 | *Dach2* | ENSMUSG00000064696 | *None* |
| ENSMUSG00000025607 | *Copg2* | ENSMUSG00000064702 | *Gm24950* |
| ENSMUSG00000025613 | *Cct8* | ENSMUSG00000064772 | *Gm25202* |
| ENSMUSG00000025616 | *Usp16* | ENSMUSG00000064844 | *Gm26202* |
| ENSMUSG00000025626 | *Phf6* | ENSMUSG00000064856 | *None* |
| ENSMUSG00000025651 | *Uqcrc1* | ENSMUSG00000064871 | *Snord58b* |
| ENSMUSG00000025745 | *Hadha* | ENSMUSG00000064923 | *Gm22042* |
| ENSMUSG00000025757 | *Hspa4l* | ENSMUSG00000064945 | *Rny3* |
| ENSMUSG00000025762 | *Larp1b* | ENSMUSG00000064968 | *Snord47* |
| ENSMUSG00000025782 | *Taf3* | ENSMUSG00000064984 | *Snord73a* |
| ENSMUSG00000025783 | *4930412O13Rik* | ENSMUSG00000064999 | *None* |
| ENSMUSG00000025808 | *Ccdc7a* | ENSMUSG00000065037 | *Rn7sk* |
| ENSMUSG00000025823 | *Pdia4* | ENSMUSG00000065041 | *Gm24044* |
| ENSMUSG00000025862 | *Stag2* | ENSMUSG00000065087 | *Snord22* |
| ENSMUSG00000025873 | *Faf2* | ENSMUSG00000065176 | *Rnu12* |
| ENSMUSG00000025878 | *Uimc1* | ENSMUSG00000065232 | *Gm22973* |
| ENSMUSG00000025885 | *Myo5b* | ENSMUSG00000065273 | *Gm25128* |
| ENSMUSG00000025894 | *Aasdhppt* | ENSMUSG00000065289 | *Gm23650* |
| ENSMUSG00000025898 | *Cwf19l2* | ENSMUSG00000065327 | *Gm22058* |
| ENSMUSG00000025907 | *Rb1cc1* | ENSMUSG00000065622 | *Gm24763* |
| ENSMUSG00000025912 | *Mybl1* | ENSMUSG00000065645 | *None* |
| ENSMUSG00000025916 | *Ppp1r42* | ENSMUSG00000065752 | *Gm23344* |
| ENSMUSG00000025925 | *Terf1* | ENSMUSG00000065767 | *Gm23849* |
| ENSMUSG00000025950 | *Idh1* | ENSMUSG00000065773 | *None* |
| ENSMUSG00000025955 | *Akr1cl* | ENSMUSG00000065808 | *Gm23037* |
| ENSMUSG00000025968 | *Ndufs1* | ENSMUSG00000065820 | *Gm26316* |
| ENSMUSG00000025977 | *Boll* | ENSMUSG00000065824 | *None* |
| ENSMUSG00000025982 | *Sf3b1* | ENSMUSG00000065853 | *Gm22354* |
| ENSMUSG00000025983 | *Ccdc150* | ENSMUSG00000065870 | *None* |
| ENSMUSG00000025986 | *Slc39a10* | ENSMUSG00000065878 | *Snord34* |
| ENSMUSG00000026018 | *Ica1l* | ENSMUSG00000065904 | *Gm26109* |

| | | | |
|---|---|---|---|
| ENSMUSG00000026020 | *Nop58* | ENSMUSG00000065905 | *Gm26110* |
| ENSMUSG00000026021 | *Sumo1* | ENSMUSG00000065911 | *Gm24447* |
| ENSMUSG00000026034 | *Clk1* | ENSMUSG00000065944 | *Rnu2-10* |
| ENSMUSG00000026049 | *Tex30* | ENSMUSG00000065954 | *Tacc1* |
| ENSMUSG00000026062 | *Slc9a2* | ENSMUSG00000066036 | *Ubr4* |
| ENSMUSG00000026074 | *Map4k4* | ENSMUSG00000066037 | *Hnrnpr* |
| ENSMUSG00000026078 | *Pdcl3* | ENSMUSG00000066043 | *Phactr4* |
| ENSMUSG00000026082 | *Rev1* | ENSMUSG00000066107 | *Gm12666* |
| ENSMUSG00000026083 | *Eif5b* | ENSMUSG00000066232 | *Ipo7* |
| ENSMUSG00000026107 | *Nabp1* | ENSMUSG00000066306 | *Numa1* |
| ENSMUSG00000026110 | *Mgat4a* | ENSMUSG00000066315 | *Gm12918* |
| ENSMUSG00000026113 | *Inpp4a* | ENSMUSG00000066406 | *Akap13* |
| ENSMUSG00000026131 | *Dst* | ENSMUSG00000066500 | *Izumo2* |
| ENSMUSG00000026134 | *Prim2* | ENSMUSG00000066551 | *Hmgb1* |
| ENSMUSG00000026142 | *Rhbdd1* | ENSMUSG00000066554 | *Gm10167* |
| ENSMUSG00000026153 | *Fam135a* | ENSMUSG00000066592 | *Gm7589* |
| ENSMUSG00000026155 | *Smap1* | ENSMUSG00000066643 | *Wdr35* |
| ENSMUSG00000026171 | *Rnf25* | ENSMUSG00000066687 | *Plzf* |
| ENSMUSG00000026173 | *Plcd4* | ENSMUSG00000066894 | *Vsig10* |
| ENSMUSG00000026179 | *Pnkd* | ENSMUSG00000066900 | *Suds3* |
| ENSMUSG00000026182 | *Tnp1* | ENSMUSG00000067017 | *Capza1-ps1* |
| ENSMUSG00000026202 | *Tuba4a* | ENSMUSG00000067158 | *Col4a4* |
| ENSMUSG00000026203 | *Dnajb2* | ENSMUSG00000067274 | *Rplp0* |
| ENSMUSG00000026213 | *Stk11ip* | ENSMUSG00000067338 | *Tuba3b* |
| ENSMUSG00000026219 | *Trip12* | ENSMUSG00000067367 | *Lyar* |
| ENSMUSG00000026226 | *Spata3* | ENSMUSG00000067438 | *Hmx1* |
| ENSMUSG00000026229 | *Psmd1* | ENSMUSG00000067608 | *Pcna-ps2* |
| ENSMUSG00000026234 | *Ncl* | ENSMUSG00000067629 | *Syngap1* |
| ENSMUSG00000026238 | *Ptma* | ENSMUSG00000067702 | *Tuba3a* |
| ENSMUSG00000026254 | *Eif4e2* | ENSMUSG00000067773 | *Defb41* |
| ENSMUSG00000026255 | *Efhd1* | ENSMUSG00000067795 | *4930444P10Rik* |
| ENSMUSG00000026274 | *Pask* | ENSMUSG00000067818 | *Myl9* |
| ENSMUSG00000026275 | *Ppp1r7* | ENSMUSG00000067848 | *4933402N22Rik* |
| ENSMUSG00000026277 | *Stk25* | ENSMUSG00000067869 | *Tcea1-ps1* |
| ENSMUSG00000026289 | *Atg16l1* | ENSMUSG00000067873 | *Htatsf1* |
| ENSMUSG00000026301 | *Iqca* | ENSMUSG00000067878 | *Map7d3* |
| ENSMUSG00000026319 | *Relch* | ENSMUSG00000067942 | *Zfp160* |
| ENSMUSG00000026331 | *Slco6c1* | ENSMUSG00000067995 | *Gtf2f2* |
| ENSMUSG00000026335 | *Pam* | ENSMUSG00000068036 | *Afdn* |
| ENSMUSG00000026336 | *Slco6d1* | ENSMUSG00000068039 | *Tcp1* |
| ENSMUSG00000026339 | *Ccdc93* | ENSMUSG00000068040 | *Tm9sf4* |
| ENSMUSG00000026341 | *Actr3* | ENSMUSG00000068062 | *Gm14164* |
| ENSMUSG00000026353 | *Ubxn4* | ENSMUSG00000068115 | *Ninl* |
| ENSMUSG00000026356 | *Dars* | ENSMUSG00000068165 | *Gm10233* |
| ENSMUSG00000026361 | *Cdc73* | ENSMUSG00000068167 | *Csnka2ip* |
| ENSMUSG00000026385 | *Dbi* | ENSMUSG00000068184 | *Ndufaf2* |
| ENSMUSG00000026401 | *Cd55b* | ENSMUSG00000068200 | *Gm5931* |
| ENSMUSG00000026409 | *Pfkfb2* | ENSMUSG00000068205 | *Macrod2* |
| ENSMUSG00000026414 | *Tnnt2* | ENSMUSG00000068290 | *Ddrgk1* |
| ENSMUSG00000026425 | *Srgap2* | ENSMUSG00000068328 | *Aup1* |
| ENSMUSG00000026429 | *Ube2t* | ENSMUSG00000068391 | *Chrac1* |
| ENSMUSG00000026434 | *Nucks1* | ENSMUSG00000068394 | *Cep152* |
| ENSMUSG00000026463 | *Atp2b4* | ENSMUSG00000068479 | *Mfap1a* |
| ENSMUSG00000026466 | *Tor1aip1* | ENSMUSG00000068522 | *Aard* |
| ENSMUSG00000026469 | *Xpr1* | ENSMUSG00000068747 | *Sort1* |
| ENSMUSG00000026470 | *Stx6* | ENSMUSG00000068794 | *Col28a1* |
| ENSMUSG00000026482 | *Rgl1* | ENSMUSG00000068823 | *Csde1* |
| ENSMUSG00000026484 | *Rnf2* | ENSMUSG00000068860 | *Gm128* |

| | | | |
|---|---|---|---|
| ENSMUSG00000026491 | *Ahctf1* | ENSMUSG00000068876 | *Cgn* |
| ENSMUSG00000026495 | *Efcab2* | ENSMUSG00000068882 | *Ssb* |
| ENSMUSG00000026496 | *Parp1* | ENSMUSG00000068917 | *Clk2* |
| ENSMUSG00000026499 | *Acbd3* | ENSMUSG00000068922 | *Msto1* |
| ENSMUSG00000026504 | *Sdccag8* | ENSMUSG00000069011 | *Gm10254* |
| ENSMUSG00000026523 | *Wdr64* | ENSMUSG00000069014 | *Gm5641* |
| ENSMUSG00000026532 | *Spta1* | ENSMUSG00000069045 | *Ddx3y* |
| ENSMUSG00000026546 | *Cfap45* | ENSMUSG00000069089 | *Cdk7* |
| ENSMUSG00000026553 | *Copa* | ENSMUSG00000069114 | *Zbtb10* |
| ENSMUSG00000026554 | *Dcaf8* | ENSMUSG00000069125 | *Rps24-ps2* |
| ENSMUSG00000026567 | *Adcy10* | ENSMUSG00000069135 | *Cep43* |
| ENSMUSG00000026571 | *Dcaf6* | ENSMUSG00000069306 | *H4c17* |
| ENSMUSG00000026572 | *Tbx19* | ENSMUSG00000069324 | *Gm5096* |
| ENSMUSG00000026576 | *Atp1b1* | ENSMUSG00000069355 | *Gm5152* |
| ENSMUSG00000026577 | *Blzf1* | ENSMUSG00000069441 | *Dsg1a* |
| ENSMUSG00000026578 | *Ccdc181* | ENSMUSG00000069565 | *Dazap1* |
| ENSMUSG00000026585 | *Kifap3* | ENSMUSG00000069601 | *Ank3* |
| ENSMUSG00000026592 | *Tex35* | ENSMUSG00000069622 | *Gm10273* |
| ENSMUSG00000026601 | *Axdnd1* | ENSMUSG00000069631 | *Strada* |
| ENSMUSG00000026605 | *Cenpf* | ENSMUSG00000069720 | *4930572O03Rik* |
| ENSMUSG00000026615 | *Eprs* | ENSMUSG00000069733 | *Ube2u* |
| ENSMUSG00000026622 | *Nek2* | ENSMUSG00000069805 | *Fbp1* |
| ENSMUSG00000026623 | *Lpgat1* | ENSMUSG00000069833 | *Ahnak* |
| ENSMUSG00000026626 | *Ppp2r5a* | ENSMUSG00000069844 | *Sco1* |
| ENSMUSG00000026643 | *Nmt2* | ENSMUSG00000069855 | *Slc47a2* |
| ENSMUSG00000026650 | *Meig1* | ENSMUSG00000069925 | *4932416K20Rik* |
| ENSMUSG00000026655 | *Fam107b* | ENSMUSG00000069971 | *4933402J07Rik* |
| ENSMUSG00000026672 | *Optn* | ENSMUSG00000070002 | *Ell* |
| ENSMUSG00000026679 | *Enkur* | ENSMUSG00000070271 | *Gm13268* |
| ENSMUSG00000026683 | *Nuf2* | ENSMUSG00000070280 | *Slc22a14* |
| ENSMUSG00000026696 | *Vamp4* | ENSMUSG00000070283 | *Ndufaf3* |
| ENSMUSG00000026712 | *Mrc1* | ENSMUSG00000070315 | *4930581F22Rik* |
| ENSMUSG00000026721 | *Rabgap1l* | ENSMUSG00000070319 | *Eif3g* |
| ENSMUSG00000026728 | *Vim* | ENSMUSG00000070327 | *Rnf213* |
| ENSMUSG00000026734 | *4921504E06Rik* | ENSMUSG00000070331 | *Qrich2* |
| ENSMUSG00000026739 | *Bmi1* | ENSMUSG00000070332 | *Trim80* |
| ENSMUSG00000026740 | *Dnajc1* | ENSMUSG00000070343 | *Gm10288* |
| ENSMUSG00000026743 | *Mllt10* | ENSMUSG00000070388 | *Fbxo39* |
| ENSMUSG00000026754 | *Golga1* | ENSMUSG00000070425 | *Xntrpc* |
| ENSMUSG00000026755 | *Arpc5l* | ENSMUSG00000070471 | *Erich6* |
| ENSMUSG00000026761 | *Orc4* | ENSMUSG00000070544 | *Top1* |
| ENSMUSG00000026764 | *Kif5c* | ENSMUSG00000070565 | *Rasal2* |
| ENSMUSG00000026774 | *Potegl* | ENSMUSG00000070568 | *Slc6a21* |
| ENSMUSG00000026775 | *Yme1l1* | ENSMUSG00000070639 | *Lrrc8b* |
| ENSMUSG00000026781 | *Acbd5* | ENSMUSG00000070697 | *Utp3* |
| ENSMUSG00000026782 | *Abi2* | ENSMUSG00000070708 | *Gtsf1l* |
| ENSMUSG00000026790 | *Odf2* | ENSMUSG00000070729 | *Gm12966* |
| ENSMUSG00000026803 | *Ttf1* | ENSMUSG00000070732 | *Rbm44* |
| ENSMUSG00000026807 | *Ak8* | ENSMUSG00000070738 | *Dgkd* |
| ENSMUSG00000026812 | *Tsc1* | ENSMUSG00000070802 | *Pnma8b* |
| ENSMUSG00000026817 | *Ak1* | ENSMUSG00000070806 | *Zmynd12* |
| ENSMUSG00000026819 | *Slc25a25* | ENSMUSG00000070883 | *Ccdc173* |
| ENSMUSG00000026821 | *Ralgds* | ENSMUSG00000070933 | *Speer4d* |
| ENSMUSG00000026822 | *Lcn2* | ENSMUSG00000070980 | *Actl7b* |
| ENSMUSG00000026827 | *Gpd2* | ENSMUSG00000071015 | *Gm136* |
| ENSMUSG00000026837 | *Col5a1* | ENSMUSG00000071054 | *Safb* |
| ENSMUSG00000026851 | *BC005624* | ENSMUSG00000071072 | *Ptges3* |
| ENSMUSG00000026853 | *Crat* | ENSMUSG00000071103 | *1700029J07Rik* |

| | | | |
|---|---|---|---|
| ENSMUSG00000026864 | *Hspa5* | ENSMUSG00000071104 | *Ccdc110* |
| ENSMUSG00000026867 | *Gapvd1* | ENSMUSG00000071138 | *Tex24* |
| ENSMUSG00000026872 | *Zeb2* | ENSMUSG00000071176 | *Arhgef10* |
| ENSMUSG00000026883 | *Dab2ip* | ENSMUSG00000071226 | *Cecr2* |
| ENSMUSG00000026894 | *Morn5* | ENSMUSG00000071252 | *2210408I21Rik* |
| ENSMUSG00000026914 | *Psmd14* | ENSMUSG00000071322 | *Tcp10a* |
| ENSMUSG00000026915 | *Strbp* | ENSMUSG00000071359 | *Tbpl1* |
| ENSMUSG00000026918 | *Brd3* | ENSMUSG00000071454 | *Dtnb* |
| ENSMUSG00000026926 | *Pmpca* | ENSMUSG00000071533 | *Pcnp* |
| ENSMUSG00000026927 | *Entr1* | ENSMUSG00000071550 | *Cfap44* |
| ENSMUSG00000026931 | *1700019N19Rik* | ENSMUSG00000071636 | *Rimbp3* |
| ENSMUSG00000026933 | *Camsap1* | ENSMUSG00000071644 | *Eef1g* |
| ENSMUSG00000026940 | *Ccdc183* | ENSMUSG00000071645 | *Tut1* |
| ENSMUSG00000026969 | *Fam166a* | ENSMUSG00000071650 | *Ganab* |
| ENSMUSG00000026970 | *Rbms1* | ENSMUSG00000071653 | *1810009A15Rik* |
| ENSMUSG00000026987 | *Baz2b* | ENSMUSG00000071655 | *Ubxn1* |
| ENSMUSG00000026994 | *Galnt3* | ENSMUSG00000071659 | *Hnrnpul2* |
| ENSMUSG00000027006 | *Dnajc10* | ENSMUSG00000071669 | *Snx29* |
| ENSMUSG00000027012 | *Dync1i2* | ENSMUSG00000071748 | *Styx-ps* |
| ENSMUSG00000027030 | *Stk39* | ENSMUSG00000071855 | *Ccdc112* |
| ENSMUSG00000027067 | *Ssrp1* | ENSMUSG00000071890 | *Mroh9* |
| ENSMUSG00000027071 | *P2rx3* | ENSMUSG00000072188 | *Gm10354* |
| ENSMUSG00000027077 | *Smtnl1* | ENSMUSG00000072214 | *Septin5* |
| ENSMUSG00000027080 | *Med19* | ENSMUSG00000072258 | *Taf1a* |
| ENSMUSG00000027086 | *Fastkd1* | ENSMUSG00000072295 | *C2cd6* |
| ENSMUSG00000027088 | *Phospho2* | ENSMUSG00000072487 | *Mroh5* |
| ENSMUSG00000027091 | *Zc3h15* | ENSMUSG00000072501 | *Phf20l1* |
| ENSMUSG00000027102 | *Hoxd8* | ENSMUSG00000072647 | *Adam1a* |
| ENSMUSG00000027108 | *Ola1* | ENSMUSG00000072663 | *Spef2* |
| ENSMUSG00000027130 | *Slc12a6* | ENSMUSG00000072692 | *Rpl37rt* |
| ENSMUSG00000027157 | *Potefam1* | ENSMUSG00000072694 | *1500011B03Rik* |
| ENSMUSG00000027160 | *Ccdc34* | ENSMUSG00000072720 | *Myo18b* |
| ENSMUSG00000027162 | *Lin7c* | ENSMUSG00000072722 | *Ccdc121rt2* |
| ENSMUSG00000027165 | *Iftap* | ENSMUSG00000072723 | *Gm10044* |
| ENSMUSG00000027176 | *Cstf3* | ENSMUSG00000072770 | *Acrbp* |
| ENSMUSG00000027180 | *Fbxo3* | ENSMUSG00000072878 | *1700123L14Rik* |
| ENSMUSG00000027184 | *Caprin1* | ENSMUSG00000072889 | *Nfxl1* |
| ENSMUSG00000027185 | *Nat10* | ENSMUSG00000072915 | *Gm12258* |
| ENSMUSG00000027189 | *Trim44* | ENSMUSG00000072952 | *Gm5878* |
| ENSMUSG00000027201 | *Myef2* | ENSMUSG00000072964 | *Bhlhb9* |
| ENSMUSG00000027209 | *Fam227b* | ENSMUSG00000072966 | *Gprasp2* |
| ENSMUSG00000027227 | *Sord* | ENSMUSG00000072974 | *Gm4787* |
| ENSMUSG00000027236 | *Eif3j1* | ENSMUSG00000073001 | *Cylc1* |
| ENSMUSG00000027245 | *Hypk* | ENSMUSG00000073079 | *Srp54a* |
| ENSMUSG00000027248 | *Pdia3* | ENSMUSG00000073102 | *Drc1* |
| ENSMUSG00000027276 | *Jag1* | ENSMUSG00000073117 | *Gm7347* |
| ENSMUSG00000027281 | *Slx4ip* | ENSMUSG00000073141 | *4930567H17Rik* |
| ENSMUSG00000027282 | *Mtch2* | ENSMUSG00000073208 | *Speer4c* |
| ENSMUSG00000027284 | *Cdan1* | ENSMUSG00000073293 | *Nudt10* |
| ENSMUSG00000027286 | *Lrrc57* | ENSMUSG00000073295 | *Nudt11* |
| ENSMUSG00000027287 | *Snap23* | ENSMUSG00000073430 | *Gm10505* |
| ENSMUSG00000027288 | *Zfp106* | ENSMUSG00000073471 | *Rsph3a* |
| ENSMUSG00000027304 | *Rtf1* | ENSMUSG00000073514 | *Dok6* |
| ENSMUSG00000027306 | *Nusap1* | ENSMUSG00000073551 | *Spink13* |
| ENSMUSG00000027309 | *Dnaaf9* | ENSMUSG00000073554 | *Gm6960* |
| ENSMUSG00000027326 | *Knl1* | ENSMUSG00000073557 | *Ppp1r12b* |
| ENSMUSG00000027331 | *Knstrn* | ENSMUSG00000073640 | *Rpl27-ps3* |
| ENSMUSG00000027344 | *Fsip1* | ENSMUSG00000073643 | *Wdfy1* |

| | | | |
|---|---|---|---|
| ENSMUSG00000027346 | *Gpcpd1* | ENSMUSG00000073647 | *Gm10557* |
| ENSMUSG00000027350 | *Chgb* | ENSMUSG00000073650 | *Catip* |
| ENSMUSG00000027355 | *Tmco5* | ENSMUSG00000073716 | *Gm13241* |
| ENSMUSG00000027363 | *Usp8* | ENSMUSG00000073722 | *4931408C20Rik* |
| ENSMUSG00000027365 | *Trpm7* | ENSMUSG00000073725 | *Lmbrd1* |
| ENSMUSG00000027366 | *Sppl2a* | ENSMUSG00000073730 | *Ppp1r14bl* |
| ENSMUSG00000027374 | *Mrps5* | ENSMUSG00000073758 | *Sh3d21* |
| ENSMUSG00000027378 | *Nphp1* | ENSMUSG00000073761 | *4933427I04Rik* |
| ENSMUSG00000027404 | *Snrpb* | ENSMUSG00000073888 | *Ccl27a* |
| ENSMUSG00000027405 | *Nop56* | ENSMUSG00000074030 | *Exoc8* |
| ENSMUSG00000027422 | *Rrbp1* | ENSMUSG00000074067 | *Gm10619* |
| ENSMUSG00000027433 | *Xrn2* | ENSMUSG00000074088 | *Snrnp40* |
| ENSMUSG00000027439 | *Gzf1* | ENSMUSG00000074093 | *Svip* |
| ENSMUSG00000027442 | *Cst8* | ENSMUSG00000074129 | *Rpl13a* |
| ENSMUSG00000027443 | *Cst12* | ENSMUSG00000074139 | *1700057G04Rik* |
| ENSMUSG00000027445 | *Cst9* | ENSMUSG00000074149 | *Gm10634* |
| ENSMUSG00000027447 | *Cst3* | ENSMUSG00000074182 | *Znhit6* |
| ENSMUSG00000027455 | *Nsfl1c* | ENSMUSG00000074212 | *Dnajb14* |
| ENSMUSG00000027469 | *Tpx2* | ENSMUSG00000074220 | *Zfp382* |
| ENSMUSG00000027475 | *Kif3b* | ENSMUSG00000074224 | *Wdr87-ps* |
| ENSMUSG00000027480 | *Sun5* | ENSMUSG00000074305 | *Peak1* |
| ENSMUSG00000027502 | *Rtf2* | ENSMUSG00000074340 | *Ovgp1* |
| ENSMUSG00000027517 | *Ankrd60* | ENSMUSG00000074404 | *Gm1698* |
| ENSMUSG00000027520 | *Zdbf2* | ENSMUSG00000074435 | *Smcp* |
| ENSMUSG00000027522 | *Stx16* | ENSMUSG00000074579 | *Lekr1* |
| ENSMUSG00000027525 | *Phactr3* | ENSMUSG00000074582 | *Arfgef2* |
| ENSMUSG00000027528 | *Fabp9* | ENSMUSG00000074589 | *4930449A18Rik* |
| ENSMUSG00000027531 | *Impa1* | ENSMUSG00000074619 | *1700034I23Rik* |
| ENSMUSG00000027550 | *Lrrcc1* | ENSMUSG00000074620 | *Gm10731* |
| ENSMUSG00000027562 | *Car2* | ENSMUSG00000074656 | *Eif2s2* |
| ENSMUSG00000027569 | *Mrgbp* | ENSMUSG00000074673 | *Ttll9* |
| ENSMUSG00000027575 | *Arfgap1* | ENSMUSG00000074734 | *Taf7l2* |
| ENSMUSG00000027582 | *Zgpat* | ENSMUSG00000074746 | *Pdzd8* |
| ENSMUSG00000027593 | *Raly* | ENSMUSG00000074748 | *Atxn7l3b* |
| ENSMUSG00000027596 | *a* | ENSMUSG00000074749 | *Kiz* |
| ENSMUSG00000027601 | *Mtfr1* | ENSMUSG00000074764 | *Sel1l2* |
| ENSMUSG00000027602 | *Map1lc3a* | ENSMUSG00000074771 | *Ankef1* |
| ENSMUSG00000027605 | *Acss2* | ENSMUSG00000074780 | *Anapc15-ps* |
| ENSMUSG00000027606 | *Dnajc5b* | ENSMUSG00000074785 | *Plxnc1* |
| ENSMUSG00000027620 | *Rbm39* | ENSMUSG00000074812 | *Spdye4c* |
| ENSMUSG00000027630 | *Tbl1xr1* | ENSMUSG00000074817 | *Papolb* |
| ENSMUSG00000027642 | *Rpn2* | ENSMUSG00000074884 | *Serf2* |
| ENSMUSG00000027649 | *Ctnnbl1* | ENSMUSG00000074903 | *Gm2058* |
| ENSMUSG00000027668 | *Mfn1* | ENSMUSG00000074978 | *Actg-ps1* |
| ENSMUSG00000027671 | *Actl6a* | ENSMUSG00000075053 | *Vdac3-ps1* |
| ENSMUSG00000027676 | *Ccdc39* | ENSMUSG00000075232 | *Amd1* |
| ENSMUSG00000027677 | *Ttc14* | ENSMUSG00000075249 | *Fsip2* |
| ENSMUSG00000027680 | *Fxr1* | ENSMUSG00000075268 | *Gm10819* |
| ENSMUSG00000027692 | *Tnik* | ENSMUSG00000075271 | *Ttc30a1* |
| ENSMUSG00000027694 | *Gm8325* | ENSMUSG00000075272 | *Ttc30a2* |
| ENSMUSG00000027699 | *Ect2* | ENSMUSG00000075302 | *Erich2* |
| ENSMUSG00000027703 | *Lrriq4* | ENSMUSG00000075376 | *Rc3h2* |
| ENSMUSG00000027706 | *Sec62* | ENSMUSG00000075415 | *Fnbp1* |
| ENSMUSG00000027708 | *Dcun1d1* | ENSMUSG00000075511 | *1700001L05Rik* |
| ENSMUSG00000027714 | *Exosc9* | ENSMUSG00000075524 | *4930407I10Rik* |
| ENSMUSG00000027719 | *Adad1* | ENSMUSG00000075528 | *Aarsd1* |
| ENSMUSG00000027722 | *Spata5* | ENSMUSG00000075569 | *Rsph10b* |
| ENSMUSG00000027751 | *Supt20* | ENSMUSG00000075581 | *Gm16409* |

| | | | |
|---|---|---|---|
| ENSMUSG00000027770 | *Dhx36* | ENSMUSG00000075590 | *Nrbp2* |
| ENSMUSG00000027778 | *Ift80* | ENSMUSG00000075595 | *Zfp652* |
| ENSMUSG00000027782 | *Kpna4* | ENSMUSG00000075701 | *Selenos* |
| ENSMUSG00000027799 | *Nbea* | ENSMUSG00000075706 | *Gpx4* |
| ENSMUSG00000027804 | *Ppid* | ENSMUSG00000075918 | *None* |
| ENSMUSG00000027823 | *Gmps* | ENSMUSG00000076433 | *Cep295nl* |
| ENSMUSG00000027849 | *Syt6* | ENSMUSG00000076436 | *Oxct2a* |
| ENSMUSG00000027855 | *Sycp1* | ENSMUSG00000077323 | *Rnu11* |
| ENSMUSG00000027867 | *Spag17* | ENSMUSG00000077457 | *Snord65* |
| ENSMUSG00000027871 | *Hsd3b1* | ENSMUSG00000077549 | *Snord71* |
| ENSMUSG00000027881 | *Prpf38b* | ENSMUSG00000077734 | *Snord83b* |
| ENSMUSG00000027886 | *Cfap276* | ENSMUSG00000078127 | *Fam170b* |
| ENSMUSG00000027893 | *Ahcyl1* | ENSMUSG00000078161 | *Erich3* |
| ENSMUSG00000027931 | *Npr1* | ENSMUSG00000078184 | *Rbm8a2* |
| ENSMUSG00000027935 | *Rab13* | ENSMUSG00000078427 | *Sarnp* |
| ENSMUSG00000027938 | *Creb3l4* | ENSMUSG00000078439 | *Smim24* |
| ENSMUSG00000027940 | *Tpm3* | ENSMUSG00000078442 | *Ccdc105* |
| ENSMUSG00000027959 | *Sass6* | ENSMUSG00000078451 | *Ppil6* |
| ENSMUSG00000027968 | *Larp7* | ENSMUSG00000078489 | *Gm17106* |
| ENSMUSG00000027981 | *Rnpc3* | ENSMUSG00000078490 | *Cfap74* |
| ENSMUSG00000028010 | *Gar1* | ENSMUSG00000078554 | *Fam229a* |
| ENSMUSG00000028028 | *Alpk1* | ENSMUSG00000078570 | *1110065P20Rik* |
| ENSMUSG00000028033 | *Kcnq5* | ENSMUSG00000078577 | *Tmco2* |
| ENSMUSG00000028034 | *Fubp1* | ENSMUSG00000078578 | *Ube2d3* |
| ENSMUSG00000028035 | *Dnajb4* | ENSMUSG00000078611 | *Gm5901* |
| ENSMUSG00000028047 | *Thbs3* | ENSMUSG00000078612 | *Fyb2* |
| ENSMUSG00000028053 | *Ash1l* | ENSMUSG00000078622 | *Ccdc47* |
| ENSMUSG00000028059 | *Arhgef2* | ENSMUSG00000078627 | *Marchf10* |
| ENSMUSG00000028060 | *Khdc4* | ENSMUSG00000078632 | *Lrrc37a* |
| ENSMUSG00000028069 | *Gpatch4* | ENSMUSG00000078652 | *Psme3* |
| ENSMUSG00000028073 | *Pear1* | ENSMUSG00000078653 | *Cntd1* |
| ENSMUSG00000028081 | *Rps3a1* | ENSMUSG00000078671 | *Chd2* |
| ENSMUSG00000028086 | *Fbxw7* | ENSMUSG00000078676 | *Casc3* |
| ENSMUSG00000028089 | *Chd1l* | ENSMUSG00000078684 | *5830417I10Rik* |
| ENSMUSG00000028099 | *Polr3c* | ENSMUSG00000078812 | *Eif5a* |
| ENSMUSG00000028104 | *Polr3gl* | ENSMUSG00000078813 | *Leng1* |
| ENSMUSG00000028109 | *Hormad1* | ENSMUSG00000078907 | *Fam186b* |
| ENSMUSG00000028126 | *Pip5k1a* | ENSMUSG00000078937 | *Cpt1b* |
| ENSMUSG00000028127 | *Abcd3* | ENSMUSG00000078941 | *Ak6* |
| ENSMUSG00000028136 | *Snx27* | ENSMUSG00000079003 | *Samd1* |
| ENSMUSG00000028137 | *Celf3* | ENSMUSG00000079005 | *Gm14147* |
| ENSMUSG00000028139 | *Riiad1* | ENSMUSG00000079006 | *Gm14151* |
| ENSMUSG00000028141 | *Oaz3* | ENSMUSG00000079019 | *Insl3* |
| ENSMUSG00000028156 | *Eif4e* | ENSMUSG00000079022 | *Col22a1* |
| ENSMUSG00000028180 | *Zranb2* | ENSMUSG00000079067 | *Hmgn2-ps1* |
| ENSMUSG00000028182 | *Lrriq3* | ENSMUSG00000079084 | *Ccdc82* |
| ENSMUSG00000028188 | *Spata1* | ENSMUSG00000079104 | *Prps1l3* |
| ENSMUSG00000028218 | *Cibar1* | ENSMUSG00000079105 | *C7* |
| ENSMUSG00000028224 | *Nbn* | ENSMUSG00000079108 | *Srp54c* |
| ENSMUSG00000028234 | *Rps20* | ENSMUSG00000079139 | *Gm4204* |
| ENSMUSG00000028248 | *Pnisr* | ENSMUSG00000079169 | *Gm15130* |
| ENSMUSG00000028274 | *Rngtt* | ENSMUSG00000079170 | *Gm13941* |
| ENSMUSG00000028277 | *Ube2j1* | ENSMUSG00000079173 | *Zan* |
| ENSMUSG00000028282 | *Casp8ap2* | ENSMUSG00000079177 | *Fam228a* |
| ENSMUSG00000028294 | *Cfap206* | ENSMUSG00000079184 | *Mphosph8* |
| ENSMUSG00000028309 | *Rnf20* | ENSMUSG00000079225 | *Gm9531* |
| ENSMUSG00000028312 | *Smc2* | ENSMUSG00000079235 | *Ccdc13* |
| ENSMUSG00000028328 | *Tmod1* | ENSMUSG00000079324 | *4932414N04Rik* |

| | | | |
|---|---|---|---|
| ENSMUSG00000028329 | *Xpa* | ENSMUSG00000079407 | *1700110I01Rik* |
| ENSMUSG00000028330 | *Ncbp1* | ENSMUSG00000079429 | *Mroh2a* |
| ENSMUSG00000028332 | *Hemgn* | ENSMUSG00000079465 | *Col4a3* |
| ENSMUSG00000028333 | *Anp32b* | ENSMUSG00000079470 | *Utp14b* |
| ENSMUSG00000028347 | *Tmeff1* | ENSMUSG00000079523 | *Tmsb10* |
| ENSMUSG00000028383 | *Hsdl2* | ENSMUSG00000079547 | *H2-DMb1* |
| ENSMUSG00000028389 | *Zfp37* | ENSMUSG00000079562 | *Maea* |
| ENSMUSG00000028392 | *Bspry* | ENSMUSG00000079579 | *Gm6760* |
| ENSMUSG00000028394 | *Pole3* | ENSMUSG00000079606 | *Fsip2l* |
| ENSMUSG00000028397 | *Kdm4c* | ENSMUSG00000079608 | *Stard6* |
| ENSMUSG00000028399 | *Ptprd* | ENSMUSG00000079652 | *Garin1a* |
| ENSMUSG00000028409 | *Smu1* | ENSMUSG00000079707 | *Dynlt2a1* |
| ENSMUSG00000028410 | *Dnaja1* | ENSMUSG00000079710 | *Dynlt2a2* |
| ENSMUSG00000028411 | *Aptx* | ENSMUSG00000079884 | *Tmed2b* |
| ENSMUSG00000028416 | *Bag1* | ENSMUSG00000080021 | *Gm5915* |
| ENSMUSG00000028419 | *Chmp5* | ENSMUSG00000080152 | *H3f4* |
| ENSMUSG00000028426 | *Rad23b* | ENSMUSG00000080237 | *Gm14239* |
| ENSMUSG00000028430 | *Nol6* | ENSMUSG00000080268 | *Brms1* |
| ENSMUSG00000028431 | *Elp1* | ENSMUSG00000080486 | *None* |
| ENSMUSG00000028433 | *Ubap2* | ENSMUSG00000080615 | *Snord99* |
| ENSMUSG00000028441 | *1110017D15Rik* | ENSMUSG00000080719 | *Gm15532* |
| ENSMUSG00000028456 | *Unc13b* | ENSMUSG00000080775 | *Gm6368* |
| ENSMUSG00000028457 | *Atp8b5* | ENSMUSG00000080776 | *Gm12174* |
| ENSMUSG00000028458 | *Tesk1* | ENSMUSG00000080778 | *Gm7199* |
| ENSMUSG00000028464 | *Tpm2* | ENSMUSG00000080806 | *Gm13005* |
| ENSMUSG00000028478 | *Clta* | ENSMUSG00000080860 | *Gm13815* |
| ENSMUSG00000028484 | *Psip1* | ENSMUSG00000080875 | *Gm7332* |
| ENSMUSG00000028494 | *Plin2* | ENSMUSG00000080902 | *Ywhaq-ps3* |
| ENSMUSG00000028495 | *Rps6* | ENSMUSG00000080968 | *Atf1-ps* |
| ENSMUSG00000028496 | *Mllt3* | ENSMUSG00000080974 | *Gm9009* |
| ENSMUSG00000028514 | *Usp24* | ENSMUSG00000081002 | *Gm14240* |
| ENSMUSG00000028519 | *Dab1* | ENSMUSG00000081044 | *AU015836* |
| ENSMUSG00000028520 | *4921539E11Rik* | ENSMUSG00000081046 | *Gm12846* |
| ENSMUSG00000028522 | *Mier1* | ENSMUSG00000081068 | *Gm12804* |
| ENSMUSG00000028530 | *Jak1* | ENSMUSG00000081087 | *Rps15a-ps7* |
| ENSMUSG00000028533 | *Izumo3* | ENSMUSG00000081094 | *Rpl19-ps11* |
| ENSMUSG00000028549 | *Itgb3bp* | ENSMUSG00000081104 | *Gm15989* |
| ENSMUSG00000028551 | *Cdkn2c* | ENSMUSG00000081111 | *Gm5913* |
| ENSMUSG00000028552 | *Eps15* | ENSMUSG00000081113 | *Gm7308* |
| ENSMUSG00000028555 | *Ttc39a* | ENSMUSG00000081125 | *Rpl26-ps5* |
| ENSMUSG00000028558 | *Calr4* | ENSMUSG00000081128 | *Gm13328* |
| ENSMUSG00000028559 | *Osbpl9* | ENSMUSG00000081151 | *Gm11448* |
| ENSMUSG00000028560 | *Usp1* | ENSMUSG00000081152 | *Gm12430* |
| ENSMUSG00000028568 | *Btf3l4* | ENSMUSG00000081159 | *Gm6023* |
| ENSMUSG00000028572 | *Hook1* | ENSMUSG00000081185 | *Gm4852* |
| ENSMUSG00000028575 | *Eqtn* | ENSMUSG00000081226 | *Gm8802* |
| ENSMUSG00000028576 | *Ift74* | ENSMUSG00000081227 | *Gm13082* |
| ENSMUSG00000028577 | *Plaa* | ENSMUSG00000081242 | *Gm15719* |
| ENSMUSG00000028580 | *Pum1* | ENSMUSG00000081243 | *Gm14037* |
| ENSMUSG00000028582 | *Cc2d1b* | ENSMUSG00000081281 | *Rpl7-ps9* |
| ENSMUSG00000028607 | *Cpt2* | ENSMUSG00000081289 | *Gm14857* |
| ENSMUSG00000028613 | *Lrp8* | ENSMUSG00000081302 | *Gm12020* |
| ENSMUSG00000028617 | *Lrrc42* | ENSMUSG00000081325 | *Gm12599* |
| ENSMUSG00000028629 | *Exo5* | ENSMUSG00000081352 | *Gm12389* |
| ENSMUSG00000028637 | *Ccdc30* | ENSMUSG00000081413 | *Uqcrh-ps2* |
| ENSMUSG00000028639 | *Ybx1* | ENSMUSG00000081434 | *Gm14165* |
| ENSMUSG00000028643 | *Svbp* | ENSMUSG00000081528 | *Gm12924* |
| ENSMUSG00000028649 | *Macf1* | ENSMUSG00000081552 | *Gm12328* |

| | | | |
|---|---|---|---|
| ENSMUSG00000028668 | *Eloa* | ENSMUSG00000081553 | *Ccdc50-ps* |
| ENSMUSG00000028676 | *Srsf10* | ENSMUSG00000081557 | *Gm5697* |
| ENSMUSG00000028677 | *Rnf220* | ENSMUSG00000081583 | *Gm14769* |
| ENSMUSG00000028678 | *Kif2c* | ENSMUSG00000081595 | *Gm13036* |
| ENSMUSG00000028689 | *Ccdc163* | ENSMUSG00000081603 | *Gm14681* |
| ENSMUSG00000028693 | *Nasp* | ENSMUSG00000081606 | *Gm9059* |
| ENSMUSG00000028698 | *Pik3r3* | ENSMUSG00000081623 | *Gm8809* |
| ENSMUSG00000028700 | *Pomgnt1* | ENSMUSG00000081627 | *Gm13158* |
| ENSMUSG00000028729 | *Ebna1bp2* | ENSMUSG00000081633 | *Gm8522* |
| ENSMUSG00000028730 | *Cfap57* | ENSMUSG00000081647 | *Gm13022* |
| ENSMUSG00000028745 | *Capzb* | ENSMUSG00000081656 | *Gm11246* |
| ENSMUSG00000028758 | *Kif17* | ENSMUSG00000081657 | *Gm15466* |
| ENSMUSG00000028759 | *Hp1bp3* | ENSMUSG00000081662 | *Gm14098* |
| ENSMUSG00000028760 | *Eif4g3* | ENSMUSG00000081670 | *Gm15697* |
| ENSMUSG00000028793 | *Rnf19b* | ENSMUSG00000081671 | *Gm13167* |
| ENSMUSG00000028795 | *Ccdc28b* | ENSMUSG00000081681 | *Gm14868* |
| ENSMUSG00000028809 | *Srrm1* | ENSMUSG00000081700 | *Atp5k-ps2* |
| ENSMUSG00000028820 | *Sfpq* | ENSMUSG00000081703 | *Gm6285* |
| ENSMUSG00000028821 | *Syf2* | ENSMUSG00000081729 | *Hspa9-ps1* |
| ENSMUSG00000028826 | *Maco1* | ENSMUSG00000081731 | *Calr-ps* |
| ENSMUSG00000028832 | *Stmn1* | ENSMUSG00000081788 | *Gm5898* |
| ENSMUSG00000028845 | *Tekt2* | ENSMUSG00000081789 | *Gm11893* |
| ENSMUSG00000028849 | *Map7d1* | ENSMUSG00000081792 | *Anp32b-ps1* |
| ENSMUSG00000028850 | *Gpatch3* | ENSMUSG00000081865 | *Gm15484* |
| ENSMUSG00000028851 | *Nudc* | ENSMUSG00000081871 | *Gm11488* |
| ENSMUSG00000028863 | *Meaf6* | ENSMUSG00000081885 | *Gm13231* |
| ENSMUSG00000028868 | *Wasf2* | ENSMUSG00000081887 | *Gm12097* |
| ENSMUSG00000028869 | *Gnl2* | ENSMUSG00000081888 | *Spcs2-ps* |
| ENSMUSG00000028873 | *Cdca8* | ENSMUSG00000081975 | *Gm12482* |
| ENSMUSG00000028878 | *Fam76a* | ENSMUSG00000081984 | *Dnajb3* |
| ENSMUSG00000028896 | *Rcc1* | ENSMUSG00000081992 | *Gm13408* |
| ENSMUSG00000028899 | *Taf12* | ENSMUSG00000082013 | *Gm13216* |
| ENSMUSG00000028901 | *Gmeb1* | ENSMUSG00000082043 | *Gm12848* |
| ENSMUSG00000028902 | *Sf3a3* | ENSMUSG00000082051 | *Gm16072* |
| ENSMUSG00000028911 | *Srsf4* | ENSMUSG00000082064 | *Rpl5-ps2* |
| ENSMUSG00000028917 | *Plekhm2* | ENSMUSG00000082078 | *Gm12098* |
| ENSMUSG00000028932 | *Psmc2* | ENSMUSG00000082100 | *Glns-ps1* |
| ENSMUSG00000028936 | *Rpl22* | ENSMUSG00000082117 | *Gm4994* |
| ENSMUSG00000028937 | *Acot7* | ENSMUSG00000082120 | *Gm15720* |
| ENSMUSG00000028938 | *Galntl5* | ENSMUSG00000082145 | *Gm12312* |
| ENSMUSG00000028943 | *Espn* | ENSMUSG00000082154 | *Gm16464* |
| ENSMUSG00000028944 | *Prkag2* | ENSMUSG00000082178 | *Gm11307* |
| ENSMUSG00000028953 | *Abcf2* | ENSMUSG00000082179 | *Gm11407* |
| ENSMUSG00000028954 | *Nub1* | ENSMUSG00000082194 | *Gm12444* |
| ENSMUSG00000028962 | *Slc4a2* | ENSMUSG00000082195 | *Gm13034* |
| ENSMUSG00000028976 | *Slc2a5* | ENSMUSG00000082253 | *Gm14639* |
| ENSMUSG00000028991 | *Mtor* | ENSMUSG00000082272 | *Gm11675* |
| ENSMUSG00000028995 | *Fam126a* | ENSMUSG00000082320 | *Gm11877* |
| ENSMUSG00000028998 | *Tomm7* | ENSMUSG00000082325 | *Gm12642* |
| ENSMUSG00000029001 | *Fbxo44* | ENSMUSG00000082348 | *Gm12807* |
| ENSMUSG00000029003 | *Mad2l2* | ENSMUSG00000082381 | *Gm12928* |
| ENSMUSG00000029004 | *Kmt2e* | ENSMUSG00000082419 | *Gm11425* |
| ENSMUSG00000029009 | *Mthfr* | ENSMUSG00000082454 | *Gm12183* |
| ENSMUSG00000029014 | *Dnajc2* | ENSMUSG00000082461 | *Gm8844* |
| ENSMUSG00000029017 | *Pmpcb* | ENSMUSG00000082491 | *Gm5909* |
| ENSMUSG00000029020 | *Mfn2* | ENSMUSG00000082530 | *Gm12168* |
| ENSMUSG00000029022 | *Miip* | ENSMUSG00000082536 | *Gm13456* |
| ENSMUSG00000029028 | *Lrrc47* | ENSMUSG00000082575 | *Eef2-ps2* |

| | | | |
|---|---|---|---|
| ENSMUSG00000029036 | *Atad3a* | ENSMUSG00000082625 | *Gm15204* |
| ENSMUSG00000029038 | *Ssu72* | ENSMUSG00000082652 | *Gm14725* |
| ENSMUSG00000029048 | *Rer1* | ENSMUSG00000082691 | *Dynlt1-ps1* |
| ENSMUSG00000029062 | *Cdk11b* | ENSMUSG00000082715 | *Gm11633* |
| ENSMUSG00000029066 | *Mrpl20* | ENSMUSG00000082774 | *Gm8805* |
| ENSMUSG00000029068 | *Ccnl2* | ENSMUSG00000082791 | *Gm4875* |
| ENSMUSG00000029071 | *Dvl1* | ENSMUSG00000082810 | *Gm8682* |
| ENSMUSG00000029074 | *Ttll10* | ENSMUSG00000082820 | *Gm13803* |
| ENSMUSG00000029076 | *Sdf4* | ENSMUSG00000082836 | *Gm13612* |
| ENSMUSG00000029086 | *Prom1* | ENSMUSG00000082872 | *Gm15773* |
| ENSMUSG00000029094 | *Afap1* | ENSMUSG00000082876 | *Gm11889* |
| ENSMUSG00000029103 | *Lrpap1* | ENSMUSG00000082878 | *Gm12583* |
| ENSMUSG00000029110 | *Rnf4* | ENSMUSG00000082896 | *Gm5844* |
| ENSMUSG00000029119 | *Man2b2* | ENSMUSG00000082930 | *Gm13462* |
| ENSMUSG00000029125 | *Stx18* | ENSMUSG00000082938 | *Gm2810* |
| ENSMUSG00000029128 | *Rab28* | ENSMUSG00000082971 | *Gm13517* |
| ENSMUSG00000029130 | *Rnf32* | ENSMUSG00000082998 | *Gm11221* |
| ENSMUSG00000029131 | *Dnajb6* | ENSMUSG00000083022 | *Rps15a-ps6* |
| ENSMUSG00000029134 | *Plb1* | ENSMUSG00000083027 | *Gm13140* |
| ENSMUSG00000029138 | *Ccdc121* | ENSMUSG00000083064 | *Gm7785* |
| ENSMUSG00000029141 | *Slc4a1ap* | ENSMUSG00000083081 | *Gm13223* |
| ENSMUSG00000029145 | *Eif2b4* | ENSMUSG00000083166 | *Gm8648* |
| ENSMUSG00000029147 | *Ppm1g* | ENSMUSG00000083175 | *Gm12206* |
| ENSMUSG00000029148 | *Nrbp1* | ENSMUSG00000083179 | *Gm12693* |
| ENSMUSG00000029155 | *Spata18* | ENSMUSG00000083218 | *Gm16425* |
| ENSMUSG00000029165 | *Agbl5* | ENSMUSG00000083240 | *Gm13453* |
| ENSMUSG00000029166 | *Mapre3* | ENSMUSG00000083242 | *Gm15659* |
| ENSMUSG00000029169 | *Dhx15* | ENSMUSG00000083258 | *Gm5939* |
| ENSMUSG00000029174 | *Tbc1d1* | ENSMUSG00000083261 | *Gm7816* |
| ENSMUSG00000029176 | *Anapc4* | ENSMUSG00000083270 | *Gm13498* |
| ENSMUSG00000029186 | *Pi4k2b* | ENSMUSG00000083307 | *AA414768* |
| ENSMUSG00000029191 | *Rfc1* | ENSMUSG00000083325 | *Gm14121* |
| ENSMUSG00000029192 | *Tbc1d14* | ENSMUSG00000083326 | *Rpl38-ps1* |
| ENSMUSG00000029196 | *Tada2b* | ENSMUSG00000083327 | *Vcp-rs* |
| ENSMUSG00000029201 | *Ugdh* | ENSMUSG00000083338 | *Gm12475* |
| ENSMUSG00000029202 | *Pds5a* | ENSMUSG00000083346 | *Gm8260* |
| ENSMUSG00000029203 | *Ube2k* | ENSMUSG00000083367 | *Gm8806* |
| ENSMUSG00000029206 | *Nsun7* | ENSMUSG00000083392 | *Gm6335* |
| ENSMUSG00000029212 | *Gabrb1* | ENSMUSG00000083396 | *Gm15542* |
| ENSMUSG00000029223 | *Uchl1* | ENSMUSG00000083429 | *Gm15198* |
| ENSMUSG00000029227 | *Fip1l1* | ENSMUSG00000083438 | *Gm5402* |
| ENSMUSG00000029229 | *Chic2* | ENSMUSG00000083453 | *Gm13338* |
| ENSMUSG00000029235 | *Pdcl2* | ENSMUSG00000083496 | *Gm11263* |
| ENSMUSG00000029238 | *Clock* | ENSMUSG00000083512 | *Gm12749* |
| ENSMUSG00000029249 | *Rest* | ENSMUSG00000083536 | *Gm15808* |
| ENSMUSG00000029250 | *Polr2b* | ENSMUSG00000083562 | *Gm13234* |
| ENSMUSG00000029267 | *Mtf2* | ENSMUSG00000083563 | *Gm13340* |
| ENSMUSG00000029276 | *Glmn* | ENSMUSG00000083579 | *Gm15538* |
| ENSMUSG00000029279 | *Brdt* | ENSMUSG00000083621 | *Gm14586* |
| ENSMUSG00000029290 | *Zfp326* | ENSMUSG00000083640 | *Gm8876* |
| ENSMUSG00000029291 | *Rufy3* | ENSMUSG00000083692 | *Gm9575* |
| ENSMUSG00000029309 | *Sparcl1* | ENSMUSG00000083720 | *Gm12901* |
| ENSMUSG00000029310 | *Nudt9* | ENSMUSG00000083736 | *Gm6039* |
| ENSMUSG00000029314 | *Gpat3* | ENSMUSG00000083815 | *Gm13435* |
| ENSMUSG00000029320 | *1700016H13Rik* | ENSMUSG00000083829 | *Gm2199* |
| ENSMUSG00000029328 | *Hnrnpdl* | ENSMUSG00000083830 | *Gm8839* |
| ENSMUSG00000029345 | *Tfip11* | ENSMUSG00000083834 | *Gm12577* |
| ENSMUSG00000029381 | *Shroom3* | ENSMUSG00000083838 | *Gm11957* |

| | | | |
|---|---|---|---|
| ENSMUSG00000029389 | *Ddx55* | ENSMUSG00000083890 | *Gm15703* |
| ENSMUSG00000029390 | *Tmed2* | ENSMUSG00000083899 | *Gm12346* |
| ENSMUSG00000029402 | *Snrnp35* | ENSMUSG00000083911 | *Gm4342* |
| ENSMUSG00000029403 | *Cdkl2* | ENSMUSG00000083912 | *Gm5391* |
| ENSMUSG00000029404 | *Arl6ip4* | ENSMUSG00000083928 | *Gm4991* |
| ENSMUSG00000029406 | *Pitpnm2* | ENSMUSG00000083937 | *Cct3-ps1* |
| ENSMUSG00000029415 | *Sdad1* | ENSMUSG00000083992 | *Gm11478* |
| ENSMUSG00000029422 | *Rsrc2* | ENSMUSG00000084006 | *Gm14245* |
| ENSMUSG00000029423 | *Piwil1* | ENSMUSG00000084067 | *Gm14269* |
| ENSMUSG00000029427 | *Zcchc8* | ENSMUSG00000084073 | *Gm11910* |
| ENSMUSG00000029428 | *Stx2* | ENSMUSG00000084077 | *Gm11952* |
| ENSMUSG00000029430 | *Ran* | ENSMUSG00000084081 | *Gm12057* |
| ENSMUSG00000029433 | *Diablo* | ENSMUSG00000084083 | *Gm15782* |
| ENSMUSG00000029439 | *Sfswap* | ENSMUSG00000084094 | *Gm13961* |
| ENSMUSG00000029440 | *Psmd9* | ENSMUSG00000084098 | *Gm13422* |
| ENSMUSG00000029442 | *Wdr66* | ENSMUSG00000084145 | *Gm12263* |
| ENSMUSG00000029447 | *Cct6a* | ENSMUSG00000084153 | *Gm12186* |
| ENSMUSG00000029458 | *Brap* | ENSMUSG00000084159 | *Gm12696* |
| ENSMUSG00000029463 | *Fam216a* | ENSMUSG00000084166 | *Gm6451* |
| ENSMUSG00000029467 | *Atp2a2* | ENSMUSG00000084183 | *Gm12009* |
| ENSMUSG00000029469 | *Ift81* | ENSMUSG00000084184 | *Gm13474* |
| ENSMUSG00000029472 | *Anapc5* | ENSMUSG00000084274 | *Ptma-ps2* |
| ENSMUSG00000029474 | *Rnf34* | ENSMUSG00000084279 | *Gm13232* |
| ENSMUSG00000029475 | *Kdm2b* | ENSMUSG00000084314 | *Rps15a-ps3* |
| ENSMUSG00000029477 | *Morn3* | ENSMUSG00000084342 | *Gm11224* |
| ENSMUSG00000029478 | *Ncor2* | ENSMUSG00000084378 | *Gm15238* |
| ENSMUSG00000029502 | *Golga3* | ENSMUSG00000084390 | *Gm15425* |
| ENSMUSG00000029505 | *Ep400* | ENSMUSG00000084729 | *Gm24139* |
| ENSMUSG00000029516 | *Cit* | ENSMUSG00000084780 | *Gm15350* |
| ENSMUSG00000029528 | *Pxn* | ENSMUSG00000084804 | *1700095J12Rik* |
| ENSMUSG00000029538 | *Srsf9* | ENSMUSG00000084817 | *Gm5526* |
| ENSMUSG00000029544 | *Cabp1* | ENSMUSG00000084833 | *Gm12649* |
| ENSMUSG00000029547 | *Ints1* | ENSMUSG00000084897 | *Gm14226* |
| ENSMUSG00000029552 | *Tes* | ENSMUSG00000084899 | *Gm15344* |
| ENSMUSG00000029554 | *Mad1l1* | ENSMUSG00000084948 | *1700061H18Rik* |
| ENSMUSG00000029564 | *4930519G04Rik* | ENSMUSG00000085006 | *BC021767* |
| ENSMUSG00000029576 | *Radil* | ENSMUSG00000085016 | *Gm11335* |
| ENSMUSG00000029580 | *Actb* | ENSMUSG00000085172 | *Gm6542* |
| ENSMUSG00000029594 | *Rbm19* | ENSMUSG00000085253 | *Gm16833* |
| ENSMUSG00000029598 | *Plbd2* | ENSMUSG00000085260 | *Med9os* |
| ENSMUSG00000029599 | *Ddx54* | ENSMUSG00000085278 | *Gm12841* |
| ENSMUSG00000029601 | *Iqcd* | ENSMUSG00000085486 | *Gm11634* |
| ENSMUSG00000029616 | *Erp29* | ENSMUSG00000085491 | *4930527E20Rik* |
| ENSMUSG00000029617 | *Ccz1* | ENSMUSG00000085757 | *Tslrn1* |
| ENSMUSG00000029623 | *Pdap1* | ENSMUSG00000085931 | *Gm12648* |
| ENSMUSG00000029629 | *Phf14* | ENSMUSG00000086046 | *1700095A21Rik* |
| ENSMUSG00000029634 | *Rnf6* | ENSMUSG00000086148 | *Gm15271* |
| ENSMUSG00000029635 | *Cdk8* | ENSMUSG00000086157 | *Gm14643* |
| ENSMUSG00000029640 | *Usp12* | ENSMUSG00000086175 | *Gm15802* |
| ENSMUSG00000029642 | *Polr1d* | ENSMUSG00000086229 | *Gm4887* |
| ENSMUSG00000029647 | *Pan3* | ENSMUSG00000086290 | *Snhg12* |
| ENSMUSG00000029655 | *N4bp2l2* | ENSMUSG00000086324 | *Gm15564* |
| ENSMUSG00000029670 | *Ing3* | ENSMUSG00000086379 | *1700026D11Rik* |
| ENSMUSG00000029686 | *Cul1* | ENSMUSG00000086459 | *1700030C12Rik* |
| ENSMUSG00000029687 | *Ezh2* | ENSMUSG00000086537 | *Gnasas1* |
| ENSMUSG00000029701 | *Rbm28* | ENSMUSG00000086654 | *Gm13165* |
| ENSMUSG00000029705 | *Cux1* | ENSMUSG00000086688 | *Gm11560* |
| ENSMUSG00000029708 | *Gcc1* | ENSMUSG00000086724 | *Gm12833* |

| | | | |
|---|---|---|---|
| ENSMUSG00000029714 | *Gigyf1* | ENSMUSG00000086732 | *Gm6117* |
| ENSMUSG00000029718 | *Pcolce* | ENSMUSG00000086788 | *1700029M20Rik* |
| ENSMUSG00000029720 | *Gm20605* | ENSMUSG00000086790 | *Gm12911* |
| ENSMUSG00000029729 | *Zkscan1* | ENSMUSG00000086841 | *2410006H16Rik* |
| ENSMUSG00000029730 | *Mcm7* | ENSMUSG00000086842 | *Gm15681* |
| ENSMUSG00000029752 | *Asns* | ENSMUSG00000086918 | *4930429F24Rik* |
| ENSMUSG00000029761 | *Cald1* | ENSMUSG00000086922 | *Gm13835* |
| ENSMUSG00000029767 | *Calu* | ENSMUSG00000086923 | *4930406D18Rik* |
| ENSMUSG00000029769 | *Ccdc136* | ENSMUSG00000086951 | *1700051A21Rik* |
| ENSMUSG00000029772 | *Ahcyl2* | ENSMUSG00000087104 | *Tmem132cos* |
| ENSMUSG00000029784 | *Ssmem1* | ENSMUSG00000087122 | *4930403D09Rik* |
| ENSMUSG00000029817 | *Tra2a* | ENSMUSG00000087153 | *Gm6483* |
| ENSMUSG00000029823 | *Luc7l2* | ENSMUSG00000087162 | *Gm14244* |
| ENSMUSG00000029828 | *4921507P07Rik* | ENSMUSG00000087177 | *E130307A14Rik* |
| ENSMUSG00000029833 | *Trim24* | ENSMUSG00000087202 | *Gm15813* |
| ENSMUSG00000029848 | *Stra8* | ENSMUSG00000087204 | *1700073E17Rik* |
| ENSMUSG00000029875 | *Ccdc184* | ENSMUSG00000087233 | *Gm43213* |
| ENSMUSG00000029883 | *Prss59* | ENSMUSG00000087479 | *Gm16835* |
| ENSMUSG00000029909 | *Prss37* | ENSMUSG00000087526 | *Gm15738* |
| ENSMUSG00000029917 | *Qrfprl* | ENSMUSG00000087635 | *Gm13414* |
| ENSMUSG00000029920 | *Smarcad1* | ENSMUSG00000088294 | *None* |
| ENSMUSG00000029922 | *Mkrn1* | ENSMUSG00000088524 | *Snord2* |
| ENSMUSG00000029992 | *Gfpt1* | ENSMUSG00000088948 | *Gm23262* |
| ENSMUSG00000030007 | *Cct7* | ENSMUSG00000089357 | *Mir2137* |
| ENSMUSG00000030016 | *Zfp638* | ENSMUSG00000089673 | *Gm16546* |
| ENSMUSG00000030030 | *1700003E16Rik* | ENSMUSG00000089682 | *Bcl2l2* |
| ENSMUSG00000030034 | *Ino80b* | ENSMUSG00000089698 | *Gm2541* |
| ENSMUSG00000030041 | *M1ap* | ENSMUSG00000089717 | *Olfr275* |
| ENSMUSG00000030051 | *Aplf* | ENSMUSG00000089730 | *1700007P06Rik* |
| ENSMUSG00000030056 | *Isy1* | ENSMUSG00000089743 | *Gm16226* |
| ENSMUSG00000030059 | *Tmf1* | ENSMUSG00000089782 | *Btf3-ps1* |
| ENSMUSG00000030062 | *Rpn1* | ENSMUSG00000089791 | *Gm3555* |
| ENSMUSG00000030077 | *Chl1* | ENSMUSG00000089808 | *Gm7241* |
| ENSMUSG00000030122 | *Ptms* | ENSMUSG00000089865 | *Gm44503* |
| ENSMUSG00000030125 | *Lrrc23* | ENSMUSG00000089887 | *4930428N03Rik* |
| ENSMUSG00000030126 | *Tmcc1* | ENSMUSG00000089911 | *Mfsd14a* |
| ENSMUSG00000030127 | *Cops7a* | ENSMUSG00000089945 | *Pakap* |
| ENSMUSG00000030137 | *Tuba8* | ENSMUSG00000089958 | *Gm16165* |
| ENSMUSG00000030138 | *Bms1* | ENSMUSG00000089975 | *Gm7420* |
| ENSMUSG00000030161 | *Gabarapl1* | ENSMUSG00000089984 | *Fbxo24* |
| ENSMUSG00000030172 | *Erc1* | ENSMUSG00000089988 | *Gm16238* |
| ENSMUSG00000030177 | *Ccdc77* | ENSMUSG00000089989 | *Gm45713* |
| ENSMUSG00000030180 | *Kdm5a* | ENSMUSG00000090000 | *Ier3ip1* |
| ENSMUSG00000030189 | *Ybx3* | ENSMUSG00000090027 | *Gm15740* |
| ENSMUSG00000030200 | *Bcl2l14* | ENSMUSG00000090077 | *Lime1* |
| ENSMUSG00000030204 | *Ddx47* | ENSMUSG00000090083 | *Rnf8* |
| ENSMUSG00000030206 | *Gsg1* | ENSMUSG00000090100 | *Ttbk2* |
| ENSMUSG00000030213 | *Atf7ip* | ENSMUSG00000090115 | *Usp49* |
| ENSMUSG00000030216 | *Wbp11* | ENSMUSG00000090119 | *Gm8185* |
| ENSMUSG00000030224 | *Strap* | ENSMUSG00000090173 | *Fbxw10* |
| ENSMUSG00000030228 | *Pik3c2g* | ENSMUSG00000090191 | *9230105E05Rik* |
| ENSMUSG00000030230 | *Plcz1* | ENSMUSG00000090202 | *4930503B20Rik* |
| ENSMUSG00000030231 | *Plekha5* | ENSMUSG00000090319 | *Gm4462* |
| ENSMUSG00000030232 | *Aebp2* | ENSMUSG00000090323 | *Gm5263* |
| ENSMUSG00000030243 | *Recql* | ENSMUSG00000090369 | *4933411K16Rik* |
| ENSMUSG00000030254 | *Rad18* | ENSMUSG00000090389 | *Cdv3-ps* |
| ENSMUSG00000030257 | *Srgap3* | ENSMUSG00000090394 | *4930523C07Rik* |
| ENSMUSG00000030265 | *Kras* | ENSMUSG00000090457 | *4930571K23Rik* |

| | | | |
|---|---|---|---|
| ENSMUSG00000030275 | *Etnk1* | ENSMUSG00000090516 | *Rps11-ps1* |
| ENSMUSG00000030276 | *Ttll3* | ENSMUSG00000090585 | *4933406F09Rik* |
| ENSMUSG00000030287 | *Itpr2* | ENSMUSG00000090602 | *Gm5611* |
| ENSMUSG00000030292 | *Smco2* | ENSMUSG00000090626 | *Tex9* |
| ENSMUSG00000030301 | *Ccdc91* | ENSMUSG00000090671 | *Gm5067* |
| ENSMUSG00000030304 | *Ergic2* | ENSMUSG00000090777 | *Ccdc188* |
| ENSMUSG00000030314 | *Atg7* | ENSMUSG00000090812 | *Samd15* |
| ENSMUSG00000030321 | *Efcab12* | ENSMUSG00000090816 | *Gm6729* |
| ENSMUSG00000030323 | *Ift122* | ENSMUSG00000090840 | *1700092M07Rik* |
| ENSMUSG00000030335 | *Mrpl51* | ENSMUSG00000090841 | *Myl6* |
| ENSMUSG00000030344 | *Akap3* | ENSMUSG00000090877 | *Hspa1b* |
| ENSMUSG00000030346 | *Rad51ap1* | ENSMUSG00000090935 | *Synj2bp* |
| ENSMUSG00000030357 | *Fkbp4* | ENSMUSG00000091002 | *Tcerg1l* |
| ENSMUSG00000030374 | *Strn4* | ENSMUSG00000091010 | *Gm17107* |
| ENSMUSG00000030397 | *Mark4* | ENSMUSG00000091017 | *Garin4* |
| ENSMUSG00000030403 | *Vasp* | ENSMUSG00000091119 | *Ccdc152* |
| ENSMUSG00000030417 | *Pdcd5* | ENSMUSG00000091255 | *Speer4e* |
| ENSMUSG00000030421 | *Uri1* | ENSMUSG00000091311 | *Spata31d1b* |
| ENSMUSG00000030435 | *U2af2* | ENSMUSG00000091337 | *Eid1* |
| ENSMUSG00000030451 | *Herc2* | ENSMUSG00000091358 | *Speer5-ps1* |
| ENSMUSG00000030465 | *Psd3* | ENSMUSG00000091415 | *Ak9* |
| ENSMUSG00000030491 | *Tdrd12* | ENSMUSG00000091416 | *Gm6327* |
| ENSMUSG00000030498 | *Gas2* | ENSMUSG00000091421 | *Gm4202* |
| ENSMUSG00000030510 | *Cers3* | ENSMUSG00000091422 | *Gm6455* |
| ENSMUSG00000030515 | *Tarsl2* | ENSMUSG00000091457 | *Gm17171* |
| ENSMUSG00000030516 | *Tjp1* | ENSMUSG00000091476 | *Catspere2* |
| ENSMUSG00000030521 | *Mphosph10* | ENSMUSG00000091537 | *Tma7* |
| ENSMUSG00000030528 | *Blm* | ENSMUSG00000091556 | *Gm14569* |
| ENSMUSG00000030534 | *Vps33b* | ENSMUSG00000091561 | *Gm6665* |
| ENSMUSG00000030536 | *Iqgap1* | ENSMUSG00000091723 | *Gm17366* |
| ENSMUSG00000030555 | *Ttc23* | ENSMUSG00000091735 | *Gpr62* |
| ENSMUSG00000030557 | *Mef2a* | ENSMUSG00000091742 | *Gm5093* |
| ENSMUSG00000030595 | *Nfkbib* | ENSMUSG00000091803 | *Cox16* |
| ENSMUSG00000030603 | *Psmc4* | ENSMUSG00000091825 | *Gm5778* |
| ENSMUSG00000030617 | *Ccdc83* | ENSMUSG00000091827 | *Speer4f2* |
| ENSMUSG00000030619 | *Eed* | ENSMUSG00000091844 | *Gm8251* |
| ENSMUSG00000030638 | *Sh3gl3* | ENSMUSG00000091896 | *Ube2d2a* |
| ENSMUSG00000030649 | *Anapc15* | ENSMUSG00000091902 | *Gm6918* |
| ENSMUSG00000030655 | *Smg1* | ENSMUSG00000091905 | *Dnajb6-ps* |
| ENSMUSG00000030659 | *Nucb2* | ENSMUSG00000091931 | *Gon7* |
| ENSMUSG00000030662 | *Ipo5* | ENSMUSG00000091933 | *Gm8857* |
| ENSMUSG00000030663 | *1110004F10Rik* | ENSMUSG00000091955 | *Gm9844* |
| ENSMUSG00000030671 | *Pde3b* | ENSMUSG00000091971 | *Hspa1a* |
| ENSMUSG00000030680 | *Pagr1a* | ENSMUSG00000091993 | *B930036N10Rik* |
| ENSMUSG00000030688 | *Stard10* | ENSMUSG00000092009 | *Myh15* |
| ENSMUSG00000030695 | *Aldoa* | ENSMUSG00000092054 | *Kif4-ps* |
| ENSMUSG00000030718 | *Ppme1* | ENSMUSG00000092056 | *Gm7289* |
| ENSMUSG00000030726 | *Pold3* | ENSMUSG00000092192 | *Dnaaf4* |
| ENSMUSG00000030727 | *Rabep2* | ENSMUSG00000092220 | *Gm20528* |
| ENSMUSG00000030737 | *Slco2b1* | ENSMUSG00000092232 | *Gm20521* |
| ENSMUSG00000030738 | *Eif3c* | ENSMUSG00000092274 | *Neat1* |
| ENSMUSG00000030761 | *Myo7a* | ENSMUSG00000092278 | *Gm8752* |
| ENSMUSG00000030768 | *Disp1* | ENSMUSG00000092305 | *Prps1l1* |
| ENSMUSG00000030779 | *Rbbp6* | ENSMUSG00000092329 | *None* |
| ENSMUSG00000030788 | *Rnf141* | ENSMUSG00000092341 | *Malat1* |
| ENSMUSG00000030792 | *Dkkl1* | ENSMUSG00000092518 | *Garin5b* |
| ENSMUSG00000030795 | *Fus* | ENSMUSG00000092522 | *Gm20389* |
| ENSMUSG00000030801 | *Kat8* | ENSMUSG00000092541 | *Gm20537* |

| | | | |
|---|---|---|---|
| ENSMUSG00000030805 | *Stx4a* | ENSMUSG00000092560 | *Gm8750* |
| ENSMUSG00000030811 | *Fbxl19* | ENSMUSG00000092599 | *1700010K23Rik* |
| ENSMUSG00000030815 | *Phkg2* | ENSMUSG00000092680 | *Snord55* |
| ENSMUSG00000030816 | *Rnf40* | ENSMUSG00000092702 | *Gm24514* |
| ENSMUSG00000030824 | *Nucb1* | ENSMUSG00000092746 | *Rn7s6* |
| ENSMUSG00000030846 | *Tial1* | ENSMUSG00000092805 | *Gm26461* |
| ENSMUSG00000030849 | *Fgfr2* | ENSMUSG00000093178 | *Snord87* |
| ENSMUSG00000030850 | *Ate1* | ENSMUSG00000093183 | *Gm25687* |
| ENSMUSG00000030851 | *Ldhc* | ENSMUSG00000093383 | *Gm20642* |
| ENSMUSG00000030877 | *Mfsd13b* | ENSMUSG00000093384 | *Gm20689* |
| ENSMUSG00000030878 | *Cdr2* | ENSMUSG00000093394 | *Gm20621* |
| ENSMUSG00000030882 | *Dnhd1* | ENSMUSG00000093445 | *Lrch4* |
| ENSMUSG00000030887 | *Pdzd9* | ENSMUSG00000093484 | *Gm20657* |
| ENSMUSG00000030889 | *Vwa3a* | ENSMUSG00000093535 | *4930533I22Rik* |
| ENSMUSG00000030924 | *Rexo5* | ENSMUSG00000093537 | *Gm7584* |
| ENSMUSG00000030942 | *Thumpd1* | ENSMUSG00000093545 | *Gm5871* |
| ENSMUSG00000030965 | *Abraxas2* | ENSMUSG00000093575 | *Gm20695* |
| ENSMUSG00000030967 | *Zranb1* | ENSMUSG00000093617 | *4933438K21Rik* |
| ENSMUSG00000030968 | *Pdilt* | ENSMUSG00000093671 | *Gm20656* |
| ENSMUSG00000030978 | *Rrm1* | ENSMUSG00000093674 | *Rpl41* |
| ENSMUSG00000030980 | *Knop1* | ENSMUSG00000093752 | *Gm20716* |
| ENSMUSG00000030983 | *Bccip* | ENSMUSG00000093793 | *Gm24091* |
| ENSMUSG00000030987 | *Stim1* | ENSMUSG00000093843 | *Gm25939* |
| ENSMUSG00000030994 | *D7Ertd443e* | ENSMUSG00000093919 | *Gm20760* |
| ENSMUSG00000031004 | *Mki67* | ENSMUSG00000093956 | *Gm24497* |
| ENSMUSG00000031010 | *Usp9x* | ENSMUSG00000094017 | *Gm13160* |
| ENSMUSG00000031015 | *Swap70* | ENSMUSG00000094036 | *Gm6465* |
| ENSMUSG00000031027 | *Stk33* | ENSMUSG00000094050 | *Gm23472* |
| ENSMUSG00000031028 | *Tub* | ENSMUSG00000094090 | *Gm8494* |
| ENSMUSG00000031068 | *Glrx3* | ENSMUSG00000094103 | *Fam177a2* |
| ENSMUSG00000031075 | *Ano1* | ENSMUSG00000094105 | *Gm8922* |
| ENSMUSG00000031078 | *Cttn* | ENSMUSG00000094205 | *Gm8926* |
| ENSMUSG00000031139 | *Mcf2* | ENSMUSG00000094230 | *Gm21847* |
| ENSMUSG00000031153 | *Gripap1* | ENSMUSG00000094251 | *Gm25632* |
| ENSMUSG00000031161 | *Hdac6* | ENSMUSG00000094344 | *Gm11942* |
| ENSMUSG00000031174 | *Rpgr* | ENSMUSG00000094377 | *Gm24407* |
| ENSMUSG00000031196 | *F8* | ENSMUSG00000094382 | *Gm22192* |
| ENSMUSG00000031226 | *Pbdc1* | ENSMUSG00000094405 | *Gm23143* |
| ENSMUSG00000031229 | *Atrx* | ENSMUSG00000094410 | *Zbed6* |
| ENSMUSG00000031233 | *Pgk2* | ENSMUSG00000094437 | *Gm9830* |
| ENSMUSG00000031245 | *Hmgn5* | ENSMUSG00000094445 | *1700015G11Rik* |
| ENSMUSG00000031256 | *Cstf2* | ENSMUSG00000094597 | *Gm4810* |
| ENSMUSG00000031292 | *Cdkl5* | ENSMUSG00000094638 | *Gm21972* |
| ENSMUSG00000031303 | *Map3k15* | ENSMUSG00000094812 | *None* |
| ENSMUSG00000031311 | *Nono* | ENSMUSG00000094826 | *Gm23804* |
| ENSMUSG00000031314 | *Taf1* | ENSMUSG00000094830 | *None* |
| ENSMUSG00000031333 | *Abcb7* | ENSMUSG00000094843 | *Ppp1r2-ps1* |
| ENSMUSG00000031351 | *Zfp185* | ENSMUSG00000094936 | *Rbm4* |
| ENSMUSG00000031353 | *Rbbp7* | ENSMUSG00000095054 | *Gm13146* |
| ENSMUSG00000031370 | *Zrsr2* | ENSMUSG00000095110 | *1700003E24Rik* |
| ENSMUSG00000031371 | *Haus7* | ENSMUSG00000095118 | *None* |
| ENSMUSG00000031446 | *Cul4a* | ENSMUSG00000095159 | *Tubb4b-ps1* |
| ENSMUSG00000031451 | *Gas6* | ENSMUSG00000095180 | *Rhox5* |
| ENSMUSG00000031452 | *1700029H14Rik* | ENSMUSG00000095260 | *None* |
| ENSMUSG00000031483 | *Erlin2* | ENSMUSG00000095296 | *Gm8906* |
| ENSMUSG00000031485 | *Plpbp* | ENSMUSG00000095403 | *Gm21092* |
| ENSMUSG00000031493 | *Ggn* | ENSMUSG00000095464 | *Gm21987* |
| ENSMUSG00000031502 | *Col4a1* | ENSMUSG00000095469 | *Gm21965* |

| | | | |
|---|---|---|---|
| ENSMUSG00000031503 | *Col4a2* | ENSMUSG00000095513 | *None* |
| ENSMUSG00000031509 | *1700016D06Rik* | ENSMUSG00000095562 | *Gm55594* |
| ENSMUSG00000031511 | *Arhgef7* | ENSMUSG00000095567 | *Noc2l* |
| ENSMUSG00000031518 | *Spata4* | ENSMUSG00000095580 | *Rnu1b1* |
| ENSMUSG00000031529 | *Tnks* | ENSMUSG00000095595 | *Fam177a* |
| ENSMUSG00000031533 | *Mrps31* | ENSMUSG00000095676 | *Gm25099* |
| ENSMUSG00000031536 | *Polb* | ENSMUSG00000095701 | *None* |
| ENSMUSG00000031540 | *Kat6a* | ENSMUSG00000095710 | *Gm8871* |
| ENSMUSG00000031545 | *Gpat4* | ENSMUSG00000095738 | *Gm25313* |
| ENSMUSG00000031552 | *Adam18* | ENSMUSG00000095762 | *Gm16378* |
| ENSMUSG00000031553 | *Adam3* | ENSMUSG00000095829 | *Speer1* |
| ENSMUSG00000031554 | *Adam5* | ENSMUSG00000095865 | *Gm13237* |
| ENSMUSG00000031559 | *4930555F03Rik* | ENSMUSG00000095869 | *Ppp1r2-ps6* |
| ENSMUSG00000031563 | *Wwc2* | ENSMUSG00000095892 | *Rnu5g* |
| ENSMUSG00000031575 | *Ash2l* | ENSMUSG00000095895 | *Gm21963* |
| ENSMUSG00000031578 | *Mak16* | ENSMUSG00000095918 | *Gm5861* |
| ENSMUSG00000031585 | *Gtf2e2* | ENSMUSG00000095969 | *Rnu1a1* |
| ENSMUSG00000031590 | *Frg1* | ENSMUSG00000096006 | *Gm21596* |
| ENSMUSG00000031592 | *Pcm1* | ENSMUSG00000096054 | *Syne1* |
| ENSMUSG00000031601 | *Cnot7* | ENSMUSG00000096141 | *Dnah7a* |
| ENSMUSG00000031617 | *Tmem184c* | ENSMUSG00000096153 | *BC061195* |
| ENSMUSG00000031620 | *Iqcm* | ENSMUSG00000096160 | *Gm3436* |
| ENSMUSG00000031622 | *Sin3b* | ENSMUSG00000096205 | *Gm22068* |
| ENSMUSG00000031629 | *Cenpu* | ENSMUSG00000096218 | *Gm2916* |
| ENSMUSG00000031631 | *Cfap97* | ENSMUSG00000096233 | *Gm13238* |
| ENSMUSG00000031644 | *Nek1* | ENSMUSG00000096243 | *Gm24265* |
| ENSMUSG00000031647 | *Mfap3l* | ENSMUSG00000096255 | *Dynlt1b* |
| ENSMUSG00000031651 | *Triml1* | ENSMUSG00000096263 | *None* |
| ENSMUSG00000031660 | *Brd7* | ENSMUSG00000096280 | *Gm23287* |
| ENSMUSG00000031666 | *Rbl2* | ENSMUSG00000096349 | *Gm22513* |
| ENSMUSG00000031682 | *1700011L22Rik* | ENSMUSG00000096370 | *Gm21992* |
| ENSMUSG00000031683 | *Lsm6* | ENSMUSG00000096401 | *Gm21811* |
| ENSMUSG00000031691 | *Tnpo2* | ENSMUSG00000096405 | *4930474N05Rik* |
| ENSMUSG00000031696 | *Vps35* | ENSMUSG00000096458 | *Moap1* |
| ENSMUSG00000031697 | *Orc6* | ENSMUSG00000096732 | *Gm8897* |
| ENSMUSG00000031701 | *Dnaja2* | ENSMUSG00000096740 | *Lbhd1* |
| ENSMUSG00000031711 | *Zfp330* | ENSMUSG00000096764 | *Gm21985* |
| ENSMUSG00000031715 | *Smarca5* | ENSMUSG00000096768 | *Gm47283* |
| ENSMUSG00000031727 | *Pmfbp1* | ENSMUSG00000096793 | *Gm3002* |
| ENSMUSG00000031728 | *Zfp821* | ENSMUSG00000096838 | *None* |
| ENSMUSG00000031729 | *Ist1* | ENSMUSG00000096846 | *Gm8890* |
| ENSMUSG00000031731 | *Ap1g1* | ENSMUSG00000096878 | *Gm21083* |
| ENSMUSG00000031756 | *Cenpn* | ENSMUSG00000096887 | *Gm20594* |
| ENSMUSG00000031774 | *Psme3ip1* | ENSMUSG00000096914 | *Galntl6* |
| ENSMUSG00000031776 | *Arl2bp* | ENSMUSG00000096916 | *Zfp850* |
| ENSMUSG00000031786 | *Drc7* | ENSMUSG00000097032 | *4930539J05Rik* |
| ENSMUSG00000031788 | *Kifc3* | ENSMUSG00000097075 | *Cdiptos* |
| ENSMUSG00000031796 | *Cfap20* | ENSMUSG00000097098 | *9330111N05Rik* |
| ENSMUSG00000031805 | *Jak3* | ENSMUSG00000097177 | *9330159M07Rik* |
| ENSMUSG00000031809 | *1700018B08Rik* | ENSMUSG00000097207 | *6030443J06Rik* |
| ENSMUSG00000031820 | *Babam1* | ENSMUSG00000097239 | *Gm27029* |
| ENSMUSG00000031831 | *Dnaaf1* | ENSMUSG00000097271 | *Gm9903* |
| ENSMUSG00000031839 | *Hsbp1* | ENSMUSG00000097375 | *6720427I07Rik* |
| ENSMUSG00000031843 | *Mphosph6* | ENSMUSG00000097392 | *Thoc2l* |
| ENSMUSG00000031845 | *Bco1* | ENSMUSG00000097447 | *Gm26630* |
| ENSMUSG00000031847 | *1700030J22Rik* | ENSMUSG00000097451 | *Rian* |
| ENSMUSG00000031849 | *Comp* | ENSMUSG00000097494 | *4933406C10Rik* |
| ENSMUSG00000031851 | *Ntpcr* | ENSMUSG00000097520 | *4930488L21Rik* |

| | | | |
|---|---|---|---|
| ENSMUSG00000031864 | *Ints10* | ENSMUSG00000097772 | *5430416N02Rik* |
| ENSMUSG00000031865 | *Dctn1* | ENSMUSG00000097901 | *Gm26680* |
| ENSMUSG00000031878 | *Nae1* | ENSMUSG00000097906 | *Gm9625* |
| ENSMUSG00000031893 | *Tsnaxip1* | ENSMUSG00000098004 | *Gm27027* |
| ENSMUSG00000031898 | *Dpep3* | ENSMUSG00000098090 | *2700099C18Rik* |
| ENSMUSG00000031913 | *Vps4a* | ENSMUSG00000098104 | *Gm6085* |
| ENSMUSG00000031922 | *Cep57* | ENSMUSG00000098112 | *Bin2* |
| ENSMUSG00000031927 | *1700012B09Rik* | ENSMUSG00000098127 | *Gm4665* |
| ENSMUSG00000031928 | *Mre11a* | ENSMUSG00000098134 | *Rnf113a2* |
| ENSMUSG00000031930 | *Wwp2* | ENSMUSG00000098198 | *Gm9169* |
| ENSMUSG00000031939 | *Taf1d* | ENSMUSG00000098234 | *Snhg6* |
| ENSMUSG00000031954 | *Cfdp1* | ENSMUSG00000098374 | *Gm28043* |
| ENSMUSG00000031960 | *Aars* | ENSMUSG00000098404 | *Mrip-ps* |
| ENSMUSG00000031971 | *Ccsap* | ENSMUSG00000098425 | *Gm24949* |
| ENSMUSG00000031979 | *Cog2* | ENSMUSG00000098530 | *Gm28051* |
| ENSMUSG00000031984 | *2810004N23Rik* | ENSMUSG00000098641 | *None* |
| ENSMUSG00000031986 | *Sprtn* | ENSMUSG00000098761 | *Gm18821* |
| ENSMUSG00000031991 | *Spata19* | ENSMUSG00000098875 | *Gm27211* |
| ENSMUSG00000031996 | *Aplp2* | ENSMUSG00000098973 | *Mir6236* |
| ENSMUSG00000032002 | *Dcun1d5* | ENSMUSG00000098985 | *Gm27219* |
| ENSMUSG00000032006 | *Pdgfd* | ENSMUSG00000099021 | *Rn7s1* |
| ENSMUSG00000032009 | *Sesn3* | ENSMUSG00000099250 | *Rn7s2* |
| ENSMUSG00000032010 | *Usp2* | ENSMUSG00000099329 | *Gm28052* |
| ENSMUSG00000032023 | *Jhy* | ENSMUSG00000099353 | *1700031M16Rik* |
| ENSMUSG00000032030 | *Cul5* | ENSMUSG00000099418 | *Gm6657* |
| ENSMUSG00000032050 | *Rdx* | ENSMUSG00000099422 | *Gm4275* |
| ENSMUSG00000032058 | *Ppp2r1b* | ENSMUSG00000099481 | *Xndc1* |
| ENSMUSG00000032064 | *Dixdc1* | ENSMUSG00000099492 | *Gm5525* |
| ENSMUSG00000032076 | *Cadm1* | ENSMUSG00000099639 | *1700084F23Rik* |
| ENSMUSG00000032078 | *Zpr1* | ENSMUSG00000099662 | *Ppp1r2-ps2* |
| ENSMUSG00000032096 | *Arcn1* | ENSMUSG00000099671 | *Gm18183* |
| ENSMUSG00000032097 | *Ddx6* | ENSMUSG00000099762 | *Gm21149* |
| ENSMUSG00000032101 | *Ddx25* | ENSMUSG00000099858 | *Gm6652* |
| ENSMUSG00000032115 | *Hyou1* | ENSMUSG00000099869 | *1700030F04Rik* |
| ENSMUSG00000032116 | *Stt3a* | ENSMUSG00000099966 | *2810402E24Rik* |
| ENSMUSG00000032177 | *Pde4a* | ENSMUSG00000099998 | *Gm6501* |
| ENSMUSG00000032178 | *Ilf3* | ENSMUSG00000100104 | *Gm5644* |
| ENSMUSG00000032186 | *Tmod2* | ENSMUSG00000100153 | *Ppp1ccb* |
| ENSMUSG00000032187 | *Smarca4* | ENSMUSG00000100253 | *1700020N18Rik* |
| ENSMUSG00000032212 | *Sltm* | ENSMUSG00000100347 | *Gm7895* |
| ENSMUSG00000032216 | *Nedd4* | ENSMUSG00000100389 | *Rbm6-ps2* |
| ENSMUSG00000032217 | *Rnf111* | ENSMUSG00000100615 | *Gm5511* |
| ENSMUSG00000032218 | *Ccnb2* | ENSMUSG00000100704 | *Gm28434* |
| ENSMUSG00000032221 | *Mns1* | ENSMUSG00000100727 | *1700003P14Rik* |
| ENSMUSG00000032231 | *Anxa2* | ENSMUSG00000100863 | *Gm12669* |
| ENSMUSG00000032232 | *Cgnl1* | ENSMUSG00000100937 | *Nscme3l* |
| ENSMUSG00000032239 | *Rp9* | ENSMUSG00000100969 | *1700030N03Rik* |
| ENSMUSG00000032244 | *Fem1b* | ENSMUSG00000101012 | *1700008K24Rik* |
| ENSMUSG00000032249 | *Anp32a* | ENSMUSG00000101111 | *Gm28437* |
| ENSMUSG00000032253 | *Phip* | ENSMUSG00000101166 | *Gm28496* |
| ENSMUSG00000032254 | *Kif23* | ENSMUSG00000101188 | *Eif4a-ps4* |
| ENSMUSG00000032258 | *Lca5* | ENSMUSG00000101227 | *Gm29506* |
| ENSMUSG00000032274 | *Cyp19a1* | ENSMUSG00000101281 | *Gm8449* |
| ENSMUSG00000032279 | *Idh3a* | ENSMUSG00000101337 | *Dnah7c* |
| ENSMUSG00000032281 | *Acsbg1* | ENSMUSG00000101589 | *Rbm6-ps1* |
| ENSMUSG00000032285 | *Dnaja4* | ENSMUSG00000101605 | *Ace3* |
| ENSMUSG00000032294 | *Pkm* | ENSMUSG00000101645 | *Gm28635* |
| ENSMUSG00000032301 | *Psma4* | ENSMUSG00000101685 | *Gm17837* |

| | | | |
|---|---|---|---|
| ENSMUSG00000032307 | *Ube2q2* | ENSMUSG00000101801 | *1700020G17Rik* |
| ENSMUSG00000032328 | *Tmem30a* | ENSMUSG00000101814 | *Gm17807* |
| ENSMUSG00000032329 | *Hmg20a* | ENSMUSG00000101854 | *1700026F02Rik* |
| ENSMUSG00000032330 | *Cox7a2* | ENSMUSG00000101895 | *Gm28981* |
| ENSMUSG00000032348 | *Gsta4* | ENSMUSG00000101904 | *Gm29427* |
| ENSMUSG00000032350 | *Gclc* | ENSMUSG00000101963 | *1700001J11Rik* |
| ENSMUSG00000032352 | *Lrrc1* | ENSMUSG00000101970 | *Chaserr* |
| ENSMUSG00000032366 | *Tpm1* | ENSMUSG00000102018 | *Iqank1* |
| ENSMUSG00000032382 | *Snx1* | ENSMUSG00000102206 | *Pcdha11* |
| ENSMUSG00000032386 | *Trip4* | ENSMUSG00000102245 | *Gm34423* |
| ENSMUSG00000032393 | *Dpp8* | ENSMUSG00000102312 | *Pcdha3* |
| ENSMUSG00000032396 | *Dis3l* | ENSMUSG00000102478 | *BC085271* |
| ENSMUSG00000032397 | *Tipin* | ENSMUSG00000102483 | *Catspere1* |
| ENSMUSG00000032398 | *Snapc5* | ENSMUSG00000102550 | *Gm8276* |
| ENSMUSG00000032399 | *Rpl4* | ENSMUSG00000102609 | *Gm18432* |
| ENSMUSG00000032405 | *Pias1* | ENSMUSG00000102636 | *Gm31466* |
| ENSMUSG00000032407 | *U2surp* | ENSMUSG00000102697 | *Pcdhac2* |
| ENSMUSG00000032409 | *Atr* | ENSMUSG00000102735 | *Gm7369* |
| ENSMUSG00000032412 | *Atp1b3* | ENSMUSG00000102740 | *Gm37599* |
| ENSMUSG00000032418 | *Me1* | ENSMUSG00000102836 | *Gm38666* |
| ENSMUSG00000032423 | *Syncrip* | ENSMUSG00000102869 | *Norad* |
| ENSMUSG00000032425 | *Zfp949* | ENSMUSG00000102976 | *Zc3h11a* |
| ENSMUSG00000032435 | *Dync1li1* | ENSMUSG00000103034 | *Gm8797* |
| ENSMUSG00000032437 | *Stt3b* | ENSMUSG00000103092 | *Pcdha5* |
| ENSMUSG00000032456 | *Nmnat3* | ENSMUSG00000103124 | *Gm37389* |
| ENSMUSG00000032463 | *Faim* | ENSMUSG00000103125 | *Gm37388* |
| ENSMUSG00000032479 | *Map4* | ENSMUSG00000103255 | *Pcdhac1* |
| ENSMUSG00000032481 | *Smarcc1* | ENSMUSG00000103310 | *Pcdha12* |
| ENSMUSG00000032489 | *Kif9* | ENSMUSG00000103433 | *Gm2272* |
| ENSMUSG00000032497 | *Lrrfip2* | ENSMUSG00000103442 | *Pcdha1* |
| ENSMUSG00000032512 | *Wdr48* | ENSMUSG00000103458 | *Gm37013* |
| ENSMUSG00000032514 | *Ttc21a* | ENSMUSG00000103548 | *Gm37670* |
| ENSMUSG00000032518 | *Rpsa* | ENSMUSG00000103705 | *4930518J20Rik* |
| ENSMUSG00000032525 | *Nktr* | ENSMUSG00000103707 | *Pcdha6* |
| ENSMUSG00000032534 | *Cep63* | ENSMUSG00000103760 | *Gm8009* |
| ENSMUSG00000032536 | *Trak1* | ENSMUSG00000103770 | *Pcdha9* |
| ENSMUSG00000032540 | *Abhd5* | ENSMUSG00000103792 | *Gm3745* |
| ENSMUSG00000032560 | *Dnajc13* | ENSMUSG00000103800 | *Pcdha8* |
| ENSMUSG00000032562 | *Gnai2* | ENSMUSG00000103845 | *Gm19026* |
| ENSMUSG00000032575 | *Manf* | ENSMUSG00000103921 | *Gm5841* |
| ENSMUSG00000032580 | *Rbm5* | ENSMUSG00000103922 | *Gm6123* |
| ENSMUSG00000032582 | *Rbm6* | ENSMUSG00000103948 | *4930594C11Rik* |
| ENSMUSG00000032594 | *Ip6k1* | ENSMUSG00000103986 | *Gm5539* |
| ENSMUSG00000032599 | *Ip6k2* | ENSMUSG00000104141 | *Gm6140* |
| ENSMUSG00000032601 | *Prkar2a* | ENSMUSG00000104148 | *Pcdha2* |
| ENSMUSG00000032611 | *1700102P08Rik* | ENSMUSG00000104252 | *Pcdha4* |
| ENSMUSG00000032612 | *Usp4* | ENSMUSG00000104318 | *Pcdha7* |
| ENSMUSG00000032621 | *Srek1* | ENSMUSG00000104529 | *Rbakdn* |
| ENSMUSG00000032637 | *Atxn2l* | ENSMUSG00000104627 | *Mir3535* |
| ENSMUSG00000032666 | *1700025G04Rik* | ENSMUSG00000104777 | *Gm4865* |
| ENSMUSG00000032671 | *A930018P22Rik* | ENSMUSG00000104892 | *Gm6615* |
| ENSMUSG00000032726 | *Bmp8a* | ENSMUSG00000104896 | *Rnu3b4* |
| ENSMUSG00000032727 | *Mier3* | ENSMUSG00000104913 | *Gm6560* |
| ENSMUSG00000032740 | *Ccdc88a* | ENSMUSG00000105103 | *Gm43191* |
| ENSMUSG00000032743 | *Katnip* | ENSMUSG00000105167 | *Snord43* |
| ENSMUSG00000032745 | *Gpbp1* | ENSMUSG00000105181 | *Gm19620* |
| ENSMUSG00000032777 | *Gtf3c1* | ENSMUSG00000105185 | *4930519H02Rik* |
| ENSMUSG00000032782 | *Cntrob* | ENSMUSG00000105223 | *Gm43433* |

| | | | |
|---|---|---|---|
| ENSMUSG00000032803 | *Cdv3* | ENSMUSG00000105340 | *Gm42878* |
| ENSMUSG00000032826 | *Ank2* | ENSMUSG00000105384 | *Gm43396* |
| ENSMUSG00000032827 | *Ppp1r9a* | ENSMUSG00000105390 | *Gm8458* |
| ENSMUSG00000032872 | *Cyb5r4* | ENSMUSG00000105531 | *4930458A03Rik* |
| ENSMUSG00000032883 | *Acsl3* | ENSMUSG00000105662 | *Gm6639* |
| ENSMUSG00000032889 | *Gm6685* | ENSMUSG00000105704 | *Gm43055* |
| ENSMUSG00000032921 | *Odf4* | ENSMUSG00000105705 | *ENSMUSG00000105705* |
| ENSMUSG00000032959 | *Pebp1* | ENSMUSG00000105787 | *Gm8099* |
| ENSMUSG00000032965 | *Ift57* | ENSMUSG00000105875 | *Gm43518* |
| ENSMUSG00000033004 | *Mycbp2* | ENSMUSG00000105975 | *Gm9831* |
| ENSMUSG00000033014 | *Trim33* | ENSMUSG00000106013 | *4933402J10Rik* |
| ENSMUSG00000033020 | *Polr2f* | ENSMUSG00000106037 | *Gm4332* |
| ENSMUSG00000033029 | *1700088E04Rik* | ENSMUSG00000106059 | *Gm9364* |
| ENSMUSG00000033032 | *Afap1l1* | ENSMUSG00000106106 | *CT010467.1* |
| ENSMUSG00000033039 | *Micall1* | ENSMUSG00000106135 | *Gm6543* |
| ENSMUSG00000033053 | *Cfap95* | ENSMUSG00000106147 | *Snord3a* |
| ENSMUSG00000033066 | *Gas7* | ENSMUSG00000106190 | *Gm20768* |
| ENSMUSG00000033099 | *Nol12* | ENSMUSG00000106204 | *Gm21009* |
| ENSMUSG00000033106 | *Slc7a6os* | ENSMUSG00000106239 | *Gm9260* |
| ENSMUSG00000033161 | *Atp1a1* | ENSMUSG00000106246 | *Gm18867* |
| ENSMUSG00000033166 | *Dis3* | ENSMUSG00000106306 | *4933401H06Rik* |
| ENSMUSG00000033207 | *Mamdc2* | ENSMUSG00000106445 | *Gm21190* |
| ENSMUSG00000033209 | *Ttc28* | ENSMUSG00000106499 | *Gm9403* |
| ENSMUSG00000033228 | *Scaf11* | ENSMUSG00000106586 | *1700013M08Rik* |
| ENSMUSG00000033233 | *Trim45* | ENSMUSG00000106622 | *Gm5554* |
| ENSMUSG00000033282 | *Rpgrip1l* | ENSMUSG00000106627 | *Gm21680* |
| ENSMUSG00000033306 | *Lpp* | ENSMUSG00000106740 | *Gm21221* |
| ENSMUSG00000033319 | *Fem1c* | ENSMUSG00000106892 | *Gm42791* |
| ENSMUSG00000033323 | *Ctdp1* | ENSMUSG00000106895 | *Gm4754* |
| ENSMUSG00000033326 | *Kdm4a* | ENSMUSG00000106917 | *Gm7832* |
| ENSMUSG00000033335 | *Dnm2* | ENSMUSG00000107008 | *Gm2762* |
| ENSMUSG00000033364 | *Usp37* | ENSMUSG00000107023 | *Gm42715* |
| ENSMUSG00000033368 | *Trim69* | ENSMUSG00000107035 | *Ybx1-ps2* |
| ENSMUSG00000033409 | *Syce1l* | ENSMUSG00000107053 | *1700021F13Rik* |
| ENSMUSG00000033444 | *Specc1l* | ENSMUSG00000107068 | *Gm42742* |
| ENSMUSG00000033486 | *Catsper2* | ENSMUSG00000107256 | *Gm13821* |
| ENSMUSG00000033487 | *Fndc3a* | ENSMUSG00000107257 | *Gm43028* |
| ENSMUSG00000033502 | *Cdc14a* | ENSMUSG00000107369 | *Gstm2-ps1* |
| ENSMUSG00000033510 | *Otud7a* | ENSMUSG00000107383 | *Gm4366* |
| ENSMUSG00000033543 | *Gtf2a2* | ENSMUSG00000107411 | *Gm19040* |
| ENSMUSG00000033577 | *Myo6* | ENSMUSG00000107453 | *Gm44145* |
| ENSMUSG00000033610 | *Pank1* | ENSMUSG00000107462 | *Gm7384* |
| ENSMUSG00000033624 | *Pdpr* | ENSMUSG00000107482 | *Etfbl* |
| ENSMUSG00000033632 | *AW554918* | ENSMUSG00000107484 | *Nasp-ps1* |
| ENSMUSG00000033644 | *Piwil2* | ENSMUSG00000107501 | *Gm29781* |
| ENSMUSG00000033658 | *Ddx19b* | ENSMUSG00000107596 | *Gm18913* |
| ENSMUSG00000033671 | *Cep350* | ENSMUSG00000107639 | *Gm4374* |
| ENSMUSG00000033701 | *Acbd6* | ENSMUSG00000107681 | *4930528H21Rik* |
| ENSMUSG00000033705 | *Stard9* | ENSMUSG00000107747 | *Gm5881* |
| ENSMUSG00000033712 | *Ccar2* | ENSMUSG00000107817 | *Gm38840* |
| ENSMUSG00000033713 | *Foxn3* | ENSMUSG00000107836 | *Gm5315* |
| ENSMUSG00000033739 | *Fkbpl* | ENSMUSG00000107872 | *Gm44511* |
| ENSMUSG00000033751 | *Gadd45gip1* | ENSMUSG00000107877 | *4933427D14Rik* |
| ENSMUSG00000033760 | *Rbm4b* | ENSMUSG00000107879 | *9330102E08Rik* |
| ENSMUSG00000033762 | *Recql4* | ENSMUSG00000107921 | *Gm6423* |
| ENSMUSG00000033769 | *Exoc6b* | ENSMUSG00000107928 | *Gm45140* |
| ENSMUSG00000033793 | *Atp6v1h* | ENSMUSG00000108078 | *Gm8744* |
| ENSMUSG00000033794 | *Lpcat2b* | ENSMUSG00000108112 | *Gm45193* |

| | | | |
|---|---|---|---|
| ENSMUSG00000033808 | *Tmem87a* | ENSMUSG00000108145 | *Gm38811* |
| ENSMUSG00000033813 | *Tcea1* | ENSMUSG00000108227 | *4933412L11Rik* |
| ENSMUSG00000033826 | *Dnah8* | ENSMUSG00000108366 | *Gm5586* |
| ENSMUSG00000033882 | *Rbm46* | ENSMUSG00000108401 | *Rpl19-ps10* |
| ENSMUSG00000033885 | *Pxk* | ENSMUSG00000108414 | *Snhg1* |
| ENSMUSG00000033902 | *Mapkbp1* | ENSMUSG00000108448 | *Gm21269* |
| ENSMUSG00000033904 | *Ccp110* | ENSMUSG00000108615 | *Gm4973* |
| ENSMUSG00000033909 | *Usp36* | ENSMUSG00000108660 | *Gm21284* |
| ENSMUSG00000033916 | *Chmp2a* | ENSMUSG00000108702 | *Gm9333* |
| ENSMUSG00000033933 | *Vhl* | ENSMUSG00000108813 | *Rpl19-ps9* |
| ENSMUSG00000033938 | *Ndufb7* | ENSMUSG00000108815 | *Gm49388* |
| ENSMUSG00000033943 | *Mga* | ENSMUSG00000108833 | *Gm17909* |
| ENSMUSG00000033949 | *Trim36* | ENSMUSG00000108878 | *Gm49493* |
| ENSMUSG00000033960 | *Jcad* | ENSMUSG00000108929 | *Cc2d2b* |
| ENSMUSG00000033983 | *Coil* | ENSMUSG00000108946 | *Gm18598* |
| ENSMUSG00000033985 | *Tesk2* | ENSMUSG00000108963 | *Gm16478* |
| ENSMUSG00000033987 | *Dnah17* | ENSMUSG00000109061 | *Gm49320* |
| ENSMUSG00000034007 | *Scaper* | ENSMUSG00000109113 | *Gm32916* |
| ENSMUSG00000034021 | *Pds5b* | ENSMUSG00000109116 | *Sycp1-ps1* |
| ENSMUSG00000034022 | *Cpsf1* | ENSMUSG00000109156 | *Gm45194* |
| ENSMUSG00000034023 | *Fancd2* | ENSMUSG00000109179 | *Gm35339* |
| ENSMUSG00000034024 | *Cct2* | ENSMUSG00000109181 | *Gm5605* |
| ENSMUSG00000034063 | *4930590J08Rik* | ENSMUSG00000109222 | *Gm10297* |
| ENSMUSG00000034083 | *Ccdc174* | ENSMUSG00000109224 | *Tmem147os* |
| ENSMUSG00000034088 | *Hdlbp* | ENSMUSG00000109228 | *Fam81b* |
| ENSMUSG00000034101 | *Ctnnd1* | ENSMUSG00000109274 | *Gm45133* |
| ENSMUSG00000034120 | *Srsf2* | ENSMUSG00000109326 | *Gm9165* |
| ENSMUSG00000034121 | *Mks1* | ENSMUSG00000109561 | *Ankrd31* |
| ENSMUSG00000034126 | *Pomt2* | ENSMUSG00000109565 | *Gm6579* |
| ENSMUSG00000034135 | *Sik3* | ENSMUSG00000109572 | *Cfap99* |
| ENSMUSG00000034151 | *Zbbx* | ENSMUSG00000109586 | *Gm6853* |
| ENSMUSG00000034154 | *Ino80* | ENSMUSG00000109657 | *1700008N11Rik* |
| ENSMUSG00000034163 | *Zfc3h1* | ENSMUSG00000109695 | *Gm31166* |
| ENSMUSG00000034194 | *R3hcc1* | ENSMUSG00000109737 | *Spem3* |
| ENSMUSG00000034206 | *Polq* | ENSMUSG00000109739 | *Gm17949* |
| ENSMUSG00000034212 | *Ankmy1* | ENSMUSG00000109750 | *Hmgb1-rs17* |
| ENSMUSG00000034219 | *Septin14* | ENSMUSG00000109757 | *Gm7128* |
| ENSMUSG00000034239 | *Gm884* | ENSMUSG00000109789 | *Gm2033* |
| ENSMUSG00000034243 | *Golgb1* | ENSMUSG00000109830 | *Gm7965* |
| ENSMUSG00000034245 | *Hdac11* | ENSMUSG00000109843 | *Gm17841* |
| ENSMUSG00000034252 | *Senp6* | ENSMUSG00000109864 | *Eid3* |
| ENSMUSG00000034269 | *Setd5* | ENSMUSG00000109894 | *Gm5904* |
| ENSMUSG00000034274 | *Thoc5* | ENSMUSG00000109933 | *Gm7600* |
| ENSMUSG00000034292 | *Traf3ip1* | ENSMUSG00000110170 | *St6galnac2* |
| ENSMUSG00000034297 | *Med13* | ENSMUSG00000110234 | *Gm45799* |
| ENSMUSG00000034303 | *Ccdc15* | ENSMUSG00000110267 | *Gm45283* |
| ENSMUSG00000034317 | *Trim59* | ENSMUSG00000110275 | *Gm5905* |
| ENSMUSG00000034342 | *Cbl* | ENSMUSG00000110331 | *Nudc-ps1* |
| ENSMUSG00000034349 | *Smc4* | ENSMUSG00000110333 | *Gm45861* |
| ENSMUSG00000034354 | *Mtmr3* | ENSMUSG00000110336 | *4933400L20Rik* |
| ENSMUSG00000034371 | *Tkfc* | ENSMUSG00000110416 | *Gm19248* |
| ENSMUSG00000034377 | *Tulp4* | ENSMUSG00000110448 | *Gm33242* |
| ENSMUSG00000034401 | *Spata6* | ENSMUSG00000110455 | *Gm45904* |
| ENSMUSG00000034403 | *Pja1* | ENSMUSG00000110587 | *Gm7344* |
| ENSMUSG00000034462 | *Pkd2* | ENSMUSG00000110610 | *Gm7769* |
| ENSMUSG00000034480 | *Diaph2* | ENSMUSG00000110617 | *Gm5910* |
| ENSMUSG00000034485 | *Uaca* | ENSMUSG00000110622 | *Iqcn* |
| ENSMUSG00000034518 | *Hmgxb4* | ENSMUSG00000110737 | *4930444F02Rik* |

| | | | |
|---|---|---|---|
| ENSMUSG00000034543 | *Morc2a* | ENSMUSG00000110768 | *Gm18541* |
| ENSMUSG00000034544 | *Rsrc1* | ENSMUSG00000110841 | *Gpx4-ps2* |
| ENSMUSG00000034563 | *Ccpg1* | ENSMUSG00000110847 | *Gm18705* |
| ENSMUSG00000034573 | *Ptpn13* | ENSMUSG00000110882 | *Gm7642* |
| ENSMUSG00000034574 | *Daam1* | ENSMUSG00000110926 | *Gm5917* |
| ENSMUSG00000034593 | *Myo5a* | ENSMUSG00000110989 | *Gm7444* |
| ENSMUSG00000034610 | *Tut4* | ENSMUSG00000111108 | *Gm17795* |
| ENSMUSG00000034621 | *Gpatch8* | ENSMUSG00000111172 | *Gm39302* |
| ENSMUSG00000034623 | *Prss55* | ENSMUSG00000111209 | *Gm7529* |
| ENSMUSG00000034647 | *Ankrd12* | ENSMUSG00000111224 | *Gm18669* |
| ENSMUSG00000034675 | *Dbn1* | ENSMUSG00000111368 | *Gm47889* |
| ENSMUSG00000034689 | *Spertl* | ENSMUSG00000111410 | *Gm49337* |
| ENSMUSG00000034706 | *Dnai2* | ENSMUSG00000111515 | *Gm5055* |
| ENSMUSG00000034723 | *Tmx4* | ENSMUSG00000111600 | *Gm8049* |
| ENSMUSG00000034731 | *Dgkh* | ENSMUSG00000111601 | *Gm18986* |
| ENSMUSG00000034761 | *Map4k5* | ENSMUSG00000111662 | *Gm7435* |
| ENSMUSG00000034764 | *1700006J14Rik* | ENSMUSG00000111734 | *Gm29825* |
| ENSMUSG00000034795 | *Ccdc122* | ENSMUSG00000111795 | *Gm47620* |
| ENSMUSG00000034813 | *Grip1* | ENSMUSG00000111815 | *Gm6018* |
| ENSMUSG00000034842 | *Art3* | ENSMUSG00000111842 | *Gm49318* |
| ENSMUSG00000034848 | *Ttc21b* | ENSMUSG00000111847 | *Gm8899* |
| ENSMUSG00000034889 | *Cactin* | ENSMUSG00000111877 | *Gm6477* |
| ENSMUSG00000034893 | *Cog3* | ENSMUSG00000111884 | *Gm19612* |
| ENSMUSG00000034898 | *Filip1* | ENSMUSG00000112218 | *Gm18726* |
| ENSMUSG00000034913 | *Cby2* | ENSMUSG00000112252 | *Gm32802* |
| ENSMUSG00000034928 | *Rnf44* | ENSMUSG00000112282 | *Gm19989* |
| ENSMUSG00000034931 | *Dhx8* | ENSMUSG00000112297 | *Gm17816* |
| ENSMUSG00000034994 | *Eef2* | ENSMUSG00000112316 | *Gm48425* |
| ENSMUSG00000035021 | *Baz1a* | ENSMUSG00000112323 | *Gm32828* |
| ENSMUSG00000035024 | *Ncapd3* | ENSMUSG00000112368 | *Gm49376* |
| ENSMUSG00000035032 | *Nek11* | ENSMUSG00000112395 | *Gm7198* |
| ENSMUSG00000035047 | *Kri1* | ENSMUSG00000112418 | *Gm9466* |
| ENSMUSG00000035048 | *Anapc13* | ENSMUSG00000112449 | *Srp54b* |
| ENSMUSG00000035049 | *Rrp12* | ENSMUSG00000112454 | *Gm48619* |
| ENSMUSG00000035051 | *Dhx57* | ENSMUSG00000112491 | *Gm19801* |
| ENSMUSG00000035057 | *Mgat4d* | ENSMUSG00000112515 | *Gm4928* |
| ENSMUSG00000035086 | *Becn1* | ENSMUSG00000112546 | *Gm6992* |
| ENSMUSG00000035125 | *Gcfc2* | ENSMUSG00000112550 | *Gm6627* |
| ENSMUSG00000035126 | *Dnai4* | ENSMUSG00000112657 | *BC106175* |
| ENSMUSG00000035129 | *Gm6781* | ENSMUSG00000112678 | *Gm47492* |
| ENSMUSG00000035133 | *Arhgap5* | ENSMUSG00000112709 | *Gm19395* |
| ENSMUSG00000035152 | *Ap2b1* | ENSMUSG00000112735 | *Gm46191* |
| ENSMUSG00000035161 | *Ints6* | ENSMUSG00000112795 | *Gm2027* |
| ENSMUSG00000035165 | *Kcne3* | ENSMUSG00000112825 | *Gm9118* |
| ENSMUSG00000035173 | *Ccdc186* | ENSMUSG00000112908 | *Gm7392* |
| ENSMUSG00000035202 | *Lars2* | ENSMUSG00000113014 | *Selenok-ps8* |
| ENSMUSG00000035211 | *Xrra1* | ENSMUSG00000113022 | *Gm9544* |
| ENSMUSG00000035227 | *Spcs2* | ENSMUSG00000113086 | *Gm47376* |
| ENSMUSG00000035247 | *Hectd1* | ENSMUSG00000113168 | *Gm7614* |
| ENSMUSG00000035248 | *Tut7* | ENSMUSG00000113183 | *Gm47664* |
| ENSMUSG00000035268 | *Pkig* | ENSMUSG00000113211 | *4921525O09Rik* |
| ENSMUSG00000035297 | *Cops4* | ENSMUSG00000113255 | *Tes3-ps* |
| ENSMUSG00000035310 | *Lin54* | ENSMUSG00000113263 | *Gm4811* |
| ENSMUSG00000035325 | *Sec31a* | ENSMUSG00000113266 | *Gm19563* |
| ENSMUSG00000035356 | *Nfkbiz* | ENSMUSG00000113285 | *Gm36128* |
| ENSMUSG00000035365 | *Parpbp* | ENSMUSG00000113419 | *Gm5789* |
| ENSMUSG00000035382 | *Pcsk7* | ENSMUSG00000113510 | *Gm7239* |
| ENSMUSG00000035392 | *Dennd1a* | ENSMUSG00000113512 | *Gm9063* |

| | | | |
|---|---|---|---|
| ENSMUSG00000035394 | *Cfap53* | ENSMUSG00000113627 | *Gm32526* |
| ENSMUSG00000035420 | *Fam170a* | ENSMUSG00000113637 | *Gm7049* |
| ENSMUSG00000035437 | *Rabgap1* | ENSMUSG00000113664 | *Gm47184* |
| ENSMUSG00000035458 | *Tnni3* | ENSMUSG00000113790 | *Gm17872* |
| ENSMUSG00000035478 | *Mbd3* | ENSMUSG00000113840 | *Gm18955* |
| ENSMUSG00000035504 | *Reep6* | ENSMUSG00000114034 | *Gm9267* |
| ENSMUSG00000035517 | *Tdrd7* | ENSMUSG00000114046 | *Snapc1l* |
| ENSMUSG00000035522 | *Tsga8* | ENSMUSG00000114066 | *Gm9202* |
| ENSMUSG00000035529 | *Prdm4* | ENSMUSG00000114073 | *Gm30302* |
| ENSMUSG00000035530 | *Eif1* | ENSMUSG00000114124 | *Gm19084* |
| ENSMUSG00000035539 | *Ccdc180* | ENSMUSG00000114133 | *Btf3l4b* |
| ENSMUSG00000035545 | *Leng8* | ENSMUSG00000114167 | *Gm7002* |
| ENSMUSG00000035569 | *Ankrd11* | ENSMUSG00000114170 | *Gm47120* |
| ENSMUSG00000035578 | *Iqcg* | ENSMUSG00000114251 | *Tmed10-ps* |
| ENSMUSG00000035601 | *Trmt10b* | ENSMUSG00000114273 | *Gm34585* |
| ENSMUSG00000035623 | *Rsf1* | ENSMUSG00000114278 | *Gm49027* |
| ENSMUSG00000035642 | *Aamdc* | ENSMUSG00000114361 | *Gm31104* |
| ENSMUSG00000035649 | *Zcchc7* | ENSMUSG00000114409 | *Gm9042* |
| ENSMUSG00000035694 | *Caps2* | ENSMUSG00000114424 | *Gm47414* |
| ENSMUSG00000035726 | *Supt16* | ENSMUSG00000114447 | *Gm47636* |
| ENSMUSG00000035769 | *Xylb* | ENSMUSG00000114557 | *Gm8345* |
| ENSMUSG00000035770 | *Dync1li2* | ENSMUSG00000114558 | *Gm9570* |
| ENSMUSG00000035785 | *Cmtm2b* | ENSMUSG00000114579 | *Gm4130* |
| ENSMUSG00000035790 | *Cep19* | ENSMUSG00000114681 | *1700026N04Rik* |
| ENSMUSG00000035834 | *Polr3g* | ENSMUSG00000114692 | *Gm48157* |
| ENSMUSG00000035842 | *Ddx11* | ENSMUSG00000114722 | *Gm31392* |
| ENSMUSG00000035851 | *Ythdc1* | ENSMUSG00000114763 | *Gm49354* |
| ENSMUSG00000035877 | *Zhx3* | ENSMUSG00000114832 | *Gm21388* |
| ENSMUSG00000035890 | *Rnf126* | ENSMUSG00000114878 | *Gm9086* |
| ENSMUSG00000035901 | *Dennd5a* | ENSMUSG00000114940 | *Gm21378* |
| ENSMUSG00000035910 | *Dcdc2a* | ENSMUSG00000114942 | *Gm49361* |
| ENSMUSG00000035984 | *Nme5* | ENSMUSG00000114951 | *Gm20784* |
| ENSMUSG00000036023 | *Parp2* | ENSMUSG00000114972 | *Gm49119* |
| ENSMUSG00000036045 | *Gm12221* | ENSMUSG00000114999 | *Gm7962* |
| ENSMUSG00000036046 | *5031439G07Rik* | ENSMUSG00000115070 | *Gm4740* |
| ENSMUSG00000036054 | *Sugp2* | ENSMUSG00000115118 | *Gm36617* |
| ENSMUSG00000036087 | *Slain2* | ENSMUSG00000115193 | *Gm6533* |
| ENSMUSG00000036097 | *Slf2* | ENSMUSG00000115194 | *Gm48909* |
| ENSMUSG00000036104 | *Rab3gap1* | ENSMUSG00000115195 | *Gm5671* |
| ENSMUSG00000036112 | *Metap2* | ENSMUSG00000115232 | *Gm49378* |
| ENSMUSG00000036155 | *Mgat5* | ENSMUSG00000115248 | *Gm49037* |
| ENSMUSG00000036160 | *Surf6* | ENSMUSG00000115271 | *4930529K09Rik* |
| ENSMUSG00000036168 | *Ccdc38* | ENSMUSG00000115286 | *4930429C20Rik* |
| ENSMUSG00000036180 | *Gatad2a* | ENSMUSG00000115312 | *Gm8518* |
| ENSMUSG00000036197 | *Gxylt1* | ENSMUSG00000115392 | *Gm17882* |
| ENSMUSG00000036199 | *Ndufa13* | ENSMUSG00000115411 | *Gm6986* |
| ENSMUSG00000036202 | *Rif1* | ENSMUSG00000115420 | *Rmrp* |
| ENSMUSG00000036211 | *H1f6* | ENSMUSG00000115448 | *Gm21178* |
| ENSMUSG00000036214 | *Polr1has* | ENSMUSG00000115463 | *Gm49356* |
| ENSMUSG00000036241 | *Ube2r2* | ENSMUSG00000115546 | *Gm49077* |
| ENSMUSG00000036257 | *Pnpla8* | ENSMUSG00000115571 | *Gm6616* |
| ENSMUSG00000036275 | *9530068E07Rik* | ENSMUSG00000115623 | *Gm41225* |
| ENSMUSG00000036281 | *Snapc4* | ENSMUSG00000115632 | *Gm17922* |
| ENSMUSG00000036292 | *Gramd1c* | ENSMUSG00000115700 | *Gm7517* |
| ENSMUSG00000036323 | *Srp72* | ENSMUSG00000115743 | *Gm7004* |
| ENSMUSG00000036333 | *Kidins220* | ENSMUSG00000115858 | *4930588J15Rik* |
| ENSMUSG00000036352 | *Ubac1* | ENSMUSG00000116016 | *Gm49496* |
| ENSMUSG00000036368 | *Rmdn2* | ENSMUSG00000116090 | *Rpl19-ps6* |

| | | | |
|---|---|---|---|
| ENSMUSG00000036371 | *Serbp1* | ENSMUSG00000116096 | *4933404G15Rik* |
| ENSMUSG00000036377 | *Cracd* | ENSMUSG00000116207 | *Nnt* |
| ENSMUSG00000036398 | *Ppp1r11* | ENSMUSG00000116244 | *Gm49461* |
| ENSMUSG00000036403 | *Cep135* | ENSMUSG00000116260 | *1700041B01Rik* |
| ENSMUSG00000036427 | *Gpi1* | ENSMUSG00000116275 | *Zc3h11a* |
| ENSMUSG00000036435 | *Exoc1* | ENSMUSG00000116284 | *Gm3787* |
| ENSMUSG00000036438 | *Calm2* | ENSMUSG00000116343 | *Gm6847* |
| ENSMUSG00000036452 | *Arhgap26* | ENSMUSG00000116347 | *Gm49416* |
| ENSMUSG00000036461 | *Elf1* | ENSMUSG00000116348 | *Gm18722* |
| ENSMUSG00000036463 | *4930544G11Rik* | ENSMUSG00000116376 | *Tmem249* |
| ENSMUSG00000036478 | *Btg1* | ENSMUSG00000116423 | *Gm7085* |
| ENSMUSG00000036499 | *Eea1* | ENSMUSG00000116461 | *Gm44502* |
| ENSMUSG00000036501 | *Fam13b* | ENSMUSG00000116520 | *4930414F18Rik* |
| ENSMUSG00000036510 | *Cdh8* | ENSMUSG00000116550 | *Gm3507* |
| ENSMUSG00000036528 | *Ppfibp2* | ENSMUSG00000116587 | *Gm49753* |
| ENSMUSG00000036533 | *Cdc42ep3* | ENSMUSG00000116594 | *Gm49601* |
| ENSMUSG00000036550 | *Cnot1* | ENSMUSG00000116615 | *4930548J01Rik* |
| ENSMUSG00000036555 | *Iqce* | ENSMUSG00000116622 | *Gm5221* |
| ENSMUSG00000036561 | *Ppp6r2* | ENSMUSG00000116676 | *Gm5768* |
| ENSMUSG00000036572 | *Upf3b* | ENSMUSG00000116689 | *4930405D01Rik* |
| ENSMUSG00000036580 | *Spg20* | ENSMUSG00000116743 | *Gm49573* |
| ENSMUSG00000036598 | *Ccdc113* | ENSMUSG00000116751 | *Gm49793* |
| ENSMUSG00000036606 | *Plxnb2* | ENSMUSG00000116780 | *Dynlt2a3* |
| ENSMUSG00000036617 | *Etl4* | ENSMUSG00000116795 | *Gm49600* |
| ENSMUSG00000036641 | *Ccdc148* | ENSMUSG00000116817 | *Gm5679* |
| ENSMUSG00000036646 | *Man1b1* | ENSMUSG00000116838 | *Gm6475* |
| ENSMUSG00000036693 | *Nop14* | ENSMUSG00000116879 | *Gm49794* |
| ENSMUSG00000036707 | *Cab39* | ENSMUSG00000116892 | *Gm49787* |
| ENSMUSG00000036743 | *Psma8* | ENSMUSG00000116908 | *Gm49599* |
| ENSMUSG00000036752 | *Tubb4b* | ENSMUSG00000116912 | *Gm5487* |
| ENSMUSG00000036768 | *Kif15* | ENSMUSG00000116925 | *Gm49776* |
| ENSMUSG00000036777 | *Anln* | ENSMUSG00000116949 | *Gm7177* |
| ENSMUSG00000036779 | *Tent4b* | ENSMUSG00000116972 | *Gm6278* |
| ENSMUSG00000036817 | *Sun1* | ENSMUSG00000116976 | *4930500H12Rik* |
| ENSMUSG00000036822 | *Topors* | ENSMUSG00000117008 | *Gm49817* |
| ENSMUSG00000036825 | *Ssx2ip* | ENSMUSG00000117049 | *Gm6540* |
| ENSMUSG00000036872 | *Abcc12* | ENSMUSG00000117071 | *Gm9598* |
| ENSMUSG00000036873 | *2410004B18Rik* | ENSMUSG00000117114 | *Gm49867* |
| ENSMUSG00000036882 | *Arhgap33* | ENSMUSG00000117120 | *Gm50238* |
| ENSMUSG00000036918 | *Ttc7* | ENSMUSG00000117197 | *Gm6618* |
| ENSMUSG00000036928 | *Stag3* | ENSMUSG00000117274 | *Gm17921* |
| ENSMUSG00000036934 | *4921524J17Rik* | ENSMUSG00000117286 | *Gm1043* |
| ENSMUSG00000036940 | *Kdm1a* | ENSMUSG00000117299 | *Gm9288* |
| ENSMUSG00000036955 | *Kifbp* | ENSMUSG00000117338 | *Gm49804* |
| ENSMUSG00000036964 | *Trim17* | ENSMUSG00000117340 | *Gm49920* |
| ENSMUSG00000036966 | *Spryd3* | ENSMUSG00000117416 | *Gm5064* |
| ENSMUSG00000036990 | *Otud4* | ENSMUSG00000117425 | *Gm7464* |
| ENSMUSG00000037001 | *Zfp39* | ENSMUSG00000117477 | *Gm50092* |
| ENSMUSG00000037012 | *Hk1* | ENSMUSG00000117507 | *Gm50045* |
| ENSMUSG00000037017 | *Zscan21* | ENSMUSG00000117518 | *Gpx4-ps1* |
| ENSMUSG00000037020 | *Wdr62* | ENSMUSG00000117563 | *Gm7575* |
| ENSMUSG00000037058 | *Paip2* | ENSMUSG00000117572 | *Gm9410* |
| ENSMUSG00000037062 | *Sh3glb1* | ENSMUSG00000117606 | *Gm5969* |
| ENSMUSG00000037075 | *Rnf139* | ENSMUSG00000117618 | *4930578E11Rik* |
| ENSMUSG00000037098 | *Rab11fip3* | ENSMUSG00000117673 | *Gm6974* |
| ENSMUSG00000037101 | *Ttc29* | ENSMUSG00000117694 | *Snhg4* |
| ENSMUSG00000037108 | *Zcwpw1* | ENSMUSG00000117720 | *Gm5249* |
| ENSMUSG00000037110 | *Ralgapa2* | ENSMUSG00000117725 | *Gm50240* |

| | | | |
|---|---|---|---|
| ENSMUSG00000037124 | *Trim58* | ENSMUSG00000117732 | *Gm50364* |
| ENSMUSG00000037138 | *Aff3* | ENSMUSG00000117771 | *Gm25432* |
| ENSMUSG00000037148 | *Arhgap10* | ENSMUSG00000117789 | *Gm50388* |
| ENSMUSG00000037149 | *Ddx1* | ENSMUSG00000117804 | *Gm6252* |
| ENSMUSG00000037152 | *Ndufc1* | ENSMUSG00000117815 | *Gm50166* |
| ENSMUSG00000037166 | *Ppp1r14a* | ENSMUSG00000117819 | *Gm50253* |
| ENSMUSG00000037174 | *Elf2* | ENSMUSG00000117822 | *Eef1a1-ps1* |
| ENSMUSG00000037196 | *Pacrg* | ENSMUSG00000117841 | *Gm50371* |
| ENSMUSG00000037197 | *Rbm17* | ENSMUSG00000117852 | *4930471G03Rik* |
| ENSMUSG00000037210 | *Fam193a* | ENSMUSG00000117874 | *Gm50318* |
| ENSMUSG00000037214 | *Thap1* | ENSMUSG00000117877 | *Gm8184* |
| ENSMUSG00000037234 | *Hook3* | ENSMUSG00000117929 | *Gm6663* |
| ENSMUSG00000037236 | *Matr3* | ENSMUSG00000117939 | *Gm9067* |
| ENSMUSG00000037243 | *Zfp692* | ENSMUSG00000117942 | *Maskbp3* |
| ENSMUSG00000037262 | *Kin* | ENSMUSG00000117975 | *Itprip* |
| ENSMUSG00000037266 | *Rsrp1* | ENSMUSG00000117988 | *Gm8663* |
| ENSMUSG00000037270 | *4932438A13Rik* | ENSMUSG00000118036 | *Gm6386* |
| ENSMUSG00000037286 | *Stag1* | ENSMUSG00000118059 | *Gm6975* |
| ENSMUSG00000037307 | *Banf2* | ENSMUSG00000118124 | *Gm50139* |
| ENSMUSG00000037313 | *Tacc3* | ENSMUSG00000118135 | *Gm50367* |
| ENSMUSG00000037331 | *Larp1* | ENSMUSG00000118203 | *Gm7798* |
| ENSMUSG00000037343 | *Taf2* | ENSMUSG00000118252 | *Gm5521* |
| ENSMUSG00000037355 | *Uvssa* | ENSMUSG00000118267 | *G630055G22Rik* |
| ENSMUSG00000037361 | *Sf3b6* | ENSMUSG00000118274 | *Gm50204* |
| ENSMUSG00000037364 | *Srrt* | ENSMUSG00000118280 | *Gm41804* |
| ENSMUSG00000037376 | *Trmt6* | ENSMUSG00000118341 | *Gm50252* |
| ENSMUSG00000037386 | *Rims2* | ENSMUSG00000118353 | *Nsa2-ps1* |
| ENSMUSG00000037395 | *Rcor3* | ENSMUSG00000118377 | *Gm9276* |
| ENSMUSG00000037432 | *Fer1l5* | ENSMUSG00000118396 | *4930517N10Rik* |
| ENSMUSG00000037437 | *Adam32* | ENSMUSG00000118454 | *Wdr88* |
| ENSMUSG00000037438 | *Uqcrh-ps1* | ENSMUSG00000118491 | *Gm44505* |
| ENSMUSG00000037443 | *Cep85* | ENSMUSG00000118504 | *Gm53012* |
| ENSMUSG00000037458 | *Azin1* | ENSMUSG00000118506 | *Cfap141* |
| ENSMUSG00000037470 | *Uggt1* | ENSMUSG00000118519 | *Gm52992* |
| ENSMUSG00000037475 | *Thoc2* | ENSMUSG00000118523 | *Gm34319* |
| ENSMUSG00000037486 | *Asxl2* | ENSMUSG00000118552 | *Rplp2-ps1* |
