## Supplemental Table 2 for "Splicing factor SRSF1 is essential for homing of precursor spermatogonial stem cells in mice"

**Table S2. Differentially expressed genes were analysed in this study.**

| gene id | cKO1 | cKO2 | cKO3 | Control1 | Control2 | Control3 | cKO | Ctrl | log2Fold Change | pvalue | padj | gene name | gene chr |
|---|---|---|---|---|---|---|---|---|---|---|---|---|---|
| ENSMUSG00000031780 | 271.85 | 308.52 | 478.91 | 59.178 | 40.766 | 72.675 | 353.09 | 57.54 | 2.6226 | 5.93E-23 | 1.07E-18 | *Ccl17* | 8 |
| ENSMUSG00000063171 | 983.48 | 838.53 | 975.99 | 1941.7 | 2948.2 | 2342.5 | 932.67 | 2410.8 | -1.3707 | 6.71E-19 | 6.07E-15 | *Rps4l* | 6 |
| ENSMUSG00000031397 | 40.935 | 83.062 | 59.864 | 287.73 | 628.84 | 283.73 | 61.287 | 400.1 | -2.706 | 1.26E-18 | 7.59E-15 | *Tktl1* | X |
| ENSMUSG00000020738 | 1406.5 | 1023.4 | 1270 | 2611 | 2830.2 | 2965.7 | 1233.3 | 2802.3 | -1.1845 | 3.09E-18 | 1.40E-14 | *Sumo2* | 11 |
| ENSMUSG00000082033 | 40.935 | 58.341 | 47.036 | 216.31 | 588.07 | 229.97 | 48.771 | 344.78 | -2.8222 | 1.72E-17 | 6.21E-14 | *Mageb10-ps* | X |
| ENSMUSG00000025480 | 30.439 | 64.274 | 52.381 | 224.47 | 497.87 | 236.94 | 49.031 | 319.76 | -2.705 | 2.41E-17 | 7.27E-14 | *Syce1* | 7 |
| ENSMUSG00000041912 | 128.05 | 183.92 | 153.94 | 422.41 | 823.13 | 503.75 | 155.3 | 583.1 | -1.9091 | 3.15E-16 | 7.35E-13 | *Tdrkh* | 3 |
| ENSMUSG00000075511 | 53.53 | 106.79 | 78.037 | 305.07 | 467.51 | 282.74 | 79.454 | 351.77 | -2.1455 | 3.25E-16 | 7.35E-13 | *1700001L05Rik* | 15 |
| ENSMUSG00000044595 | 114.41 | 86.029 | 95.14 | 295.89 | 520.42 | 322.56 | 98.525 | 379.62 | -1.9486 | 4.86E-16 | 8.65E-13 | *Dnd1* | 18 |
| ENSMUSG00000021098 | 152.19 | 189.86 | 175.31 | 416.29 | 664.4 | 503.75 | 172.45 | 528.15 | -1.6155 | 5.20E-16 | 8.65E-13 | *4930447C04Rik* | 12 |
| ENSMUSG00000044303 | 52.48 | 79.107 | 55.588 | 232.63 | 409.4 | 230.97 | 62.392 | 291 | -2.2212 | 5.26E-16 | 8.65E-13 | *Cdkn2a* | 4 |
| ENSMUSG00000018379 | 4251.9 | 4365.7 | 4869.3 | 7745.2 | 8402.2 | 8339.7 | 4495.6 | 8162.4 | -0.8605 | 1.47E-15 | 2.16E-12 | *Srsf1* | 11 |
| ENSMUSG00000031765 | 424.04 | 587.37 | 654.22 | 1233.6 | 1995.8 | 1471.4 | 555.21 | 1566.9 | -1.4973 | 1.55E-15 | 2.16E-12 | *Mt1* | 8 |
| ENSMUSG00000051920 | 3.1488 | 9.8884 | 3.207 | 78.564 | 154.39 | 70.684 | 5.4147 | 101.21 | -4.2099 | 2.53E-15 | 3.27E-12 | *Rspo2* | 15 |
| ENSMUSG00000046774 | 40.935 | 69.218 | 54.519 | 243.86 | 514.35 | 194.13 | 54.891 | 317.44 | -2.5321 | 3.55E-15 | 4.28E-12 | *8030474K03Rik* | X |
| ENSMUSG00000031384 | 9.4464 | 19.777 | 17.104 | 86.727 | 209.03 | 120.46 | 15.442 | 138.74 | -3.1672 | 5.28E-15 | 5.96E-12 | *Asb9* | X |
| ENSMUSG00000108981 | 112.31 | 165.14 | 101.55 | 404.05 | 667.87 | 356.41 | 126.33 | 476.11 | -1.9136 | 1.32E-14 | 1.40E-11 | *Gm15262* | X |
| ENSMUSG00000073894 | 39.885 | 95.917 | 82.312 | 278.55 | 510.01 | 267.8 | 72.705 | 352.12 | -2.2758 | 2.63E-14 | 2.64E-11 | *Rbmxl2* | 7 |
| ENSMUSG00000028434 | 78.72 | 108.77 | 109.04 | 283.65 | 546.44 | 303.64 | 98.843 | 377.91 | -1.9363 | 5.63E-14 | 5.36E-11 | *Epb41l4b* | 4 |
| ENSMUSG00000046637 | 2.0992 | 18.788 | 8.5519 | 71.422 | 189.95 | 118.47 | 9.813 | 126.62 | -3.6791 | 6.35E-14 | 5.74E-11 | *Ttc34* | 4 |
| ENSMUSG00000031430 | 64.026 | 107.78 | 42.76 | 297.93 | 435.42 | 234.95 | 71.523 | 322.77 | -2.1715 | 2.10E-13 | 1.81E-10 | *Vsig1* | X |
| ENSMUSG00000068245 | 35.687 | 34.609 | 31.001 | 160.19 | 229.85 | 113.49 | 33.766 | 167.84 | -2.3153 | 2.65E-13 | 2.18E-10 | *Phf11d* | 14 |
| ENSMUSG00000033007 | 385.2 | 402.46 | 375.22 | 156.11 | 117.09 | 171.23 | 387.63 | 148.15 | 1.3911 | 3.25E-13 | 2.50E-10 | *Asic4* | 1 |
| ENSMUSG00000084838 | 1450.6 | 1562.4 | 1406.8 | 682.59 | 843.95 | 606.29 | 1473.2 | 710.94 | 1.0502 | 3.31E-13 | 2.50E-10 | *Gm10241* | 16 |
| ENSMUSG00000030041 | 26.24 | 40.542 | 37.415 | 162.23 | 324.39 | 121.46 | 34.732 | 202.69 | -2.5463 | 3.70E-13 | 2.68E-10 | *M1ap* | 6 |
| ENSMUSG00000074397 | 22.042 | 26.699 | 21.38 | 122.44 | 294.04 | 88.604 | 23.373 | 168.36 | -2.85 | 1.50E-12 | 1.02E-09 | *Foxr1* | 9 |
| ENSMUSG00000028610 | 5.248 | 6.9218 | 7.483 | 59.178 | 148.32 | 56.746 | 6.5509 | 88.081 | -3.7529 | 1.55E-12 | 1.02E-09 | *Dmrtb1* | 4 |
| ENSMUSG00000028591 | 37.786 | 58.341 | 48.105 | 456.08 | 1185.7 | 425.1 | 48.077 | 688.96 | -3.8407 | 1.57E-12 | 1.02E-09 | *Pramef12* | 4 |
| ENSMUSG00000057836 | 44.083 | 37.576 | 36.346 | 181.62 | 355.62 | 112.5 | 39.335 | 216.58 | -2.4636 | 3.00E-12 | 1.87E-09 | *Xlr3a* | X |
| ENSMUSG00000025519 | 15.744 | 39.553 | 35.277 | 121.42 | 259.34 | 138.38 | 30.191 | 173.05 | -2.5191 | 3.85E-12 | 2.32E-09 | *Tktl2* | 8 |
| ENSMUSG00000079536 | 11.546 | 30.654 | 27.794 | 106.11 | 238.53 | 114.49 | 23.331 | 153.04 | -2.7138 | 4.27E-12 | 2.49E-09 | *Gm6880* | X |
| ENSMUSG00000058328 | 12.595 | 32.632 | 26.725 | 121.42 | 264.55 | 99.555 | 23.984 | 161.84 | -2.7541 | 6.11E-12 | 3.45E-09 | *Xlr5a* | X |
| ENSMUSG00000054889 | 974.03 | 1060 | 892.61 | 480.57 | 405.93 | 545.56 | 975.56 | 477.35 | 1.0326 | 7.04E-12 | 3.85E-09 | *Dsp* | 13 |
| ENSMUSG00000007888 | 35.687 | 63.285 | 60.933 | 169.37 | 203.83 | 190.15 | 53.302 | 187.78 | -1.8163 | 1.18E-11 | 6.27E-09 | *Crlf1* | 8 |
| ENSMUSG00000072249 | 0 | 2.9665 | 6.414 | 42.853 | 107.55 | 52.764 | 3.1268 | 67.724 | -4.4478 | 3.51E-11 | 1.77E-08 | *Fthl17f* | X |

| | | | | | | | | | | | | | |
|---|---|---|---|---|---|---|---|---|---|---|---|---|---|
| ENSMUSG00000006310 | 15.744 | 32.632 | 21.38 | 120.4 | 217.71 | 83.626 | 23.252 | 140.58 | -2.5945 | 3.53E-11 | 1.77E-08 | *Zbtb32* | 7 |
| ENSMUSG00000010796 | 102.86 | 168.1 | 103.69 | 309.16 | 706.03 | 347.45 | 124.89 | 454.21 | -1.8628 | 4.08E-11 | 1.98E-08 | *Asz1* | 6 |
| ENSMUSG00000022487 | 46.183 | 73.174 | 47.036 | 211.21 | 357.35 | 143.36 | 55.464 | 237.31 | -2.0968 | 4.22E-11 | 1.98E-08 | *Gtsf1* | 15 |
| ENSMUSG00000063804 | 26.24 | 39.553 | 39.553 | 123.46 | 269.75 | 122.45 | 35.115 | 171.89 | -2.2935 | 4.26E-11 | 1.98E-08 | *Lin28b* | 10 |
| ENSMUSG00000010651 | 17.843 | 16.81 | 24.587 | 91.828 | 176.94 | 80.639 | 19.747 | 116.47 | -2.5662 | 5.35E-11 | 2.42E-08 | *Acaa1b* | 9 |
| ENSMUSG00000056436 | 15.744 | 45.486 | 28.863 | 113.26 | 188.22 | 142.36 | 30.031 | 147.95 | -2.2973 | 5.87E-11 | 2.59E-08 | *Cyct* | 2 |
| ENSMUSG00000073130 | 11.546 | 13.844 | 10.69 | 63.26 | 148.32 | 67.697 | 12.026 | 93.092 | -2.954 | 6.40E-11 | 2.75E-08 | *Gm1141* | X |
| ENSMUSG00000044576 | 15.744 | 42.52 | 23.518 | 106.11 | 216.84 | 123.45 | 27.261 | 148.8 | -2.4454 | 7.37E-11 | 3.10E-08 | *Garem2* | 5 |
| ENSMUSG00000096883 | 450.28 | 516.17 | 315.35 | 1091.7 | 954.97 | 834.27 | 427.27 | 960.33 | -1.1675 | 8.25E-11 | 3.39E-08 | *Shisa8* | 15 |
| ENSMUSG00000039431 | 87.117 | 87.017 | 117.59 | 253.04 | 402.46 | 232.96 | 97.241 | 296.15 | -1.6097 | 9.73E-11 | 3.91E-08 | *Mtmr7* | 8 |
| ENSMUSG00000055403 | 28.339 | 37.576 | 21.38 | 112.23 | 312.25 | 100.55 | 29.098 | 175.01 | -2.5885 | 1.14E-10 | 4.49E-08 | *4933427D06Rik* | 6 |
| ENSMUSG00000039209 | 76.621 | 142.39 | 111.18 | 290.79 | 642.72 | 278.75 | 110.06 | 404.09 | -1.8768 | 2.21E-10 | 8.50E-08 | *Rpl39l* | 16 |
| ENSMUSG00000071341 | 112.31 | 234.35 | 248.01 | 554.03 | 780.63 | 458.95 | 198.22 | 597.87 | -1.5931 | 2.56E-10 | 9.62E-08 | *Egr4* | 6 |
| ENSMUSG00000016626 | 3.1488 | 1.9777 | 4.276 | 57.138 | 127.5 | 22.898 | 3.1342 | 69.179 | -4.4769 | 2.71E-10 | 1.00E-07 | *Nlrp14* | 7 |
| ENSMUSG00000027315 | 46.183 | 88.995 | 62.002 | 192.84 | 361.69 | 184.18 | 65.726 | 246.24 | -1.9053 | 2.99E-10 | 1.08E-07 | *Spint1* | 2 |
| ENSMUSG00000022763 | 18.893 | 45.486 | 44.898 | 118.36 | 294.04 | 140.37 | 36.426 | 184.26 | -2.3399 | 3.16E-10 | 1.12E-07 | *Aifm3* | 16 |
| ENSMUSG00000083411 | 231.96 | 382.68 | 271.52 | 54.077 | 91.073 | 123.45 | 295.39 | 89.533 | 1.7217 | 3.63E-10 | 1.26E-07 | *Rpl30-ps10* | X |
| ENSMUSG00000041324 | 3994.8 | 3565.7 | 3652.8 | 2140.6 | 2315.9 | 2439.1 | 3737.8 | 2298.5 | 0.7014 | 4.63E-10 | 1.58E-07 | *Inhba* | 13 |
| ENSMUSG00000067768 | 8.3968 | 20.766 | 12.828 | 75.503 | 164.8 | 62.72 | 13.997 | 101.01 | -2.8489 | 4.85E-10 | 1.63E-07 | *Xlr4b* | X |
| ENSMUSG00000034690 | 99.712 | 159.2 | 128.28 | 561.17 | 1236.9 | 625.2 | 129.06 | 807.75 | -2.6458 | 5.50E-10 | 1.81E-07 | *Nlrp4c* | 7 |
| ENSMUSG00000115483 | 16.794 | 24.721 | 19.242 | 92.849 | 122.3 | 76.657 | 20.252 | 97.268 | -2.2632 | 6.88E-10 | 2.22E-07 | *Gm9732* | 14 |
| ENSMUSG00000005251 | 32.538 | 35.598 | 49.174 | 136.72 | 322.66 | 106.52 | 39.103 | 188.64 | -2.2743 | 7.98E-10 | 2.53E-07 | *Ripk4* | 16 |
| ENSMUSG00000020059 | 22.042 | 44.498 | 27.794 | 97.95 | 338.27 | 114.49 | 31.444 | 183.57 | -2.5452 | 8.87E-10 | 2.76E-07 | *Sycp3* | 10 |
| ENSMUSG00000053398 | 2543.2 | 2838 | 2689.6 | 4387.4 | 4662.1 | 3991.2 | 2690.2 | 4346.9 | -0.6923 | 9.13E-10 | 2.80E-07 | *Phgdh* | 3 |
| ENSMUSG00000073460 | 93.415 | 115.69 | 98.347 | 278.55 | 497.87 | 210.06 | 102.49 | 328.82 | -1.6832 | 9.95E-10 | 3.00E-07 | *Pnldc1* | 17 |
| ENSMUSG00000025038 | 4.1984 | 25.71 | 18.173 | 76.524 | 147.45 | 92.586 | 16.027 | 105.52 | -2.7152 | 1.47E-09 | 4.31E-07 | *Efhc2* | X |
| ENSMUSG00000072944 | 106.01 | 152.28 | 119.73 | 253.04 | 623.64 | 330.52 | 126.01 | 402.4 | -1.6761 | 1.48E-09 | 4.31E-07 | *Nup62cl* | X |
| ENSMUSG00000049387 | 17.843 | 24.721 | 22.449 | 70.402 | 189.95 | 91.591 | 21.671 | 117.32 | -2.439 | 1.51E-09 | 4.34E-07 | *Cox7b2* | 5 |
| ENSMUSG00000009248 | 3.1488 | 6.9218 | 11.759 | 40.813 | 114.49 | 58.737 | 7.2765 | 71.347 | -3.3009 | 1.77E-09 | 5.00E-07 | *Ascl2* | 7 |
| ENSMUSG00000062170 | 12.595 | 15.821 | 17.104 | 65.3 | 119.7 | 69.688 | 15.174 | 84.895 | -2.4876 | 1.83E-09 | 5.09E-07 | *Fmr1nb* | X |
| ENSMUSG00000034555 | 5.248 | 17.799 | 20.311 | 63.26 | 170.87 | 76.657 | 14.453 | 103.6 | -2.8437 | 1.86E-09 | 5.10E-07 | *Tex16* | X |
| ENSMUSG00000046167 | 217.27 | 196.78 | 268.32 | 58.158 | 53.777 | 107.52 | 227.45 | 73.151 | 1.6395 | 2.04E-09 | 5.52E-07 | *Gldn* | 9 |
| ENSMUSG00000045179 | 9.4464 | 25.71 | 7.483 | 176.51 | 489.19 | 189.15 | 14.213 | 284.95 | -4.3212 | 2.27E-09 | 6.02E-07 | *Sox3* | X |
| ENSMUSG00000024406 | 37.786 | 48.453 | 66.278 | 186.72 | 315.72 | 114.49 | 50.839 | 205.64 | -2.0193 | 2.35E-09 | 6.15E-07 | *Pou5f1* | 17 |
| ENSMUSG00000072437 | 39.885 | 41.531 | 47.036 | 121.42 | 261.08 | 121.46 | 42.817 | 167.98 | -1.9757 | 2.39E-09 | 6.18E-07 | *Nanos1* | 19 |
| ENSMUSG00000028089 | 416.69 | 425.2 | 363.46 | 666.27 | 838.74 | 740.69 | 401.78 | 748.57 | -0.8981 | 2.95E-09 | 7.47E-07 | *Chd1l* | 3 |
| ENSMUSG00000079387 | 3.1488 | 15.821 | 9.6209 | 56.117 | 131.84 | 55.751 | 9.5304 | 81.236 | -3.0867 | 2.98E-09 | 7.47E-07 | *Luzp4* | X |
| ENSMUSG00000037910 | 12.595 | 41.531 | 36.346 | 102.03 | 220.31 | 122.45 | 30.157 | 148.26 | -2.2975 | 3.23E-09 | 8.00E-07 | *1700018B24Rik* | 3 |

| | | | | | | | | | | | | | |
|---|---|---|---|---|---|---|---|---|---|---|---|---|---|
| ENSMUSG00000045394 | 215.17 | 309.51 | 366.66 | 668.31 | 754.61 | 556.51 | 297.11 | 659.81 | -1.1517 | 3.39E-09 | 8.28E-07 | *Epcam* | 17 |
| ENSMUSG00000026709 | 370.51 | 368.84 | 363.46 | 644.84 | 753.74 | 618.24 | 367.6 | 672.27 | -0.8717 | 4.01E-09 | 9.66E-07 | *Dars2* | 1 |
| ENSMUSG00000036832 | 5.248 | 24.721 | 21.38 | 76.524 | 117.96 | 95.573 | 17.116 | 96.686 | -2.4955 | 4.38E-09 | 1.04E-06 | *Lpar3* | 3 |
| ENSMUSG00000109561 | 29.389 | 71.196 | 56.657 | 168.35 | 294.04 | 142.36 | 52.414 | 201.58 | -1.9433 | 4.47E-09 | 1.05E-06 | *Ankrd31* | 13 |
| ENSMUSG00000034645 | 19.942 | 35.598 | 18.173 | 128.56 | 142.25 | 70.684 | 24.571 | 113.83 | -2.208 | 5.29E-09 | 1.22E-06 | *Zyg11a* | 4 |
| ENSMUSG00000038305 | 1852.6 | 1749.2 | 1892.1 | 1117.2 | 1108.5 | 1213.6 | 1831.3 | 1146.4 | 0.6758 | 5.33E-09 | 1.22E-06 | *Spats2l* | 1 |
| ENSMUSG00000060985 | 123.85 | 148.33 | 110.11 | 302.01 | 636.65 | 255.86 | 127.43 | 398.17 | -1.6447 | 5.47E-09 | 1.24E-06 | *Tdrd5* | 1 |
| ENSMUSG00000037605 | 362.11 | 305.55 | 311.08 | 140.8 | 160.46 | 162.27 | 326.25 | 154.51 | 1.0772 | 6.55E-09 | 1.46E-06 | *Adgrl3* | 5 |
| ENSMUSG00000040612 | 176.33 | 159.2 | 169.97 | 315.28 | 477.92 | 340.48 | 168.5 | 377.89 | -1.1673 | 7.20E-09 | 1.59E-06 | *Ildr2* | 1 |
| ENSMUSG00000028698 | 243.51 | 197.77 | 171.04 | 446.9 | 861.29 | 382.29 | 204.1 | 563.49 | -1.4664 | 7.35E-09 | 1.60E-06 | *Pik3r3* | 4 |
| ENSMUSG00000042498 | 35.687 | 37.576 | 36.346 | 90.808 | 299.24 | 117.47 | 36.536 | 169.17 | -2.2142 | 7.88E-09 | 1.70E-06 | *D330045A20Rik* | X |
| ENSMUSG00000090470 | 2.0992 | 5.933 | 6.414 | 34.691 | 84.134 | 43.804 | 4.8154 | 54.21 | -3.4948 | 9.03E-09 | 1.92E-06 | *Prr23a3* | 9 |
| ENSMUSG00000022949 | 25.191 | 13.844 | 18.173 | 62.239 | 235.06 | 70.684 | 19.069 | 122.66 | -2.6913 | 1.11E-08 | 2.32E-06 | *Clic6* | 16 |
| ENSMUSG00000040649 | 115.46 | 121.63 | 129.35 | 246.92 | 575.93 | 267.8 | 122.14 | 363.55 | -1.5758 | 1.17E-08 | 2.43E-06 | *Rimklb* | 6 |
| ENSMUSG00000079584 | 38.835 | 92.95 | 37.415 | 176.51 | 317.46 | 159.29 | 56.4 | 217.75 | -1.9465 | 1.46E-08 | 3.00E-06 | *Gm364* | X |
| ENSMUSG00000058897 | 638.16 | 552.76 | 535.57 | 314.26 | 232.45 | 329.53 | 575.49 | 292.08 | 0.9805 | 1.52E-08 | 3.08E-06 | *Col25a1* | 3 |
| ENSMUSG00000024805 | 298.09 | 370.81 | 338.87 | 557.09 | 1110.2 | 669.01 | 335.92 | 778.78 | -1.214 | 1.75E-08 | 3.49E-06 | *Pcgf5* | 19 |
| ENSMUSG00000039639 | 322.23 | 282.81 | 403.01 | 139.78 | 113.62 | 178.2 | 336.02 | 143.87 | 1.2256 | 1.76E-08 | 3.49E-06 | *Kcne1* | 16 |
| ENSMUSG00000049493 | 32.538 | 38.565 | 32.07 | 89.788 | 218.58 | 107.52 | 34.391 | 138.63 | -2.0135 | 1.80E-08 | 3.53E-06 | *Pls1* | 9 |
| ENSMUSG00000078206 | 0 | 7.9107 | 4.276 | 43.874 | 83.267 | 33.849 | 4.0622 | 53.663 | -3.7115 | 1.96E-08 | 3.82E-06 | *Fthl17d* | X |
| ENSMUSG00000112858 | 9.4464 | 11.866 | 10.69 | 51.016 | 102.35 | 49.777 | 10.667 | 67.714 | -2.6689 | 2.22E-08 | 4.26E-06 | *Rnf212b* | 14 |
| ENSMUSG00000055780 | 55.629 | 138.44 | 73.761 | 474.45 | 882.11 | 455.96 | 89.275 | 604.17 | -2.7578 | 2.53E-08 | 4.82E-06 | *Usp26* | X |
| ENSMUSG00000036766 | 136.45 | 191.83 | 161.42 | 320.38 | 327 | 399.22 | 163.23 | 348.86 | -1.0943 | 2.59E-08 | 4.88E-06 | *Dner* | 1 |
| ENSMUSG00000036304 | 37.786 | 72.185 | 29.932 | 151.01 | 307.91 | 117.47 | 46.634 | 192.13 | -2.0411 | 3.05E-08 | 5.58E-06 | *Zdhhc23* | 16 |
| ENSMUSG00000031383 | 157.44 | 203.7 | 149.66 | 322.42 | 545.57 | 343.46 | 170.27 | 403.82 | -1.2466 | 3.05E-08 | 5.58E-06 | *Dusp9* | X |
| ENSMUSG00000016239 | 376.81 | 471.67 | 280.08 | 700.96 | 1359.2 | 718.79 | 376.19 | 926.3 | -1.3003 | 3.06E-08 | 5.58E-06 | *Lonrf3* | X |
| ENSMUSG00000095872 | 0 | 4.9442 | 6.414 | 26.528 | 104.08 | 38.826 | 3.786 | 56.479 | -3.9019 | 3.14E-08 | 5.67E-06 | *Gm15128* | X |
| ENSMUSG00000037625 | 714.78 | 1069.9 | 751.5 | 2179.4 | 1395.6 | 1553.1 | 845.4 | 1709.3 | -1.015 | 3.44E-08 | 6.15E-06 | *Cldn11* | 3 |
| ENSMUSG00000109860 | 60.877 | 68.23 | 62.002 | 144.88 | 366.9 | 151.32 | 63.703 | 221.03 | -1.7973 | 3.47E-08 | 6.15E-06 | *Gm17737* | 7 |
| ENSMUSG00000070348 | 176.33 | 248.2 | 234.11 | 425.47 | 477.05 | 410.17 | 219.55 | 437.56 | -0.995 | 3.70E-08 | 6.49E-06 | *Ccnd1* | 7 |
| ENSMUSG00000039224 | 48.282 | 78.118 | 39.553 | 124.48 | 285.36 | 169.24 | 55.318 | 193.03 | -1.8023 | 4.27E-08 | 7.43E-06 | *D1Pas1* | 1 |
| ENSMUSG00000041020 | 65.075 | 93.939 | 72.692 | 181.62 | 360.82 | 166.26 | 77.235 | 236.23 | -1.6141 | 5.33E-08 | 9.18E-06 | *Map7d2* | X |
| ENSMUSG00000025977 | 18.893 | 32.632 | 18.173 | 79.585 | 197.76 | 69.688 | 23.232 | 115.68 | -2.3155 | 5.39E-08 | 9.19E-06 | *Boll* | 1 |
| ENSMUSG00000073125 | 66.125 | 51.419 | 85.519 | 241.81 | 194.29 | 140.37 | 67.688 | 192.16 | -1.5074 | 5.56E-08 | 9.39E-06 | *Xlr3b* | X |
| ENSMUSG00000037979 | 31.488 | 30.654 | 42.76 | 99.991 | 196.02 | 93.582 | 34.967 | 129.87 | -1.898 | 6.05E-08 | 1.01E-05 | *Ccdc92* | 5 |
| ENSMUSG00000008489 | 129.1 | 162.17 | 104.76 | 239.77 | 439.75 | 308.62 | 132.01 | 329.38 | -1.3195 | 6.43E-08 | 1.07E-05 | *Elavl2* | 4 |
| ENSMUSG00000040434 | 122.8 | 179.97 | 194.56 | 305.07 | 470.11 | 360.39 | 165.78 | 378.52 | -1.1924 | 6.53E-08 | 1.07E-05 | *Large2* | 2 |
| ENSMUSG00000035427 | 78.72 | 87.017 | 67.347 | 369.35 | 1034.8 | 348.44 | 77.695 | 584.19 | -2.9107 | 7.22E-08 | 1.18E-05 | *Mageb4* | X |

| | | | | | | | | | | | | | |
|---|---|---|---|---|---|---|---|---|---|---|---|---|---|
| ENSMUSG00000078249 | 15.744 | 36.587 | 24.587 | 81.625 | 139.65 | 90.595 | 25.639 | 103.96 | -2.0179 | 7.55E-08 | 1.22E-05 | *Hmga1b* | 11 |
| ENSMUSG00000024640 | 1387.6 | 1322.1 | 1483.8 | 2285.5 | 2220.5 | 1997.1 | 1397.8 | 2167.7 | -0.6332 | 8.32E-08 | 1.33E-05 | *Psat1* | 19 |
| ENSMUSG00000045662 | 6.2976 | 13.844 | 8.5519 | 36.731 | 131.84 | 48.782 | 9.5644 | 72.451 | -2.9203 | 8.54E-08 | 1.34E-05 | *Henmt1* | 3 |
| ENSMUSG00000027722 | 651.8 | 929.5 | 775.02 | 1203 | 2441.6 | 1479.4 | 785.44 | 1708 | -1.1211 | 8.56E-08 | 1.34E-05 | *Spata5* | 3 |
| ENSMUSG00000097599 | 0 | 2.9665 | 1.069 | 29.589 | 60.716 | 27.875 | 1.3452 | 39.393 | -4.8475 | 8.90E-08 | 1.39E-05 | *Gm26678* | 3 |
| ENSMUSG00000009108 | 28.339 | 50.431 | 43.829 | 93.869 | 169.14 | 141.37 | 40.866 | 134.79 | -1.7221 | 9.63E-08 | 1.49E-05 | *Gnat2* | 3 |
| ENSMUSG00000034640 | 450.28 | 449.92 | 399.8 | 636.68 | 919.41 | 811.37 | 433.33 | 789.15 | -0.8655 | 9.86E-08 | 1.51E-05 | *Tiparp* | 3 |
| ENSMUSG00000067360 | 11.546 | 44.498 | 31.001 | 93.869 | 212.5 | 98.559 | 29.015 | 134.98 | -2.2168 | 1.02E-07 | 1.54E-05 | *Pramel3* | X |
| ENSMUSG00000006362 | 992.93 | 1027.4 | 1036.9 | 600.97 | 462.31 | 675.98 | 1019.1 | 579.75 | 0.8151 | 1.11E-07 | 1.67E-05 | *Cbfa2t3* | 8 |
| ENSMUSG00000070802 | 653.9 | 705.04 | 814.57 | 432.61 | 365.16 | 442.02 | 724.51 | 413.27 | 0.8108 | 1.18E-07 | 1.77E-05 | *Pnmal2* | 7 |
| ENSMUSG00000073177 | 19.942 | 30.654 | 31.001 | 92.849 | 182.15 | 68.693 | 27.199 | 114.56 | -2.0773 | 1.23E-07 | 1.83E-05 | *Gm773* | X |
| ENSMUSG00000031842 | 23.091 | 19.777 | 22.449 | 64.28 | 126.64 | 75.662 | 21.772 | 88.859 | -2.0339 | 1.37E-07 | 2.01E-05 | *Pde4c* | 8 |
| ENSMUSG00000039187 | 226.71 | 222.49 | 195.63 | 394.86 | 470.11 | 364.37 | 214.94 | 409.78 | -0.9316 | 1.40E-07 | 2.04E-05 | *Fanci* | 7 |
| ENSMUSG00000038028 | 167.94 | 150.3 | 156.07 | 260.18 | 373.83 | 344.46 | 158.1 | 326.16 | -1.0463 | 1.46E-07 | 2.12E-05 | *Tigar* | 6 |
| ENSMUSG00000074269 | 19.942 | 27.687 | 32.07 | 73.463 | 165.67 | 83.626 | 26.567 | 107.59 | -2.0218 | 1.50E-07 | 2.16E-05 | *Rec114* | 9 |
| ENSMUSG00000028278 | 286.54 | 406.41 | 316.42 | 612.19 | 995.73 | 564.48 | 336.46 | 724.13 | -1.1063 | 1.55E-07 | 2.21E-05 | *Rragd* | 4 |
| ENSMUSG00000028626 | 1116.8 | 1246.9 | 1593.9 | 819.31 | 725.12 | 751.64 | 1319.2 | 765.36 | 0.7856 | 1.76E-07 | 2.48E-05 | *Col9a2* | 4 |
| ENSMUSG00000036111 | 44.083 | 17.799 | 21.38 | 105.09 | 185.62 | 71.68 | 27.754 | 120.8 | -2.1268 | 1.86E-07 | 2.60E-05 | *Lmo1* | 7 |
| ENSMUSG00000009670 | 66.125 | 126.57 | 79.106 | 405.07 | 983.59 | 400.21 | 90.6 | 596.29 | -2.7181 | 1.95E-07 | 2.71E-05 | *Tex11* | X |
| ENSMUSG00000041359 | 43.034 | 75.151 | 29.932 | 217.33 | 208.17 | 99.555 | 49.372 | 175.02 | -1.8224 | 1.98E-07 | 2.73E-05 | *Tcl1* | 12 |
| ENSMUSG00000015365 | 101.81 | 185.9 | 107.97 | 644.84 | 1450.2 | 537.6 | 131.89 | 877.56 | -2.7339 | 2.09E-07 | 2.86E-05 | *Mov10l1* | 15 |
| ENSMUSG00000028587 | 172.14 | 160.19 | 204.18 | 356.09 | 490.93 | 297.67 | 178.83 | 381.56 | -1.0956 | 2.41E-07 | 3.27E-05 | *Orc1* | 4 |
| ENSMUSG00000051176 | 7.3472 | 13.844 | 8.5519 | 54.077 | 71.124 | 43.804 | 9.9143 | 56.335 | -2.5027 | 2.46E-07 | 3.31E-05 | *Zfp42* | 8 |
| ENSMUSG00000056155 | 33.587 | 27.687 | 25.656 | 80.605 | 168.27 | 79.644 | 28.977 | 109.51 | -1.9219 | 2.49E-07 | 3.33E-05 | *Nanos3* | 8 |
| ENSMUSG00000037224 | 857.53 | 698.12 | 602.91 | 429.55 | 404.19 | 372.34 | 719.52 | 402.03 | 0.8398 | 2.53E-07 | 3.36E-05 | *Zfyve28* | 5 |
| ENSMUSG00000054321 | 220.42 | 260.06 | 188.14 | 392.82 | 680.01 | 401.21 | 222.87 | 491.35 | -1.1413 | 2.59E-07 | 3.42E-05 | *Taf4b* | 18 |
| ENSMUSG00000072479 | 24.141 | 24.721 | 25.656 | 55.097 | 124.03 | 109.51 | 24.839 | 96.214 | -1.9565 | 2.63E-07 | 3.45E-05 | *Xlr5b* | X |
| ENSMUSG00000079033 | 20.992 | 29.665 | 38.484 | 64.28 | 182.15 | 110.51 | 29.714 | 118.98 | -2.0056 | 2.87E-07 | 3.72E-05 | *Mef2b* | 8 |
| ENSMUSG00000071815 | 5.248 | 15.821 | 14.966 | 49.995 | 107.55 | 50.773 | 12.012 | 69.441 | -2.5324 | 2.88E-07 | 3.72E-05 | *Fthl17e* | X |
| ENSMUSG00000025090 | 9.4464 | 38.565 | 35.277 | 78.564 | 184.75 | 107.52 | 27.763 | 123.61 | -2.1548 | 3.47E-07 | 4.44E-05 | *Ccdc172* | 19 |
| ENSMUSG00000062542 | 98.663 | 178.98 | 87.657 | 320.38 | 353.88 | 238.93 | 121.77 | 304.4 | -1.3199 | 3.74E-07 | 4.76E-05 | *Syt9* | 7 |
| ENSMUSG00000026398 | 27.29 | 24.721 | 49.174 | 153.05 | 83.267 | 114.49 | 33.728 | 116.93 | -1.7949 | 3.80E-07 | 4.80E-05 | *Nr5a2* | 1 |
| ENSMUSG00000091490 | 2.0992 | 13.844 | 11.759 | 70.402 | 60.716 | 43.804 | 9.2339 | 58.307 | -2.6533 | 4.00E-07 | 4.99E-05 | *Triml2* | 8 |
| ENSMUSG00000031162 | 90.266 | 84.051 | 80.174 | 186.72 | 209.9 | 170.24 | 84.83 | 188.95 | -1.1564 | 4.03E-07 | 4.99E-05 | *Gata1* | X |
| ENSMUSG00000096520 | 11.546 | 17.799 | 12.828 | 47.955 | 125.77 | 50.773 | 14.058 | 74.832 | -2.4138 | 4.06E-07 | 4.99E-05 | *Gm3376* | Y |
| ENSMUSG00000079432 | 7.3472 | 20.766 | 7.483 | 60.199 | 91.073 | 47.786 | 11.865 | 66.353 | -2.4758 | 4.06E-07 | 4.99E-05 | *Gm15023* | X |
| ENSMUSG00000046999 | 456.58 | 345.1 | 383.77 | 218.35 | 130.97 | 213.05 | 395.15 | 187.46 | 1.0788 | 4.12E-07 | 5.03E-05 | *1110032F04Rik* | 3 |
| ENSMUSG00000046000 | 3.1488 | 4.9442 | 2.138 | 26.528 | 58.981 | 30.862 | 3.4103 | 38.79 | -3.5021 | 4.15E-07 | 5.04E-05 | *Naa11* | 5 |

| | | | | | | | | | | | | | |
|---|---|---|---|---|---|---|---|---|---|---|---|---|---|
| ENSMUSG00000040957 | 1073.7 | 941.37 | 1157.7 | 547.91 | 456.23 | 734.72 | 1057.6 | 579.62 | 0.8684 | 4.34E-07 | 5.23E-05 | *Cables1* | 18 |
| ENSMUSG00000031558 | 1288.9 | 1220.2 | 1311.7 | 777.48 | 693.89 | 904.95 | 1273.6 | 792.11 | 0.6857 | 4.49E-07 | 5.38E-05 | *Slit2* | 5 |
| ENSMUSG00000070529 | 784.05 | 1014.5 | 894.75 | 1558 | 1313.2 | 1494.3 | 897.78 | 1455.2 | -0.6961 | 4.68E-07 | 5.56E-05 | *Wfdc10* | 2 |
| ENSMUSG00000046337 | 26.24 | 21.754 | 32.07 | 72.442 | 154.39 | 76.657 | 26.688 | 101.16 | -1.9286 | 4.89E-07 | 5.78E-05 | *Fam178b* | 1 |
| ENSMUSG00000029223 | 813.44 | 763.38 | 765.4 | 1888.6 | 3846.8 | 2079.7 | 780.74 | 2605 | -1.7385 | 4.94E-07 | 5.79E-05 | *Uchl1* | 5 |
| ENSMUSG00000036181 | 910.01 | 719.87 | 912.92 | 554.03 | 452.76 | 489.81 | 847.6 | 498.87 | 0.7652 | 5.04E-07 | 5.88E-05 | *Hist1h1c* | 13 |
| ENSMUSG00000043549 | 10.496 | 15.821 | 18.173 | 38.772 | 189.95 | 51.769 | 14.83 | 93.498 | -2.6608 | 5.17E-07 | 5.99E-05 | *Fam90a1b* | X |
| ENSMUSG00000064194 | 9.4464 | 17.799 | 31.001 | 62.239 | 168.27 | 64.711 | 19.415 | 98.406 | -2.3474 | 5.29E-07 | 6.09E-05 | *Zfp936* | 7 |
| ENSMUSG00000026110 | 144.85 | 126.57 | 137.9 | 293.85 | 298.37 | 230.97 | 136.44 | 274.4 | -1.0092 | 5.37E-07 | 6.15E-05 | *Mgat4a* | 1 |
| ENSMUSG00000046942 | 5.248 | 10.877 | 11.759 | 53.056 | 84.134 | 33.849 | 9.2947 | 57.013 | -2.6192 | 5.53E-07 | 6.29E-05 | *Mageb16* | X |
| ENSMUSG00000028345 | 1222.8 | 1371.5 | 1175.9 | 1845.8 | 2562.2 | 1868.6 | 1256.7 | 2092.2 | -0.7356 | 6.13E-07 | 6.93E-05 | *Tex10* | 4 |
| ENSMUSG00000027577 | 22.042 | 50.431 | 67.347 | 154.07 | 142.25 | 137.39 | 46.606 | 144.57 | -1.6334 | 6.18E-07 | 6.93E-05 | *Chrna4* | 2 |
| ENSMUSG00000037572 | 856.48 | 870.17 | 835.95 | 1345.8 | 1986.3 | 1229.5 | 854.2 | 1520.5 | -0.8325 | 6.44E-07 | 7.18E-05 | *Wdhd1* | 14 |
| ENSMUSG00000034220 | 3063.8 | 2383.1 | 2752.7 | 1714.1 | 1473.7 | 1912.5 | 2733.2 | 1700.1 | 0.6852 | 6.63E-07 | 7.35E-05 | *Gpc1* | 1 |
| ENSMUSG00000020396 | 17.843 | 43.509 | 27.794 | 83.666 | 176.08 | 84.622 | 29.715 | 114.79 | -1.9491 | 7.17E-07 | 7.90E-05 | *Nefh* | 11 |
| ENSMUSG00000021257 | 147.99 | 150.3 | 138.97 | 271.4 | 302.71 | 262.83 | 145.76 | 278.98 | -0.9371 | 7.51E-07 | 8.18E-05 | *Angel1* | 12 |
| ENSMUSG00000038080 | 344.27 | 301.59 | 287.56 | 538.73 | 817.92 | 480.85 | 311.14 | 612.5 | -0.9785 | 7.51E-07 | 8.18E-05 | *Kdm1b* | 13 |
| ENSMUSG00000002324 | 23.091 | 12.855 | 23.518 | 78.564 | 132.71 | 48.782 | 19.821 | 86.684 | -2.1361 | 8.13E-07 | 8.79E-05 | *Rec8* | 14 |
| ENSMUSG00000072244 | 173.18 | 311.48 | 204.18 | 428.53 | 559.45 | 441.03 | 229.62 | 476.34 | -1.0521 | 8.17E-07 | 8.79E-05 | *Trim6* | 7 |
| ENSMUSG00000049164 | 1006.6 | 1160.9 | 910.78 | 1543.7 | 2595.2 | 1592.9 | 1026.1 | 1910.6 | -0.8972 | 8.47E-07 | 9.06E-05 | *Zfp518a* | 19 |
| ENSMUSG00000071470 | 19.942 | 31.643 | 28.863 | 74.483 | 152.66 | 72.675 | 26.816 | 99.938 | -1.9004 | 8.82E-07 | 9.37E-05 | *Ccnb1ip1* | 14 |
| ENSMUSG00000053046 | 11.546 | 37.576 | 21.38 | 65.3 | 144.85 | 87.608 | 23.5 | 99.253 | -2.076 | 9.75E-07 | 0.000103 | *Brsk2* | 7 |
| ENSMUSG00000078932 | 17.843 | 34.609 | 25.656 | 80.605 | 201.23 | 59.733 | 26.036 | 113.86 | -2.1298 | 9.77E-07 | 0.000103 | *CN725425* | 15 |
| ENSMUSG00000017724 | 141.7 | 183.92 | 197.76 | 326.5 | 322.66 | 344.46 | 174.46 | 331.21 | -0.9246 | 9.88E-07 | 0.000103 | *Etv4* | 11 |
| ENSMUSG00000026024 | 467.07 | 463.76 | 383.77 | 739.73 | 951.5 | 645.12 | 438.2 | 778.78 | -0.8302 | 1.03E-06 | 0.000107 | *Als2* | 1 |
| ENSMUSG00000108801 | 300.19 | 202.71 | 225.56 | 121.42 | 85.869 | 122.45 | 242.82 | 109.91 | 1.1463 | 1.05E-06 | 0.000109 | *Gm39090* | 7 |
| ENSMUSG00000073000 | 17.843 | 10.877 | 25.656 | 85.707 | 98.88 | 47.786 | 18.125 | 77.457 | -2.1028 | 1.07E-06 | 0.00011 | *Gm10451* | 12 |
| ENSMUSG00000032261 | 566.79 | 505.29 | 422.25 | 778.5 | 1018.3 | 785.49 | 498.11 | 860.76 | -0.7897 | 1.28E-06 | 0.00013 | *Sh3bgrl2* | 9 |
| ENSMUSG00000018830 | 2375.3 | 2286.2 | 2456.5 | 4392.5 | 3341.1 | 3332.1 | 2372.7 | 3688.6 | -0.6364 | 1.32E-06 | 0.000134 | *Myh11* | 16 |
| ENSMUSG00000043099 | 1859.9 | 1924.3 | 2271.6 | 1374.4 | 1230.8 | 1364.9 | 2018.6 | 1323.4 | 0.6094 | 1.34E-06 | 0.000135 | *Hic1* | 11 |
| ENSMUSG00000067702 | 252.95 | 706.03 | 475.7 | 2000.8 | 4418.4 | 2350.5 | 478.23 | 2923.2 | -2.6117 | 1.43E-06 | 0.000144 | *Tuba3a* | 6 |
| ENSMUSG00000073243 | 1.0496 | 3.9553 | 2.138 | 20.406 | 50.307 | 28.871 | 2.381 | 33.195 | -3.7926 | 1.44E-06 | 0.000144 | *Gm9* | X |
| ENSMUSG00000096966 | 23.091 | 58.341 | 56.657 | 114.28 | 265.41 | 109.51 | 46.03 | 163.07 | -1.8264 | 1.55E-06 | 0.000154 | *Gm18336* | X |
| ENSMUSG00000011589 | 77.671 | 76.14 | 66.278 | 154.07 | 343.48 | 135.39 | 73.363 | 210.98 | -1.5266 | 1.56E-06 | 0.000154 | *Fsd1* | 17 |
| ENSMUSG00000080069 | 14.694 | 22.743 | 34.208 | 166.31 | 454.5 | 113.49 | 23.882 | 244.77 | -3.3587 | 1.68E-06 | 0.000165 | *Gm41* | X |
| ENSMUSG00000079105 | 319.08 | 364.88 | 195.63 | 127.54 | 122.3 | 150.33 | 293.2 | 133.39 | 1.1379 | 1.69E-06 | 0.000165 | *C7* | 15 |
| ENSMUSG00000032939 | 1045.4 | 879.07 | 952.47 | 1606 | 1810.2 | 1278.3 | 958.98 | 1564.8 | -0.707 | 1.75E-06 | 0.000169 | *Nup93* | 8 |
| ENSMUSG00000031458 | 198.38 | 193.81 | 222.35 | 306.09 | 556.85 | 388.26 | 204.85 | 417.07 | -1.0279 | 1.75E-06 | 0.000169 | *Coprs* | 8 |

| | | | | | | | | | | | | | |
|---|---|---|---|---|---|---|---|---|---|---|---|---|---|
| ENSMUSG00000036752 | 4249.8 | 4214.4 | 5727.7 | 6668.8 | 9568.8 | 7460.6 | 4730.6 | 7899.4 | -0.7399 | 1.76E-06 | 0.000169 | *Tubb4b* | 2 |
| ENSMUSG00000078498 | 23.091 | 60.319 | 50.243 | 116.32 | 249.8 | 101.55 | 44.551 | 155.89 | -1.8078 | 1.78E-06 | 0.000171 | *Zfp988* | 4 |
| ENSMUSG00000066687 | 53.53 | 122.62 | 55.588 | 309.16 | 608.89 | 316.58 | 77.244 | 411.54 | -2.4126 | 2.14E-06 | 0.000203 | *Zbtb16* | 9 |
| ENSMUSG00000072923 | 7.3472 | 13.844 | 5.345 | 145.91 | 283.63 | 39.822 | 8.8453 | 156.45 | -4.1416 | 2.15E-06 | 0.000203 | *Gm10439* | X |
| ENSMUSG00000070732 | 32.538 | 85.04 | 48.105 | 259.16 | 600.22 | 244.91 | 55.227 | 368.09 | -2.7359 | 2.31E-06 | 0.000217 | *Rbm44* | 1 |
| ENSMUSG00000046709 | 81.869 | 96.906 | 72.692 | 146.93 | 259.34 | 185.17 | 83.822 | 197.15 | -1.2351 | 2.46E-06 | 0.000231 | *Mapk10* | 5 |
| ENSMUSG00000061911 | 9.4464 | 25.71 | 19.242 | 67.341 | 122.3 | 48.782 | 18.133 | 79.474 | -2.1316 | 2.74E-06 | 0.000255 | *Myt1l* | 12 |
| ENSMUSG00000002076 | 14.694 | 35.598 | 18.173 | 67.341 | 103.22 | 78.648 | 22.822 | 83.069 | -1.8599 | 2.95E-06 | 0.000274 | *Hsf2bp* | 17 |
| ENSMUSG00000024647 | 78.72 | 71.196 | 66.278 | 125.5 | 202.96 | 175.22 | 72.065 | 167.89 | -1.2222 | 2.97E-06 | 0.000274 | *Cbln2* | 18 |
| ENSMUSG00000039109 | 200.47 | 155.25 | 190.28 | 333.64 | 334.8 | 316.58 | 182 | 328.34 | -0.8524 | 3.05E-06 | 0.000277 | *F13a1* | 13 |
| ENSMUSG00000038299 | 1139.9 | 1243 | 1204.8 | 1620.3 | 2123.3 | 1767.1 | 1195.9 | 1836.9 | -0.6196 | 3.05E-06 | 0.000277 | *Wdr36* | 18 |
| ENSMUSG00000038508 | 596.18 | 431.13 | 576.19 | 270.38 | 248.07 | 355.41 | 534.5 | 291.29 | 0.8764 | 3.06E-06 | 0.000277 | *Gdf15* | 8 |
| ENSMUSG00000026893 | 546.84 | 656.59 | 409.42 | 995.83 | 848.28 | 947.76 | 537.62 | 930.62 | -0.7905 | 3.07E-06 | 0.000277 | *Gca* | 2 |
| ENSMUSG00000042195 | 342.17 | 377.73 | 242.66 | 547.91 | 671.34 | 529.63 | 320.86 | 582.96 | -0.8613 | 3.11E-06 | 0.000279 | *Slc35f2* | 9 |
| ENSMUSG00000045330 | 441.88 | 880.06 | 683.09 | 1130.5 | 1344.4 | 1270.3 | 668.34 | 1248.4 | -0.9011 | 3.27E-06 | 0.000293 | *4933402E13Rik* | X |
| ENSMUSG00000033948 | 55.629 | 51.419 | 42.76 | 96.93 | 287.97 | 107.52 | 49.936 | 164.14 | -1.7198 | 3.32E-06 | 0.000296 | *Zswim5* | 4 |
| ENSMUSG00000035395 | 38.835 | 33.62 | 39.553 | 88.767 | 165.67 | 83.626 | 37.336 | 112.69 | -1.5984 | 3.36E-06 | 0.000298 | *Pet2* | X |
| ENSMUSG00000060176 | 30.439 | 35.598 | 28.863 | 80.605 | 153.52 | 72.675 | 31.633 | 102.27 | -1.6954 | 3.51E-06 | 0.000309 | *Kif27* | 13 |
| ENSMUSG00000025757 | 306.48 | 389.6 | 302.53 | 529.54 | 780.63 | 520.67 | 332.87 | 610.28 | -0.8751 | 3.82E-06 | 0.000335 | *Hspa4l* | 3 |
| ENSMUSG00000025001 | 1780.1 | 2253.6 | 2187.2 | 2814 | 5113.1 | 3238.5 | 2073.6 | 3721.9 | -0.8441 | 3.85E-06 | 0.000337 | *Hells* | 19 |
| ENSMUSG00000060461 | 15.744 | 14.833 | 11.759 | 39.792 | 79.798 | 59.733 | 14.112 | 59.774 | -2.0849 | 3.96E-06 | 0.000343 | *Dppa5a* | 9 |
| ENSMUSG00000071562 | 34.637 | 35.598 | 21.38 | 5.1016 | 2.6021 | 0 | 30.538 | 2.5679 | 3.5795 | 3.97E-06 | 0.000343 | *Stfa1* | 16 |
| ENSMUSG00000094004 | 5.248 | 7.9107 | 6.414 | 18.366 | 82.4 | 41.813 | 6.5242 | 47.526 | -2.8673 | 4.13E-06 | 0.000355 | *Gm5128* | X |
| ENSMUSG00000049539 | 0 | 1.9777 | 4.276 | 35.711 | 27.756 | 23.893 | 2.0845 | 29.12 | -3.8108 | 4.22E-06 | 0.000361 | *Hist1h1a* | 13 |
| ENSMUSG00000062327 | 64.026 | 50.431 | 40.622 | 10.203 | 15.613 | 6.9688 | 51.693 | 10.928 | 2.2297 | 4.24E-06 | 0.000361 | *T* | 17 |
| ENSMUSG00000029149 | 66.125 | 67.241 | 84.45 | 147.95 | 236.79 | 137.39 | 72.605 | 174.04 | -1.2649 | 4.40E-06 | 0.000373 | *Krtcap3* | 5 |
| ENSMUSG00000030158 | 602.47 | 847.43 | 687.36 | 1379.5 | 1092 | 1074.2 | 712.42 | 1181.9 | -0.7295 | 4.48E-06 | 0.000378 | *Clec12b* | 6 |
| ENSMUSG00000025876 | 23.091 | 36.587 | 38.484 | 71.422 | 111.02 | 110.51 | 32.721 | 97.65 | -1.5784 | 4.60E-06 | 0.000387 | *Unc5a* | 13 |
| ENSMUSG00000030747 | 129.1 | 132.5 | 118.66 | 239.77 | 536.9 | 196.12 | 126.75 | 324.27 | -1.3571 | 4.65E-06 | 0.000389 | *Dgat2* | 7 |
| ENSMUSG00000074637 | 29.389 | 47.464 | 38.484 | 2.0406 | 8.6736 | 5.9733 | 38.446 | 5.5625 | 2.7676 | 4.88E-06 | 0.000407 | *Sox2* | 3 |
| ENSMUSG00000016763 | 533.2 | 483.54 | 536.63 | 278.55 | 300.98 | 356.41 | 517.79 | 311.98 | 0.7308 | 5.07E-06 | 0.000418 | *Scube1* | 15 |
| ENSMUSG00000037315 | 1196.5 | 1382.4 | 1121.4 | 2064.1 | 2044.4 | 1625.7 | 1233.4 | 1911.4 | -0.6318 | 5.08E-06 | 0.000418 | *Jade3* | X |
| ENSMUSG00000025586 | 41.984 | 65.263 | 31.001 | 139.78 | 183.88 | 86.613 | 46.083 | 136.76 | -1.5678 | 5.09E-06 | 0.000418 | *Cpeb1* | 7 |
| ENSMUSG00000047674 | 6.2976 | 16.81 | 27.794 | 61.219 | 119.7 | 51.769 | 16.967 | 77.561 | -2.1977 | 5.12E-06 | 0.000419 | *Pdha2* | 3 |
| ENSMUSG00000032854 | 12.595 | 15.821 | 17.104 | 55.097 | 88.471 | 42.809 | 15.174 | 62.126 | -2.0374 | 5.33E-06 | 0.000434 | *Ugt8a* | 3 |
| ENSMUSG00000026274 | 535.3 | 529.03 | 463.94 | 868.29 | 815.32 | 717.79 | 509.42 | 800.47 | -0.6517 | 5.40E-06 | 0.000437 | *Pask* | 1 |
| ENSMUSG00000040111 | 2695.4 | 2524.5 | 2216 | 1816.2 | 1390.4 | 1659.6 | 2478.6 | 1622 | 0.6123 | 5.44E-06 | 0.000438 | *Gramd1b* | 9 |
| ENSMUSG00000071568 | 2.0992 | 0.9888 | 2.138 | 44.894 | 6.0716 | 48.782 | 1.742 | 33.249 | -4.2592 | 5.45E-06 | 0.000438 | *Gm5874* | 6 |

| | | | | | | | | | | | | | |
|---|---|---|---|---|---|---|---|---|---|---|---|---|---|
| ENSMUSG00000067338 | 224.62 | 739.65 | 516.32 | 2016.1 | 3831.1 | 2170.3 | 493.53 | 2672.5 | -2.4369 | 5.51E-06 | 0.000439 | *Tuba3b* | 6 |
| ENSMUSG00000026915 | 820.79 | 890.94 | 894.75 | 1220.3 | 1605.5 | 1242.4 | 868.83 | 1356.1 | -0.6429 | 5.52E-06 | 0.000439 | *Strbp* | 2 |
| ENSMUSG00000081137 | 53.53 | 104.82 | 48.105 | 169.37 | 267.15 | 134.4 | 68.817 | 190.31 | -1.4662 | 5.54E-06 | 0.000439 | *BC022960* | X |
| ENSMUSG00000047281 | 501.71 | 354 | 390.18 | 210.19 | 269.75 | 181.19 | 415.3 | 220.38 | 0.9117 | 5.64E-06 | 0.000445 | *Sfn* | 4 |
| ENSMUSG00000026669 | 607.72 | 511.23 | 593.29 | 923.39 | 1058.2 | 780.51 | 570.75 | 920.69 | -0.6908 | 5.85E-06 | 0.00046 | *Mcm10* | 2 |
| ENSMUSG00000024462 | 874.32 | 973.01 | 930.02 | 613.21 | 542.97 | 658.06 | 925.79 | 604.75 | 0.6153 | 6.39E-06 | 0.000499 | *Gabbr1* | 17 |
| ENSMUSG00000025857 | 314.88 | 300.61 | 259.77 | 476.49 | 523.02 | 448.99 | 291.75 | 482.83 | -0.7271 | 6.40E-06 | 0.000499 | *Dnaaf5* | 5 |
| ENSMUSG00000040629 | 184.73 | 373.78 | 225.56 | 774.42 | 1840.5 | 951.75 | 261.36 | 1188.9 | -2.1854 | 6.54E-06 | 0.000507 | *Mael* | 1 |
| ENSMUSG00000043453 | 6.2976 | 16.81 | 21.38 | 47.955 | 106.69 | 49.777 | 14.829 | 68.139 | -2.204 | 6.57E-06 | 0.000508 | *Magea10* | X |
| ENSMUSG00000051257 | 27.29 | 89.984 | 63.071 | 123.46 | 270.62 | 161.28 | 60.115 | 185.12 | -1.6223 | 6.66E-06 | 0.000512 | *Trap1a* | X |
| ENSMUSG00000085577 | 1.0496 | 5.933 | 2.138 | 29.589 | 49.44 | 18.915 | 3.0402 | 32.648 | -3.411 | 6.73E-06 | 0.000515 | *Mageb6-ps* | X |
| ENSMUSG00000029778 | 2159 | 2121.1 | 2023.6 | 1565.2 | 1085.9 | 1403.7 | 2101.2 | 1351.6 | 0.6373 | 6.77E-06 | 0.000517 | *Adcyap1r1* | 6 |
| ENSMUSG00000085146 | 1.0496 | 3.9553 | 2.138 | 16.325 | 55.511 | 21.902 | 2.381 | 31.246 | -3.7076 | 7.09E-06 | 0.000539 | *Eif2c5* | X |
| ENSMUSG00000049382 | 918.4 | 717.89 | 589.02 | 395.88 | 483.99 | 405.19 | 741.77 | 428.35 | 0.7913 | 7.16E-06 | 0.000542 | *Krt8* | 15 |
| ENSMUSG00000031362 | 15.744 | 20.766 | 9.6209 | 38.772 | 77.195 | 74.666 | 15.377 | 63.545 | -2.0442 | 7.22E-06 | 0.000544 | *Xlr4c* | X |
| ENSMUSG00000074595 | 78.72 | 94.928 | 101.55 | 211.21 | 250.67 | 144.35 | 91.734 | 202.08 | -1.141 | 7.27E-06 | 0.000545 | *Wfdc6a* | 2 |
| ENSMUSG00000044349 | 75.572 | 87.017 | 65.209 | 157.13 | 228.98 | 136.39 | 75.933 | 174.17 | -1.1989 | 7.63E-06 | 0.00057 | *Snhg11* | 2 |
| ENSMUSG00000040013 | 181.58 | 394.55 | 285.42 | 771.36 | 1498.8 | 948.76 | 287.18 | 1073 | -1.9015 | 7.68E-06 | 0.00057 | *Fkbp6* | 5 |
| ENSMUSG00000030110 | 204.67 | 342.14 | 168.9 | 424.45 | 665.27 | 432.07 | 238.57 | 507.26 | -1.0879 | 7.70E-06 | 0.00057 | *Ret* | 6 |
| ENSMUSG00000043342 | 1886.1 | 1768 | 2409.5 | 1359.1 | 1269 | 1332 | 2021.2 | 1320 | 0.6146 | 7.73E-06 | 0.000571 | *Hoxd9* | 2 |
| ENSMUSG00000029135 | 1041.2 | 924.56 | 840.23 | 589.74 | 539.5 | 664.03 | 935.33 | 597.76 | 0.6466 | 7.82E-06 | 0.000572 | *Fosl2* | 5 |
| ENSMUSG00000074129 | 958.29 | 975.98 | 901.16 | 634.64 | 667 | 602.31 | 945.14 | 634.65 | 0.5743 | 7.82E-06 | 0.000572 | *Rpl13a* | 7 |
| ENSMUSG00000027547 | 93.415 | 177 | 114.38 | 217.33 | 516.08 | 245.9 | 128.27 | 326.44 | -1.3482 | 7.87E-06 | 0.000574 | *Sall4* | 2 |
| ENSMUSG00000036743 | 56.679 | 57.352 | 40.622 | 103.05 | 179.54 | 115.48 | 51.551 | 132.69 | -1.3657 | 8.02E-06 | 0.000583 | *Psma8* | 18 |
| ENSMUSG00000013415 | 100.76 | 251.16 | 173.18 | 332.62 | 492.66 | 354.42 | 175.03 | 393.23 | -1.1672 | 8.31E-06 | 0.000601 | *Igf2bp1* | 11 |
| ENSMUSG00000031966 | 348.47 | 427.18 | 451.12 | 809.11 | 586.34 | 665.03 | 408.92 | 686.83 | -0.7473 | 8.54E-06 | 0.000615 | *Glb1l3* | 9 |
| ENSMUSG00000045004 | 1041.2 | 931.48 | 1071.1 | 405.07 | 439.75 | 781.51 | 1014.6 | 542.11 | 0.9047 | 8.69E-06 | 0.000623 | *Spata21* | 4 |
| ENSMUSG00000037541 | 409.35 | 552.76 | 378.42 | 268.34 | 202.96 | 266.81 | 446.84 | 246.04 | 0.8633 | 8.73E-06 | 0.000624 | *Shank2* | 7 |
| ENSMUSG00000048647 | 40.935 | 32.632 | 17.104 | 73.463 | 197.76 | 68.693 | 30.223 | 113.3 | -1.9087 | 9.03E-06 | 0.00064 | *Exd1* | 2 |
| ENSMUSG00000040688 | 795.6 | 679.33 | 771.81 | 1124.4 | 1452 | 1023.4 | 748.91 | 1199.9 | -0.681 | 9.03E-06 | 0.00064 | *Tbl3* | 17 |
| ENSMUSG00000001985 | 8.3968 | 15.821 | 6.414 | 44.894 | 63.318 | 38.826 | 10.211 | 49.013 | -2.2573 | 9.54E-06 | 0.000672 | *Grik3* | 4 |
| ENSMUSG00000090744 | 12.595 | 25.71 | 9.6209 | 46.935 | 122.3 | 48.782 | 15.975 | 72.672 | -2.1831 | 9.55E-06 | 0.000672 | *Gm6871* | 7 |
| ENSMUSG00000067547 | 112.31 | 101.85 | 146.45 | 389.76 | 143.98 | 368.35 | 120.2 | 300.7 | -1.3217 | 9.77E-06 | 0.000685 | *Gm7666* | 15 |
| ENSMUSG00000031283 | 129.1 | 135.47 | 109.04 | 766.26 | 679.15 | 243.91 | 124.54 | 563.1 | -2.1768 | 1.01E-05 | 0.000702 | *Chrdl1* | X |
| ENSMUSG00000061728 | 7.3472 | 4.9442 | 3.207 | 16.325 | 90.206 | 25.884 | 5.1661 | 44.138 | -3.1 | 1.04E-05 | 0.000721 | *Btnl7-ps* | 17 |
| ENSMUSG00000010342 | 71.373 | 165.14 | 94.071 | 459.14 | 949.76 | 361.38 | 110.19 | 590.1 | -2.4206 | 1.05E-05 | 0.000729 | *Tex14* | 11 |
| ENSMUSG00000041438 | 979.28 | 1149 | 986.68 | 1553.9 | 2281.2 | 1396.8 | 1038.3 | 1744 | -0.7485 | 1.06E-05 | 0.000734 | *Utp4* | 8 |
| ENSMUSG00000013584 | 333.77 | 405.42 | 489.6 | 787.68 | 611.49 | 673.99 | 409.6 | 691.05 | -0.7542 | 1.10E-05 | 0.000758 | *Aldh1a2* | 9 |

| | | | | | | | | | | | | | |
|---|---|---|---|---|---|---|---|---|---|---|---|---|---|
| ENSMUSG00000052688 | 537.4 | 482.55 | 335.66 | 785.64 | 708.64 | 802.41 | 451.87 | 765.56 | -0.7599 | 1.14E-05 | 0.000781 | *Rab7b* | 1 |
| ENSMUSG00000020534 | 758.86 | 711.96 | 750.43 | 1079.5 | 1255.9 | 1005.5 | 740.42 | 1113.6 | -0.5896 | 1.17E-05 | 0.000796 | *Shmt1* | 11 |
| ENSMUSG00000100009 | 5.248 | 0.9888 | 1.069 | 17.345 | 65.052 | 15.929 | 2.4353 | 32.776 | -3.7636 | 1.21E-05 | 0.000825 | *Gm7967* | 1 |
| ENSMUSG00000000365 | 58.778 | 165.14 | 101.55 | 442.82 | 1194.4 | 479.85 | 108.49 | 705.68 | -2.7012 | 1.23E-05 | 0.000836 | *Rnf17* | 14 |
| ENSMUSG00000032101 | 136.45 | 139.43 | 143.25 | 212.23 | 576.8 | 238.93 | 139.71 | 342.65 | -1.2966 | 1.25E-05 | 0.000843 | *Ddx25* | 9 |
| ENSMUSG00000020083 | 72.423 | 62.297 | 68.416 | 120.4 | 195.16 | 142.36 | 67.712 | 152.64 | -1.176 | 1.30E-05 | 0.000876 | *Fam241b* | 10 |
| ENSMUSG00000024087 | 828.14 | 619.01 | 767.54 | 459.14 | 410.26 | 504.74 | 738.23 | 458.05 | 0.6889 | 1.35E-05 | 0.000905 | *Cyp1b1* | 17 |
| ENSMUSG00000058806 | 673.85 | 561.66 | 659.57 | 456.08 | 303.58 | 366.36 | 631.69 | 375.34 | 0.7525 | 1.36E-05 | 0.000909 | *Col13a1* | 10 |
| ENSMUSG00000025732 | 1231.2 | 1204.4 | 1311.7 | 852.98 | 636.65 | 906.95 | 1249.1 | 798.86 | 0.6458 | 1.41E-05 | 0.000939 | *Mcrip2* | 17 |
| ENSMUSG00000000120 | 669.65 | 548.8 | 808.16 | 326.5 | 308.78 | 486.82 | 675.54 | 374.04 | 0.8533 | 1.43E-05 | 0.000945 | *Ngfr* | 11 |
| ENSMUSG00000020021 | 154.29 | 205.68 | 156.07 | 330.58 | 372.1 | 258.84 | 172.01 | 320.51 | -0.8977 | 1.43E-05 | 0.000945 | *Fgd6* | 10 |
| ENSMUSG00000000037 | 356.87 | 610.11 | 467.15 | 1213.2 | 2392.2 | 1195.7 | 478.04 | 1600.3 | -1.7432 | 1.44E-05 | 0.000948 | *Scml2* | X |
| ENSMUSG00000052374 | 8.3968 | 15.821 | 17.104 | 57.138 | 79.798 | 35.84 | 13.774 | 57.592 | -2.0662 | 1.50E-05 | 0.000983 | *Actn2* | 13 |
| ENSMUSG00000001642 | 5801.2 | 4119.5 | 3926.4 | 6765.7 | 7934.7 | 6877.3 | 4615.7 | 7192.5 | -0.64 | 1.52E-05 | 0.000991 | *Akr1b3* | 6 |
| ENSMUSG00000028532 | 658.1 | 389.6 | 452.18 | 277.53 | 180.41 | 306.63 | 499.96 | 254.86 | 0.9738 | 1.55E-05 | 0.001009 | *Cachd1* | 4 |
| ENSMUSG00000033732 | 2984 | 2672.8 | 3077.6 | 4256.8 | 4633.5 | 3702.4 | 2911.5 | 4197.6 | -0.528 | 1.62E-05 | 0.001047 | *Sf3b3* | 8 |
| ENSMUSG00000029769 | 239.31 | 263.03 | 187.07 | 348.95 | 623.64 | 376.32 | 229.8 | 449.63 | -0.9693 | 1.67E-05 | 0.001075 | *Ccdc136* | 6 |
| ENSMUSG00000061186 | 140.65 | 213.59 | 138.97 | 356.09 | 342.61 | 254.86 | 164.4 | 317.85 | -0.9498 | 1.74E-05 | 0.001122 | *Sfmbt2* | 2 |
| ENSMUSG00000006998 | 4172.2 | 4105.6 | 4256.7 | 5894.4 | 7184.4 | 5251.5 | 4178.2 | 6110.1 | -0.5485 | 1.77E-05 | 0.00113 | *Psmd2* | 16 |
| ENSMUSG00000096834 | 1.0496 | 1.9777 | 3.207 | 21.427 | 39.899 | 16.924 | 2.0781 | 26.083 | -3.6584 | 1.77E-05 | 0.00113 | *Gm15127* | X |
| ENSMUSG00000003051 | 132.25 | 144.37 | 188.14 | 66.321 | 70.257 | 80.639 | 154.92 | 72.406 | 1.097 | 1.77E-05 | 0.00113 | *Elf3* | 1 |
| ENSMUSG00000050424 | 57.728 | 38.565 | 52.381 | 225.49 | 480.52 | 127.43 | 49.558 | 277.81 | -2.4882 | 1.78E-05 | 0.001131 | *Pnma5* | X |
| ENSMUSG00000009941 | 172.14 | 311.48 | 249.08 | 849.92 | 2497.1 | 822.32 | 244.23 | 1389.8 | -2.5086 | 1.79E-05 | 0.001131 | *Nxf2* | X |
| ENSMUSG00000048636 | 575.18 | 515.18 | 605.05 | 383.64 | 268.88 | 367.36 | 565.14 | 339.96 | 0.7349 | 1.80E-05 | 0.001131 | *A730049H05Rik* | 6 |
| ENSMUSG00000024837 | 1692 | 2537.4 | 2058.9 | 2999.7 | 3629.9 | 3155.9 | 2096.1 | 3261.8 | -0.638 | 1.80E-05 | 0.001131 | *Dmrt1* | 19 |
| ENSMUSG00000086841 | 993.98 | 1251.9 | 927.89 | 708.1 | 648.79 | 700.87 | 1057.9 | 685.92 | 0.6259 | 1.85E-05 | 0.001158 | *2410006H16Rik* | 11 |
| ENSMUSG00000087522 | 5.248 | 10.877 | 4.276 | 23.467 | 69.389 | 33.849 | 6.8004 | 42.235 | -2.6307 | 1.87E-05 | 0.001164 | *Gm371* | X |
| ENSMUSG00000031778 | 830.24 | 972.02 | 819.92 | 575.46 | 398.99 | 612.26 | 874.06 | 528.9 | 0.7264 | 1.90E-05 | 0.001183 | *Cx3cl1* | 8 |
| ENSMUSG00000037266 | 9169.3 | 8760.1 | 8203.5 | 6420.8 | 6599.8 | 6269 | 8711 | 6429.9 | 0.438 | 1.94E-05 | 0.001197 | *Rsrp1* | 4 |
| ENSMUSG00000038932 | 86.068 | 138.44 | 79.106 | 283.65 | 720.78 | 325.54 | 101.2 | 443.32 | -2.131 | 1.94E-05 | 0.001197 | *Tcfl5* | 2 |
| ENSMUSG00000041801 | 942.54 | 988.84 | 1155.6 | 604.03 | 700.83 | 727.75 | 1029 | 677.53 | 0.6024 | 1.98E-05 | 0.001216 | *Phlda3* | 1 |
| ENSMUSG00000067835 | 2.0992 | 0.9888 | 2.138 | 15.305 | 41.634 | 18.915 | 1.742 | 25.285 | -3.875 | 1.99E-05 | 0.001216 | *Gm14661* | X |
| ENSMUSG00000001334 | 957.24 | 978.95 | 786.78 | 1418.2 | 1242.1 | 1392.8 | 907.65 | 1351 | -0.5732 | 1.99E-05 | 0.001216 | *Fndc5* | 4 |
| ENSMUSG00000090626 | 153.24 | 176.01 | 144.31 | 240.79 | 319.19 | 290.7 | 157.86 | 283.56 | -0.8454 | 2.10E-05 | 0.00128 | *Tex9* | 9 |
| ENSMUSG00000010592 | 545.79 | 1322.1 | 913.99 | 2962 | 7580.8 | 3739.3 | 927.29 | 4760.7 | -2.3601 | 2.14E-05 | 0.001299 | *Dazl* | 17 |
| ENSMUSG00000036478 | 2092.9 | 1833.3 | 2141.2 | 1333.6 | 1304.5 | 1549.1 | 2022.5 | 1395.7 | 0.5352 | 2.15E-05 | 0.001299 | *Btg1* | 10 |
| ENSMUSG00000028527 | 141.7 | 218.53 | 179.59 | 280.59 | 562.92 | 298.66 | 179.94 | 380.72 | -1.0824 | 2.34E-05 | 0.001408 | *Ak4* | 4 |
| ENSMUSG00000074887 | 2.0992 | 5.933 | 8.5519 | 21.427 | 62.45 | 29.866 | 5.5281 | 37.914 | -2.7843 | 2.34E-05 | 0.001408 | *EU599041* | 7 |

| | | | | | | | | | | | | | |
|---|---|---|---|---|---|---|---|---|---|---|---|---|---|
| ENSMUSG00000045010 | 72.423 | 132.5 | 70.554 | 295.89 | 684.35 | 282.74 | 91.827 | 420.99 | -2.1965 | 2.39E-05 | 0.001428 | *Gm4779* | X |
| ENSMUSG00000039476 | 1037 | 953.24 | 948.2 | 661.16 | 570.73 | 726.75 | 979.48 | 652.88 | 0.586 | 2.45E-05 | 0.001456 | *Prrx2* | 2 |
| ENSMUSG00000000531 | 864.87 | 794.03 | 817.78 | 581.58 | 425.88 | 574.43 | 825.56 | 527.3 | 0.6481 | 2.45E-05 | 0.001456 | *Grasp* | 15 |
| ENSMUSG00000047139 | 1134.6 | 927.53 | 1106.4 | 592.8 | 721.65 | 754.63 | 1056.2 | 689.69 | 0.6142 | 2.50E-05 | 0.001484 | *Cd24a* | 10 |
| ENSMUSG00000039911 | 1447.4 | 1276.6 | 1368.3 | 773.4 | 751.14 | 1084.2 | 1364.1 | 869.56 | 0.65 | 2.54E-05 | 0.001501 | *Spsb1* | 4 |
| ENSMUSG00000028024 | 2318.6 | 2168.5 | 2091 | 1615.2 | 1563.9 | 1119 | 2192.7 | 1432.7 | 0.6138 | 2.55E-05 | 0.001501 | *Enpep* | 3 |
| ENSMUSG00000045822 | 80.82 | 82.073 | 67.347 | 135.7 | 229.85 | 142.36 | 76.746 | 169.31 | -1.1437 | 2.59E-05 | 0.001517 | *Zswim3* | 2 |
| ENSMUSG00000030031 | 107.06 | 103.83 | 114.38 | 161.21 | 365.16 | 201.1 | 108.42 | 242.49 | -1.1643 | 2.59E-05 | 0.001517 | *Kbtbd8* | 6 |
| ENSMUSG00000033405 | 104.96 | 87.017 | 68.416 | 162.23 | 203.83 | 166.26 | 86.798 | 177.44 | -1.0325 | 2.72E-05 | 0.001585 | *Nudt15* | 14 |
| ENSMUSG00000036564 | 255.05 | 261.05 | 244.8 | 364.25 | 747.67 | 391.25 | 253.64 | 501.06 | -0.9838 | 2.73E-05 | 0.001585 | *Ndrg4* | 8 |
| ENSMUSG00000047220 | 26.24 | 49.442 | 53.45 | 72.442 | 196.02 | 114.49 | 43.044 | 127.65 | -1.5711 | 2.80E-05 | 0.001622 | *Ccdc36* | 9 |
| ENSMUSG00000022432 | 38.835 | 154.26 | 63.071 | 365.27 | 1078.1 | 564.48 | 85.388 | 669.29 | -2.9699 | 3.05E-05 | 0.001755 | *Smc1b* | 15 |
| ENSMUSG00000035653 | 200.47 | 98.884 | 137.9 | 51.016 | 51.175 | 76.657 | 145.75 | 59.616 | 1.2904 | 3.05E-05 | 0.001755 | *Lrfn5* | 12 |
| ENSMUSG00000028525 | 221.47 | 154.26 | 176.38 | 405.07 | 279.29 | 322.56 | 184.04 | 335.64 | -0.8664 | 3.07E-05 | 0.001761 | *Pde4b* | 4 |
| ENSMUSG00000004661 | 172.14 | 187.88 | 119.73 | 313.24 | 390.31 | 233.95 | 159.91 | 312.5 | -0.9667 | 3.18E-05 | 0.001822 | *Arid3b* | 9 |
| ENSMUSG00000069044 | 46.183 | 88.006 | 33.139 | 119.38 | 287.97 | 103.54 | 55.776 | 170.29 | -1.6097 | 3.58E-05 | 0.002043 | *Usp9y* | Y |
| ENSMUSG00000025608 | 419.84 | 402.46 | 564.43 | 802.99 | 608.89 | 933.83 | 462.24 | 781.9 | -0.7581 | 3.80E-05 | 0.002155 | *Podxl* | 6 |
| ENSMUSG00000079479 | 1.0496 | 3.9553 | 1.069 | 18.366 | 32.96 | 20.907 | 2.0246 | 24.077 | -3.5549 | 3.80E-05 | 0.002155 | *Gm9112* | X |
| ENSMUSG00000022724 | 331.67 | 310.49 | 258.7 | 494.85 | 628.84 | 411.16 | 300.29 | 511.62 | -0.7696 | 3.84E-05 | 0.002169 | *Riox2* | 16 |
| ENSMUSG00000095478 | 427.19 | 358.95 | 339.94 | 458.12 | 761.55 | 734.72 | 375.36 | 651.46 | -0.7963 | 3.86E-05 | 0.002169 | *Gm9824* | 10 |
| ENSMUSG00000034685 | 648.66 | 606.16 | 720.5 | 462.2 | 367.76 | 446.01 | 658.44 | 425.32 | 0.6315 | 3.86E-05 | 0.002169 | *Fam171a2* | 11 |
| ENSMUSG00000026219 | 4620.4 | 5164.7 | 4380.7 | 6786.1 | 7341.4 | 5899.6 | 4721.9 | 6675.7 | -0.4996 | 3.88E-05 | 0.002172 | *Trip12* | 1 |
| ENSMUSG00000041567 | 192.08 | 109.76 | 165.69 | 93.869 | 47.705 | 51.769 | 155.84 | 64.448 | 1.2776 | 3.93E-05 | 0.00219 | *Serpina12* | 12 |
| ENSMUSG00000051316 | 699.04 | 972.02 | 942.85 | 1167.2 | 1903.9 | 1316.1 | 871.3 | 1462.4 | -0.7476 | 4.04E-05 | 0.002245 | *Taf7* | 18 |
| ENSMUSG00000044338 | 313.83 | 288.74 | 360.25 | 169.37 | 143.12 | 224 | 320.94 | 178.83 | 0.8454 | 4.10E-05 | 0.002275 | *Aplnr* | 2 |
| ENSMUSG00000095599 | 1.0496 | 4.9442 | 4.276 | 18.366 | 66.787 | 13.938 | 3.4233 | 33.03 | -3.2714 | 4.15E-05 | 0.002294 | *Gm6215* | X |
| ENSMUSG00000032936 | 20.992 | 36.587 | 17.104 | 62.239 | 125.77 | 58.737 | 24.894 | 82.248 | -1.7233 | 4.24E-05 | 0.002334 | *Camkv* | 9 |
| ENSMUSG00000059625 | 58.778 | 79.107 | 54.519 | 217.33 | 788.43 | 220.02 | 64.134 | 408.59 | -2.6716 | 4.25E-05 | 0.002335 | *Sohlh1* | 2 |
| ENSMUSG00000054626 | 24.141 | 23.732 | 14.966 | 52.036 | 85.002 | 60.729 | 20.946 | 65.922 | -1.6548 | 4.38E-05 | 0.002399 | *Xlr* | X |
| ENSMUSG00000094378 | 4.1984 | 12.855 | 1.069 | 15.305 | 72.859 | 44.8 | 6.0408 | 44.321 | -2.8634 | 4.40E-05 | 0.002402 | *Gm15097* | X |
| ENSMUSG00000092060 | 240.36 | 321.37 | 195.63 | 474.45 | 627.1 | 340.48 | 252.45 | 480.68 | -0.9291 | 4.74E-05 | 0.002582 | *Bend4* | 5 |
| ENSMUSG00000056300 | 31.488 | 88.006 | 54.519 | 108.15 | 275.82 | 121.46 | 58.004 | 168.48 | -1.5385 | 4.87E-05 | 0.002645 | *Zfp981* | 4 |
| ENSMUSG00000054932 | 48.282 | 118.66 | 58.795 | 160.19 | 220.31 | 158.29 | 75.246 | 179.6 | -1.253 | 4.93E-05 | 0.00267 | *Afp* | 5 |
| ENSMUSG00000026121 | 889.01 | 947.3 | 942.85 | 682.59 | 532.56 | 656.07 | 926.39 | 623.74 | 0.5718 | 5.01E-05 | 0.002702 | *Sema4c* | 1 |
| ENSMUSG00000032291 | 37.786 | 66.252 | 29.932 | 152.03 | 596.75 | 198.11 | 44.657 | 315.63 | -2.8208 | 5.20E-05 | 0.002799 | *Crabp1* | 9 |
| ENSMUSG00000069310 | 13.645 | 7.9107 | 14.966 | 42.853 | 59.848 | 38.826 | 12.174 | 47.176 | -1.9621 | 5.32E-05 | 0.002854 | *Hist1h3c* | 13 |
| ENSMUSG00000040570 | 155.34 | 203.7 | 131.49 | 224.47 | 501.34 | 301.65 | 163.51 | 342.49 | -1.0676 | 5.34E-05 | 0.002855 | *Rundc3b* | 5 |
| ENSMUSG00000043557 | 574.13 | 714.93 | 582.6 | 458.12 | 244.6 | 362.38 | 623.89 | 355.03 | 0.8157 | 5.47E-05 | 0.002918 | *Mdga1* | 17 |

| | | | | | | | | | | | | | |
|---|---|---|---|---|---|---|---|---|---|---|---|---|---|
| ENSMUSG00000027719 | 35.687 | 52.408 | 22.449 | 70.402 | 163.93 | 90.595 | 36.848 | 108.31 | -1.5552 | 5.54E-05 | 0.002944 | *Adad1* | 3 |
| ENSMUSG00000025089 | 117.56 | 232.38 | 146.45 | 287.73 | 405.06 | 289.7 | 165.46 | 327.5 | -0.9844 | 5.56E-05 | 0.00295 | *Gfra1* | 19 |
| ENSMUSG00000052942 | 79.77 | 164.15 | 95.14 | 212.23 | 327.86 | 199.11 | 113.02 | 246.4 | -1.1239 | 5.58E-05 | 0.00295 | *Glis3* | 19 |
| ENSMUSG00000043661 | 6.2976 | 10.877 | 10.69 | 32.65 | 85.002 | 23.893 | 9.2882 | 47.182 | -2.349 | 5.62E-05 | 0.002959 | *Gm9785* | X |
| ENSMUSG00000039329 | 238.26 | 300.61 | 242.66 | 703 | 1602 | 584.39 | 260.51 | 963.14 | -1.8866 | 5.63E-05 | 0.002959 | *Tex19.1* | 11 |
| ENSMUSG00000038095 | 2406.7 | 2920 | 2166.8 | 3564 | 4401 | 3279.3 | 2497.9 | 3748.1 | -0.5855 | 5.65E-05 | 0.002962 | *Sbno1* | 5 |
| ENSMUSG00000052684 | 1709.8 | 1903.5 | 2118.7 | 1336.6 | 1181.4 | 1448.5 | 1910.7 | 1322.2 | 0.5315 | 5.73E-05 | 0.002995 | *Jun* | 4 |
| ENSMUSG00000071350 | 189.98 | 152.28 | 138.97 | 257.12 | 385.11 | 250.88 | 160.41 | 297.7 | -0.8941 | 5.97E-05 | 0.003109 | *Setdb2* | 14 |
| ENSMUSG00000045672 | 902.66 | 667.46 | 857.33 | 551.99 | 469.24 | 551.53 | 809.15 | 524.26 | 0.6265 | 6.04E-05 | 0.003135 | *Col27a1* | 4 |
| ENSMUSG00000030309 | 716.88 | 822.71 | 677.74 | 1020.3 | 1377.4 | 1011.5 | 739.11 | 1136.4 | -0.621 | 6.09E-05 | 0.003149 | *Caprin2* | 6 |
| ENSMUSG00000043289 | 3.1488 | 4.9442 | 7.483 | 17.345 | 58.113 | 26.88 | 5.192 | 34.113 | -2.7243 | 6.10E-05 | 0.003149 | *Mei4* | 9 |
| ENSMUSG00000028358 | 1198.6 | 1194.5 | 946.06 | 822.37 | 671.34 | 739.69 | 1113.1 | 744.47 | 0.5811 | 6.17E-05 | 0.00318 | *Zfp618* | 4 |
| ENSMUSG00000060499 | 107.06 | 210.62 | 136.83 | 412.21 | 872.57 | 397.22 | 151.5 | 560.67 | -1.8877 | 6.36E-05 | 0.003268 | *Rpl10l* | 12 |
| ENSMUSG00000051220 | 574.13 | 553.75 | 605.05 | 750.95 | 1112 | 841.24 | 577.64 | 901.38 | -0.6431 | 6.43E-05 | 0.003291 | *Ercc6l* | X |
| ENSMUSG00000038116 | 1050.7 | 1233.1 | 1024.1 | 1517.2 | 2086 | 1464.5 | 1102.6 | 1689.2 | -0.6158 | 6.45E-05 | 0.003291 | *Phf20* | 2 |
| ENSMUSG00000024968 | 262.4 | 251.16 | 266.18 | 379.56 | 640.12 | 372.34 | 259.91 | 464 | -0.838 | 6.70E-05 | 0.003411 | *Rcor2* | 19 |
| ENSMUSG00000091497 | 1.0496 | 8.8995 | 9.6209 | 51.016 | 224.65 | 53.76 | 6.5234 | 109.81 | -4.073 | 6.73E-05 | 0.003414 | *Gm44* | X |
| ENSMUSG00000010064 | 106.01 | 167.11 | 146.45 | 71.422 | 58.113 | 68.693 | 139.86 | 66.076 | 1.0848 | 6.74E-05 | 0.003414 | *Slc38a3* | 9 |
| ENSMUSG00000095698 | 2.0992 | 0 | 2.138 | 26.528 | 24.286 | 12.942 | 1.4124 | 21.252 | -3.9359 | 6.86E-05 | 0.003464 | *Rhox2d* | X |
| ENSMUSG00000025995 | 1171.4 | 1130.2 | 1179.1 | 1452.9 | 2477.2 | 1646.6 | 1160.2 | 1858.9 | -0.6807 | 6.97E-05 | 0.003511 | *Wdr75* | 1 |
| ENSMUSG00000032902 | 768.31 | 1067.9 | 1091.4 | 1412.1 | 1695.7 | 1378.8 | 975.9 | 1495.6 | -0.6161 | 7.03E-05 | 0.003528 | *Slc16a1* | 3 |
| ENSMUSG00000091705 | 4.1984 | 5.933 | 2.138 | 15.305 | 71.124 | 15.929 | 4.0898 | 34.119 | -3.0593 | 7.10E-05 | 0.003554 | *H2-Q2* | 17 |
| ENSMUSG00000035378 | 212.02 | 247.21 | 277.94 | 363.23 | 551.64 | 367.36 | 245.72 | 427.41 | -0.8004 | 7.12E-05 | 0.003554 | *Shq1* | 6 |
| ENSMUSG00000029168 | 651.8 | 747.56 | 615.74 | 489.75 | 435.42 | 410.17 | 671.7 | 445.11 | 0.5944 | 7.18E-05 | 0.003574 | *Dpysl5* | 5 |
| ENSMUSG00000114816 | 4.1984 | 3.9553 | 1.069 | 16.325 | 42.501 | 20.907 | 3.0743 | 26.577 | -3.1094 | 7.23E-05 | 0.00359 | *Gm10323* | 13 |
| ENSMUSG00000041805 | 28.339 | 91.962 | 32.07 | 192.84 | 344.34 | 227.98 | 50.79 | 255.05 | -2.3265 | 7.28E-05 | 0.003603 | *Pramel1* | 4 |
| ENSMUSG00000106734 | 435.59 | 658.56 | 466.08 | 300.99 | 324.39 | 322.56 | 520.08 | 315.98 | 0.7193 | 7.30E-05 | 0.003603 | *Gm20559* | 6 |
| ENSMUSG00000030946 | 285.49 | 271.93 | 330.32 | 532.6 | 451.03 | 423.11 | 295.91 | 468.91 | -0.6643 | 7.33E-05 | 0.003609 | *Lhpp* | 7 |
| ENSMUSG00000035232 | 787.2 | 851.39 | 899.02 | 1116.2 | 1308 | 1188.7 | 845.87 | 1204.3 | -0.5102 | 7.67E-05 | 0.003754 | *Pdk3* | X |
| ENSMUSG00000018548 | 1349.8 | 1320.1 | 1310.6 | 1747.8 | 2248.2 | 1735.2 | 1326.8 | 1910.4 | -0.5264 | 7.67E-05 | 0.003754 | *Trim37* | 11 |
| ENSMUSG00000097248 | 494.36 | 324.34 | 446.84 | 170.39 | 265.41 | 267.8 | 421.85 | 234.54 | 0.8449 | 7.69E-05 | 0.003754 | *Gm2694* | 8 |
| ENSMUSG00000024451 | 405.15 | 337.19 | 365.6 | 242.84 | 228.12 | 225.99 | 369.31 | 232.31 | 0.6688 | 7.70E-05 | 0.003754 | *Arap3* | 18 |
| ENSMUSG00000028633 | 1354 | 1476.3 | 1512.6 | 1950.8 | 2644.6 | 1869.6 | 1447.6 | 2155 | -0.5745 | 7.81E-05 | 0.003786 | *Ctps* | 4 |
| ENSMUSG00000079465 | 722.13 | 930.49 | 594.36 | 1373.3 | 962.77 | 1331 | 748.99 | 1222.4 | -0.7056 | 7.82E-05 | 0.003786 | *Col4a3* | 1 |
| ENSMUSG00000020419 | 9.4464 | 23.732 | 7.483 | 36.731 | 85.002 | 46.791 | 13.554 | 56.175 | -2.0465 | 7.83E-05 | 0.003786 | *Hormad2* | 11 |
| ENSMUSG00000031665 | 24.141 | 39.553 | 37.415 | 83.666 | 162.2 | 55.751 | 33.703 | 100.54 | -1.5796 | 7.87E-05 | 0.003795 | *Sall1* | 8 |
| ENSMUSG00000060967 | 390.45 | 654.61 | 660.64 | 925.43 | 934.15 | 930.84 | 568.57 | 930.14 | -0.7099 | 7.91E-05 | 0.003803 | *Etd* | X |
| ENSMUSG00000019312 | 33.587 | 37.576 | 40.622 | 54.077 | 228.12 | 79.644 | 37.262 | 120.61 | -1.6988 | 8.20E-05 | 0.003932 | *Grb7* | 11 |

| | | | | | | | | | | | | | |
|---|---|---|---|---|---|---|---|---|---|---|---|---|---|
| ENSMUSG00000039200 | 68.224 | 75.151 | 76.968 | 126.52 | 222.05 | 126.43 | 73.448 | 158.33 | -1.1113 | 8.26E-05 | 0.003947 | *Atf7ip2* | 16 |
| ENSMUSG00000024266 | 46.183 | 74.163 | 26.725 | 80.605 | 244.6 | 113.49 | 49.023 | 146.23 | -1.5765 | 8.28E-05 | 0.003947 | *Adad2* | 8 |
| ENSMUSG00000031594 | 6.2976 | 23.732 | 25.656 | 55.097 | 68.522 | 67.697 | 18.562 | 63.772 | -1.7798 | 8.36E-05 | 0.003975 | *Fgl1* | 8 |
| ENSMUSG00000031239 | 10276 | 8822.4 | 10741 | 7144.3 | 7602.5 | 7241.6 | 9946.4 | 7329.4 | 0.4404 | 8.58E-05 | 0.004072 | *Itm2a* | X |
| ENSMUSG00000069041 | 193.13 | 412.34 | 242.66 | 840.74 | 1928.2 | 796.44 | 282.71 | 1188.4 | -2.0716 | 8.62E-05 | 0.004072 | *Slc25a31* | 3 |
| ENSMUSG00000029112 | 8.3968 | 16.81 | 12.828 | 44.894 | 117.96 | 19.911 | 12.678 | 60.922 | -2.2667 | 8.63E-05 | 0.004072 | *Nkx1-1* | 5 |
| ENSMUSG00000026832 | 159.54 | 178.98 | 126.14 | 276.51 | 234.19 | 310.61 | 154.89 | 273.77 | -0.8194 | 8.73E-05 | 0.004111 | *Cytip* | 2 |
| ENSMUSG00000038384 | 529 | 732.73 | 579.39 | 421.39 | 330.47 | 406.18 | 613.71 | 386.01 | 0.6706 | 8.76E-05 | 0.004112 | *Setd1b* | 5 |
| ENSMUSG00000041936 | 1836.8 | 2020.2 | 1685.8 | 1315.2 | 1129.3 | 1424.6 | 1847.6 | 1289.7 | 0.5192 | 8.80E-05 | 0.004123 | *Agrn* | 4 |
| ENSMUSG00000024437 | 9.4464 | 6.9218 | 9.6209 | 365.27 | 157.86 | 14.933 | 8.6631 | 179.36 | -4.3725 | 8.98E-05 | 0.004196 | *Gm8615* | 5 |
| ENSMUSG00000030757 | 47.232 | 64.274 | 56.657 | 122.44 | 129.24 | 105.53 | 56.054 | 119.07 | -1.0865 | 9.02E-05 | 0.004202 | *Zkscan2* | 7 |
| ENSMUSG00000053909 | 17.843 | 17.799 | 17.104 | 32.65 | 124.9 | 45.795 | 17.582 | 67.782 | -1.9521 | 9.04E-05 | 0.004202 | *Rhox10* | X |
| ENSMUSG00000042386 | 27.29 | 99.872 | 74.83 | 286.71 | 612.36 | 251.87 | 67.331 | 383.65 | -2.5102 | 9.32E-05 | 0.00432 | *Tex13b* | X |
| ENSMUSG00000034171 | 58.778 | 84.051 | 59.864 | 123.46 | 185.62 | 124.44 | 67.564 | 144.51 | -1.0974 | 9.37E-05 | 0.004333 | *Faah* | 4 |
| ENSMUSG00000005268 | 1408.6 | 1425.9 | 901.16 | 859.11 | 736.39 | 762.59 | 1245.2 | 786.03 | 0.6643 | 9.57E-05 | 0.004414 | *Prlr* | 15 |
| ENSMUSG00000009596 | 171.09 | 408.39 | 277.94 | 924.41 | 2336.7 | 866.13 | 285.8 | 1375.7 | -2.2671 | 9.72E-05 | 0.00447 | *Taf7l* | X |
| ENSMUSG00000027444 | 360.01 | 354.99 | 482.12 | 588.72 | 610.62 | 652.08 | 399.04 | 617.14 | -0.6297 | 0.000101 | 0.004655 | *8030411F24Rik* | 2 |
| ENSMUSG00000024597 | 801.9 | 837.54 | 866.95 | 1142.8 | 1229.9 | 1116 | 835.46 | 1162.9 | -0.4774 | 0.000102 | 0.004669 | *Slc12a2* | 18 |
| ENSMUSG00000061353 | 6064.6 | 5365.4 | 6338.1 | 4171.1 | 3218.8 | 4728.9 | 5922.7 | 4039.6 | 0.5522 | 0.000103 | 0.004669 | *Cxcl12* | 6 |
| ENSMUSG00000039899 | 915.25 | 1501.1 | 1164.1 | 1882.5 | 2464.2 | 1554.1 | 1193.5 | 1966.9 | -0.7208 | 0.000103 | 0.004669 | *Fgl2* | 5 |
| ENSMUSG00000079009 | 12.595 | 19.777 | 13.897 | 33.67 | 80.665 | 50.773 | 15.423 | 55.036 | -1.8365 | 0.000103 | 0.004669 | *Gm14139* | 2 |
| ENSMUSG00000040010 | 6121.3 | 6319.6 | 6479.2 | 4816.9 | 4971.7 | 4614.4 | 6306.7 | 4801 | 0.3935 | 0.000104 | 0.004703 | *Slc7a5* | 8 |
| ENSMUSG00000074731 | 7.3472 | 17.799 | 11.759 | 30.609 | 101.48 | 32.853 | 12.302 | 54.981 | -2.1613 | 0.000106 | 0.004787 | *Zfp345* | 2 |
| ENSMUSG00000028884 | 1429.6 | 1374.5 | 1683.7 | 1962.1 | 2488.5 | 2024 | 1495.9 | 2158.2 | -0.5293 | 0.000106 | 0.004787 | *Rpa2* | 4 |
| ENSMUSG00000024163 | 2101.3 | 2224.9 | 1811.9 | 1569.2 | 1404.3 | 1416.7 | 2046 | 1463.4 | 0.4839 | 0.000107 | 0.004807 | *Mapk8ip3* | 17 |
| ENSMUSG00000070570 | 4.1984 | 3.9553 | 10.69 | 25.508 | 47.705 | 27.875 | 6.2812 | 33.696 | -2.4353 | 0.000107 | 0.004812 | *Slc17a7* | 7 |
| ENSMUSG00000022602 | 22.042 | 26.699 | 13.897 | 45.914 | 96.277 | 56.746 | 20.879 | 66.313 | -1.6678 | 0.000108 | 0.004844 | *Arc* | 15 |
| ENSMUSG00000036306 | 59.827 | 83.062 | 64.14 | 128.56 | 145.72 | 140.37 | 69.01 | 138.22 | -1.0009 | 0.000109 | 0.004846 | *Lzts1* | 8 |
| ENSMUSG00000046338 | 79.77 | 166.12 | 93.002 | 355.07 | 737.26 | 290.7 | 112.97 | 461.01 | -2.0287 | 0.000109 | 0.004848 | *Gpat2* | 2 |
| ENSMUSG00000047013 | 70.323 | 91.962 | 45.967 | 122.44 | 286.23 | 118.47 | 69.417 | 175.71 | -1.3408 | 0.00011 | 0.004906 | *Fbxo41* | 6 |
| ENSMUSG00000033147 | 358.96 | 289.73 | 292.9 | 191.82 | 176.08 | 206.08 | 313.87 | 191.32 | 0.7149 | 0.000112 | 0.004983 | *Slc22a15* | 3 |
| ENSMUSG00000061666 | 192.08 | 277.86 | 198.83 | 308.14 | 538.63 | 370.34 | 222.92 | 405.7 | -0.8645 | 0.000113 | 0.004997 | *Gdpd1* | 11 |
| ENSMUSG00000020380 | 939.4 | 757.45 | 707.67 | 1078.5 | 1386.9 | 1142.9 | 801.51 | 1202.8 | -0.5861 | 0.000114 | 0.005019 | *Rad50* | 11 |
| ENSMUSG00000031546 | 795.6 | 774.26 | 847.71 | 1023.4 | 1246.4 | 1162.8 | 805.86 | 1144.2 | -0.5064 | 0.000114 | 0.005035 | *Gins4* | 8 |
| ENSMUSG00000031134 | 3169.8 | 2963.5 | 2822.1 | 2288.6 | 2004.5 | 2293.7 | 2985.2 | 2195.6 | 0.4435 | 0.000119 | 0.005223 | *Rbmx* | X |
| ENSMUSG00000036427 | 2128.6 | 2165.5 | 2341.1 | 3005.9 | 3293.4 | 2756.7 | 2211.7 | 3018.6 | -0.449 | 0.000119 | 0.005223 | *Gpi1* | 7 |
| ENSMUSG00000053559 | 59.827 | 91.962 | 95.14 | 110.19 | 338.27 | 162.27 | 82.31 | 203.58 | -1.309 | 0.00012 | 0.005243 | *Smagp* | 15 |
| ENSMUSG00000032221 | 461.83 | 387.62 | 565.5 | 677.49 | 835.27 | 693.9 | 471.65 | 735.55 | -0.6425 | 0.000122 | 0.005301 | *Mns1* | 9 |

| | | | | | | | | | | | | | |
|---|---|---|---|---|---|---|---|---|---|---|---|---|---|
| ENSMUSG00000022199 | 3373.4 | 3615.2 | 3420.8 | 2677.3 | 2100.8 | 2678 | 3469.8 | 2485.4 | 0.4818 | 0.000122 | 0.00531 | *Slc22a17* | 14 |
| ENSMUSG00000019301 | 4849.2 | 4161 | 5043.5 | 2781.4 | 2187.5 | 3922.5 | 4684.6 | 2963.8 | 0.6607 | 0.000123 | 0.005321 | *Hsd17b1* | 11 |
| ENSMUSG00000068551 | 1189.2 | 1260.8 | 1053 | 804.01 | 675.68 | 903.96 | 1167.6 | 794.55 | 0.5564 | 0.000123 | 0.005321 | *Zfp467* | 6 |
| ENSMUSG00000026049 | 458.68 | 545.84 | 585.81 | 737.69 | 1048.6 | 724.76 | 530.11 | 837.03 | -0.66 | 0.000124 | 0.005357 | *Tex30* | 1 |
| ENSMUSG00000058147 | 20.992 | 59.33 | 55.588 | 177.53 | 404.19 | 143.36 | 45.303 | 241.7 | -2.4156 | 0.000126 | 0.005418 | *Xlr3c* | X |
| ENSMUSG00000019817 | 7768.1 | 7532 | 7647.6 | 6261.7 | 5278.8 | 5684.6 | 7649.2 | 5741.7 | 0.414 | 0.000127 | 0.005467 | *Plagl1* | 10 |
| ENSMUSG00000036395 | 145.89 | 177 | 198.83 | 260.18 | 360.82 | 278.75 | 173.91 | 299.92 | -0.788 | 0.000128 | 0.005472 | *Glb1l2* | 9 |
| ENSMUSG00000021981 | 600.37 | 542.87 | 618.95 | 831.56 | 1052.1 | 763.59 | 587.4 | 882.42 | -0.5882 | 0.00013 | 0.005541 | *Cab39l* | 14 |
| ENSMUSG00000020696 | 130.15 | 234.35 | 164.62 | 270.38 | 429.35 | 298.66 | 176.38 | 332.8 | -0.9162 | 0.00013 | 0.005544 | *Rffl* | 11 |
| ENSMUSG00000070476 | 67.175 | 80.096 | 82.312 | 128.56 | 228.98 | 128.43 | 76.528 | 161.99 | -1.0849 | 0.000131 | 0.005583 | *Fam217b* | 2 |
| ENSMUSG00000073294 | 15.744 | 17.799 | 14.966 | 52.036 | 65.052 | 38.826 | 16.17 | 51.972 | -1.6856 | 0.000133 | 0.005649 | *AU022751* | X |
| ENSMUSG00000045763 | 1113.6 | 1310.2 | 1326.6 | 2204.9 | 1506.6 | 1871.6 | 1250.2 | 1861 | -0.5735 | 0.000134 | 0.005649 | *Basp1* | 15 |
| ENSMUSG00000041729 | 506.96 | 739.65 | 580.46 | 369.35 | 214.24 | 432.07 | 609.02 | 338.55 | 0.8494 | 0.000134 | 0.005649 | *Coro2b* | 9 |
| ENSMUSG00000026576 | 573.08 | 684.27 | 643.53 | 881.55 | 894.25 | 926.86 | 633.63 | 900.89 | -0.5074 | 0.000136 | 0.005732 | *Atp1b1* | 1 |
| ENSMUSG00000039987 | 540.55 | 628.9 | 527.01 | 752.99 | 995.73 | 796.44 | 565.49 | 848.39 | -0.5857 | 0.000136 | 0.005732 | *Phtf2* | 5 |
| ENSMUSG00000074505 | 392.55 | 330.27 | 422.25 | 274.46 | 185.62 | 230.97 | 381.69 | 230.35 | 0.7304 | 0.000141 | 0.005919 | *Fat3* | 9 |
| ENSMUSG00000031654 | 6081.4 | 5226 | 7251 | 2164.1 | 1519.6 | 3410.8 | 6186.1 | 2364.8 | 1.3874 | 0.000144 | 0.006018 | *Cbln1* | 8 |
| ENSMUSG00000053289 | 812.39 | 1067.9 | 925.75 | 1144.8 | 2270.8 | 1364.9 | 935.36 | 1593.5 | -0.7691 | 0.000145 | 0.006069 | *Ddx10* | 9 |
| ENSMUSG00000026249 | 6639.8 | 8302.3 | 6501.6 | 5160.8 | 5256.2 | 5185.8 | 7147.9 | 5200.9 | 0.4588 | 0.000146 | 0.006086 | *Serpine2* | 1 |
| ENSMUSG00000057000 | 120.7 | 138.44 | 62.002 | 338.74 | 227.25 | 153.31 | 107.05 | 239.77 | -1.1612 | 0.000146 | 0.006086 | *Nxf3* | X |
| ENSMUSG00000028214 | 942.54 | 848.42 | 943.92 | 707.08 | 622.77 | 568.46 | 911.63 | 632.77 | 0.5268 | 0.000148 | 0.006106 | *Gem* | 4 |
| ENSMUSG00000050471 | 440.83 | 456.84 | 518.46 | 662.19 | 856.09 | 635.16 | 472.05 | 717.81 | -0.6059 | 0.000148 | 0.006106 | *Fam118b* | 9 |
| ENSMUSG00000022652 | 103.91 | 141.4 | 59.864 | 307.12 | 765.02 | 295.68 | 101.73 | 455.94 | -2.164 | 0.000148 | 0.006106 | *Morc1* | 16 |
| ENSMUSG00000107488 | 8.3968 | 7.9107 | 6.414 | 11.223 | 111.02 | 22.898 | 7.5738 | 48.381 | -2.6806 | 0.000148 | 0.006106 | *Gm31579* | 6 |
| ENSMUSG00000021270 | 11252 | 12569 | 12506 | 15130 | 23387 | 15702 | 12109 | 18073 | -0.5778 | 0.000149 | 0.006127 | *Hsp90aa1* | 12 |
| ENSMUSG00000022610 | 65.075 | 94.928 | 89.795 | 183.66 | 137.91 | 163.27 | 83.266 | 161.61 | -0.9543 | 0.000154 | 0.006312 | *Mapk12* | 15 |
| ENSMUSG00000043753 | 442.93 | 625.93 | 420.11 | 922.37 | 747.67 | 699.87 | 496.33 | 789.97 | -0.6694 | 0.000155 | 0.006355 | *Dmrta1* | 4 |
| ENSMUSG00000019214 | 331.67 | 297.64 | 447.91 | 524.44 | 704.3 | 529.63 | 359.07 | 586.12 | -0.7087 | 0.000156 | 0.006378 | *Chtf18* | 17 |
| ENSMUSG00000110331 | 214.12 | 196.78 | 287.56 | 560.15 | 247.2 | 574.43 | 232.82 | 460.59 | -0.9834 | 0.000157 | 0.006378 | *Nudc-ps1* | 8 |
| ENSMUSG00000028572 | 742.07 | 1033.3 | 692.71 | 1134.6 | 1551.7 | 1187.7 | 822.7 | 1291.3 | -0.6504 | 0.000158 | 0.00642 | *Hook1* | 4 |
| ENSMUSG00000028487 | 168.99 | 217.54 | 107.97 | 288.75 | 416.34 | 253.87 | 164.83 | 319.65 | -0.9553 | 0.00016 | 0.006478 | *Bnc2* | 4 |
| ENSMUSG00000018920 | 325.38 | 265.01 | 286.49 | 173.45 | 196.02 | 167.25 | 292.29 | 178.91 | 0.7065 | 0.00016 | 0.006479 | *Cxcl16* | 11 |
| ENSMUSG00000024366 | 1.0496 | 0 | 2.138 | 7.1422 | 42.501 | 12.942 | 1.0625 | 20.862 | -4.329 | 0.000161 | 0.006487 | *Gfra3* | 18 |
| ENSMUSG00000038576 | 55.629 | 85.04 | 91.933 | 136.72 | 189.95 | 143.36 | 77.534 | 156.68 | -1.0166 | 0.000161 | 0.006487 | *Susd4* | 1 |
| ENSMUSG00000041117 | 2379.5 | 1997.4 | 2149.7 | 1646.8 | 1265.5 | 1647.6 | 2175.5 | 1520 | 0.5178 | 0.000162 | 0.006502 | *Ccdc8* | 7 |
| ENSMUSG00000026418 | 372.61 | 249.19 | 337.8 | 115.3 | 228.98 | 163.27 | 319.87 | 169.18 | 0.9149 | 0.000164 | 0.006578 | *Tnni1* | 1 |
| ENSMUSG00000022066 | 75.572 | 116.68 | 48.105 | 43.874 | 9.541 | 7.9644 | 80.12 | 20.46 | 1.9812 | 0.000165 | 0.006578 | *Entpd4b* | 14 |
| ENSMUSG00000053080 | 2694.3 | 2437.5 | 2407.4 | 1875.3 | 1370.4 | 1947.3 | 2513.1 | 1731 | 0.5383 | 0.000165 | 0.006578 | *2700081O15Rik* | 19 |

| | | | | | | | | | | | | | |
|---|---|---|---|---|---|---|---|---|---|---|---|---|---|
| ENSMUSG00000021710 | 609.82 | 624.94 | 675.6 | 896.86 | 1065.1 | 816.35 | 636.79 | 926.11 | -0.5412 | 0.000168 | 0.006686 | *Nln* | 13 |
| ENSMUSG00000042686 | 36.736 | 29.665 | 34.208 | 64.28 | 147.45 | 63.715 | 33.536 | 91.816 | -1.4585 | 0.000168 | 0.006686 | *Jph1* | 1 |
| ENSMUSG00000036902 | 245.61 | 190.85 | 235.18 | 322.42 | 513.48 | 327.54 | 223.88 | 387.81 | -0.7951 | 0.00017 | 0.006733 | *Neto2* | 8 |
| ENSMUSG00000079019 | 1155.6 | 734.7 | 983.47 | 588.72 | 396.39 | 708.83 | 957.93 | 564.65 | 0.7634 | 0.00017 | 0.006733 | *Insl3* | 8 |
| ENSMUSG00000036620 | 3533 | 3350.2 | 4032.2 | 2306.9 | 2431.2 | 2997.6 | 3638.5 | 2578.6 | 0.4967 | 0.000175 | 0.00688 | *Mgat4b* | 11 |
| ENSMUSG00000020656 | 15.744 | 8.8995 | 17.104 | 36.731 | 86.736 | 33.849 | 13.916 | 52.439 | -1.9233 | 0.000175 | 0.00688 | *Grhl1* | 12 |
| ENSMUSG00000033565 | 2868.6 | 3184 | 2848.9 | 2203.9 | 2011.4 | 2363.4 | 2967.2 | 2192.9 | 0.4366 | 0.000181 | 0.007103 | *Rbfox2* | 15 |
| ENSMUSG00000048899 | 32.538 | 73.174 | 29.932 | 90.808 | 175.21 | 94.577 | 45.214 | 120.2 | -1.409 | 0.000181 | 0.007103 | *Rimkla* | 4 |
| ENSMUSG00000055874 | 2.0992 | 2.9665 | 0 | 14.284 | 27.756 | 17.92 | 1.6886 | 19.987 | -3.5491 | 0.000182 | 0.007103 | *Foxi3* | 6 |
| ENSMUSG00000021182 | 204.67 | 265.01 | 202.04 | 353.03 | 652.26 | 288.71 | 223.91 | 431.33 | -0.9471 | 0.000182 | 0.007103 | *Ccdc88c* | 12 |
| ENSMUSG00000049916 | 14.694 | 22.743 | 27.794 | 51.016 | 99.747 | 48.782 | 21.744 | 66.515 | -1.6177 | 0.000182 | 0.007109 | *2610318N02Rik* | 16 |
| ENSMUSG00000078955 | 0 | 7.9107 | 8.5519 | 29.589 | 71.124 | 13.938 | 5.4875 | 38.217 | -2.8015 | 0.000186 | 0.007227 | *Gm14222* | 2 |
| ENSMUSG00000028953 | 1498.8 | 1381.4 | 1364 | 2071.2 | 2621.2 | 1653.6 | 1414.8 | 2115.3 | -0.5808 | 0.000187 | 0.00724 | *Abcf2* | 5 |
| ENSMUSG00000001661 | 455.53 | 387.62 | 350.63 | 259.16 | 227.25 | 272.78 | 397.93 | 253.06 | 0.6542 | 0.000189 | 0.007317 | *Hoxc6* | 15 |
| ENSMUSG00000045319 | 196.28 | 181.95 | 238.39 | 308.14 | 320.92 | 364.37 | 205.54 | 331.14 | -0.6888 | 0.000191 | 0.007376 | *Proser2* | 2 |
| ENSMUSG00000041577 | 87.117 | 106.79 | 113.31 | 191.82 | 158.73 | 207.07 | 102.41 | 185.87 | -0.8585 | 0.000195 | 0.007507 | *Prelp* | 1 |
| ENSMUSG00000078532 | 97.613 | 116.68 | 83.381 | 164.27 | 267.15 | 155.31 | 99.226 | 195.57 | -0.9803 | 0.000196 | 0.007533 | *Nkain1* | 4 |
| ENSMUSG00000026204 | 45.133 | 72.185 | 43.829 | 19.386 | 6.9389 | 20.907 | 53.716 | 15.744 | 1.7851 | 0.000197 | 0.007533 | *Ptprn* | 1 |
| ENSMUSG00000054727 | 2.0992 | 10.877 | 21.38 | 41.833 | 83.267 | 31.858 | 11.452 | 52.319 | -2.1981 | 0.000197 | 0.007533 | *1700013H16Rik* | X |
| ENSMUSG00000025503 | 1756 | 1439.7 | 1815.2 | 2314.1 | 2621.2 | 2130.5 | 1670.3 | 2355.2 | -0.4962 | 0.000201 | 0.007686 | *Taldo1* | 7 |
| ENSMUSG00000021953 | 5.248 | 2.9665 | 3.207 | 29.589 | 39.031 | 10.951 | 3.8072 | 26.524 | -2.8066 | 0.000202 | 0.007688 | *Tdh* | 14 |
| ENSMUSG00000041974 | 458.68 | 421.24 | 455.39 | 650.96 | 693.02 | 588.37 | 445.1 | 644.12 | -0.5339 | 0.000202 | 0.007688 | *Spidr* | 16 |
| ENSMUSG00000024247 | 4739 | 4205.5 | 5928.6 | 3724.2 | 2683.6 | 3607.9 | 4957.7 | 3338.6 | 0.5706 | 0.000207 | 0.007864 | *Pkdcc* | 17 |
| ENSMUSG00000074830 | 2.0992 | 9.8884 | 3.207 | 10.203 | 62.45 | 31.858 | 5.0648 | 34.837 | -2.7747 | 0.000208 | 0.007864 | *Zfp640* | 13 |
| ENSMUSG00000022415 | 652.85 | 719.87 | 601.84 | 469.35 | 456.23 | 443.02 | 658.19 | 456.2 | 0.5293 | 0.000208 | 0.007864 | *Syngr1* | 15 |
| ENSMUSG00000029848 | 11.546 | 84.051 | 69.485 | 300.99 | 1660.1 | 382.29 | 55.027 | 781.14 | -3.8272 | 0.000209 | 0.007879 | *Stra8* | 6 |
| ENSMUSG00000022119 | 2184.2 | 2419.7 | 2275.9 | 1835.5 | 1537 | 1696.4 | 2293.3 | 1689.6 | 0.4411 | 0.000211 | 0.007938 | *Rbm26* | 14 |
| ENSMUSG00000044312 | 2.0992 | 1.9777 | 0 | 12.244 | 45.103 | 6.9688 | 1.359 | 21.439 | -3.9729 | 0.000214 | 0.00804 | *Neurog3* | 10 |
| ENSMUSG00000025743 | 1728.7 | 1731.5 | 1740.3 | 1384.6 | 949.76 | 1257.4 | 1733.5 | 1197.2 | 0.5348 | 0.000216 | 0.008078 | *Sdc3* | 4 |
| ENSMUSG00000004415 | 895.31 | 755.47 | 1002.7 | 624.43 | 529.09 | 645.12 | 884.5 | 599.55 | 0.5614 | 0.000216 | 0.008078 | *Col26a1* | 5 |
| ENSMUSG00000025020 | 57.728 | 39.553 | 49.174 | 18.366 | 7.8063 | 17.92 | 48.818 | 14.697 | 1.7433 | 0.000218 | 0.008124 | *Slit1* | 19 |
| ENSMUSG00000050197 | 14.694 | 24.721 | 31.001 | 85.707 | 96.277 | 35.84 | 23.472 | 72.608 | -1.6316 | 0.000219 | 0.008162 | *Rhox13* | X |
| ENSMUSG00000057722 | 450.28 | 412.34 | 470.36 | 279.57 | 289.7 | 319.57 | 444.33 | 296.28 | 0.5845 | 0.000223 | 0.008278 | *Lepr* | 4 |
| ENSMUSG00000030652 | 494.36 | 398.5 | 563.36 | 655.04 | 826.6 | 729.74 | 485.41 | 737.13 | -0.6041 | 0.000224 | 0.008307 | *Coq7* | 7 |
| ENSMUSG00000033762 | 239.31 | 202.71 | 194.56 | 326.5 | 458.84 | 289.7 | 212.19 | 358.35 | -0.7578 | 0.000227 | 0.008384 | *Recql4* | 15 |
| ENSMUSG00000039055 | 229.86 | 230.4 | 213.8 | 405.07 | 364.29 | 299.66 | 224.69 | 356.34 | -0.6651 | 0.000227 | 0.008384 | *Eme1* | 11 |
| ENSMUSG00000021758 | 398.85 | 787.11 | 469.29 | 1308 | 3240.5 | 1490.3 | 551.75 | 2013 | -1.8672 | 0.000232 | 0.008574 | *Ddx4* | 13 |
| ENSMUSG00000079259 | 108.11 | 269.95 | 189.21 | 569.34 | 967.11 | 448.99 | 189.09 | 661.81 | -1.8072 | 0.000234 | 0.008608 | *Trim71* | 9 |

| | | | | | | | | | | | | | |
|---|---|---|---|---|---|---|---|---|---|---|---|---|---|
| ENSMUSG00000028702 | 408.3 | 433.11 | 456.46 | 694.84 | 769.35 | 521.67 | 432.62 | 661.95 | -0.6144 | 0.000236 | 0.008671 | *Rad54l* | 4 |
| ENSMUSG00000070407 | 250.86 | 303.57 | 317.49 | 178.56 | 137.91 | 202.1 | 290.64 | 172.85 | 0.7521 | 0.000237 | 0.00868 | *Hs3st3b1* | 11 |
| ENSMUSG00000055322 | 2872.8 | 3692.3 | 3084 | 2270.2 | 2191.8 | 2511.8 | 3216.4 | 2324.6 | 0.4687 | 0.000238 | 0.008717 | *Tns1* | 1 |
| ENSMUSG00000061633 | 0 | 11.866 | 2.138 | 19.386 | 56.379 | 26.88 | 4.668 | 34.215 | -2.8576 | 0.000239 | 0.008717 | *1700029P11Rik* | 15 |
| ENSMUSG00000026568 | 893.21 | 790.08 | 1030.5 | 1165.2 | 1473.7 | 1281.3 | 904.6 | 1306.7 | -0.5314 | 0.00024 | 0.008732 | *Mpc2* | 1 |
| ENSMUSG00000095082 | 0 | 2.9665 | 1.069 | 6.1219 | 39.031 | 16.924 | 1.3452 | 20.693 | -3.9265 | 0.000248 | 0.009028 | *Gm15091* | X |
| ENSMUSG00000026697 | 304.39 | 240.29 | 203.11 | 136.72 | 162.2 | 132.41 | 249.26 | 143.78 | 0.7922 | 0.00025 | 0.009065 | *Myoc* | 1 |
| ENSMUSG00000055407 | 153.24 | 165.14 | 161.42 | 307.12 | 259.34 | 226.99 | 159.93 | 264.48 | -0.7251 | 0.00025 | 0.009075 | *Map6* | 7 |
| ENSMUSG00000027111 | 2129.6 | 2024.1 | 1849.4 | 2869.1 | 2571.7 | 2606.3 | 2001.1 | 2682.4 | -0.4226 | 0.000252 | 0.009101 | *Itga6* | 2 |
| ENSMUSG00000042489 | 730.52 | 643.73 | 731.19 | 940.73 | 1202.2 | 913.91 | 701.82 | 1018.9 | -0.5389 | 0.000254 | 0.009159 | *Clspn* | 4 |
| ENSMUSG00000021477 | 33080 | 34351 | 35037 | 27297 | 21698 | 27191 | 34156 | 25395 | 0.4276 | 0.000257 | 0.009271 | *Ctsl* | 13 |
| ENSMUSG00000114003 | 1475.7 | 1462.5 | 1499.8 | 1030.5 | 1112 | 1155.8 | 1479.3 | 1099.4 | 0.428 | 0.000258 | 0.009275 | *Gm9616* | 13 |
| ENSMUSG00000037731 | 24.141 | 26.699 | 13.897 | 52.036 | 91.941 | 47.786 | 21.579 | 63.921 | -1.5674 | 0.000261 | 0.009345 | *Themis2* | 4 |
| ENSMUSG00000046794 | 201.52 | 290.72 | 259.77 | 87.747 | 82.4 | 193.14 | 250.67 | 121.09 | 1.0522 | 0.000265 | 0.009465 | *Ppp1r3b* | 8 |
| ENSMUSG00000078185 | 334.82 | 367.85 | 302.53 | 477.51 | 698.23 | 445.01 | 335.07 | 540.25 | -0.6902 | 0.000266 | 0.009465 | *Chml* | 1 |
| ENSMUSG00000089715 | 2715.3 | 3037.7 | 3603.6 | 2333.5 | 1949 | 2358.5 | 3118.9 | 2213.6 | 0.4948 | 0.000266 | 0.009465 | *Cbx6* | 15 |
| ENSMUSG00000043029 | 33.587 | 22.743 | 44.898 | 3.0609 | 7.8063 | 10.951 | 33.743 | 7.2728 | 2.2074 | 0.000266 | 0.009465 | *Trpv3* | 11 |
| ENSMUSG00000026090 | 36.736 | 53.397 | 40.622 | 86.727 | 109.29 | 90.595 | 43.585 | 95.537 | -1.1317 | 0.000269 | 0.009552 | *2010300C02Rik* | 1 |
| ENSMUSG00000039676 | 3.1488 | 0.9888 | 3.207 | 17.345 | 30.358 | 14.933 | 2.4482 | 20.879 | -3.1084 | 0.00027 | 0.009566 | *Capsl* | 15 |
| ENSMUSG00000107496 | 4.1984 | 2.9665 | 2.138 | 19.386 | 35.562 | 13.938 | 3.101 | 22.962 | -2.8927 | 0.000272 | 0.009616 | *4933431M02Rik* | 6 |
| ENSMUSG00000024534 | 525.85 | 431.13 | 517.39 | 326.5 | 316.59 | 347.45 | 491.46 | 330.18 | 0.5738 | 0.000273 | 0.009642 | *Sncaip* | 18 |
| ENSMUSG00000006273 | 1563.9 | 1661.2 | 1769.2 | 2198.8 | 2383.5 | 2110.6 | 1664.8 | 2231 | -0.4226 | 0.000277 | 0.009753 | *Atp6v1b2* | 8 |
| ENSMUSG00000033350 | 636.06 | 607.14 | 649.95 | 430.57 | 382.51 | 487.82 | 631.05 | 433.63 | 0.5422 | 0.000278 | 0.009776 | *Chst2* | 9 |
| ENSMUSG00000070691 | 8.3968 | 5.933 | 4.276 | 22.447 | 53.777 | 19.911 | 6.2019 | 32.045 | -2.3746 | 0.000279 | 0.009782 | *Runx3* | 4 |
| ENSMUSG00000049521 | 1042.3 | 1021.5 | 965.3 | 638.72 | 754.61 | 772.55 | 1009.7 | 721.96 | 0.4835 | 0.000281 | 0.009836 | *Cdc42ep1* | 15 |
| ENSMUSG00000067780 | 1422.2 | 1116.4 | 1456 | 932.57 | 836.14 | 1015.5 | 1331.5 | 928.06 | 0.521 | 0.000284 | 0.009924 | *Pi15* | 1 |
| ENSMUSG00000026207 | 591.98 | 564.62 | 572.98 | 419.35 | 333.07 | 423.11 | 576.53 | 391.84 | 0.5586 | 0.000285 | 0.00996 | *Speg* | 1 |
| ENSMUSG00000034472 | 124.9 | 174.03 | 198.83 | 228.55 | 425.01 | 272.78 | 165.92 | 308.78 | -0.8981 | 0.000287 | 0.009992 | *Rasd2* | 8 |
| ENSMUSG00000060445 | 85.018 | 167.11 | 103.69 | 278.55 | 708.64 | 317.58 | 118.61 | 434.92 | -1.8745 | 0.000288 | 0.009992 | *Sycp2* | 2 |
| ENSMUSG00000032006 | 709.53 | 759.43 | 640.33 | 509.14 | 459.7 | 508.73 | 703.09 | 492.52 | 0.5145 | 0.000288 | 0.009992 | *Pdgfd* | 9 |
| ENSMUSG00000002346 | 477.57 | 708.01 | 391.25 | 366.29 | 253.27 | 301.65 | 525.61 | 307.07 | 0.7774 | 0.000291 | 0.010072 | *Slc25a42* | 8 |
| ENSMUSG00000079532 | 4.1984 | 6.9218 | 5.345 | 25.508 | 43.368 | 17.92 | 5.4884 | 28.932 | -2.3992 | 0.000291 | 0.010072 | *Gm6890* | X |
| ENSMUSG00000035284 | 389.4 | 356.97 | 217.01 | 538.73 | 720.78 | 429.08 | 321.13 | 562.86 | -0.81 | 0.000292 | 0.010078 | *Vps13c* | 9 |
| ENSMUSG00000079606 | 11.546 | 26.699 | 12.828 | 44.894 | 91.941 | 37.831 | 17.024 | 58.222 | -1.7727 | 0.000297 | 0.010225 | *Gm595* | X |
| ENSMUSG00000053007 | 323.28 | 451.9 | 349.56 | 232.63 | 200.36 | 263.82 | 374.91 | 232.27 | 0.6928 | 0.000299 | 0.010247 | *Creb5* | 6 |
| ENSMUSG00000031431 | 795.6 | 849.41 | 904.37 | 1037.7 | 1572.5 | 1179.7 | 849.79 | 1263.3 | -0.5728 | 0.000299 | 0.010247 | *Tsc22d3* | X |
| ENSMUSG00000071773 | 4.1984 | 8.8995 | 5.345 | 28.569 | 37.297 | 22.898 | 6.1476 | 29.588 | -2.2628 | 0.000305 | 0.010458 | *Rhox1* | X |
| ENSMUSG00000023411 | 617.17 | 732.73 | 832.75 | 474.45 | 423.27 | 555.52 | 727.55 | 484.41 | 0.5875 | 0.000306 | 0.010464 | *Nfatc4* | 14 |

| | | | | | | | | | | | | | |
|---|---|---|---|---|---|---|---|---|---|---|---|---|---|
| ENSMUSG00000034173 | 375.76 | 334.23 | 386.98 | 461.18 | 762.41 | 533.61 | 365.65 | 585.74 | -0.6815 | 0.000308 | 0.010506 | *Zbed5* | 5 |
| ENSMUSG00000070780 | 96.564 | 111.74 | 91.933 | 49.995 | 45.97 | 49.777 | 100.08 | 48.581 | 1.0446 | 0.00031 | 0.010542 | *Rbm47* | 5 |
| ENSMUSG00000034957 | 1706.7 | 1654.3 | 1770.3 | 1122.3 | 620.17 | 1355.9 | 1710.4 | 1032.8 | 0.7286 | 0.000314 | 0.010655 | *Cebpa* | 7 |
| ENSMUSG00000051855 | 3316.7 | 2719.3 | 3339.5 | 2333.5 | 2353.2 | 2266.9 | 3125.2 | 2317.8 | 0.431 | 0.000318 | 0.01075 | *Mest* | 6 |
| ENSMUSG00000054405 | 1825.3 | 1924.3 | 2180.7 | 2398.8 | 3177.2 | 2707.9 | 1976.8 | 2761.3 | -0.4826 | 0.000318 | 0.01075 | *Dnajc8* | 4 |
| ENSMUSG00000059493 | 615.07 | 546.83 | 586.88 | 413.23 | 410.26 | 402.2 | 582.92 | 408.56 | 0.5125 | 0.000319 | 0.010764 | *Nhs* | X |
| ENSMUSG00000039774 | 89.216 | 119.65 | 83.381 | 151.01 | 289.7 | 152.32 | 97.416 | 197.68 | -1.0224 | 0.000322 | 0.010867 | *Galnt12* | 4 |
| ENSMUSG00000047242 | 244.56 | 298.63 | 225.56 | 547.91 | 1135.4 | 477.86 | 256.25 | 720.38 | -1.4915 | 0.000323 | 0.010876 | *Taf9b* | X |
| ENSMUSG00000028927 | 633.96 | 595.28 | 365.6 | 975.42 | 910.73 | 704.85 | 531.61 | 863.67 | -0.6997 | 0.000325 | 0.010876 | *Padi2* | 4 |
| ENSMUSG00000036030 | 28.339 | 37.576 | 24.587 | 66.321 | 143.98 | 47.786 | 30.167 | 86.03 | -1.5142 | 0.000325 | 0.010876 | *Prtg* | 9 |
| ENSMUSG00000046711 | 231.96 | 439.04 | 342.08 | 714.22 | 1640.2 | 857.17 | 337.69 | 1070.5 | -1.6646 | 0.000325 | 0.010876 | *Hmga1* | 17 |
| ENSMUSG00000028776 | 542.65 | 484.53 | 575.12 | 334.66 | 301.84 | 418.13 | 534.1 | 351.55 | 0.6042 | 0.000326 | 0.010907 | *Tinagl1* | 4 |
| ENSMUSG00000051769 | 6.2976 | 18.788 | 22.449 | 39.792 | 71.124 | 48.782 | 15.845 | 53.233 | -1.7508 | 0.000331 | 0.011027 | *Wfdc15a* | 2 |
| ENSMUSG00000026950 | 87.117 | 105.81 | 70.554 | 175.49 | 180.41 | 136.39 | 87.825 | 164.1 | -0.9008 | 0.000331 | 0.011027 | *Neb* | 2 |
| ENSMUSG00000040749 | 250.86 | 291.71 | 330.32 | 382.62 | 621.9 | 429.08 | 290.96 | 477.87 | -0.7175 | 0.000338 | 0.011232 | *Siah1b* | X |
| ENSMUSG00000006494 | 1044.4 | 1341.8 | 1067.9 | 1559 | 1958.5 | 1482.4 | 1151.4 | 1666.6 | -0.5337 | 0.000339 | 0.011256 | *Pdk1* | 2 |
| ENSMUSG00000116295 | 13.645 | 26.699 | 11.759 | 48.975 | 75.461 | 40.818 | 17.367 | 55.084 | -1.6624 | 0.00034 | 0.011269 | *Gm32885* | 15 |
| ENSMUSG00000031026 | 16.794 | 16.81 | 10.69 | 71.422 | 38.164 | 36.835 | 14.765 | 48.807 | -1.7198 | 0.000341 | 0.011285 | *Trim66* | 7 |
| ENSMUSG00000022325 | 403.05 | 338.18 | 385.91 | 688.71 | 632.31 | 443.02 | 375.71 | 588.01 | -0.6468 | 0.000344 | 0.01134 | *Pop1* | 15 |
| ENSMUSG00000051827 | 4.1984 | 1.9777 | 5.345 | 22.447 | 23.419 | 23.893 | 3.8404 | 23.253 | -2.6077 | 0.000346 | 0.011403 | *Rhox2a* | X |
| ENSMUSG00000025287 | 487.02 | 459.81 | 520.6 | 584.64 | 993.13 | 724.76 | 489.14 | 767.51 | -0.6513 | 0.000348 | 0.011452 | *Acot9* | X |
| ENSMUSG00000010277 | 3323 | 3483.7 | 3057.3 | 4750.6 | 4823.4 | 3836.8 | 3288 | 4470.3 | -0.4432 | 0.000352 | 0.011535 | *2610507B11Rik* | 11 |
| ENSMUSG00000114865 | 2.0992 | 3.9553 | 3.207 | 19.386 | 46.838 | 7.9644 | 3.0872 | 24.729 | -3.0044 | 0.000355 | 0.011628 | *Gm29776* | 14 |
| ENSMUSG00000044313 | 361.06 | 218.53 | 198.83 | 141.82 | 90.206 | 168.25 | 259.48 | 133.43 | 0.9623 | 0.000363 | 0.011857 | *Mab21l3* | 3 |
| ENSMUSG00000020549 | 457.63 | 516.17 | 556.95 | 695.86 | 840.48 | 681.95 | 510.25 | 739.43 | -0.536 | 0.000364 | 0.011878 | *Elac2* | 11 |
| ENSMUSG00000054418 | 11.546 | 8.8995 | 9.6209 | 33.67 | 50.307 | 27.875 | 10.022 | 37.284 | -1.9003 | 0.000367 | 0.011957 | *2900041M22Rik* | 11 |
| ENSMUSG00000024989 | 897.41 | 777.22 | 863.75 | 1140.7 | 1382.6 | 1066.2 | 846.13 | 1196.5 | -0.5007 | 0.000368 | 0.011957 | *Cep55* | 19 |
| ENSMUSG00000000325 | 295.99 | 266.99 | 277.94 | 344.87 | 587.21 | 434.06 | 280.3 | 455.38 | -0.7019 | 0.000371 | 0.012026 | *Arvcf* | 16 |
| ENSMUSG00000053825 | 59.827 | 56.364 | 74.83 | 23.467 | 27.756 | 24.889 | 63.674 | 25.371 | 1.3242 | 0.000371 | 0.012026 | *Ppfia2* | 10 |
| ENSMUSG00000024180 | 146.94 | 150.3 | 164.62 | 292.83 | 315.72 | 190.15 | 153.96 | 266.23 | -0.7915 | 0.000373 | 0.012055 | *Tmem8* | 17 |
| ENSMUSG00000023169 | 965.64 | 934.45 | 820.99 | 385.68 | 436.28 | 789.47 | 907.02 | 537.14 | 0.7564 | 0.000373 | 0.012055 | *Slc38a1* | 15 |
| ENSMUSG00000029366 | 431.39 | 443 | 552.67 | 660.14 | 769.35 | 657.06 | 475.68 | 695.52 | -0.5491 | 0.000374 | 0.012064 | *Dck* | 5 |
| ENSMUSG00000053441 | 619.27 | 575.5 | 686.29 | 478.53 | 392.92 | 426.1 | 627.02 | 432.51 | 0.5364 | 0.000375 | 0.012077 | *Adamts19* | 18 |
| ENSMUSG00000037108 | 362.11 | 433.11 | 394.46 | 497.91 | 922.88 | 557.51 | 396.56 | 659.43 | -0.7348 | 0.000377 | 0.012088 | *Zcwpw1* | 5 |
| ENSMUSG00000032058 | 679.09 | 826.67 | 717.29 | 981.54 | 1465.8 | 948.76 | 741.02 | 1132 | -0.612 | 0.000377 | 0.012088 | *Ppp2r1b* | 9 |
| ENSMUSG00000109032 | 27.29 | 56.364 | 37.415 | 82.646 | 156.99 | 68.693 | 40.356 | 102.78 | -1.3495 | 0.000381 | 0.012181 | *Gm7972* | 9 |
| ENSMUSG00000034023 | 534.25 | 514.19 | 548.39 | 737.69 | 1077.3 | 657.06 | 532.28 | 824.01 | -0.6317 | 0.000382 | 0.012199 | *Fancd2* | 6 |
| ENSMUSG00000100816 | 3.1488 | 0 | 2.138 | 5.1016 | 49.44 | 12.942 | 1.7623 | 22.495 | -3.6986 | 0.000383 | 0.012199 | *Gm28321* | 1 |

| | | | | | | | | | | | | | |
|---|---|---|---|---|---|---|---|---|---|---|---|---|---|
| ENSMUSG00000032380 | 29.389 | 37.576 | 32.07 | 52.036 | 156.99 | 64.711 | 33.011 | 91.247 | -1.4706 | 0.000383 | 0.012199 | *Dapk2* | 9 |
| ENSMUSG00000020773 | 4329.6 | 5691.7 | 4756 | 3336.4 | 2356.6 | 4045.9 | 4925.8 | 3246.3 | 0.6018 | 0.000385 | 0.012241 | *Trim47* | 11 |
| ENSMUSG00000025081 | 136.45 | 215.57 | 110.11 | 412.21 | 929.81 | 347.45 | 154.04 | 563.16 | -1.8702 | 0.000388 | 0.012318 | *Tdrd1* | 19 |
| ENSMUSG00000020908 | 8.3968 | 19.777 | 9.6209 | 49.995 | 59.848 | 26.88 | 12.598 | 45.574 | -1.8508 | 0.000391 | 0.012374 | *Myh3* | 11 |
| ENSMUSG00000035834 | 696.94 | 859.3 | 667.05 | 985.63 | 1195.2 | 1008.5 | 741.1 | 1063.1 | -0.5206 | 0.000392 | 0.012374 | *Polr3g* | 13 |
| ENSMUSG00000000915 | 109.16 | 173.05 | 140.04 | 224.47 | 387.71 | 196.12 | 140.75 | 269.43 | -0.9381 | 0.000395 | 0.012448 | *Hip1r* | 5 |
| ENSMUSG00000028799 | 1832.6 | 2331.7 | 2195.7 | 1384.6 | 1229.9 | 1771.1 | 2120 | 1461.9 | 0.5367 | 0.000402 | 0.012658 | *Zfp362* | 4 |
| ENSMUSG00000066000 | 209.92 | 237.32 | 262.97 | 357.11 | 490.06 | 311.61 | 236.74 | 386.26 | -0.708 | 0.000403 | 0.012664 | *Zfp979* | 4 |
| ENSMUSG00000045349 | 39.885 | 44.498 | 35.277 | 82.646 | 124.03 | 67.697 | 39.886 | 91.459 | -1.1994 | 0.000404 | 0.012675 | *Sh2d5* | 4 |
| ENSMUSG00000038128 | 12.595 | 26.699 | 24.587 | 55.097 | 84.134 | 44.8 | 21.294 | 61.344 | -1.528 | 0.000411 | 0.012887 | *Camk4* | 18 |
| ENSMUSG00000061048 | 1515.6 | 1541.6 | 1553.2 | 1111.1 | 1182.2 | 1192.7 | 1536.8 | 1162 | 0.4032 | 0.000413 | 0.012909 | *Cdh3* | 8 |
| ENSMUSG00000031584 | 1315.2 | 1640.5 | 1456 | 1700.9 | 2487.6 | 2192.2 | 1470.5 | 2126.9 | -0.5326 | 0.000414 | 0.012909 | *Gsr* | 8 |
| ENSMUSG00000033644 | 234.06 | 405.42 | 196.69 | 871.35 | 1832.7 | 604.3 | 278.73 | 1102.8 | -1.9842 | 0.000414 | 0.012909 | *Piwil2* | 14 |
| ENSMUSG00000037280 | 36.736 | 71.196 | 43.829 | 94.889 | 182.15 | 84.622 | 50.587 | 120.55 | -1.2535 | 0.000428 | 0.013311 | *Galnt6* | 15 |
| ENSMUSG00000079008 | 11.546 | 18.788 | 9.6209 | 28.569 | 65.052 | 43.804 | 13.318 | 45.808 | -1.7813 | 0.00044 | 0.01365 | *Gm14124* | 2 |
| ENSMUSG00000043065 | 618.22 | 625.93 | 576.19 | 766.26 | 968.85 | 832.28 | 606.78 | 855.79 | -0.4967 | 0.000441 | 0.01365 | *Spice1* | 16 |
| ENSMUSG00000018166 | 72.423 | 107.78 | 72.692 | 129.58 | 186.48 | 163.27 | 84.299 | 159.78 | -0.922 | 0.000441 | 0.01365 | *Erbb3* | 10 |
| ENSMUSG00000031303 | 74.522 | 81.084 | 40.622 | 103.05 | 274.95 | 104.53 | 65.409 | 160.85 | -1.2997 | 0.000443 | 0.013676 | *Map3k15* | X |
| ENSMUSG00000038550 | 55.629 | 46.475 | 39.553 | 91.828 | 117.96 | 87.608 | 47.219 | 99.133 | -1.0718 | 0.000444 | 0.013691 | *Ciart* | 3 |
| ENSMUSG00000021175 | 660.2 | 668.45 | 408.36 | 898.9 | 993.13 | 804.4 | 579 | 898.81 | -0.6342 | 0.000454 | 0.013989 | *Cdca7l* | 12 |
| ENSMUSG00000021831 | 554.19 | 575.5 | 562.29 | 703 | 927.21 | 781.51 | 563.99 | 803.91 | -0.5122 | 0.000457 | 0.014051 | *Ero1l* | 14 |
| ENSMUSG00000040183 | 115.46 | 118.66 | 106.9 | 157.13 | 300.98 | 180.19 | 113.67 | 212.77 | -0.907 | 0.00046 | 0.014133 | *Ankrd6* | 4 |
| ENSMUSG00000028496 | 257.15 | 255.12 | 249.08 | 367.31 | 366.03 | 393.24 | 253.78 | 375.53 | -0.5651 | 0.000462 | 0.014156 | *Mllt3* | 4 |
| ENSMUSG00000035131 | 117.56 | 88.006 | 122.93 | 66.321 | 25.154 | 51.769 | 109.5 | 47.748 | 1.2046 | 0.000466 | 0.014251 | *Brinp3* | 1 |
| ENSMUSG00000039414 | 558.39 | 517.16 | 389.11 | 1026.4 | 777.16 | 574.43 | 488.22 | 792.68 | -0.6987 | 0.000473 | 0.014442 | *Heatr5b* | 17 |
| ENSMUSG00000039990 | 656 | 680.32 | 585.81 | 882.57 | 1129.3 | 794.45 | 640.71 | 935.44 | -0.5466 | 0.000474 | 0.014457 | *Edrf1* | 7 |
| ENSMUSG00000107879 | 37.786 | 28.676 | 28.863 | 62.239 | 135.31 | 53.76 | 31.775 | 83.769 | -1.4037 | 0.000476 | 0.014495 | *9330102E08Rik* | 6 |
| ENSMUSG00000041649 | 201.52 | 240.29 | 168.9 | 355.07 | 453.63 | 247.89 | 203.57 | 352.2 | -0.7914 | 0.000484 | 0.014717 | *Klf8* | X |
| ENSMUSG00000029403 | 64.026 | 97.895 | 58.795 | 123.46 | 155.26 | 148.34 | 73.572 | 142.35 | -0.9505 | 0.000491 | 0.01486 | *Cdkl2* | 5 |
| ENSMUSG00000048142 | 37.786 | 44.498 | 29.932 | 66.321 | 113.62 | 78.648 | 37.405 | 86.198 | -1.2057 | 0.000492 | 0.01486 | *Nat8l* | 5 |
| ENSMUSG00000019849 | 1255.3 | 1200.4 | 1243.2 | 1572.3 | 1664.5 | 1622.7 | 1233 | 1619.8 | -0.3939 | 0.000492 | 0.01486 | *Prep* | 10 |
| ENSMUSG00000064128 | 399.9 | 456.84 | 405.15 | 521.38 | 796.24 | 602.31 | 420.63 | 639.98 | -0.6064 | 0.000494 | 0.01492 | *Cenpj* | 14 |
| ENSMUSG00000032826 | 828.14 | 774.26 | 673.47 | 540.77 | 477.05 | 578.41 | 758.62 | 532.08 | 0.5127 | 0.000495 | 0.01492 | *Ank2* | 3 |
| ENSMUSG00000041141 | 368.41 | 343.13 | 329.25 | 232.63 | 212.5 | 245.9 | 346.93 | 230.35 | 0.5919 | 0.000496 | 0.01492 | *Pnmal1* | 7 |
| ENSMUSG00000050700 | 4906.9 | 4982.7 | 4842.5 | 3564 | 2781.6 | 4165.4 | 4910.7 | 3503.7 | 0.4873 | 0.000499 | 0.014994 | *Emilin3* | 2 |
| ENSMUSG00000027018 | 1539.8 | 1826.4 | 2027.9 | 2222.2 | 3370.6 | 2378.4 | 1798 | 2657.1 | -0.5638 | 0.0005 | 0.014999 | *Hat1* | 2 |
| ENSMUSG00000029648 | 159.54 | 187.88 | 179.59 | 117.34 | 95.41 | 92.586 | 175.67 | 101.78 | 0.7892 | 0.000506 | 0.015158 | *Flt1* | 5 |
| ENSMUSG00000074817 | 5.248 | 12.855 | 7.483 | 24.488 | 43.368 | 34.844 | 8.5286 | 34.233 | -2.0014 | 0.00051 | 0.01523 | *Papolb* | 5 |

| | | | | | | | | | | | | | |
|---|---|---|---|---|---|---|---|---|---|---|---|---|---|
| ENSMUSG00000025878 | 660.2 | 622.97 | 633.91 | 804.01 | 1138 | 844.23 | 639.03 | 928.74 | -0.5404 | 0.000511 | 0.01523 | *Uimc1* | 13 |
| ENSMUSG00000021573 | 204.67 | 266 | 235.18 | 145.91 | 130.97 | 155.31 | 235.28 | 144.06 | 0.7096 | 0.000516 | 0.015351 | *Tppp* | 13 |
| ENSMUSG00000006154 | 22.042 | 18.788 | 20.311 | 32.65 | 100.61 | 51.769 | 20.38 | 61.678 | -1.6035 | 0.000516 | 0.015351 | *Eps8l1* | 7 |
| ENSMUSG00000000215 | 14.694 | 19.777 | 7.483 | 39.792 | 47.705 | 46.791 | 13.985 | 44.763 | -1.6732 | 0.000521 | 0.015478 | *Ins2* | 7 |
| ENSMUSG00000031963 | 334.82 | 220.51 | 382.7 | 137.74 | 191.69 | 207.07 | 312.68 | 178.83 | 0.8041 | 0.000523 | 0.015505 | *Bmper* | 9 |
| ENSMUSG00000041406 | 852.28 | 855.34 | 792.12 | 1080.5 | 1573.4 | 1038.4 | 833.25 | 1230.8 | -0.5635 | 0.000526 | 0.015565 | *BC055324* | 1 |
| ENSMUSG00000005370 | 1935.5 | 2049.9 | 2109.1 | 2501.8 | 3107.8 | 2580.5 | 2031.5 | 2730 | -0.4267 | 0.000529 | 0.01562 | *Msh6* | 17 |
| ENSMUSG00000058741 | 29.389 | 25.71 | 14.966 | 32.65 | 117.09 | 64.711 | 23.355 | 71.485 | -1.6169 | 0.000532 | 0.015694 | *Prr19* | 7 |
| ENSMUSG00000087166 | 18.893 | 13.844 | 26.725 | 3.0609 | 2.6021 | 2.9866 | 19.82 | 2.8832 | 2.7833 | 0.000535 | 0.015757 | *L1td1* | 4 |
| ENSMUSG00000050394 | 857.53 | 765.36 | 749.36 | 553.01 | 431.08 | 631.18 | 790.75 | 538.42 | 0.5557 | 0.00054 | 0.015869 | *Armcx6* | X |
| ENSMUSG00000024081 | 1371.8 | 1351.7 | 1404.7 | 1691.7 | 2297.6 | 1757.1 | 1376.1 | 1915.5 | -0.4777 | 0.000542 | 0.015893 | *Cebpz* | 17 |
| ENSMUSG00000002289 | 1623.7 | 1366.6 | 1113.9 | 898.9 | 782.36 | 1090.1 | 1368.1 | 923.8 | 0.5671 | 0.000547 | 0.016021 | *Angptl4* | 17 |
| ENSMUSG00000074704 | 34.637 | 37.576 | 31.001 | 49.995 | 133.57 | 76.657 | 34.404 | 86.742 | -1.3377 | 0.00055 | 0.01608 | *Rad21l* | 2 |
| ENSMUSG00000032511 | 41.984 | 40.542 | 35.277 | 41.833 | 185.62 | 95.573 | 39.268 | 107.67 | -1.4589 | 0.000551 | 0.016094 | *Scn5a* | 9 |
| ENSMUSG00000020836 | 65.075 | 73.174 | 58.795 | 35.711 | 16.48 | 25.884 | 65.681 | 26.025 | 1.3451 | 0.000557 | 0.016228 | *Coro6* | 11 |
| ENSMUSG00000072930 | 0 | 4.9442 | 1.069 | 2.0406 | 45.97 | 25.884 | 2.0044 | 24.632 | -3.6012 | 0.000573 | 0.016674 | *Gm15107* | X |
| ENSMUSG00000025791 | 806.1 | 880.06 | 881.92 | 1083.6 | 1260.3 | 1134.9 | 856.03 | 1159.6 | -0.4383 | 0.000574 | 0.016674 | *Pgm2* | 4 |
| ENSMUSG00000103408 | 2.0992 | 8.8995 | 1.069 | 19.386 | 37.297 | 19.911 | 4.0226 | 25.531 | -2.6505 | 0.000576 | 0.016714 | *Gm37933* | 3 |
| ENSMUSG00000029730 | 3232.8 | 2972.4 | 3261.5 | 4315.9 | 4703.7 | 3661.6 | 3155.6 | 4227.1 | -0.422 | 0.000582 | 0.01687 | *Mcm7* | 5 |
| ENSMUSG00000015869 | 1546.1 | 1878.8 | 1746.7 | 2164.1 | 2364.4 | 2364.4 | 1723.9 | 2297.7 | -0.4145 | 0.000588 | 0.017021 | *Prpsap1* | 11 |
| ENSMUSG00000069170 | 36.736 | 39.553 | 29.932 | 59.178 | 98.012 | 84.622 | 35.407 | 80.604 | -1.188 | 0.000596 | 0.017199 | *Adgrv1* | 13 |
| ENSMUSG00000054994 | 11.546 | 18.788 | 16.035 | 33.67 | 91.941 | 31.858 | 15.456 | 52.49 | -1.7678 | 0.0006 | 0.017292 | *AV320801* | X |
| ENSMUSG00000017737 | 213.07 | 200.73 | 85.519 | 57.138 | 98.012 | 74.666 | 166.44 | 76.605 | 1.117 | 0.000603 | 0.017361 | *Mmp9* | 2 |
| ENSMUSG00000033799 | 1235.4 | 1367.6 | 832.75 | 1548.8 | 2256.9 | 1541.1 | 1145.2 | 1782.3 | -0.6382 | 0.000607 | 0.017414 | *Fam208b* | 13 |
| ENSMUSG00000021904 | 367.36 | 275.88 | 354.91 | 245.9 | 175.21 | 204.09 | 332.72 | 208.4 | 0.6763 | 0.000607 | 0.017414 | *Sema3g* | 14 |
| ENSMUSG00000079626 | 1.0496 | 10.877 | 4.276 | 29.589 | 26.021 | 28.871 | 5.4009 | 28.16 | -2.3679 | 0.000616 | 0.017635 | *Rhox7b* | X |
| ENSMUSG00000032598 | 564.69 | 611.1 | 642.46 | 402 | 405.93 | 472.89 | 606.08 | 426.94 | 0.5058 | 0.000617 | 0.01765 | *Nckipsd* | 9 |
| ENSMUSG00000054079 | 602.47 | 654.61 | 720.5 | 810.13 | 1177 | 900.97 | 659.19 | 962.71 | -0.5474 | 0.000622 | 0.017759 | *Utp18* | 11 |
| ENSMUSG00000073568 | 45.133 | 69.218 | 66.278 | 98.971 | 258.47 | 82.631 | 60.21 | 146.69 | -1.2878 | 0.000625 | 0.017813 | *Arl14epl* | 18 |
| ENSMUSG00000036352 | 923.65 | 914.67 | 1051.9 | 1215.2 | 1409.5 | 1288.2 | 963.4 | 1304.3 | -0.4376 | 0.000626 | 0.017813 | *Ubac1* | 2 |
| ENSMUSG00000027510 | 475.47 | 535.95 | 422.25 | 624.43 | 1003.5 | 623.21 | 477.89 | 750.4 | -0.6518 | 0.000634 | 0.017997 | *Rbm38* | 2 |
| ENSMUSG00000027463 | 1282.6 | 1228.1 | 1134.2 | 1837.6 | 1509.2 | 1580.9 | 1215 | 1642.6 | -0.4346 | 0.000634 | 0.017997 | *Slc52a3* | 2 |
| ENSMUSG00000040950 | 17.843 | 9.8884 | 8.5519 | 23.467 | 69.389 | 38.826 | 12.095 | 43.894 | -1.8657 | 0.000639 | 0.018093 | *Mgl2* | 11 |
| ENSMUSG00000064370 | 59537 | 53554 | 51133 | 66104 | 85415 | 68790 | 54741 | 73436 | -0.4239 | 0.00064 | 0.018102 | *mt-Cytb* | MT |
| ENSMUSG00000028391 | 98.663 | 110.75 | 79.106 | 173.45 | 195.16 | 143.36 | 96.173 | 170.66 | -0.8273 | 0.000641 | 0.018102 | *Wdr31* | 4 |
| ENSMUSG00000052928 | 687.49 | 873.14 | 782.5 | 584.64 | 425.01 | 582.4 | 781.05 | 530.68 | 0.5591 | 0.000642 | 0.018115 | *Ctif* | 18 |
| ENSMUSG00000062995 | 86.068 | 88.995 | 101.55 | 146.93 | 244.6 | 132.41 | 92.206 | 174.64 | -0.9249 | 0.000647 | 0.018212 | *Ica1* | 6 |
| ENSMUSG00000026098 | 228.81 | 220.51 | 187.07 | 322.42 | 384.24 | 285.72 | 212.13 | 330.8 | -0.6418 | 0.000651 | 0.018253 | *Pms1* | 1 |

| | | | | | | | | | | | | | |
|---|---|---|---|---|---|---|---|---|---|---|---|---|---|
| ENSMUSG00000020255 | 1173.5 | 1129.2 | 1185.5 | 1432.5 | 1903 | 1485.4 | 1162.7 | 1607 | -0.4675 | 0.000651 | 0.018253 | *D10Wsu102e* | 10 |
| ENSMUSG00000027855 | 74.522 | 251.16 | 131.49 | 421.39 | 945.43 | 509.72 | 152.39 | 625.51 | -2.037 | 0.000651 | 0.018253 | *Sycp1* | 3 |
| ENSMUSG00000092203 | 857.53 | 678.34 | 811.37 | 539.75 | 598.48 | 523.66 | 782.41 | 553.96 | 0.4972 | 0.000656 | 0.018359 | *1110038B12Rik* | 17 |
| ENSMUSG00000028577 | 1872.5 | 2199.2 | 2053.5 | 2492.6 | 3362.8 | 2617.3 | 2041.7 | 2824.2 | -0.4683 | 0.000657 | 0.018368 | *Plaa* | 4 |
| ENSMUSG00000004667 | 1286.8 | 1212.3 | 1476.3 | 1838.6 | 2184.9 | 1568 | 1325.1 | 1863.8 | -0.4927 | 0.000661 | 0.018429 | *Polr2e* | 10 |
| ENSMUSG00000041642 | 1004.5 | 1110.5 | 1096.8 | 860.13 | 666.14 | 792.46 | 1070.6 | 772.91 | 0.471 | 0.000665 | 0.018511 | *Kif21b* | 1 |
| ENSMUSG00000027173 | 59.827 | 72.185 | 102.62 | 118.36 | 262.81 | 122.45 | 78.212 | 167.87 | -1.1059 | 0.000671 | 0.018648 | *Depdc7* | 2 |
| ENSMUSG00000029361 | 20.992 | 28.676 | 17.104 | 52.036 | 60.716 | 58.737 | 22.257 | 57.163 | -1.358 | 0.00068 | 0.018888 | *Nos1* | 5 |
| ENSMUSG00000078190 | 2187.4 | 1790.8 | 1675.1 | 1465.2 | 1419.9 | 1143.9 | 1884.4 | 1343 | 0.4885 | 0.000683 | 0.018946 | *Dnm3os* | 1 |
| ENSMUSG00000109764 | 68.224 | 72.185 | 120.8 | 29.589 | 19.949 | 53.76 | 87.068 | 34.433 | 1.3434 | 0.000685 | 0.018962 | *Klkb1* | 8 |
| ENSMUSG00000029656 | 7.3472 | 9.8884 | 3.207 | 16.325 | 46.838 | 30.862 | 6.8142 | 31.342 | -2.1989 | 0.000687 | 0.018993 | *C8b* | 4 |
| ENSMUSG00000001674 | 1072.7 | 1196.5 | 1271 | 1463.1 | 1936.8 | 1528.2 | 1180.1 | 1642.7 | -0.4777 | 0.000698 | 0.019256 | *Ddx18* | 1 |
| ENSMUSG00000041245 | 554.19 | 673.4 | 373.08 | 1136.6 | 808.38 | 685.93 | 533.56 | 876.98 | -0.716 | 0.000702 | 0.019342 | *Wnk3* | X |
| ENSMUSG00000092341 | 6162.2 | 8059 | 11803 | 6392.3 | 4713.3 | 5744.3 | 8674.7 | 5616.6 | 0.6272 | 0.000704 | 0.019354 | *Malat1* | 19 |
| ENSMUSG00000078653 | 8.3968 | 4.9442 | 5.345 | 28.569 | 31.225 | 21.902 | 6.2287 | 27.232 | -2.1319 | 0.000705 | 0.019354 | *Cntd1* | 11 |
| ENSMUSG00000018209 | 1392.8 | 1485.2 | 1337.3 | 1949.8 | 2113.8 | 1646.6 | 1405.1 | 1903.4 | -0.4381 | 0.000706 | 0.019354 | *Stk4* | 2 |
| ENSMUSG00000094685 | 1.0496 | 1.9777 | 3.207 | 7.1422 | 53.777 | 5.9733 | 2.0781 | 22.297 | -3.435 | 0.000719 | 0.019699 | *Gm5900* | 7 |
| ENSMUSG00000051343 | 1189.2 | 1172.8 | 1043.3 | 790.74 | 772.82 | 927.85 | 1135.1 | 830.47 | 0.4513 | 0.00072 | 0.019699 | *Rab11fip5* | 6 |
| ENSMUSG00000030491 | 22.042 | 83.062 | 43.829 | 157.13 | 321.79 | 157.3 | 49.644 | 212.07 | -2.094 | 0.000732 | 0.019947 | *Tdrd12* | 7 |
| ENSMUSG00000028683 | 426.14 | 359.94 | 420.11 | 555.05 | 666.14 | 524.65 | 402.06 | 581.95 | -0.5349 | 0.000732 | 0.019947 | *Eif2b3* | 4 |
| ENSMUSG00000060098 | 714.78 | 754.48 | 757.92 | 936.65 | 1299.3 | 944.78 | 742.39 | 1060.2 | -0.515 | 0.000734 | 0.019988 | *Prmt7* | 8 |
| ENSMUSG00000030254 | 625.56 | 618.02 | 572.98 | 775.44 | 885.58 | 819.34 | 605.52 | 826.79 | -0.4497 | 0.000745 | 0.020212 | *Rad18* | 6 |
| ENSMUSG00000008682 | 20735 | 19735 | 20326 | 15711 | 15874 | 16802 | 20265 | 16129 | 0.3294 | 0.000746 | 0.020212 | *Rpl10* | X |
| ENSMUSG00000107709 | 4.1984 | 7.9107 | 5.345 | 27.549 | 51.175 | 10.951 | 5.818 | 29.891 | -2.3619 | 0.000746 | 0.020212 | *9430018G01Rik* | 6 |
| ENSMUSG00000029014 | 931 | 1049.2 | 1252.9 | 1362.1 | 2131.1 | 1383.8 | 1077.7 | 1625.7 | -0.5938 | 0.000749 | 0.020279 | *Dnajc2* | 5 |
| ENSMUSG00000072774 | 77.671 | 48.453 | 94.071 | 80.605 | 300.11 | 147.34 | 73.398 | 176.02 | -1.2664 | 0.000751 | 0.020285 | *Zfp951* | 5 |
| ENSMUSG00000031241 | 4.1984 | 26.699 | 13.897 | 68.361 | 42.501 | 43.804 | 14.931 | 51.555 | -1.7791 | 0.000753 | 0.020305 | *Tbx22* | X |
| ENSMUSG00000037759 | 82.919 | 95.917 | 57.726 | 39.792 | 24.286 | 38.826 | 78.854 | 34.302 | 1.2081 | 0.000765 | 0.02061 | *Ptger2* | 14 |
| ENSMUSG00000047022 | 153.24 | 178.98 | 151.8 | 240.79 | 324.39 | 224 | 161.34 | 263.06 | -0.7065 | 0.000766 | 0.020614 | *Mipol1* | 12 |
| ENSMUSG00000005470 | 329.58 | 313.46 | 416.91 | 452 | 659.2 | 510.72 | 353.31 | 540.64 | -0.6155 | 0.000773 | 0.020751 | *Asf1b* | 8 |
| ENSMUSG00000007379 | 141.7 | 147.34 | 98.347 | 206.1 | 337.4 | 173.23 | 129.13 | 238.91 | -0.8891 | 0.000781 | 0.020939 | *Dennd2c* | 3 |
| ENSMUSG00000052371 | 113.36 | 97.895 | 96.209 | 53.056 | 55.511 | 50.773 | 102.49 | 53.114 | 0.9473 | 0.000789 | 0.021121 | *Hoxd3os1* | 2 |
| ENSMUSG00000050050 | 22.042 | 41.531 | 18.173 | 53.056 | 127.5 | 50.773 | 27.249 | 77.111 | -1.5008 | 0.000792 | 0.02118 | *Ccdc158* | 5 |
| ENSMUSG00000020848 | 671.75 | 907.75 | 772.88 | 486.69 | 463.17 | 641.13 | 784.13 | 530.33 | 0.5651 | 0.000796 | 0.021242 | *Doc2b* | 11 |
| ENSMUSG00000036928 | 120.7 | 272.92 | 200.97 | 524.44 | 1160.5 | 433.06 | 198.2 | 706.01 | -1.8328 | 0.000804 | 0.021437 | *Stag3* | 5 |
| ENSMUSG00000040728 | 66.125 | 214.58 | 103.69 | 366.29 | 749.4 | 383.29 | 128.13 | 499.66 | -1.9629 | 0.000808 | 0.021519 | *Esrp1* | 4 |
| ENSMUSG00000012123 | 14.694 | 20.766 | 13.897 | 39.792 | 92.808 | 27.875 | 16.452 | 53.492 | -1.7039 | 0.000827 | 0.021981 | *Crybg2* | 4 |
| ENSMUSG00000031478 | 92.365 | 77.129 | 109.04 | 138.76 | 220.31 | 149.33 | 92.844 | 169.47 | -0.8722 | 0.00083 | 0.022029 | *Nek3* | 8 |

| | | | | | | | | | | | | | |
|---|---|---|---|---|---|---|---|---|---|---|---|---|---|
| ENSMUSG00000022849 | 201.52 | 235.34 | 209.52 | 308.14 | 466.64 | 279.75 | 215.46 | 351.51 | -0.7077 | 0.000832 | 0.022051 | *Hspbap1* | 16 |
| ENSMUSG00000049608 | 47.232 | 48.453 | 43.829 | 159.17 | 100.61 | 58.737 | 46.505 | 106.17 | -1.1896 | 0.000849 | 0.022467 | *Gpr55* | 1 |
| ENSMUSG00000022557 | 749.42 | 700.1 | 837.02 | 1074.4 | 1170.1 | 922.87 | 762.18 | 1055.8 | -0.4708 | 0.000854 | 0.022571 | *Bop1* | 15 |
| ENSMUSG00000024556 | 2145.4 | 1768 | 1846.2 | 2505.9 | 2753.9 | 2392.3 | 1919.9 | 2550.7 | -0.4102 | 0.000863 | 0.022777 | *Me2* | 18 |
| ENSMUSG00000001630 | 586.73 | 634.83 | 540.91 | 810.13 | 1065.1 | 705.84 | 587.49 | 860.37 | -0.551 | 0.00087 | 0.022917 | *Stk38l* | 6 |
| ENSMUSG00000041378 | 379.96 | 370.81 | 437.22 | 244.88 | 296.64 | 262.83 | 396 | 268.11 | 0.5609 | 0.000877 | 0.023045 | *Cldn5* | 16 |
| ENSMUSG00000044863 | 822.89 | 730.75 | 605.05 | 1107 | 1124.1 | 867.12 | 719.56 | 1032.8 | -0.5214 | 0.000877 | 0.023045 | *Defb36* | 2 |
| ENSMUSG00000025420 | 30.439 | 27.687 | 23.518 | 48.975 | 75.461 | 69.688 | 27.215 | 64.708 | -1.2512 | 0.000888 | 0.023294 | *Katnal2* | 18 |
| ENSMUSG00000040721 | 556.29 | 568.58 | 434.01 | 385.68 | 342.61 | 342.47 | 519.63 | 356.92 | 0.5428 | 0.000892 | 0.02336 | *Zfhx2* | 14 |
| ENSMUSG00000038201 | 162.69 | 135.47 | 156.07 | 97.95 | 47.705 | 91.591 | 151.41 | 79.082 | 0.9424 | 0.000893 | 0.02336 | *Kcna7* | 7 |
| ENSMUSG00000040841 | 1428.5 | 1453.6 | 1496.6 | 1173.4 | 860.43 | 1137.9 | 1459.6 | 1057.2 | 0.4661 | 0.000896 | 0.023414 | *Six5* | 7 |
| ENSMUSG00000082088 | 19.942 | 20.766 | 21.38 | 44.894 | 67.654 | 47.786 | 20.696 | 53.445 | -1.372 | 0.0009 | 0.023486 | *Gm15753* | 5 |
| ENSMUSG00000023232 | 303.34 | 227.43 | 306.8 | 192.84 | 165.67 | 173.23 | 279.19 | 177.24 | 0.6557 | 0.000905 | 0.023569 | *Serinc2* | 4 |
| ENSMUSG00000049804 | 1440.1 | 1369.5 | 1477.3 | 1201.9 | 890.78 | 1039.4 | 1429 | 1044 | 0.4535 | 0.000917 | 0.02386 | *Armcx4* | X |
| ENSMUSG00000014226 | 1786.4 | 1930.2 | 2283.4 | 2371.2 | 3450.4 | 2670.1 | 2000 | 2830.6 | -0.5015 | 0.000922 | 0.023954 | *Cacybp* | 1 |
| ENSMUSG00000026496 | 2939.9 | 2955.6 | 2905.5 | 3494.6 | 4123.5 | 3732.3 | 2933.7 | 3783.4 | -0.3672 | 0.000926 | 0.024028 | *Parp1* | 1 |
| ENSMUSG00000006678 | 1088.4 | 1012.6 | 1011.3 | 1359.1 | 1478 | 1281.3 | 1037.4 | 1372.8 | -0.4045 | 0.000939 | 0.02431 | *Pola1* | X |
| ENSMUSG00000095440 | 298.09 | 345.1 | 260.83 | 221.41 | 146.58 | 196.12 | 301.34 | 188.04 | 0.6836 | 0.000944 | 0.024405 | *Fignl2* | 15 |
| ENSMUSG00000022471 | 977.18 | 861.28 | 969.58 | 1139.7 | 1642.8 | 1208.6 | 936.01 | 1330.4 | -0.5081 | 0.00095 | 0.024525 | *Xrcc6* | 15 |
| ENSMUSG00000020387 | 173.18 | 319.39 | 196.69 | 350.99 | 535.16 | 318.58 | 229.76 | 401.58 | -0.8056 | 0.000951 | 0.024529 | *Jade2* | 11 |
| ENSMUSG00000050105 | 82.919 | 77.129 | 96.209 | 42.853 | 32.96 | 47.786 | 85.419 | 41.2 | 1.0552 | 0.000955 | 0.024582 | *Grrp1* | 4 |
| ENSMUSG00000033788 | 364.21 | 266 | 362.39 | 242.84 | 167.4 | 213.05 | 330.87 | 207.76 | 0.6728 | 0.000962 | 0.024738 | *Dysf* | 6 |
| ENSMUSG00000022577 | 301.24 | 170.08 | 265.11 | 121.42 | 134.44 | 168.25 | 245.48 | 141.37 | 0.7954 | 0.000965 | 0.02477 | *Ly6h* | 15 |
| ENSMUSG00000042724 | 12.595 | 39.553 | 25.656 | 53.056 | 156.99 | 37.831 | 25.935 | 82.627 | -1.6729 | 0.000967 | 0.024789 | *Map3k9* | 12 |
| ENSMUSG00000002985 | 5120 | 4765.2 | 4149.8 | 3800.7 | 2617.7 | 3617.8 | 4678.3 | 3345.4 | 0.4841 | 0.000972 | 0.024876 | *Apoe* | 7 |
| ENSMUSG00000042708 | 22.042 | 53.397 | 38.484 | 58.158 | 143.98 | 81.635 | 37.974 | 94.592 | -1.3178 | 0.000973 | 0.024887 | *Shcbp1l* | 1 |
| ENSMUSG00000034463 | 326.43 | 332.25 | 279.01 | 490.77 | 491.8 | 392.25 | 312.56 | 458.27 | -0.552 | 0.000982 | 0.025027 | *Scara3* | 14 |
| ENSMUSG00000024044 | 537.4 | 934.45 | 507.77 | 945.83 | 1249.9 | 960.71 | 659.87 | 1052.1 | -0.6728 | 0.000984 | 0.025027 | *Epb41l3* | 17 |
| ENSMUSG00000034591 | 59.827 | 35.598 | 66.278 | 92.849 | 156.13 | 92.586 | 53.901 | 113.85 | -1.0845 | 0.000984 | 0.025027 | *Slc41a2* | 10 |
| ENSMUSG00000038119 | 2785.6 | 2257.5 | 2099.5 | 1531.5 | 1469.3 | 2035.9 | 2380.9 | 1678.9 | 0.5042 | 0.000985 | 0.025027 | *Cdon* | 9 |
| ENSMUSG00000020717 | 843.88 | 752.5 | 828.47 | 648.92 | 559.45 | 564.48 | 808.29 | 590.95 | 0.4522 | 0.000986 | 0.025027 | *Pecam1* | 11 |
| ENSMUSG00000032323 | 4817.7 | 3098 | 3650.6 | 2086.5 | 2745.2 | 3044.4 | 3855.4 | 2625.4 | 0.5542 | 0.000989 | 0.025073 | *Cyp11a1* | 9 |
| ENSMUSG00000005087 | 389.4 | 388.61 | 623.22 | 329.56 | 245.46 | 300.66 | 467.08 | 291.89 | 0.6789 | 0.000999 | 0.025299 | *Cd44* | 2 |
| ENSMUSG00000040136 | 20.992 | 23.732 | 27.794 | 57.138 | 130.97 | 28.871 | 24.173 | 72.327 | -1.5863 | 0.001003 | 0.025371 | *Abcc8* | 7 |
| ENSMUSG00000063810 | 504.86 | 603.19 | 404.08 | 766.26 | 809.25 | 646.11 | 504.04 | 740.54 | -0.5547 | 0.001008 | 0.025446 | *Alms1* | 6 |
| ENSMUSG00000039183 | 600.37 | 456.84 | 540.91 | 722.38 | 866.5 | 690.91 | 532.71 | 759.93 | -0.5137 | 0.001013 | 0.02554 | *Nubp2* | 17 |
| ENSMUSG00000031962 | 20.992 | 31.643 | 36.346 | 43.874 | 120.56 | 65.706 | 29.66 | 76.715 | -1.3756 | 0.001018 | 0.025608 | *Cdh15* | 8 |
| ENSMUSG00000021095 | 24.141 | 44.498 | 45.967 | 7.1422 | 3.4695 | 18.915 | 38.202 | 9.8424 | 1.97 | 0.001019 | 0.025608 | *Gsc* | 12 |

| | | | | | | | | | | | | | |
|---|---|---|---|---|---|---|---|---|---|---|---|---|---|
| ENSMUSG00000042700 | 653.9 | 701.08 | 500.29 | 894.82 | 1120.6 | 747.66 | 618.43 | 921.04 | -0.5749 | 0.001021 | 0.025636 | *Sipa1l1* | 12 |
| ENSMUSG00000049103 | 55.629 | 59.33 | 62.002 | 29.589 | 23.419 | 21.902 | 58.987 | 24.97 | 1.2423 | 0.001024 | 0.025668 | *Ccr2* | 9 |
| ENSMUSG00000029777 | 1532.4 | 1498.1 | 1959.5 | 2067.2 | 2497.1 | 2261.9 | 1663.3 | 2275.4 | -0.4525 | 0.001025 | 0.025668 | *Gars* | 6 |
| ENSMUSG00000037151 | 152.19 | 118.66 | 165.69 | 218.35 | 220.31 | 266.81 | 145.52 | 235.16 | -0.6934 | 0.001049 | 0.02623 | *Lrrc20* | 10 |
| ENSMUSG00000109176 | 10.496 | 2.9665 | 7.483 | 25.508 | 30.358 | 29.866 | 6.9818 | 28.577 | -2.0416 | 0.001051 | 0.026231 | *Zfp264* | 7 |
| ENSMUSG00000024151 | 1288.9 | 1498.1 | 1487 | 1637.6 | 2154.5 | 2029.9 | 1424.7 | 1940.7 | -0.4462 | 0.001053 | 0.026267 | *Msh2* | 17 |
| ENSMUSG00000022177 | 625.56 | 549.79 | 668.12 | 970.32 | 873.44 | 742.68 | 614.49 | 862.15 | -0.4889 | 0.00106 | 0.026405 | *Haus4* | 14 |
| ENSMUSG00000031976 | 327.48 | 437.07 | 268.32 | 571.38 | 578.53 | 447 | 344.29 | 532.3 | -0.6279 | 0.00107 | 0.026617 | *Urb2* | 8 |
| ENSMUSG00000015880 | 1046.5 | 927.53 | 999.51 | 1165.2 | 1596.8 | 1338 | 991.16 | 1366.7 | -0.4642 | 0.001075 | 0.026689 | *Ncapg* | 5 |
| ENSMUSG00000079350 | 24.141 | 58.341 | 29.932 | 93.869 | 111.89 | 61.724 | 37.471 | 89.161 | -1.2481 | 0.001077 | 0.026701 | *Magea8* | X |
| ENSMUSG00000091475 | 428.24 | 398.5 | 459.67 | 311.2 | 268.02 | 313.6 | 428.8 | 297.6 | 0.5278 | 0.00108 | 0.026753 | *2810468N07Rik* | 17 |
| ENSMUSG00000022938 | 2.0992 | 10.877 | 10.69 | 30.609 | 56.379 | 16.924 | 7.8888 | 34.638 | -2.1362 | 0.001089 | 0.026929 | *Fam3b* | 16 |
| ENSMUSG00000061082 | 141.7 | 177 | 145.38 | 271.4 | 206.43 | 264.82 | 154.69 | 247.55 | -0.6758 | 0.001094 | 0.027013 | *Plac1* | X |
| ENSMUSG00000094497 | 27.29 | 14.833 | 13.897 | 27.549 | 167.4 | 23.893 | 18.673 | 72.948 | -1.9715 | 0.001108 | 0.027317 | *Gm8210* | 1 |
| ENSMUSG00000027102 | 1401.2 | 1425.9 | 1659.1 | 1174.4 | 1105 | 1112 | 1495.4 | 1130.5 | 0.4036 | 0.001112 | 0.027382 | *Hoxd8* | 2 |
| ENSMUSG00000028132 | 12.595 | 18.788 | 10.69 | 27.549 | 70.257 | 36.835 | 14.024 | 44.88 | -1.6795 | 0.00112 | 0.027532 | *Tmem56* | 3 |
| ENSMUSG00000082450 | 257.15 | 310.49 | 267.25 | 413.23 | 486.59 | 350.43 | 278.3 | 416.75 | -0.5831 | 0.001121 | 0.027532 | *Mageb7-ps* | X |
| ENSMUSG00000037426 | 514.31 | 559.68 | 475.7 | 801.97 | 761.55 | 619.23 | 516.56 | 727.58 | -0.4939 | 0.001122 | 0.027532 | *Depdc5* | 5 |
| ENSMUSG00000059775 | 133.3 | 180.96 | 316.42 | 106.11 | 71.991 | 137.39 | 210.23 | 105.16 | 1.0013 | 0.001146 | 0.028082 | *Rps26-ps1* | 8 |
| ENSMUSG00000037447 | 136.45 | 130.53 | 116.52 | 214.27 | 257.61 | 166.26 | 127.83 | 212.71 | -0.7361 | 0.001151 | 0.028157 | *Arid5a* | 1 |
| ENSMUSG00000028647 | 479.67 | 550.78 | 520.6 | 646.88 | 958.44 | 669.01 | 517.02 | 758.11 | -0.5532 | 0.001153 | 0.028158 | *Mycbp* | 4 |
| ENSMUSG00000006527 | 568.89 | 586.38 | 605.05 | 812.17 | 967.11 | 694.89 | 586.77 | 824.73 | -0.492 | 0.00116 | 0.028296 | *Sfmbt1* | 14 |
| ENSMUSG00000032999 | 39.885 | 58.341 | 14.966 | 108.15 | 294.04 | 126.43 | 37.731 | 176.21 | -2.2226 | 0.001168 | 0.02846 | *Nlrp4f* | 13 |
| ENSMUSG00000051900 | 0 | 4.9442 | 7.483 | 14.284 | 39.031 | 20.907 | 4.1424 | 24.741 | -2.5837 | 0.001175 | 0.02859 | *Abca16* | 7 |
| ENSMUSG00000003038 | 6970.4 | 5743.2 | 8031.3 | 4418 | 6027.3 | 4125.6 | 6915 | 4856.9 | 0.5094 | 0.001183 | 0.028744 | *Hmgn2* | 4 |
| ENSMUSG00000040732 | 80.82 | 111.74 | 126.14 | 167.33 | 344.34 | 131.41 | 106.23 | 214.36 | -1.0156 | 0.001192 | 0.028913 | *Erg* | 16 |
| ENSMUSG00000041488 | 154.29 | 147.34 | 205.25 | 254.06 | 376.44 | 222.01 | 168.96 | 284.17 | -0.7529 | 0.001199 | 0.029047 | *Stx3* | 19 |
| ENSMUSG00000028519 | 13.645 | 33.62 | 42.76 | 64.28 | 90.206 | 67.697 | 30.008 | 74.061 | -1.3054 | 0.001208 | 0.029243 | *Dab1* | 4 |
| ENSMUSG00000041431 | 2292.3 | 2051.8 | 2318.6 | 2648.7 | 3591.8 | 2794.5 | 2220.9 | 3011.7 | -0.4398 | 0.001215 | 0.029332 | *Ccnb1* | 13 |
| ENSMUSG00000006587 | 7.3472 | 27.687 | 12.828 | 34.691 | 105.82 | 29.866 | 15.954 | 56.792 | -1.8309 | 0.001215 | 0.029332 | *Snai3* | 8 |
| ENSMUSG00000032089 | 64.026 | 97.895 | 49.174 | 146.93 | 141.38 | 116.48 | 70.365 | 134.93 | -0.9364 | 0.001221 | 0.029414 | *Il10ra* | 9 |
| ENSMUSG00000097258 | 26.24 | 25.71 | 19.242 | 66.321 | 78.063 | 36.835 | 23.731 | 60.406 | -1.3492 | 0.001222 | 0.029414 | *Gm26767* | 9 |
| ENSMUSG00000021311 | 1450.6 | 1606.9 | 1377.9 | 2064.1 | 1990.6 | 1780 | 1478.4 | 1944.9 | -0.3955 | 0.001224 | 0.029414 | *Mtr* | 13 |
| ENSMUSG00000017733 | 95.514 | 53.397 | 59.864 | 147.95 | 114.49 | 133.4 | 69.592 | 131.95 | -0.9229 | 0.001231 | 0.029561 | *Eppin* | 2 |
| ENSMUSG00000097124 | 514.31 | 529.03 | 385.91 | 361.19 | 302.71 | 303.64 | 476.41 | 322.51 | 0.5641 | 0.001235 | 0.029607 | *A530020G20Rik* | 3 |
| ENSMUSG00000027353 | 417.74 | 389.6 | 467.15 | 538.73 | 659.2 | 594.34 | 424.83 | 597.42 | -0.4931 | 0.00125 | 0.029919 | *Mcm8* | 2 |
| ENSMUSG00000007833 | 561.54 | 514.19 | 521.67 | 687.69 | 735.53 | 733.72 | 532.47 | 718.98 | -0.4336 | 0.001253 | 0.029919 | *Aldh16a1* | 7 |
| ENSMUSG00000030528 | 638.16 | 581.43 | 555.88 | 754.01 | 910.73 | 771.55 | 591.82 | 812.1 | -0.4572 | 0.001255 | 0.029919 | *Blm* | 7 |

| | | | | | | | | | | | | | |
|---|---|---|---|---|---|---|---|---|---|---|---|---|---|
| ENSMUSG00000078867 | 9.4464 | 2.9665 | 3.207 | 21.427 | 39.899 | 15.929 | 5.2066 | 25.751 | -2.315 | 0.001256 | 0.029919 | *Gm14418* | 2 |
| ENSMUSG00000022191 | 979.28 | 1176.7 | 1142.8 | 1324.4 | 1738.2 | 1451.5 | 1099.6 | 1504.7 | -0.4529 | 0.001257 | 0.029919 | *Drosha* | 15 |
| ENSMUSG00000020929 | 1571.3 | 1755.2 | 1816.2 | 2120.2 | 2549.2 | 2135.5 | 1714.2 | 2268.3 | -0.4044 | 0.001261 | 0.029919 | *Eftud2* | 11 |
| ENSMUSG00000040938 | 28.339 | 63.285 | 33.139 | 62.239 | 155.26 | 84.622 | 41.588 | 100.71 | -1.2761 | 0.001261 | 0.029919 | *Slc16a11* | 11 |
| ENSMUSG00000025955 | 12056 | 12406 | 10908 | 7854.4 | 10172 | 8919.1 | 11790 | 8981.7 | 0.3924 | 0.001261 | 0.029919 | *Akr1cl* | 1 |
| ENSMUSG00000028630 | 733.67 | 803.92 | 793.19 | 561.17 | 502.2 | 633.17 | 776.93 | 565.52 | 0.4591 | 0.001266 | 0.029994 | *Dyrk2* | 10 |
| ENSMUSG00000030144 | 66.125 | 79.107 | 39.553 | 25.508 | 20.817 | 28.871 | 61.595 | 25.065 | 1.3023 | 0.00127 | 0.030039 | *Clec4d* | 6 |
| ENSMUSG00000002797 | 140.65 | 202.71 | 159.28 | 220.39 | 325.26 | 270.79 | 167.55 | 272.15 | -0.7003 | 0.001275 | 0.030122 | *Ggct* | 6 |
| ENSMUSG00000022286 | 25.191 | 39.553 | 28.863 | 53.056 | 126.64 | 54.755 | 31.202 | 78.149 | -1.3273 | 0.001279 | 0.030195 | *Grhl2* | 15 |
| ENSMUSG00000031548 | 4620.4 | 4982.7 | 4605.2 | 3377.2 | 3575.3 | 4062.8 | 4736.1 | 3671.8 | 0.3673 | 0.001282 | 0.030214 | *Sfrp1* | 8 |
| ENSMUSG00000053332 | 3479.4 | 3481.7 | 3675.2 | 2654.9 | 3045.3 | 2522.7 | 3545.4 | 2741 | 0.371 | 0.00129 | 0.030323 | *Gas5* | 1 |
| ENSMUSG00000045216 | 2054.1 | 2267.4 | 1917.8 | 1564.1 | 1073.8 | 1721.3 | 2079.7 | 1453.1 | 0.518 | 0.001291 | 0.030323 | *Hs6st1* | 1 |
| ENSMUSG00000055723 | 1614.3 | 1744.3 | 1721.1 | 2203.9 | 2449.4 | 1999.1 | 1693.2 | 2217.5 | -0.3894 | 0.001292 | 0.030323 | *Rras2* | 7 |
| ENSMUSG00000110344 | 59.827 | 83.062 | 117.59 | 136.72 | 207.3 | 145.35 | 86.826 | 163.12 | -0.9127 | 0.001294 | 0.030341 | *Gm45716* | 11 |
| ENSMUSG00000026229 | 2382.6 | 2319.8 | 2267.3 | 2733.4 | 3244.8 | 2999.6 | 2323.2 | 2992.6 | -0.3655 | 0.001304 | 0.030538 | *Psmd1* | 1 |
| ENSMUSG00000031125 | 33.587 | 24.721 | 12.828 | 49.995 | 71.124 | 59.733 | 23.712 | 60.284 | -1.3465 | 0.001316 | 0.030775 | *3830403N18Rik* | X |
| ENSMUSG00000035517 | 389.4 | 391.58 | 373.08 | 570.36 | 654.86 | 449.99 | 384.69 | 558.4 | -0.5385 | 0.001319 | 0.030789 | *Tdrd7* | 4 |
| ENSMUSG00000039759 | 366.31 | 368.84 | 404.08 | 266.3 | 259.34 | 270.79 | 379.74 | 265.48 | 0.5165 | 0.00132 | 0.030789 | *Thap3* | 4 |
| ENSMUSG00000019558 | 1277.4 | 1460.5 | 1374.7 | 1054 | 892.52 | 1118 | 1370.9 | 1021.5 | 0.4251 | 0.001331 | 0.03101 | *Slc6a8* | X |
| ENSMUSG00000073968 | 318.03 | 399.49 | 349.56 | 611.17 | 468.38 | 476.87 | 355.69 | 518.8 | -0.5433 | 0.001343 | 0.031235 | *Trim68* | 7 |
| ENSMUSG00000009628 | 429.29 | 744.59 | 331.39 | 1049.9 | 2311.5 | 1170.8 | 501.76 | 1510.7 | -1.5901 | 0.001346 | 0.031278 | *Tex15* | 8 |
| ENSMUSG00000061119 | 715.83 | 691.2 | 607.19 | 427.51 | 463.17 | 546.56 | 671.4 | 479.08 | 0.4872 | 0.001354 | 0.03141 | *Prcp* | 7 |
| ENSMUSG00000074793 | 564.69 | 593.3 | 748.3 | 485.67 | 361.69 | 461.93 | 635.43 | 436.43 | 0.543 | 0.001359 | 0.031507 | *Hspa12b* | 2 |
| ENSMUSG00000032741 | 2075.1 | 2419.7 | 1794.8 | 1622.3 | 1210.8 | 1653.6 | 2096.5 | 1495.6 | 0.488 | 0.001365 | 0.031598 | *Tpcn1* | 5 |
| ENSMUSG00000030986 | 976.13 | 1019.5 | 829.54 | 1354 | 1241.2 | 1191.7 | 941.72 | 1262.3 | -0.4222 | 0.001369 | 0.031621 | *Dhx32* | 7 |
| ENSMUSG00000022951 | 1792.7 | 1519.8 | 1907.1 | 1331.5 | 1228.2 | 1367.9 | 1739.9 | 1309.2 | 0.4103 | 0.00137 | 0.031621 | *Rcan1* | 16 |
| ENSMUSG00000056749 | 311.73 | 558.69 | 442.56 | 281.61 | 246.33 | 303.64 | 437.66 | 277.19 | 0.6604 | 0.001374 | 0.031686 | *Nfil3* | 13 |
| ENSMUSG00000039410 | 55.629 | 73.174 | 54.519 | 108.15 | 167.4 | 87.608 | 61.107 | 121.05 | -0.9879 | 0.001376 | 0.031689 | *Prdm16* | 4 |
| ENSMUSG00000020712 | 33.587 | 24.721 | 25.656 | 67.341 | 84.134 | 45.795 | 27.988 | 65.757 | -1.2358 | 0.001381 | 0.031756 | *Tcam1* | 11 |
| ENSMUSG00000078864 | 15.744 | 8.8995 | 9.6209 | 34.691 | 39.899 | 32.853 | 11.422 | 35.814 | -1.6521 | 0.001395 | 0.032047 | *Gm14322* | 2 |
| ENSMUSG00000057156 | 291.79 | 315.44 | 249.08 | 444.86 | 421.54 | 377.31 | 285.43 | 414.57 | -0.5378 | 0.001397 | 0.032058 | *Homez* | 14 |
| ENSMUSG00000042660 | 478.62 | 370.81 | 398.73 | 590.76 | 706.03 | 514.7 | 416.06 | 603.83 | -0.5386 | 0.001409 | 0.032261 | *Wdr55* | 18 |
| ENSMUSG00000031093 | 496.46 | 567.59 | 616.81 | 741.77 | 875.17 | 716.8 | 560.29 | 777.91 | -0.4742 | 0.00141 | 0.032261 | *Dock11* | X |
| ENSMUSG00000042997 | 319.08 | 318.4 | 272.59 | 502 | 392.92 | 426.1 | 303.36 | 440.34 | -0.5362 | 0.001412 | 0.032261 | *Nhlrc3* | 3 |
| ENSMUSG00000032175 | 369.46 | 402.46 | 341.01 | 533.63 | 528.23 | 490.81 | 370.98 | 517.55 | -0.48 | 0.001414 | 0.032271 | *Tyk2* | 9 |
| ENSMUSG00000023403 | 180.53 | 356.97 | 230.9 | 504.04 | 1589.9 | 656.07 | 256.13 | 916.66 | -1.8395 | 0.001421 | 0.032397 | *Stk31* | 6 |
| ENSMUSG00000034825 | 6.2976 | 27.687 | 3.207 | 30.609 | 68.522 | 44.8 | 12.397 | 47.977 | -1.9435 | 0.001432 | 0.032602 | *Nrip3* | 7 |
| ENSMUSG00000062760 | 71.373 | 32.632 | 44.898 | 12.244 | 16.48 | 23.893 | 49.634 | 17.539 | 1.4996 | 0.001441 | 0.032739 | *Shisal1* | 15 |

| | | | | | | | | | | | | |
|---|---|---|---|---|---|---|---|---|---|---|---|---|
| ENSMUSG00000073888 | 250.86 | 205.68 | 207.38 | 133.66 | 143.12 | 147.34 | 221.31 | 141.37 | 0.646 | 0.001443 | 0.032739 | *Ccl27a* | 4 |
| ENSMUSG00000043419 | 249.81 | 314.45 | 345.28 | 365.27 | 640.98 | 439.04 | 303.18 | 481.76 | -0.6697 | 0.001444 | 0.032739 | *Rnf227* | 11 |
| ENSMUSG00000040666 | 318.03 | 306.54 | 302.53 | 157.13 | 235.06 | 211.06 | 309.03 | 201.08 | 0.6176 | 0.001445 | 0.032739 | *Sh3bgr* | 16 |
| ENSMUSG00000033389 | 158.49 | 201.72 | 210.59 | 262.22 | 333.07 | 282.74 | 190.27 | 292.68 | -0.6224 | 0.001467 | 0.033181 | *Arhgap44* | 11 |
| ENSMUSG00000037025 | 14.694 | 18.788 | 9.6209 | 2.0406 | 1.7347 | 0.9955 | 14.368 | 1.5903 | 3.1756 | 0.001502 | 0.033928 | *Foxa2* | 2 |
| ENSMUSG00000079349 | 14.694 | 36.587 | 20.311 | 49.995 | 98.88 | 44.8 | 23.864 | 64.558 | -1.4355 | 0.001503 | 0.033928 | *Magea5* | X |
| ENSMUSG00000040325 | 769.36 | 813.81 | 669.19 | 1126.4 | 1125 | 869.11 | 750.79 | 1040.2 | -0.4703 | 0.001512 | 0.034069 | *Dcaf1* | 9 |
| ENSMUSG00000025328 | 29.389 | 27.687 | 19.242 | 49.995 | 100.61 | 43.804 | 25.439 | 64.805 | -1.3525 | 0.001518 | 0.034165 | *Padi3* | 4 |
| ENSMUSG00000033581 | 669.65 | 731.74 | 813.5 | 491.79 | 537.77 | 582.4 | 738.3 | 537.32 | 0.4582 | 0.00152 | 0.034165 | *Igf2bp2* | 16 |
| ENSMUSG00000028121 | 709.53 | 744.59 | 690.57 | 562.19 | 502.2 | 529.63 | 714.9 | 531.34 | 0.4288 | 0.001521 | 0.034165 | *Bcar3* | 3 |
| ENSMUSG00000086077 | 113.36 | 108.77 | 63.071 | 49.995 | 49.44 | 39.822 | 95.067 | 46.419 | 1.0345 | 0.001532 | 0.034351 | *Gm14396* | 2 |
| ENSMUSG00000029838 | 4765.2 | 2961.6 | 4748.5 | 2923.2 | 2882.3 | 2985.7 | 4158.4 | 2930.4 | 0.5048 | 0.001535 | 0.03438 | *Ptn* | 6 |
| ENSMUSG00000022672 | 635.01 | 496.4 | 417.98 | 823.39 | 882.98 | 609.28 | 516.46 | 771.88 | -0.5801 | 0.001538 | 0.034383 | *Prkdc* | 16 |
| ENSMUSG00000083114 | 0 | 2.9665 | 2.138 | 10.203 | 30.358 | 8.9599 | 1.7015 | 16.507 | -3.274 | 0.001539 | 0.034383 | *Gm14790* | X |
| ENSMUSG00000031314 | 1359.2 | 1351.7 | 1268.9 | 1761.1 | 2252.5 | 1502.3 | 1326.6 | 1838.6 | -0.4713 | 0.001546 | 0.034475 | *Taf1* | X |
| ENSMUSG00000037275 | 732.62 | 728.77 | 649.95 | 1005 | 1154.5 | 800.42 | 703.78 | 986.63 | -0.4879 | 0.001547 | 0.034475 | *Gemin5* | 11 |
| ENSMUSG00000021392 | 710.58 | 881.05 | 683.09 | 902.98 | 1457.2 | 1002.5 | 758.24 | 1120.9 | -0.5644 | 0.001567 | 0.034896 | *Nol8* | 13 |
| ENSMUSG00000050071 | 251.91 | 240.29 | 237.32 | 380.58 | 326.13 | 347.45 | 243.17 | 351.38 | -0.5303 | 0.001573 | 0.034977 | *Bex1* | X |
| ENSMUSG00000062353 | 643.41 | 727.78 | 791.06 | 6539.2 | 721.65 | 6471.1 | 720.75 | 4577.3 | -2.6669 | 0.001592 | 0.035359 | *Gm15772* | 5 |
| ENSMUSG00000029591 | 870.12 | 977.96 | 1008.1 | 1148.9 | 1340.1 | 1297.2 | 952.05 | 1262.1 | -0.407 | 0.001598 | 0.035434 | *Ung* | 5 |
| ENSMUSG00000020032 | 911.06 | 830.62 | 901.16 | 600.97 | 595.01 | 743.68 | 880.95 | 646.55 | 0.4466 | 0.0016 | 0.035438 | *Nuak1* | 10 |
| ENSMUSG00000018451 | 925.75 | 744.59 | 980.27 | 669.33 | 608.02 | 652.08 | 883.54 | 643.14 | 0.4581 | 0.001609 | 0.035609 | *6330403K07Rik* | 11 |
| ENSMUSG00000028826 | 811.34 | 1056.1 | 931.09 | 1190.7 | 1698.3 | 1137.9 | 932.84 | 1342.3 | -0.5255 | 0.001623 | 0.035857 | *Maco1* | 4 |
| ENSMUSG00000005667 | 614.02 | 457.83 | 689.5 | 819.31 | 916.8 | 779.52 | 587.12 | 838.54 | -0.5153 | 0.001643 | 0.036231 | *Mthfd2* | 6 |
| ENSMUSG00000026414 | 109.16 | 117.67 | 109.04 | 184.68 | 194.29 | 161.28 | 111.96 | 180.08 | -0.6861 | 0.001643 | 0.036231 | *Tnnt2* | 1 |
| ENSMUSG00000020490 | 1.0496 | 9.8884 | 2.138 | 46.935 | 15.613 | 14.933 | 4.3586 | 25.827 | -2.5467 | 0.001647 | 0.036264 | *Btnl10* | 11 |
| ENSMUSG00000078137 | 120.7 | 126.57 | 129.35 | 58.158 | 63.318 | 88.604 | 125.54 | 70.027 | 0.8432 | 0.001665 | 0.036617 | *Ankrd63* | 2 |
| ENSMUSG00000028138 | 4898.5 | 4786 | 5070.2 | 3998.6 | 3768.7 | 4034 | 4918.2 | 3933.8 | 0.3223 | 0.00167 | 0.036672 | *Adh5* | 3 |
| ENSMUSG00000016496 | 27.29 | 29.665 | 19.242 | 52.036 | 111.02 | 37.831 | 25.399 | 66.963 | -1.4018 | 0.001679 | 0.036824 | *Cd274* | 19 |
| ENSMUSG00000031442 | 535.3 | 547.81 | 508.84 | 414.25 | 369.5 | 374.33 | 530.65 | 386.02 | 0.4598 | 0.001687 | 0.036968 | *Mcf2l* | 8 |
| ENSMUSG00000115536 | 3.1488 | 1.9777 | 6.414 | 17.345 | 26.888 | 16.924 | 3.8468 | 20.386 | -2.4191 | 0.001711 | 0.037406 | *Gm32857* | 14 |
| ENSMUSG00000069053 | 325.38 | 424.21 | 243.73 | 784.62 | 1462.4 | 554.52 | 331.11 | 933.84 | -1.4959 | 0.001712 | 0.037406 | *Uba1y* | Y |
| ENSMUSG00000043439 | 33.587 | 74.163 | 75.899 | 119.38 | 147.45 | 101.55 | 61.216 | 122.79 | -1.0049 | 0.001713 | 0.037406 | *Epop* | 11 |
| ENSMUSG00000079184 | 926.8 | 952.25 | 957.82 | 1145.8 | 1693.1 | 1158.8 | 945.62 | 1332.6 | -0.4957 | 0.001721 | 0.037537 | *Mphosph8* | 14 |
| ENSMUSG00000021703 | 6229.4 | 6733 | 5398.4 | 5124 | 4355 | 4629.3 | 6120.3 | 4702.8 | 0.3803 | 0.001751 | 0.038105 | *Serinc5* | 13 |
| ENSMUSG00000109017 | 1.0496 | 2.9665 | 4.276 | 14.284 | 22.551 | 15.929 | 2.764 | 17.588 | -2.6743 | 0.001752 | 0.038105 | *Gm38979* | 7 |
| ENSMUSG00000043811 | 77.671 | 107.78 | 99.416 | 48.975 | 26.888 | 60.729 | 94.957 | 45.531 | 1.0675 | 0.001755 | 0.038133 | *Rtn4r* | 16 |
| ENSMUSG00000040964 | 529 | 487.5 | 525.94 | 386.7 | 344.34 | 387.27 | 514.15 | 372.77 | 0.4645 | 0.001779 | 0.038616 | *Arhgef10l* | 4 |

| | | | | | | | | | | | | | |
|---|---|---|---|---|---|---|---|---|---|---|---|---|---|
| ENSMUSG00000021494 | 746.27 | 617.03 | 640.33 | 874.41 | 1160.5 | 804.4 | 667.88 | 946.45 | -0.5039 | 0.001797 | 0.038909 | *Ddx41* | 13 |
| ENSMUSG00000032076 | 1064.3 | 1393.3 | 946.06 | 1678.4 | 1491 | 1541.1 | 1134.5 | 1570.2 | -0.4682 | 0.001798 | 0.038909 | *Cadm1* | 9 |
| ENSMUSG00000064345 | 27210 | 24592 | 21378 | 29717 | 35781 | 30250 | 24393 | 31916 | -0.3878 | 0.001799 | 0.038909 | *mt-Nd2* | MT |
| ENSMUSG00000041134 | 57.728 | 52.408 | 60.933 | 23.467 | 25.154 | 26.88 | 57.023 | 25.167 | 1.1792 | 0.001811 | 0.039106 | *Cyyr1* | 16 |
| ENSMUSG00000056305 | 820.79 | 808.87 | 912.92 | 1058.1 | 1236.9 | 1076.2 | 847.53 | 1123.7 | -0.4076 | 0.001816 | 0.039142 | *Usp39* | 6 |
| ENSMUSG00000034239 | 27.29 | 41.531 | 17.104 | 62.239 | 109.29 | 45.795 | 28.642 | 72.441 | -1.3384 | 0.001817 | 0.039142 | *Gm884* | 11 |
| ENSMUSG00000063060 | 103.91 | 83.062 | 135.76 | 64.28 | 49.44 | 55.751 | 107.58 | 56.49 | 0.9304 | 0.001819 | 0.039142 | *Sox7* | 14 |
| ENSMUSG00000073889 | 1382.3 | 1371.5 | 1445.3 | 1013.2 | 1195.2 | 984.6 | 1399.7 | 1064.3 | 0.3945 | 0.001839 | 0.039516 | *Il11ra1* | 4 |
| ENSMUSG00000001829 | 705.33 | 702.07 | 647.81 | 934.61 | 991.4 | 822.32 | 685.07 | 916.11 | -0.4195 | 0.001841 | 0.039516 | *Clpb* | 7 |
| ENSMUSG00000037892 | 1196.5 | 962.14 | 1077.5 | 870.33 | 782.36 | 760.6 | 1078.7 | 804.43 | 0.4233 | 0.001846 | 0.039585 | *Pcdh18* | 3 |
| ENSMUSG00000022528 | 738.92 | 651.64 | 742.95 | 567.3 | 481.39 | 522.66 | 711.17 | 523.78 | 0.4417 | 0.001852 | 0.03966 | *Hes1* | 16 |
| ENSMUSG00000024269 | 389.4 | 299.62 | 399.8 | 502 | 699.1 | 445.01 | 362.94 | 548.7 | -0.5982 | 0.001856 | 0.03966 | *Tpgs2* | 18 |
| ENSMUSG00000083977 | 2.0992 | 0.9888 | 3.207 | 9.1828 | 32.96 | 8.9599 | 2.0983 | 17.034 | -3.0399 | 0.001856 | 0.03966 | *Mageb8-ps* | X |
| ENSMUSG00000025902 | 224.62 | 147.34 | 228.76 | 111.21 | 135.31 | 113.49 | 200.24 | 120.01 | 0.7357 | 0.001862 | 0.039691 | *Sox17* | 1 |
| ENSMUSG00000067764 | 7.3472 | 45.486 | 23.518 | 94.889 | 318.32 | 96.568 | 25.45 | 169.93 | -2.7382 | 0.001862 | 0.039691 | *Xlr5c* | X |
| ENSMUSG00000038187 | 498.56 | 590.33 | 549.46 | 607.09 | 974.05 | 798.43 | 546.12 | 793.19 | -0.5393 | 0.001874 | 0.039894 | *Btbd10* | 7 |
| ENSMUSG00000021418 | 329.58 | 344.11 | 319.63 | 394.86 | 657.46 | 449.99 | 331.11 | 500.77 | -0.5984 | 0.001887 | 0.040088 | *Rpp40* | 13 |
| ENSMUSG00000038112 | 772.51 | 628.9 | 767.54 | 504.04 | 418.94 | 599.32 | 722.98 | 507.43 | 0.5115 | 0.001888 | 0.040088 | *AW551984* | 9 |
| ENSMUSG00000017561 | 735.77 | 731.74 | 699.12 | 852.98 | 1251.6 | 929.84 | 722.21 | 1011.5 | -0.4869 | 0.001891 | 0.040088 | *Crlf3* | 11 |
| ENSMUSG00000040363 | 721.08 | 800.96 | 847.71 | 605.05 | 592.41 | 575.43 | 789.92 | 590.96 | 0.4186 | 0.001892 | 0.040088 | *Bcor* | X |
| ENSMUSG00000022994 | 818.69 | 974 | 911.85 | 627.49 | 525.62 | 769.56 | 901.52 | 640.89 | 0.4933 | 0.001901 | 0.040232 | *Adcy6* | 15 |
| ENSMUSG00000021718 | 3.1488 | 14.833 | 10.69 | 23.467 | 48.572 | 31.858 | 9.5571 | 34.632 | -1.856 | 0.001906 | 0.04029 | *4933425L06Rik* | 13 |
| ENSMUSG00000000276 | 235.11 | 287.75 | 217.01 | 365.27 | 505.67 | 294.68 | 246.62 | 388.54 | -0.6566 | 0.001924 | 0.040632 | *Dgke* | 11 |
| ENSMUSG00000045092 | 1010.8 | 962.14 | 863.75 | 794.83 | 529.96 | 687.92 | 945.55 | 670.9 | 0.4963 | 0.001926 | 0.040636 | *S1pr1* | 3 |
| ENSMUSG00000038644 | 1143 | 1013.6 | 1464.5 | 1607 | 1730.4 | 1605.8 | 1207 | 1647.7 | -0.4495 | 0.001951 | 0.04111 | *Pold1* | 7 |
| ENSMUSG00000036185 | 119.65 | 91.962 | 134.69 | 82.646 | 46.838 | 50.773 | 115.44 | 60.085 | 0.9453 | 0.001963 | 0.041301 | *Sapcd1* | 17 |
| ENSMUSG00000047879 | 1594.3 | 1562.4 | 1731.8 | 1940.6 | 2336.7 | 2063.8 | 1629.5 | 2113.7 | -0.3758 | 0.00197 | 0.041415 | *Usp14* | 18 |
| ENSMUSG00000035529 | 573.08 | 632.85 | 560.15 | 811.15 | 969.71 | 683.94 | 588.7 | 821.6 | -0.4815 | 0.001985 | 0.041637 | *Prdm4* | 10 |
| ENSMUSG00000009894 | 1254.3 | 1035.3 | 1062.6 | 1458 | 1713 | 1340 | 1117.4 | 1503.7 | -0.4289 | 0.001985 | 0.041637 | *Snap47* | 11 |
| ENSMUSG00000029703 | 716.88 | 689.22 | 693.78 | 881.55 | 1039.1 | 873.1 | 699.96 | 931.25 | -0.4126 | 0.00199 | 0.041692 | *Lrwd1* | 5 |
| ENSMUSG00000026213 | 708.48 | 659.55 | 665.98 | 905.02 | 1036.5 | 804.4 | 678.01 | 915.31 | -0.4337 | 0.002006 | 0.041971 | *Stk11ip* | 1 |
| ENSMUSG00000027082 | 1572.3 | 1464.5 | 1453.8 | 1213.2 | 1147.5 | 1142.9 | 1496.9 | 1167.9 | 0.3582 | 0.002021 | 0.042196 | *Tfpi* | 2 |
| ENSMUSG00000006289 | 860.68 | 745.58 | 953.54 | 1204 | 1090.3 | 1124 | 853.27 | 1139.4 | -0.4174 | 0.002021 | 0.042196 | *Osgep* | 14 |
| ENSMUSG00000026615 | 3540.3 | 3242.4 | 3111.8 | 4209.8 | 5062.8 | 3780.1 | 3298.2 | 4350.9 | -0.3998 | 0.002024 | 0.042209 | *Eprs* | 1 |
| ENSMUSG00000021149 | 1455.8 | 1778.9 | 1581 | 1885.5 | 3132.9 | 1969.2 | 1605.3 | 2329.2 | -0.5374 | 0.002054 | 0.04276 | *Gtpbp4* | 13 |
| ENSMUSG00000079845 | 18.893 | 33.62 | 25.656 | 47.955 | 84.134 | 53.76 | 26.056 | 61.95 | -1.2507 | 0.002056 | 0.04276 | *Xlr4a* | X |
| ENSMUSG00000005683 | 1857.8 | 1904.5 | 2290.9 | 2617.1 | 2668.9 | 2549.6 | 2017.7 | 2611.9 | -0.3725 | 0.002067 | 0.042901 | *Cs* | 10 |
| ENSMUSG00000017692 | 213.07 | 268.96 | 315.35 | 188.76 | 155.26 | 169.24 | 265.8 | 171.09 | 0.6367 | 0.002067 | 0.042901 | *Rhbdl3* | 11 |

| | | | | | | | | | | | | | |
|---|---|---|---|---|---|---|---|---|---|---|---|---|---|
| ENSMUSG00000039313 | 290.74 | 317.42 | 246.94 | 203.04 | 117.96 | 206.08 | 285.03 | 175.69 | 0.7018 | 0.002074 | 0.043 | *AF529169* | 9 |
| ENSMUSG00000052516 | 607.72 | 568.58 | 638.19 | 448.94 | 359.96 | 487.82 | 604.83 | 432.24 | 0.4859 | 0.00209 | 0.043282 | *Robo2* | 16 |
| ENSMUSG00000068522 | 12067 | 16682 | 14304 | 20419 | 17114 | 19551 | 14351 | 19028 | -0.4069 | 0.002095 | 0.043333 | *Aard* | 15 |
| ENSMUSG00000066415 | 1154.6 | 1258.8 | 1287.1 | 965.22 | 904.66 | 983.6 | 1233.5 | 951.16 | 0.3753 | 0.002098 | 0.043337 | *Msl2* | 9 |
| ENSMUSG00000000399 | 1192.4 | 1133.2 | 1248.6 | 1393.8 | 1857.9 | 1524.2 | 1191.4 | 1591.9 | -0.4188 | 0.002106 | 0.043455 | *Ndufa9* | 6 |
| ENSMUSG00000004100 | 741.02 | 754.48 | 899.02 | 1015.2 | 1263.8 | 989.58 | 798.18 | 1089.5 | -0.4498 | 0.002111 | 0.043483 | *Ppan* | 9 |
| ENSMUSG00000040345 | 70.323 | 73.174 | 76.968 | 35.711 | 37.297 | 36.835 | 73.488 | 36.614 | 1.0045 | 0.002112 | 0.043483 | *Arhgap9* | 10 |
| ENSMUSG00000053070 | 178.43 | 221.5 | 161.42 | 237.73 | 366.9 | 280.74 | 187.12 | 295.12 | -0.6582 | 0.002119 | 0.043575 | *Cfap300* | 9 |
| ENSMUSG00000037697 | 598.27 | 690.21 | 525.94 | 931.55 | 791.04 | 777.52 | 604.81 | 833.37 | -0.4616 | 0.002126 | 0.043663 | *Ddhd1* | 14 |
| ENSMUSG00000026134 | 726.33 | 603.19 | 636.05 | 825.44 | 976.65 | 848.21 | 655.19 | 883.43 | -0.432 | 0.002137 | 0.043818 | *Prim2* | 1 |
| ENSMUSG00000018861 | 4010.5 | 4230.2 | 4730.3 | 3293.6 | 2228.3 | 3693.5 | 4323.7 | 3071.8 | 0.4935 | 0.002138 | 0.043818 | *Fdxr* | 11 |
| ENSMUSG00000048915 | 608.77 | 514.19 | 694.85 | 425.47 | 345.21 | 486.82 | 605.94 | 419.17 | 0.5324 | 0.002144 | 0.043887 | *Efna5* | 17 |
| ENSMUSG00000008129 | 53.53 | 69.218 | 66.278 | 101.01 | 156.99 | 95.573 | 63.009 | 117.86 | -0.9061 | 0.002153 | 0.043977 | *4930432K21Rik* | 8 |
| ENSMUSG00000028837 | 1698.3 | 1481.3 | 1993.7 | 2083.5 | 2458.1 | 2363.4 | 1724.4 | 2301.7 | -0.417 | 0.002153 | 0.043977 | *Psmb2* | 4 |
| ENSMUSG00000085925 | 112.31 | 124.59 | 131.49 | 60.199 | 73.726 | 77.653 | 122.8 | 70.526 | 0.7987 | 0.002165 | 0.044145 | *Rtl1* | 12 |
| ENSMUSG00000021367 | 361.06 | 366.86 | 291.84 | 424.45 | 572.46 | 474.88 | 339.92 | 490.6 | -0.53 | 0.002166 | 0.044145 | *Edn1* | 13 |
| ENSMUSG00000038774 | 584.63 | 629.89 | 471.43 | 736.67 | 913.33 | 715.8 | 561.98 | 788.6 | -0.489 | 0.002172 | 0.044218 | *Ascc3* | 10 |
| ENSMUSG00000030619 | 704.28 | 751.51 | 818.85 | 963.18 | 1340.1 | 901.97 | 758.22 | 1068.4 | -0.4957 | 0.002181 | 0.044319 | *Eed* | 7 |
| ENSMUSG00000024966 | 2076.1 | 2130.9 | 2501.4 | 2846.7 | 2996.7 | 2776.6 | 2236.2 | 2873.3 | -0.3619 | 0.002182 | 0.044319 | *Stip1* | 19 |
| ENSMUSG00000029524 | 353.72 | 352.03 | 327.11 | 227.53 | 216.84 | 270.79 | 344.28 | 238.39 | 0.5314 | 0.002187 | 0.044377 | *Sirt4* | 5 |
| ENSMUSG00000027932 | 2552.6 | 2778.6 | 2839.2 | 2080.4 | 1544.8 | 2346.5 | 2723.5 | 1990.6 | 0.4527 | 0.00219 | 0.044377 | *Slc27a3* | 3 |
| ENSMUSG00000032078 | 630.81 | 625.93 | 799.61 | 862.17 | 1137.1 | 877.08 | 685.45 | 958.79 | -0.4852 | 0.002194 | 0.044421 | *Zpr1* | 9 |
| ENSMUSG00000063972 | 131.2 | 228.42 | 152.87 | 263.24 | 406.79 | 216.03 | 170.83 | 295.36 | -0.7904 | 0.002198 | 0.044441 | *Nr6a1* | 2 |
| ENSMUSG00000000197 | 135.4 | 152.28 | 161.42 | 226.51 | 233.32 | 221.01 | 149.7 | 226.95 | -0.6006 | 0.002202 | 0.044477 | *Nalcn* | 14 |
| ENSMUSG00000052504 | 1006.6 | 728.77 | 909.71 | 634.64 | 670.47 | 621.22 | 881.69 | 642.11 | 0.4568 | 0.002208 | 0.044507 | *Epha3* | 16 |
| ENSMUSG00000063663 | 569.94 | 681.31 | 404.08 | 905.02 | 823.13 | 707.84 | 551.77 | 811.99 | -0.5567 | 0.002209 | 0.044507 | *Brwd3* | X |
| ENSMUSG00000004356 | 1256.4 | 1354.7 | 1218.7 | 1865.1 | 2306.3 | 1319.1 | 1276.6 | 1830.2 | -0.52 | 0.002235 | 0.044999 | *Utp20* | 10 |
| ENSMUSG00000043924 | 23.091 | 49.442 | 23.518 | 74.483 | 94.543 | 53.76 | 32.017 | 74.262 | -1.2113 | 0.002263 | 0.04545 | *Ncmap* | 4 |
| ENSMUSG00000083714 | 2.0992 | 7.9107 | 2.138 | 10.203 | 35.562 | 20.907 | 4.0493 | 22.224 | -2.4479 | 0.002263 | 0.04545 | *Fthl17-ps1* | X |
| ENSMUSG00000032594 | 2429.8 | 2695.6 | 2475.8 | 2018.2 | 1955 | 2052.8 | 2533.7 | 2008.7 | 0.3352 | 0.002273 | 0.045611 | *Ip6k1* | 9 |
| ENSMUSG00000036678 | 555.24 | 524.08 | 529.15 | 641.78 | 837.87 | 714.8 | 536.16 | 731.49 | -0.4492 | 0.00228 | 0.045695 | *Aaas* | 15 |
| ENSMUSG00000066652 | 23.091 | 14.833 | 24.587 | 3.0609 | 6.9389 | 2.9866 | 20.837 | 4.3288 | 2.2457 | 0.002283 | 0.045698 | *Lefty2* | 1 |
| ENSMUSG00000030788 | 690.64 | 698.12 | 597.57 | 811.15 | 1038.2 | 854.18 | 662.11 | 901.19 | -0.4453 | 0.0023 | 0.045958 | *Rnf141* | 7 |
| ENSMUSG00000020282 | 1560.8 | 1753.2 | 1861.1 | 1363.1 | 1069.5 | 1420.6 | 1725 | 1284.4 | 0.4261 | 0.002304 | 0.045958 | *Rhbdf1* | 11 |
| ENSMUSG00000086598 | 38.835 | 82.073 | 41.691 | 78.564 | 204.7 | 86.613 | 54.2 | 123.29 | -1.1865 | 0.002304 | 0.045958 | *Btbd18* | 2 |
| ENSMUSG00000041720 | 728.43 | 663.51 | 628.57 | 892.78 | 948.9 | 836.26 | 673.5 | 892.64 | -0.4067 | 0.002306 | 0.045958 | *Pi4ka* | 16 |
| ENSMUSG00000024993 | 529 | 527.05 | 537.7 | 681.57 | 740.73 | 689.92 | 531.25 | 704.07 | -0.4068 | 0.00231 | 0.045989 | *Fam45a* | 19 |
| ENSMUSG00000095567 | 1109.4 | 1333.9 | 1460.2 | 1718.2 | 2010.6 | 1556 | 1301.2 | 1761.6 | -0.4374 | 0.002315 | 0.04604 | *Noc2l* | 4 |

| | | | | | | | | | | | | | |
|---|---|---|---|---|---|---|---|---|---|---|---|---|---|
| ENSMUSG00000050812 | 2596.7 | 2822.1 | 2402 | 3463 | 3531 | 3043.4 | 2607 | 3345.8 | -0.36 | 0.002321 | 0.046085 | *Ecpas* | 4 |
| ENSMUSG00000032816 | 2287.1 | 2266.4 | 2096.3 | 1808 | 1354 | 1823.8 | 2216.6 | 1661.9 | 0.4161 | 0.002324 | 0.046085 | *Igdcc4* | 9 |
| ENSMUSG00000026646 | 393.6 | 366.86 | 434.01 | 439.76 | 757.21 | 584.39 | 398.16 | 593.78 | -0.5782 | 0.002327 | 0.046085 | *Suv39h2* | 2 |
| ENSMUSG00000041794 | 17.843 | 31.643 | 16.035 | 51.016 | 58.981 | 49.777 | 21.84 | 53.258 | -1.2822 | 0.002328 | 0.046085 | *Myrip* | 9 |
| ENSMUSG00000027905 | 818.69 | 910.72 | 880.85 | 1076.4 | 1355.7 | 1066.2 | 870.09 | 1166.1 | -0.4231 | 0.002334 | 0.046116 | *Ddx20* | 3 |
| ENSMUSG00000045294 | 1073.7 | 1042.2 | 1097.9 | 1525.4 | 1944.6 | 1123 | 1071.3 | 1531 | -0.5157 | 0.002337 | 0.046116 | *Insig1* | 5 |
| ENSMUSG00000010080 | 251.91 | 229.41 | 279.01 | 344.87 | 362.56 | 376.32 | 253.44 | 361.25 | -0.5121 | 0.002337 | 0.046116 | *Epn3* | 11 |
| ENSMUSG00000046714 | 82.919 | 104.82 | 116.52 | 164.27 | 175.21 | 157.3 | 101.42 | 165.59 | -0.708 | 0.002357 | 0.046387 | *Foxc2* | 8 |
| ENSMUSG00000063626 | 3.1488 | 6.9218 | 3.207 | 23.467 | 26.021 | 13.938 | 4.4259 | 21.142 | -2.2493 | 0.002358 | 0.046387 | *Unc5d* | 8 |
| ENSMUSG00000087516 | 44.083 | 26.699 | 52.381 | 19.386 | 14.745 | 7.9644 | 41.054 | 14.032 | 1.5473 | 0.002358 | 0.046387 | *Tbx3os1* | 5 |
| ENSMUSG00000021732 | 38.835 | 48.453 | 37.415 | 15.305 | 14.745 | 17.92 | 41.568 | 15.99 | 1.381 | 0.002373 | 0.046635 | *Fgf10* | 13 |
| ENSMUSG00000020075 | 1902.9 | 1738.4 | 1779.9 | 2401.8 | 2923 | 1995.1 | 1807.1 | 2440 | -0.4336 | 0.00238 | 0.046704 | *Ddx21* | 10 |
| ENSMUSG00000030609 | 436.64 | 384.66 | 403.01 | 518.32 | 693.02 | 517.69 | 408.1 | 576.34 | -0.4994 | 0.002398 | 0.046982 | *Aen* | 7 |
| ENSMUSG00000039137 | 688.54 | 690.21 | 594.36 | 494.85 | 438.02 | 518.68 | 657.7 | 483.85 | 0.4439 | 0.002399 | 0.046982 | *Whrn* | 4 |
| ENSMUSG00000024530 | 100.76 | 102.84 | 96.209 | 161.21 | 206.43 | 132.41 | 99.937 | 166.68 | -0.74 | 0.002405 | 0.047034 | *Prelid3a* | 18 |
| ENSMUSG00000029446 | 377.86 | 356.97 | 337.8 | 469.35 | 667 | 435.06 | 357.54 | 523.8 | -0.5523 | 0.002407 | 0.047034 | *Psph* | 5 |
| ENSMUSG00000004562 | 1695.1 | 1797.7 | 2030 | 1366.2 | 1256.8 | 1568 | 1840.9 | 1397 | 0.3984 | 0.002414 | 0.047123 | *Arhgef40* | 14 |
| ENSMUSG00000042426 | 642.36 | 594.29 | 699.12 | 893.8 | 911.6 | 780.51 | 645.26 | 861.97 | -0.4184 | 0.002418 | 0.047125 | *Dhx29* | 13 |
| ENSMUSG00000075028 | 751.52 | 782.17 | 692.71 | 633.62 | 459.7 | 518.68 | 742.13 | 537.33 | 0.4672 | 0.002422 | 0.047125 | *Prdm11* | 2 |
| ENSMUSG00000004892 | 3365 | 4198.6 | 2817.9 | 2846.7 | 2047.8 | 2549.6 | 3460.5 | 2481.4 | 0.4802 | 0.002425 | 0.047125 | *Bcan* | 3 |
| ENSMUSG00000035064 | 1093.7 | 1072.9 | 1013.4 | 729.53 | 831.8 | 863.14 | 1060 | 808.16 | 0.3911 | 0.002426 | 0.047125 | *Eef2k* | 7 |
| ENSMUSG00000030527 | 1915.5 | 2369.2 | 2079.2 | 1667.2 | 1393 | 1747.2 | 2121.3 | 1602.5 | 0.4052 | 0.00243 | 0.047125 | *Crtc3* | 7 |
| ENSMUSG00000022324 | 2999.8 | 2531.4 | 2665 | 1808 | 2133.7 | 2306.7 | 2732.1 | 2082.8 | 0.3913 | 0.002432 | 0.047125 | *Matn2* | 15 |
| ENSMUSG00000023348 | 1645.8 | 1648.4 | 1944.5 | 1371.3 | 1251.6 | 1408.7 | 1746.2 | 1343.9 | 0.378 | 0.002432 | 0.047125 | *Trip6* | 5 |
| ENSMUSG00000036825 | 466.02 | 552.76 | 595.43 | 662.19 | 921.14 | 699.87 | 538.07 | 761.07 | -0.5012 | 0.002439 | 0.047207 | *Ssx2ip* | 3 |
| ENSMUSG00000018537 | 644.46 | 838.53 | 899.02 | 604.03 | 499.6 | 599.32 | 794 | 567.65 | 0.4849 | 0.002447 | 0.047286 | *Pcgf2* | 11 |
| ENSMUSG00000074594 | 8.3968 | 6.9218 | 14.966 | 37.752 | 39.031 | 21.902 | 10.095 | 32.895 | -1.7109 | 0.002449 | 0.047286 | *Wfdc9* | 2 |
| ENSMUSG00000104960 | 544.74 | 465.74 | 695.91 | 388.74 | 353.02 | 437.05 | 568.8 | 392.93 | 0.5337 | 0.002451 | 0.047286 | *Snhg8* | 3 |
| ENSMUSG00000022479 | 124.9 | 209.63 | 136.83 | 243.86 | 241.13 | 269.79 | 157.12 | 251.59 | -0.6771 | 0.00246 | 0.04735 | *Vdr* | 15 |
| ENSMUSG00000038046 | 163.74 | 132.5 | 151.8 | 211.21 | 287.97 | 204.09 | 149.35 | 234.42 | -0.6531 | 0.002462 | 0.04735 | *Mrm3* | 11 |
| ENSMUSG00000048701 | 684.34 | 846.44 | 815.64 | 1024.4 | 1371.3 | 913.91 | 782.14 | 1103.2 | -0.4968 | 0.002462 | 0.04735 | *Ccdc6* | 10 |
| ENSMUSG00000018848 | 1393.9 | 1503 | 1599.2 | 1783.5 | 2399.1 | 1839.8 | 1498.7 | 2007.5 | -0.4222 | 0.00249 | 0.047781 | *Rars* | 11 |
| ENSMUSG00000031539 | 116.51 | 131.52 | 100.49 | 174.47 | 223.78 | 165.26 | 116.17 | 187.84 | -0.694 | 0.002492 | 0.047781 | *Ap3m2* | 8 |
| ENSMUSG00000031320 | 24103 | 25263 | 27477 | 19346 | 22222 | 19540 | 25614 | 20369 | 0.3305 | 0.002493 | 0.047781 | *Rps4x* | X |
| ENSMUSG00000040118 | 1672 | 1797.7 | 1669.8 | 1435.6 | 1112.8 | 1362.9 | 1713.2 | 1303.8 | 0.3947 | 0.002514 | 0.048133 | *Cacna2d1* | 5 |
| ENSMUSG00000097303 | 12.595 | 21.754 | 18.173 | 34.691 | 78.93 | 32.853 | 17.508 | 48.825 | -1.4831 | 0.002532 | 0.04844 | *3110083C13Rik* | 14 |
| ENSMUSG00000031935 | 1209.1 | 968.07 | 959.96 | 1309.1 | 1626.3 | 1308.2 | 1045.7 | 1414.5 | -0.4364 | 0.002543 | 0.048603 | *Med17* | 9 |
| ENSMUSG00000006333 | 19432 | 18090 | 21583 | 13935 | 13878 | 17602 | 19702 | 15139 | 0.3801 | 0.002569 | 0.048951 | *Rps9* | 7 |

| | | | | | | | | | | | | | |
|---|---|---|---|---|---|---|---|---|---|---|---|---|---|
| ENSMUSG00000038860 | 442.93 | 525.07 | 353.84 | 688.71 | 640.98 | 558.5 | 440.61 | 629.4 | -0.5137 | 0.002569 | 0.048951 | *Garnl3* | 2 |
| ENSMUSG00000035726 | 2529.5 | 2587.8 | 2173.3 | 3095.6 | 3864.1 | 2792.5 | 2430.2 | 3250.8 | -0.4199 | 0.00257 | 0.048951 | *Supt16* | 14 |
| ENSMUSG00000030878 | 262.4 | 248.2 | 313.22 | 327.52 | 558.58 | 381.3 | 274.6 | 422.47 | -0.6237 | 0.002573 | 0.048952 | *Cdr2* | 7 |
| ENSMUSG00000106223 | 4.1984 | 4.9442 | 4.276 | 15.305 | 26.021 | 19.911 | 4.4729 | 20.412 | -2.1919 | 0.002578 | 0.049 | *2400006E01Rik* | 3 |
| ENSMUSG00000027359 | 8.3968 | 14.833 | 7.483 | 21.427 | 53.777 | 27.875 | 10.237 | 34.36 | -1.7469 | 0.002591 | 0.049164 | *Slc27a2* | 2 |
| ENSMUSG00000030538 | 376.81 | 329.28 | 432.94 | 437.72 | 610.62 | 596.33 | 379.68 | 548.22 | -0.5314 | 0.002597 | 0.049164 | *Cib1* | 7 |
| ENSMUSG00000028789 | 531.1 | 572.54 | 598.64 | 453.02 | 382.51 | 415.14 | 567.42 | 416.89 | 0.4457 | 0.002598 | 0.049164 | *Azin2* | 4 |
| ENSMUSG00000076430 | 3.1488 | 9.8884 | 3.207 | 16.325 | 49.44 | 12.942 | 5.4147 | 26.236 | -2.2727 | 0.002599 | 0.049164 | *Hus1b* | 13 |
| ENSMUSG00000078652 | 1498.8 | 1724.5 | 1777.7 | 1866.2 | 2591.7 | 2248 | 1667 | 2235.3 | -0.4235 | 0.0026 | 0.049164 | *Psme3* | 11 |
| ENSMUSG00000078350 | 1848.4 | 1530.7 | 1566.1 | 1272.3 | 1186.6 | 1338 | 1648.4 | 1265.6 | 0.3814 | 0.002606 | 0.049235 | *Smim1* | 4 |
| ENSMUSG00000020733 | 262.4 | 300.61 | 408.36 | 435.67 | 563.79 | 446.01 | 323.79 | 481.82 | -0.5749 | 0.002612 | 0.049283 | *Slc9a3r1* | 11 |
| ENSMUSG00000058248 | 505.91 | 390.59 | 462.87 | 248.96 | 157.86 | 400.21 | 453.12 | 269.01 | 0.754 | 0.002618 | 0.049345 | *Kcnh1* | 1 |
| ENSMUSG00000021178 | 2751 | 2267.4 | 2766.6 | 3204.8 | 3819 | 3154.9 | 2595 | 3392.9 | -0.3871 | 0.002624 | 0.049403 | *Psmc1* | 12 |
| ENSMUSG00000087428 | 0 | 4.9442 | 2.138 | 9.1828 | 33.827 | 10.951 | 2.3607 | 17.987 | -2.9205 | 0.002629 | 0.04944 | *Gm15017* | X |
| ENSMUSG00000092390 | 2.0992 | 8.8995 | 1.069 | 22.447 | 20.817 | 19.911 | 4.0226 | 21.058 | -2.3691 | 0.002631 | 0.04944 | *Gm20541* | 17 |
| ENSMUSG00000024990 | 29.389 | 65.263 | 52.381 | 75.503 | 124.9 | 99.555 | 49.011 | 99.986 | -1.0289 | 0.002638 | 0.04944 | *Rbp4* | 19 |
| ENSMUSG00000031226 | 641.31 | 650.65 | 784.64 | 820.33 | 1243.8 | 889.03 | 692.2 | 984.39 | -0.5091 | 0.002645 | 0.04944 | *Pbdc1* | X |
| ENSMUSG00000027270 | 112.31 | 29.665 | 48.105 | 23.467 | 31.225 | 9.9555 | 63.359 | 21.549 | 1.5487 | 0.002645 | 0.04944 | *Lamp5* | 2 |
| ENSMUSG00000086748 | 0 | 9.8884 | 4.276 | 19.386 | 45.103 | 12.942 | 4.7214 | 25.81 | -2.4433 | 0.002646 | 0.04944 | *Gm13261* | 2 |
| ENSMUSG00000059810 | 1497.8 | 1606.9 | 1415.3 | 1213.2 | 1072.1 | 1216.6 | 1506.7 | 1167.3 | 0.3688 | 0.002647 | 0.04944 | *Rgs3* | 4 |
| ENSMUSG00000028693 | 4250.9 | 4269.8 | 5005 | 5442.4 | 7243.4 | 5340.1 | 4508.6 | 6008.6 | -0.4146 | 0.00265 | 0.04944 | *Nasp* | 4 |
| ENSMUSG00000055093 | 301.24 | 188.87 | 323.9 | 167.33 | 111.89 | 202.1 | 271.34 | 160.44 | 0.7597 | 0.00265 | 0.04944 | *Gm8430* | 6 |
| ENSMUSG00000072295 | 5.248 | 19.777 | 3.207 | 30.609 | 52.909 | 23.893 | 9.4106 | 35.804 | -1.9191 | 0.002671 | 0.049776 | *C2cd6* | 1 |
| ENSMUSG00000030556 | 147.99 | 185.9 | 191.35 | 242.84 | 361.69 | 227.98 | 175.08 | 277.5 | -0.6665 | 0.002677 | 0.049829 | *Lrrc28* | 7 |
| ENSMUSG00000084184 | 17.843 | 9.8884 | 12.828 | 3.0609 | 0.8674 | 0.9955 | 13.52 | 1.6413 | 3.0657 | 0.00268 | 0.049836 | *Gm13474* | 2 |
| ENSMUSG00000026567 | 14.694 | 18.788 | 8.5519 | 27.549 | 91.073 | 21.902 | 14.011 | 46.841 | -1.7438 | 0.002686 | 0.0499 | *Adcy10* | 1 |
