## Supplemental Table 3 for "Splicing factor SRSF1 is essential for homing of precursor spermatogonial stem cells in mice"

**Table S3. AS events were analysed in cKO and Ctrl testes.**

| GeneID | geneSymbol | chr | type | IncFormLe | SkipFormLe | PValue | FDR | IncLevel1 | IncLevel2 | IncLevelDifferenc |
|---|---|---|---|---|---|---|---|---|---|---|
| ENSMUSG00000045672 | *Col27a1* | 4 | A3SS | 152 | 149 | 1.80E-05 | 0.033264 | 0.893,0.728,0.648 | 1.0,1.0,1.0 | -0.244 |
| ENSMUSG00000064179 | *Tnnt1* | 7 | A3SS | 152 | 149 | 2.79E-08 | 0.000103 | 0.329,0.662,0.927 | 0.0,0.0,0.0 | 0.639 |
| ENSMUSG00000014767 | *Tbp* | 17 | A5SS | 304 | 149 | 5.48E-09 | 5.94E-06 | 0.0,0.076,0.0 | 0.318,0.51,0.62 | -0.457 |
| ENSMUSG00000051331 | *Cacna1c* | 6 | A5SS | 323 | 149 | 2.59E-14 | 5.61E-11 | 1.0,1.0,1.0 | 0.187,0.0,0.0 | 0.938 |
| ENSMUSG00000060985 | *Tdrd5* | 1 | A5SS | 380 | 149 | 2.42E-05 | 0.013133 | 0.541,0.0,0.44 | 0.836,0.897,1.0 | -0.584 |
| ENSMUSG00000073125 | *Xlr3b* | X | A5SS | 164 | 149 | 1.18E-05 | 0.008499 | 1.0,1.0,1.0 | 0.0,0.0,1.0 | 0.667 |
| ENSMUSG00000028478 | *Clta* | 4 | MXE | 203 | 185 | 2.85E-05 | 0.00966 | 1.0,0.477,1.0 | 0.0,0.313,0.0 | 0.721 |
| ENSMUSG00000051331 | *Cacna1c* | 6 | MXE | 233 | 182 | 2.03E-06 | 0.001234 | 1.0,1.0,1.0 | 0.0,0.0,1.0 | 0.667 |
| ENSMUSG00000007279 | *Scube2* | 7 | MXE | 233 | 254 | 8.86E-12 | 1.35E-08 | 1.0,1.0,1.0 | 0.154,0.466,0.353 | 0.676 |
| ENSMUSG00000033720 | *Sfxn5* | 6 | MXE | 240 | 265 | 0.000118 | 0.032722 | 0.0,0.0,0.0 | 0.424,0.408,0.321 | -0.384 |
| ENSMUSG00000014353 | *Tmem87b* | 2 | MXE | 221 | 221 | 0.000158 | 0.036935 | 0.518,0.549,0.63 | 0.357,0.301,0.333 | 0.235 |
| ENSMUSG00000048402 | *Gli2* | 1 | MXE | 316 | 335 | 1.06E-05 | 0.004602 | 0.096,0.0,0.0 | 0.597,0.414,0.298 | -0.404 |
| ENSMUSG00000041670 | *Rims1* | 1 | MXE | 230 | 218 | 0 | 0 | 0.0,0.0,0.0 | 1.0,0.895,1.0 | -0.965 |
| ENSMUSG00000028256 | *Odf2l* | 3 | MXE | 227 | 308 | 1.30E-07 | 0.000132 | 0.0,0.0,0.0 | 1.0,0.352,1.0 | -0.784 |
| ENSMUSG00000022938 | *Fam3b* | 16 | MXE | 273 | 208 | 3.07E-06 | 0.001559 | 1.0,1.0,0.873 | 0.087,0.588,0.696 | 0.501 |
| ENSMUSG00000068758 | *Il3ra* | 14 | MXE | 268 | 179 | 0.000149 | 0.036935 | 0.958,1.0,0.923 | 0.829,0.715,0.736 | 0.2 |
| ENSMUSG00000008658 | *Rbfox1* | 16 | MXE | 189 | 214 | 0.000103 | 0.031259 | 0.0,0.773,0.0 | 1.0,0.85,1.0 | -0.692 |
| ENSMUSG00000030846 | *Tial1* | 7 | MXE | 218 | 217 | 1.90E-05 | 0.007224 | 0.033,0.142,0.025 | 0.326,0.241,0.27 | -0.212 |
| ENSMUSG00000039997 | *Ifi203* | 1 | MXE | 293 | 1391 | 2.08E-07 | 0.000158 | 0.133,0.0,0.0 | 0.783,0.177,0.564 | -0.464 |
| ENSMUSG00000019320 | *Noxo1* | 17 | RI | 389 | 149 | 1.68E-05 | 0.008599 | 1.0,1.0,0.365 | 0.0,0.354,0.296 | 0.572 |
| ENSMUSG00000044551 | *9930012K11Rik* | 14 | RI | 288 | 149 | 6.94E-06 | 0.007559 | 0.376,0.368,0.376 | 0.129,0.045,0.0 | 0.315 |
| ENSMUSG00000041679 | *Lrrc29* | 8 | RI | 307 | 149 | 9.12E-06 | 0.007559 | 0.421,0.108,0.66 | 0.708,1.0,1.0 | -0.506 |
| ENSMUSG00000027804 | *Ppid* | 3 | RI | 2525 | 149 | 1.21E-05 | 0.007559 | 0.329,0.352,0.257 | 0.163,0.204,0.152 | 0.14 |
| ENSMUSG00000034928 | *Rnf44* | 13 | RI | 336 | 149 | 2.11E-05 | 0.009269 | 0.578,1.0,1.0 | 0.372,0.331,0.307 | 0.523 |
| ENSMUSG00000026754 | *Golga1* | 2 | RI | 600 | 149 | 2.99E-05 | 0.011494 | 1.0,0.897,1.0 | 0.546,0.695,0.851 | 0.268 |
| ENSMUSG00000029032 | *Arhgef16* | 4 | RI | 1028 | 149 | 6.87E-05 | 0.023472 | 1.0,1.0,1.0 | 0.0,0.367,1.0 | 0.544 |
| ENSMUSG00000078485 | *Plekhn1* | 4 | RI | 2693 | 149 | 1.23E-05 | 0.007559 | 1.0,1.0,1.0 | 0.529,0.583,0.81 | 0.359 |
| ENSMUSG00000020059 | *Sycp3* | 10 | RI | 273 | 149 | 0.000108 | 0.033103 | 0.766,0.577,0.947 | 0.443,0.263,0.392 | 0.397 |
| ENSMUSG00000070000 | *Fcho1* | 8 | RI | 388 | 149 | 7.28E-06 | 0.007559 | 0.248,0.133,0.227 | 0.578,0.633,0.569 | -0.391 |
| ENSMUSG00000024985 | *Tcf7l2* | 19 | SE | 200 | 149 | 1.55E-05 | 0.00597 | 0.598,0.516,0.701 | 1.0,1.0,0.882 | -0.356 |
| ENSMUSG00000024743 | *Syt7* | 19 | SE | 377 | 149 | 0 | 0 | 1.0,1.0,1.0 | 0.165,0.0,0.145 | 0.897 |
| ENSMUSG00000060450 | *Rnf14* | 18 | SE | 225 | 149 | 1.48E-07 | 0.000173 | 0.0,0.0,0.0 | 1.0,1.0,0.498 | -0.833 |
| ENSMUSG00000024044 | *Epb41l3* | 17 | SE | 224 | 149 | 7.87E-09 | 1.45E-05 | 1.0,0.842,1.0 | 0.25,0.195,0.0 | 0.799 |
| ENSMUSG00000025735 | *Rhbdl1* | 17 | SE | 323 | 149 | 8.71E-05 | 0.023629 | 0.639,0.356,0.504 | 1.0,0.822,0.866 | -0.396 |
| ENSMUSG00000025735 | *Rhbdl1* | 17 | SE | 311 | 149 | 9.61E-05 | 0.025139 | 0.736,0.564,0.678 | 1.0,0.888,0.913 | -0.274 |
| ENSMUSG00000014767 | *Tbp* | 17 | SE | 353 | 149 | 0.000225 | 0.049929 | 0.0,0.013,0.0 | 0.11,0.13,0.099 | -0.109 |

| | | | | | | | | | | |
|---|---|---|---|---|---|---|---|---|---|---|
| ENSMUSG00000014767 | *Tbp* | 17 | SE | 195 | 149 | 1.20E-06 | 0.001006 | 0.113,0.0,0.0 | 0.256,0.344,0.505 | -0.331 |
| ENSMUSG00000022791 | *Tnk2* | 16 | SE | 245 | 149 | 1.74E-08 | 2.71E-05 | 0.93,0.86,0.792 | 1.0,1.0,0.976 | -0.131 |
| ENSMUSG00000022556 | *Hsf1* | 15 | SE | 286 | 149 | 4.04E-05 | 0.013137 | 0.973,0.978,0.972 | 0.83,0.803,0.845 | 0.148 |
| ENSMUSG00000022378 | *Fam49b* | 15 | SE | 232 | 149 | 5.93E-08 | 7.97E-05 | 0.0,0.097,0.0 | 0.342,0.264,0.252 | -0.254 |
| ENSMUSG00000022378 | *Fam49b* | 15 | SE | 232 | 149 | 8.98E-05 | 0.024136 | 0.005,0.029,0.0 | 0.093,0.115,0.106 | -0.093 |
| ENSMUSG00000024381 | *Bin1* | 18 | SE | 278 | 149 | 4.89E-06 | 0.002678 | 0.115,0.235,0.162 | 0.339,0.479,0.42 | -0.242 |
| ENSMUSG00000022197 | *Pdzd2* | 15 | SE | 305 | 149 | 3.16E-06 | 0.001867 | 1.0,1.0,1.0 | 0.661,0.0,0.268 | 0.69 |
| ENSMUSG00000041481 | *Serpina3g* | 12 | SE | 241 | 149 | 9.47E-14 | 4.67E-10 | 0.0,0.0,0.481 | 1.0,1.0,1.0 | -0.84 |
| ENSMUSG00000054894 | *Atp5s* | 12 | SE | 249 | 149 | 9.44E-05 | 0.02493 | 0.467,0.295,0.318 | 0.725,0.7,0.749 | -0.365 |
| ENSMUSG00000022607 | *Ptk2* | 15 | SE | 170 | 149 | 1.16E-06 | 0.001006 | 0.012,0.02,0.043 | 0.117,0.236,0.178 | -0.152 |
| ENSMUSG00000079179 | *Rab10os* | 12 | SE | 325 | 149 | 5.32E-10 | 1.35E-06 | 0.957,1.0,1.0 | 0.468,0.777,0.612 | 0.367 |
| ENSMUSG00000033102 | *Cdc14b* | 13 | SE | 215 | 149 | 0.00015 | 0.036595 | 0.055,0.131,0.148 | 0.409,0.4,0.291 | -0.255 |
| ENSMUSG00000021458 | *2010111I01Rik* | 13 | SE | 338 | 149 | 3.98E-09 | 8.40E-06 | 1.0,1.0,1.0 | 0.469,0.128,0.306 | 0.699 |
| ENSMUSG00000020814 | *Mxra7* | 11 | SE | 237 | 149 | 2.70E-05 | 0.009387 | 0.812,0.711,0.891 | 0.322,0.575,0.416 | 0.367 |
| ENSMUSG00000001036 | *Epn2* | 11 | SE | 284 | 149 | 1.51E-06 | 0.001127 | 1.0,0.512,1.0 | 0.0,0.208,0.208 | 0.699 |
| ENSMUSG00000020389 | *Cdkl3* | 11 | SE | 223 | 149 | 7.60E-05 | 0.021194 | 1.0,0.501,1.0 | 0.539,0.334,0.401 | 0.409 |
| ENSMUSG00000020287 | *Mpg* | 11 | SE | 519 | 149 | 2.31E-07 | 0.000253 | 0.28,0.541,0.203 | 0.801,0.884,1.0 | -0.554 |
| ENSMUSG00000049800 | *Sertad2* | 11 | SE | 234 | 149 | 3.80E-14 | 2.24E-10 | 0.0,0.0,0.0 | 1.0,1.0,1.0 | -1 |
| ENSMUSG00000038708 | *Golga4* | 9 | SE | 248 | 149 | 4.63E-06 | 0.002583 | 0.3,0.462,0.26 | 0.029,0.059,0.082 | 0.284 |
| ENSMUSG00000033419 | *Snap91* | 9 | SE | 233 | 149 | 1.44E-09 | 3.28E-06 | 0.561,0.242,0.299 | 1.0,1.0,1.0 | -0.633 |
| ENSMUSG00000032076 | *Cadm1* | 9 | SE | 182 | 149 | 0.000112 | 0.028798 | 0.165,0.296,0.347 | 0.466,0.588,0.504 | -0.25 |
| ENSMUSG00000032386 | *Trip4* | 9 | SE | 301 | 149 | 1.37E-06 | 0.001065 | 1.0,1.0,0.271 | 0.0,0.331,0.0 | 0.647 |
| ENSMUSG00000032034 | *Kcnj5* | 9 | SE | 452 | 149 | 1.82E-05 | 0.006734 | 1.0,0.569,1.0 | 0.248,0.0,0.622 | 0.566 |
| ENSMUSG00000021843 | *Ktn1* | 14 | SE | 218 | 149 | 0.000193 | 0.044622 | 0.457,0.478,0.426 | 0.724,0.595,0.694 | -0.217 |
| ENSMUSG00000039599 | *Fam149b* | 14 | SE | 266 | 149 | 2.56E-12 | 1.08E-08 | 0.0,0.123,0.0 | 1.0,1.0,1.0 | -0.959 |
| ENSMUSG00000021012 | *Zc3h14* | 12 | SE | 309 | 149 | 9.89E-06 | 0.004363 | 0.493,0.466,0.423 | 0.633,0.666,0.63 | -0.182 |
| ENSMUSG00000003824 | *Syce2* | 8 | SE | 303 | 149 | 1.17E-05 | 0.004873 | 0.337,0.409,0.442 | 0.555,0.688,0.585 | -0.213 |
| ENSMUSG00000028634 | *Hivep3* | 4 | SE | 262 | 149 | 0.00019 | 0.044268 | 0.416,0.245,0.666 | 0.603,1.0,1.0 | -0.425 |
| ENSMUSG00000067367 | *Lyar* | 5 | SE | 273 | 149 | 1.86E-05 | 0.006807 | 0.163,0.0,0.0 | 0.302,0.308,0.148 | -0.198 |
| ENSMUSG00000066113 | *Adamtsl1* | 4 | SE | 200 | 149 | 5.10E-06 | 0.002743 | 0.0,0.0,0.0 | 0.598,1.0,1.0 | -0.866 |
| ENSMUSG00000008575 | *Nfib* | 4 | SE | 238 | 149 | 2.42E-05 | 0.008632 | 0.065,0.0,0.0 | 0.556,0.594,0.135 | -0.407 |
| ENSMUSG00000054752 | *Fsd1l* | 4 | SE | 172 | 149 | 2.00E-06 | 0.001317 | 0.0,0.634,0.0 | 1.0,1.0,1.0 | -0.789 |
| ENSMUSG00000053279 | *Aldh1a1* | 19 | SE | 347 | 149 | 1.29E-05 | 0.00514 | 1.0,1.0,1.0 | 0.417,0.256,0.75 | 0.526 |
| ENSMUSG00000069755 | *Zfp125* | 12 | SE | 232 | 149 | 0 | 0 | 0.0,0.097,0.0 | 1.0,1.0,1.0 | -0.968 |
| ENSMUSG00000029328 | *Hnrnpdl* | 5 | SE | 320 | 149 | 1.48E-05 | 0.005833 | 0.9,0.917,0.889 | 0.988,0.99,0.977 | -0.083 |
| ENSMUSG00000068917 | *Clk2* | 3 | SE | 237 | 149 | 1.28E-05 | 0.00514 | 0.672,0.667,0.642 | 0.839,0.886,0.918 | -0.221 |
| ENSMUSG00000068917 | *Clk2* | 3 | SE | 234 | 149 | 1.53E-06 | 0.001127 | 0.564,0.601,0.619 | 0.802,0.869,0.906 | -0.264 |
| ENSMUSG00000035401 | *Emsy* | 7 | SE | 485 | 149 | 0.000103 | 0.026739 | 0.819,1.0,1.0 | 0.673,0.818,0.646 | 0.227 |

| | | | | | | | | | | |
|---|---|---|---|---|---|---|---|---|---|---|
| ENSMUSG00000021280 | *Exoc3l4* | 12 | SE | 254 | 149 | 5.51E-07 | 0.000543 | 0.128,0.089,0.0 | 1.0,0.518,1.0 | -0.767 |
| ENSMUSG00000021357 | *Exoc2* | 13 | SE | 208 | 149 | 5.00E-08 | 7.17E-05 | 0.0,0.014,0.0 | 0.155,0.217,0.237 | -0.198 |
| ENSMUSG00000040928 | *S100pbp* | 4 | SE | 266 | 149 | 9.33E-05 | 0.024838 | 0.766,0.728,0.883 | 1.0,0.956,1.0 | -0.193 |
| ENSMUSG00000030629 | *Zfand6* | 7 | SE | 310 | 149 | 0.000196 | 0.044875 | 0.852,1.0,1.0 | 0.325,0.325,0.684 | 0.506 |
| ENSMUSG00000030314 | *Atg7* | 6 | SE | 270 | 149 | 3.75E-05 | 0.012313 | 0.0,0.0,0.0 | 0.0,1.0,1.0 | -0.667 |
| ENSMUSG00000031458 | *Coprs* | 8 | SE | 365 | 149 | 0.000131 | 0.032446 | 0.62,0.569,0.786 | 0.936,0.891,0.862 | -0.238 |
| ENSMUSG00000041199 | *Rpusd1* | 17 | SE | 338 | 149 | 1.96E-06 | 0.001317 | 1.0,1.0,1.0 | 0.897,0.767,0.806 | 0.177 |
| ENSMUSG00000004947 | *Dtx2* | 5 | SE | 287 | 149 | 0.000217 | 0.048629 | 0.63,0.72,0.801 | 0.433,0.345,0.558 | 0.272 |
| ENSMUSG00000078515 | *Ddi2* | 4 | SE | 211 | 149 | 1.03E-05 | 0.004396 | 1.0,1.0,0.973 | 0.393,0.713,0.874 | 0.331 |
| ENSMUSG00000078515 | *Ddi2* | 4 | SE | 211 | 149 | 6.19E-05 | 0.018124 | 0.962,1.0,1.0 | 0.0,0.463,0.87 | 0.543 |
| ENSMUSG00000078515 | *Ddi2* | 4 | SE | 339 | 149 | 8.24E-05 | 0.022776 | 0.966,1.0,1.0 | 0.796,0.694,0.951 | 0.175 |
| ENSMUSG00000026672 | *Optn* | 2 | SE | 352 | 149 | 3.53E-05 | 0.011724 | 1.0,1.0,1.0 | 0.714,0.894,0.691 | 0.234 |
| ENSMUSG00000026672 | *Optn* | 2 | SE | 326 | 149 | 5.48E-05 | 0.016535 | 1.0,1.0,1.0 | 0.794,0.809,0.814 | 0.194 |
| ENSMUSG00000024266 | *Adad2* | 8 | SE | 290 | 149 | 2.02E-05 | 0.007281 | 1.0,1.0,1.0 | 0.698,0.629,0.814 | 0.286 |
| ENSMUSG00000026723 | *Trdmt1* | 2 | SE | 259 | 149 | 1.76E-05 | 0.006582 | 0.955,1.0,1.0 | 0.817,0.788,0.771 | 0.193 |
| ENSMUSG00000032966 | *Fkbp1a* | 2 | SE | 240 | 149 | 3.43E-05 | 0.011537 | 0.037,0.094,0.0 | 0.276,0.261,0.271 | -0.226 |
| ENSMUSG00000006315 | *Tmem147* | 7 | SE | 219 | 149 | 4.55E-05 | 0.014321 | 0.731,0.689,0.362 | 0.731,1.0,1.0 | -0.316 |
| ENSMUSG00000051985 | *Igfn1* | 1 | SE | 4634 | 149 | 6.76E-05 | 0.019228 | 1.0,1.0,1.0 | 0.088,0.114,1.0 | 0.599 |
| ENSMUSG00000028483 | *Snapc3* | 4 | SE | 234 | 149 | 6.61E-05 | 0.018972 | 0.813,0.74,0.925 | 1.0,0.954,0.988 | -0.155 |
| ENSMUSG00000053333 | *Dis3l2* | 1 | SE | 191 | 149 | 1.22E-06 | 0.001006 | 0.084,0.0,0.0 | 0.245,0.324,0.247 | -0.244 |
| ENSMUSG00000099913 | *Gm28551* | 1 | SE | 346 | 149 | 6.81E-06 | 0.003345 | 0.365,0.564,0.528 | 1.0,1.0,1.0 | -0.514 |
| ENSMUSG00000026219 | *Trip12* | 1 | SE | 275 | 149 | 0.000174 | 0.041576 | 0.234,0.142,0.0 | 0.359,0.306,0.368 | -0.219 |
| ENSMUSG00000024566 | *Atp9b* | 18 | SE | 226 | 149 | 5.09E-08 | 7.17E-05 | 0.928,1.0,0.896 | 0.755,0.654,0.446 | 0.323 |
| ENSMUSG00000087177 | *E130307A14Rik* | 10 | SE | 216 | 149 | 1.08E-07 | 0.000139 | 1.0,1.0,0.805 | 0.315,0.133,0.58 | 0.592 |
| ENSMUSG00000087177 | *E130307A14Rik* | 10 | SE | 352 | 149 | 3.90E-06 | 0.002261 | 1.0,1.0,0.899 | 0.76,0.669,0.798 | 0.224 |
| ENSMUSG00000029769 | *Ccdc136* | 6 | SE | 335 | 149 | 4.33E-05 | 0.013761 | 0.0,0.727,0.527 | 1.0,1.0,0.83 | -0.525 |
| ENSMUSG00000029769 | *Ccdc136* | 6 | SE | 431 | 149 | 2.39E-11 | 7.07E-08 | 0.195,0.28,0.257 | 1.0,0.818,1.0 | -0.695 |
| ENSMUSG00000029769 | *Ccdc136* | 6 | SE | 428 | 149 | 8.36E-12 | 3.09E-08 | 1.0,1.0,1.0 | 0.723,0.474,0.701 | 0.367 |
| ENSMUSG00000051285 | *Pcmtd1* | 1 | SE | 341 | 149 | 4.19E-05 | 0.013458 | 0.561,0.476,0.466 | 0.341,0.132,0.256 | 0.258 |
| ENSMUSG00000022995 | *Enah* | 1 | SE | 212 | 149 | 1.56E-06 | 0.001127 | 0.014,0.05,0.021 | 0.097,0.271,0.162 | -0.148 |
| ENSMUSG00000028919 | *Arhgef19* | 4 | SE | 298 | 149 | 1.00E-11 | 3.28E-08 | 1.0,0.765,1.0 | 0.077,0.152,0.36 | 0.725 |
| ENSMUSG00000028919 | *Arhgef19* | 4 | SE | 256 | 149 | 8.71E-05 | 0.023629 | 1.0,0.368,1.0 | 0.185,0.143,0.068 | 0.657 |
| ENSMUSG00000049606 | *Zfp644* | 5 | SE | 712 | 149 | 1.78E-15 | 1.75E-11 | 1.0,1.0,1.0 | 0.334,0.255,0.611 | 0.6 |
| ENSMUSG00000049606 | *Zfp644* | 5 | SE | 206 | 149 | 0.000199 | 0.045168 | 0.685,1.0,1.0 | 1.0,0.083,0.126 | 0.492 |
| ENSMUSG00000049606 | *Zfp644* | 5 | SE | 206 | 149 | 3.33E-15 | 2.46E-11 | 1.0,1.0,1.0 | 0.126,0.062,0.52 | 0.764 |
| ENSMUSG00000052373 | *Mpp3* | 11 | SE | 170 | 149 | 1.36E-08 | 2.23E-05 | 0.549,0.513,0.672 | 0.141,0.137,0.0 | 0.485 |
| ENSMUSG00000040728 | *Esrp1* | 4 | SE | 259 | 149 | 0.000136 | 0.033487 | 0.0,0.071,0.0 | 0.333,0.165,0.087 | -0.171 |
| ENSMUSG00000023044 | *Csad* | 15 | SE | 239 | 149 | 6.43E-05 | 0.018629 | 0.903,0.652,0.804 | 1.0,0.961,1.0 | -0.201 |

| | | | | | | | | | | |
|---|---|---|---|---|---|---|---|---|---|---|
| ENSMUSG00000023044 | *Csad* | 15 | SE | 201 | 149 | 1.54E-05 | 0.00597 | 0.863,0.608,0.797 | 1.0,0.942,1.0 | -0.225 |
| ENSMUSG00000028693 | *Nasp* | 4 | SE | 1124 | 149 | 6.06E-09 | 1.19E-05 | 0.359,0.396,0.381 | 0.538,0.59,0.57 | -0.187 |
| ENSMUSG00000041757 | *Plekha6* | 1 | SE | 295 | 149 | 2.64E-06 | 0.001663 | 0.202,0.0,0.0 | 0.735,0.414,1.0 | -0.649 |
| ENSMUSG00000021947 | *Cryl1* | 14 | SE | 254 | 149 | 6.89E-06 | 0.003345 | 1.0,1.0,0.37 | 0.0,0.0,0.37 | 0.667 |
| ENSMUSG00000039262 | *Prrc2b* | 2 | SE | 231 | 149 | 5.90E-05 | 0.017443 | 0.222,0.443,0.241 | 0.537,0.576,0.726 | -0.311 |
| ENSMUSG00000058402 | *Zfp420* | 7 | SE | 202 | 149 | 6.90E-06 | 0.003345 | 0.0,0.0,0.425 | 0.633,1.0,0.596 | -0.601 |
| ENSMUSG00000030041 | *M1ap* | 6 | SE | 290 | 149 | 6.68E-07 | 0.000637 | 1.0,1.0,1.0 | 0.855,0.637,0.782 | 0.242 |
| ENSMUSG00000031389 | *Arhgap4* | X | SE | 212 | 149 | 2.01E-06 | 0.001317 | 0.484,0.0,0.26 | 1.0,1.0,0.738 | -0.665 |
| ENSMUSG00000032582 | *Rbm6* | 9 | SE | 219 | 149 | 1.21E-05 | 0.00496 | 1.0,0.576,0.405 | 0.095,0.036,0.026 | 0.608 |
| ENSMUSG00000074899 | *Sptbn5* | 2 | SE | 232 | 149 | 1.52E-07 | 0.000173 | 0.0,0.391,0.0 | 0.72,1.0,0.562 | -0.63 |
| ENSMUSG00000004043 | *Stat5a* | 11 | SE | 201 | 149 | 5.12E-05 | 0.015777 | 0.763,0.426,0.553 | 1.0,0.952,0.87 | -0.36 |
| ENSMUSG00000043962 | *Thrap3* | 4 | SE | 845 | 149 | 5.47E-10 | 1.35E-06 | 0.994,0.954,0.98 | 0.735,0.693,0.893 | 0.202 |
| ENSMUSG00000068758 | *Il3ra* | 14 | SE | 179 | 149 | 9.35E-06 | 0.004186 | 0.143,0.0,0.454 | 0.4,1.0,1.0 | -0.601 |
| ENSMUSG00000013973 | *Dedd* | 1 | SE | 182 | 149 | 1.01E-05 | 0.004382 | 0.539,0.353,0.399 | 1.0,1.0,0.724 | -0.478 |
| ENSMUSG00000014592 | *Camta1* | 4 | SE | 170 | 149 | 2.45E-06 | 0.001576 | 0.0,0.0,0.305 | 1.0,0.637,0.305 | -0.546 |
| ENSMUSG00000019790 | *Stxbp5* | 10 | SE | 197 | 149 | 0.000156 | 0.037913 | 1.0,0.532,1.0 | 0.602,0.201,0.0 | 0.576 |
| ENSMUSG00000034164 | *Emid1* | 11 | SE | 240 | 149 | 0.000123 | 0.031014 | 0.134,0.383,0.0 | 0.262,0.383,0.347 | -0.158 |
| ENSMUSG00000037605 | *Adgrl3* | 5 | SE | 353 | 149 | 2.86E-06 | 0.001727 | 0.655,0.53,0.747 | 1.0,1.0,1.0 | -0.356 |
| ENSMUSG00000006262 | *Mob1b* | 5 | SE | 258 | 149 | 0.000163 | 0.0391 | 0.634,0.464,0.722 | 1.0,1.0,1.0 | -0.393 |
| ENSMUSG00000029298 | *Gbp9* | 5 | SE | 252 | 149 | 2.91E-05 | 0.010018 | 0.639,0.542,0.825 | 0.0,1.0,0.0 | 0.335 |
| ENSMUSG00000033209 | *Ttc28* | 5 | SE | 173 | 149 | 1.23E-06 | 0.001006 | 0.223,1.0,1.0 | 0.0,0.0,0.0 | 0.741 |
| ENSMUSG00000028573 | *Fggy* | 4 | SE | 211 | 149 | 5.95E-06 | 0.003142 | 1.0,1.0,0.931 | 0.712,0.647,0.647 | 0.308 |
| ENSMUSG00000057337 | *Chst3* | 10 | SE | 302 | 149 | 0.000214 | 0.048214 | 1.0,1.0,1.0 | 0.152,1.0,0.33 | 0.506 |
| ENSMUSG00000036555 | *Iqce* | 5 | SE | 284 | 149 | 7.46E-06 | 0.003501 | 0.677,0.655,0.84 | 1.0,1.0,1.0 | -0.276 |
| ENSMUSG00000066640 | *Fbxl18* | 5 | SE | 1693 | 149 | 5.31E-05 | 0.016171 | 0.937,1.0,0.893 | 0.656,0.655,0.837 | 0.227 |
| ENSMUSG00000007812 | *Zfp655* | 5 | SE | 235 | 149 | 1.09E-05 | 0.004598 | 1.0,1.0,1.0 | 0.727,0.769,0.538 | 0.322 |
| ENSMUSG00000029823 | *Luc7l2* | 6 | SE | 250 | 149 | 0.000183 | 0.043247 | 1.0,1.0,1.0 | 0.749,0.641,0.573 | 0.346 |
| ENSMUSG00000029823 | *Luc7l2* | 6 | SE | 217 | 149 | 8.69E-09 | 1.51E-05 | 0.089,0.407,0.654 | 1.0,1.0,1.0 | -0.617 |
| ENSMUSG00000036879 | *Phkb* | 8 | SE | 258 | 149 | 1.32E-06 | 0.001053 | 1.0,0.882,1.0 | 0.743,0.536,0.548 | 0.352 |
| ENSMUSG00000036879 | *Phkb* | 8 | SE | 241 | 149 | 1.66E-06 | 0.001167 | 1.0,0.867,1.0 | 0.724,0.382,0.587 | 0.391 |
| ENSMUSG00000039678 | *Tbc1d13* | 2 | SE | 293 | 149 | 9.02E-06 | 0.004102 | 1.0,1.0,0.869 | 0.484,0.604,0.921 | 0.287 |
| ENSMUSG00000022487 | *Gtsf1* | 15 | SE | 230 | 149 | 0.000122 | 0.030986 | 0.463,1.0,1.0 | 0.257,0.162,0.404 | 0.547 |
| ENSMUSG00000031864 | *Ints10* | 8 | SE | 227 | 149 | 5.73E-05 | 0.0171 | 1.0,0.868,1.0 | 0.663,0.766,0.441 | 0.333 |
| ENSMUSG00000032254 | *Kif23* | 9 | SE | 461 | 149 | 4.18E-06 | 0.002378 | 0.481,0.543,0.506 | 0.738,0.671,0.66 | -0.18 |
| ENSMUSG00000008140 | *Emc10* | 7 | SE | 236 | 149 | 0.000124 | 0.031014 | 0.557,0.503,0.508 | 0.374,0.301,0.389 | 0.168 |
| ENSMUSG00000040189 | *Ccdc114* | 7 | SE | 228 | 149 | 3.35E-05 | 0.011388 | 1.0,1.0,0.907 | 0.753,0.805,0.649 | 0.233 |
| ENSMUSG00000040189 | *Ccdc114* | 7 | SE | 224 | 149 | 8.97E-07 | 0.000828 | 1.0,1.0,0.823 | 0.624,0.608,0.1 | 0.497 |
| ENSMUSG00000075555 | *Gm10863* | 15 | SE | 208 | 149 | 1.31E-07 | 0.000162 | 0.571,0.648,0.626 | 0.917,1.0,1.0 | -0.357 |

| | | | | | | | | | | |
|---|---|---|---|---|---|---|---|---|---|---|
| ENSMUSG00000029267 | *Mtf2* | 5 | SE | 262 | 149 | 4.80E-07 | 0.000503 | 1.0,1.0,1.0 | 0.363,0.645,0.561 | 0.477 |
| ENSMUSG00000029267 | *Mtf2* | 5 | SE | 297 | 149 | 8.76E-06 | 0.004047 | 1.0,1.0,1.0 | 0.501,0.707,0.681 | 0.37 |
| ENSMUSG00000062797 | *Hikeshi* | 7 | SE | 387 | 149 | 6.87E-06 | 0.003345 | 0.688,0.682,0.623 | 0.786,0.843,0.883 | -0.173 |
| ENSMUSG00000006310 | *Zbtb32* | 7 | SE | 218 | 149 | 0.000188 | 0.044192 | 0.406,0.0,0.0 | 0.421,0.381,0.732 | -0.376 |
| ENSMUSG00000030849 | *Fgfr2* | 7 | SE | 416 | 149 | 7.25E-05 | 0.020409 | 0.642,0.668,0.617 | 0.0,0.207,0.417 | 0.434 |
| ENSMUSG00000038695 | *Josd2* | 7 | SE | 261 | 149 | 2.59E-05 | 0.009127 | 0.624,0.826,0.66 | 0.268,0.363,0.311 | 0.389 |
| ENSMUSG00000030846 | *Tial1* | 7 | SE | 218 | 149 | 7.25E-06 | 0.003456 | 0.054,0.339,0.05 | 0.506,0.785,0.533 | -0.46 |
| ENSMUSG00000031808 | *Slc27a1* | 8 | SE | 256 | 149 | 4.94E-05 | 0.01537 | 0.088,0.085,0.026 | 0.431,0.256,0.327 | -0.272 |
| ENSMUSG00000052446 | *Zfp961* | 8 | SE | 272 | 149 | 2.76E-06 | 0.0017 | 1.0,0.751,1.0 | 0.512,0.354,0.558 | 0.442 |
| ENSMUSG00000052446 | *Zfp961* | 8 | SE | 276 | 149 | 4.93E-07 | 0.000503 | 1.0,0.829,1.0 | 0.612,0.674,0.654 | 0.296 |
| ENSMUSG00000052446 | *Zfp961* | 8 | SE | 346 | 149 | 6.25E-06 | 0.003241 | 1.0,0.66,1.0 | 0.418,0.334,0.498 | 0.47 |
| ENSMUSG00000020027 | *Socs2* | 10 | SE | 253 | 149 | 1.71E-05 | 0.006465 | 0.078,0.0,0.0 | 0.32,0.371,0.841 | -0.485 |
