## Supplemental Table 4 for "Splicing factor SRSF1 is essential for homing of precursor spermatogonial stem cells in mice"

**Table S4. The SRSF1 interacting proteins were displayed through IP-MS.**

| Protein | ScoreRep1 | ScoreRep2 | Protein | ScoreRep1 | ScoreRep2 | Protein | ScoreRep1 | ScoreRep2 |
|---|---|---|---|---|---|---|---|---|
| SRSF1 | 2882 | 2396 | GM10709 | 21 | 21 | MME | 22 | 21 |
| IGKV4-91 | 158 | 150 | DAZAP1 | 21 | 41 | GEMIN4 | 51 | 121 |
| IGHV1-43 | 327 | 347 | IDH2 | 35 | 29 | DHX38 | 40 | 18 |
| IGKV19-93 | 31 | 31 | XAB2 | 36 | 36 | F830045P16RIK | 36 | 35 |
| SRRM2 | 59 | 53 | SPATS2L | 26 | 23 | KIF2B | 25 | 16 |
| L3MBTL4 | 20 | 20 | COL12A1 | 14 | 22 | CCAR1 | 34 | 106 |
| PCDHGA7 | 16 | 15 | ZC3H13 | 28 | 28 | LRRC45 | 15 | 16 |
| CYTH1 | 20 | 20 | ACIN1 | 144 | 171 | SLC25A1 | 16 | 15 |
| PTPRF | 26 | 26 | GM9493 | 119 | 68 | TWF1 | 24 | 24 |
| DOCK7 | 25 | 25 | GIGYF2 | 16 | 17 | WDR7 | 20 | 29 |
| GEMIN5 | 291 | 361 | GM9825 | 31 | 30 | 44987 | 17 | 17 |
| MED12 | 15 | 27 | SNRPA1 | 25 | 18 | PRPF3 | 39 | 54 |
| AQR | 15 | 24 | PAXBP1 | 21 | 17 | DDX50 | 36 | 31 |
| PPIG | 99 | 65 | SF3B6 | 43 | 55 | COX6C | 17 | 15 |
| CWC22 | 31 | 21 | CDC42 | 21 | 16 | GEMIN2 | 83 | 29 |
| NRK | 44 | 28 | SNRPE | 46 | 34 | PRPF4 | 43 | 52 |
| TBX1 | 24 | 20 | ARF5 | 55 | 78 | CDC40 | 20 | 37 |
| SF3A2 | 23 | 18 | RBM15 | 102 | 71 | CRIP2 | 32 | 41 |
| HADHB | 19 | 21 | SRSF10 | 204 | 256 | TFIP11 | 56 | 50 |
| SAP18 | 38 | 84 | USP39 | 18 | 15 | DDX20 | 54 | 67 |
| RPL5 | 243 | 145 | LUC7L3 | 101 | 84 | GOLGA5 | 14 | 15 |
| DDX23 | 32 | 30 | PDLIM4 | 14 | 22 | SART1 | 85 | 91 |
| GYS1 | 16 | 16 | | | | | | |
