## Supplemental Table 5 for "Splicing factor SRSF1 is essential for homing of precursor spermatogonial stem cells in mice"

**Table S5. Primer sequences were used in this study.**

| Target | Sequence(5' to 3') | Application |
|---|---|---|
| *Srsf1* -GT-F | ACTAATGTGGGAAGAATGGC | Genotyping |
| *Srsf1* -GT-FKO | TTCGCCTTCGTTGAGTTC | |
| *Srsf1* -GT-R | AAACTATTGCTCCCATCTGC | |
| *Vasa* -Cre-MutF | CACGTGCAGCCGTTTAAGCCGCGT | |
| *Vasa* -Cre-MutR | TTCCCATTCTAAACAACACCCTGAA | |
| *Vasa-Cre-* WtF | CTAGGCCACAGAATTGAAAGATCT | |
| *Vasa* -Cre-WtR | GTAGGTGGAAATTCTAGCATCATCC | |
| *Srsf1* -F | AGGAGGATTGAGGAGGATCAG | RT-qPCR & RIP-qPCR |
| *Srsf1* -R | CGCTCCATGAATCCTGGTAA | |
| *Gapdh* -F | GGGTCCCAGCTTAGGTTCAT | |
| *Gapdh* -R | CCCAATACGGCCAAATCCGT | |
| *Gfra1* -F | AGCCACTCTGTACTTCGTGC | |
| *Gfra1* -R | TCAGTGTGCGGTACTTGGTG | |
| *Pou5f1* -F | GAGAACCGTGTGAGGTGGAG | |
| *Pou5f1* -R | GATCTTTTGCCCTTCTGGCG | |
| *Plzf* -F | GGAGACGCACTACAGGGTTC | |
| *Plzf* -R | GGTACACTGGTATGGCGAGG | |
| *Nanos3* -F | GTCTACTGCTACACCACCCG | |
| *Nanos3* -R | CTGCCACTTTTGGAACCTGC | |
| *Dnd1* -F | CTTAAACCGCCGAGCGTTAC | |
| *Dnd1* -R | GAGGCGTGACTGACCTTCTA | |
| *Stra8* -F | ACCGTGGTGGCCTTAAAGATTA | |
| *Stra8* -R | TGAAGAGCCCTACCAGGGTG | |
| *Taf4b* -F | CGACAATCCAACTACCGGCT | |
| *Taf4b* -R | CTTGTGGTCTTGGCTCCTGT | |
| *Sall4* -F | AACTTCTCGTCTGCCAGTGC | |
| *Sall4* -R | GCGGGCGGAGTTATTGTTG | |
| *Tial1* -F | TCGGGGCTGACAGATCAACTT | |
| *Tial1* -R | GTTCACAGAGACAATGGCGTG | |
| *Tial1* -seF | TTGTTGGGGATTTGAGTCCAGA | RT-PCR |
| *Tial1* -seR | TCTATCATCATTTTGGAGTTTGGGC | |
| SRSF1-OEF | GTGGCTCGAGCAGGCGCGCCATGTCGGGAGGTGGTGTGAT | Plasmid construction |
| SRSF1-OER | GGGCCCTCTAGATTAATTAATTATGTACGAGAGCGAGATC | |
| SRSF1-RRM1F | GCCCGGCGGGGAACAACGACGGCCGCGGGACCGGCCGAGG | |
| SRSF1-RRM1R | CCTCGGCCGGTCCCGCGGCCGTCGTTGTTCCCCGCCGGGC | |
| SRSF1-RRM2F | CGCCGTCCAGGCGGTCCGAGGGGCCCAGAAGTCCAAGTTA | |
| SRSF1-RRM2R | TAACTTGGACTTCTGGGCCCCTCGGACCGCCTGGACGGCG | |
| SRSF1-RSR | GGGCCCTCTAGATTAATTAATTATGTGGGCCCATCAACTTTAACCC | |
| SRSF10-F | GTGGCTCGAGCAGGCGCGCCATGTCCCGATACCTGCGCCC | |
| SRSF10-R | GGGCCCTCTAGATTAATTAATCAGATCTTTCTTGAAGTGT | |
| SART1-F | GTGGCTCGAGCAGGCGCGCCATGGGGTCGTCAAAGAAGCA | |
| SART1-R | GGGCCCTCTAGATTAATTAATCATTTGGTGATGGTGTTCG | |
| RBM15-F | GTGGCTCGAGCAGGCGCGCCATGAGGTCTGCGGGGCGGGA | |
| RBM15-R | GGGCCCTCTAGATTAATTAATCATCCGCTGTTCACCAGTT | |
