## Supplemental Table 6 for "Splicing factor SRSF1 is essential for homing of precursor spermatogonial stem cells in mice"

**Table S6. Antibodies were used in this study.**

| Name | Cat.NO. | Source | Company | Dilution |
|---|---|---|---|---|
| DDX4 | ab13840 | Rabbit polyclonal | abcam | IF, 1: 800 |
| DDX4 | ab27591 | Mouse monoclonal | abcam | IF, 1: 800 |
| ACTB | 81115-1-RR | Rabbit monoclonal | Proteintech | WB, 1: 20000 |
| SRSF1 | 12929-2-AP | Rabbit polyclonal | Proteintech | WB, 1:1000 IF, 1:200 |
| SRSF1 | sc-33652 | Mouse monoclonal | Santa Cruz | IF, 1:500 |
| γH2AX (AlexaFluor® 555) | 05-636-AF555 | Mouse monoclonal | Millipore | IF, 1:800 |
| HA | M20003 | Mouse monoclonal | Abmart | WB, 1: 5000 |
| FLAG | bsm-33346M | Mouse monoclonal | Bioss | WB, 1: 5000 |
| PLZF | sc-28319 | Mouse monoclonal | Santa Cruz | IF, 1: 200 |
| SOX9 (AlexaFluor® 555) | AB5535-AF555 | Rabbit polyclonal | Millipore | IF, 1: 100 |
| TIAL1/TIAR | 66907-1-Ig | Mouse monoclonal | Proteintech | IF, 1: 200 |
| SRSF10 | bs-13229R | Rabbit polyclonal | Bioss | WB, 1: 1000 |
| SART1 | ab181867 | Rabbit monoclonal | abcam | WB, 1: 20000 |
| RBM15 | bs-12402R | Rabbit polyclonal | Bioss | WB, 1: 1000 |
| TRA98 | ab82527 | Rat monoclonal | abcam | IF, 1:400 |
| GAR-488 | A11034 | Goat polyclonal | Life Technologies | IF, 1: 500 |
| GAM-594 | A32742 | Goat polyclonal | Life Technologies | IF, 1: 500 |
| DAR-594 | A-21207 | Donkey polyclonal | Life Technologies | IF, 1: 500 |
| DAM-488 | A-21202 | Donkey polyclonal | Life Technologies | IF, 1: 500 |
| GAR-HRP | 7074P2 | Goat polyclonal | CST | WB, 1: 2000 |
| GAM-HRP | 7076P2 | Goat polyclonal | CST | WB, 1: 2000 |
